# Higher functional resilience of temperate forests at intermediate latitudes of a large latitudinal gradient in South America

**DOI:** 10.1101/2024.05.31.596814

**Authors:** Xiongjie Deng, Danny E. Carvajal, Rocío Urrutia-Jalabert, Waira S. Machida, Alice Rosen, Huanyuan Zhang-Zheng, David Galbraith, Sandra Díaz, Yadvinder Malhi, Jesús Aguirre-Gutiérrez

## Abstract

Accurately mapping and assessing plant functional composition across space and time is pivotal for understanding environmental change impacts on the biodiversity and functioning of forests. Here, we test the capabilities of a combination of *in-situ* and remote sensing approaches to deliver accurate estimates of the functional composition of temperate forest ecosystems considering leaf and stem morphological, nutrient, hydraulic, and photosynthetic traits. We identify hydrological stress, soil, and topography as key drivers of plant functional traits. Further, hydrological stress and soil are key determinants of functional dispersion and redundancy in temperate forests distributed across a large latitudinal (30°S to 53°S) gradient in Chile. Functional dispersion peaks across Mediterranean forests, woodlands, and scrub, occupying between 30°S to 35°S. Conversely, functional redundancy peaks between 42°S and 53°S, corresponding to Magellanic subpolar forests. Although functional dispersion and redundancy peak at different latitudes corresponding to distinct forest types; they are both high at latitudes between 35°S and 42°S, coinciding with Valdivian temperate rainforests. Our results highlight areas in temperate forests in South America where both tree functional dispersion and redundancy are high, and hence could potentially be more resilient to environmental changes.

## Main

Temperate forests show remarkable diversity in terms of species, soil composition, and the carbon dynamics within their ecosystems^1–3^. In particular, due to their geographic isolation, temperate forests in Chile are highly endemic at species and higher^4,5^, and thus have become a priority for conservation^6^. These temperate forests also provide essential contributions to people^7^, protecting watersheds, absorbing and storing large amounts of carbon dioxide^8–11^, accumulating carbon in the soil, which further contributes to long-term carbon sequestration^12–14^, and influencing temperature, humidity, and precipitation patterns^15^. However, these forests are under persistent threat from replacement by other land uses such as agriculture and exotic plantations, degradation^6^, unsustainable logging, warming, and precipitation decrease^16–20^. As a result, the interactions that underpin ecosystem functions and benefits to people have become progressively fragile^21,22^. Therefore, a comprehensive understanding of biodiversity dynamics, drivers of biodiversity, and the potential impact of environmental changes on biodiversity^23,24^ in such temperate forests becomes imperative for better anticipating the consequences of climate change and biodiversity loss on ecosystem functioning and derived societal benefits^25–27^.

Functional diversity is a vital component of biodiversity that captures the range and distribution of plant functional traits across spatial scales^28^ and is a fundamental component of ecosystem processes^29^, ecosystem resilience to environmental changes^30^, and provision of nature’s benefits to people^31,32^. Functional diversity metrics like functional dispersion quantifies the extent of trait dissimilarity within plant communities^33^, elucidating how divergent or convergent species are in terms of their functional traits. There is a vast literature on in what circumstances and how taxonomic or functional richness contribute to ecosystem functioning^25,34^, but there is a widely held opinion that the larger the variety, the higher the chances that at least some species would be able to thrive under changing conditions^29^. At the same time, functional redundancy is the degree to which multiple species within an ecosystem show similar functional attributes and therefore are assumed to perform similar ecological roles or functions, so that, when one species disappears, some other very similar ones would still be able to play its role^35–37^. It is thought to be important for ecosystem stability and resilience in the face of disturbances, including extreme climatic events^13,38,39^. Ecosystems with low functional redundancy may be more vulnerable to environmental changes because the roles of lost species are less likely to be compensated by remaining functionally similar species^37^. Hence, assessing an ecosystem’s functional dispersion and redundancy is particularly relevant for understanding how biodiversity contributes to ecosystem functioning and benefits to people^40,41^, for tracking responses of species to changing environmental conditions and disturbance regimes^42,43^, and for predicting ecosystem productivity and stability^44^.

Previous studies exploring relationships between biodiversity and the environment predominantly relied on taxonomic methods to assess the impact of biodiversity on ecosystem functioning^45,46^. Unlike taxonomic methods, trait-based approaches focus on plant characteristics, which represent their morphological, physiological, biochemical or phenological characteristics that are considered relevant to the response of such plants to the environment and/or their impacts on ecosystem properties^47,48^. By analysing functional traits we can better understand how plants adapt to different environmental conditions^49,50^ and shed light on the role of plants in ecosystem processes, such as nutrient cycling^51^, biomass production^52^, and carbon sequestration^53^.

Trait-based approaches that measure the characteristics of individuals have been valuable in describing and predicting ecosystem functions at the individual-to-plot scales. However, their upscaling to whole ecosystems and beyond has proven challenging^54^. Still, recent advances in remote sensing techniques have allowed upscaling of some functional traits from local to global scales and assessment of functional diversity continuously in space and time^44,55–57^. Despite the capabilities of remote sensing approaches to upscale and predict some functional traits, there is a lack of understanding about mapping and predicting variability in plant functional trait distributions^58^ and the distribution of plant functional diversity and redundancy at large spatial scales^59^, particularly in temperate forests beyond North America and Western Europe^60^.

Here, we use leaf and stem trait data measured *in*-*situ* from 8,104 individual trees across 16 permanent forest plots in Chile (Fig.1a, b, Extended Data Table 1), including leaf and stem morphology, leaf nutrients, and hydraulic and photosynthetic traits (Fig.1c). The plots are distributed across a large latitudinal gradient extending from Mediterranean climate ∼30°S to cold temperate climate at ∼53°S, covering a large annual rainfall (450 mm-4,500 mm) and mean annual temperature gradient (5.7°C-13°C). Our main objectives are to 1) test how community-level tree functional traits and functional dispersion and redundancy vary across the latitudinal gradient, 2) identify and quantify the main environmental drivers of distributions of functional traits and functional dispersion and redundancy, and 3) test if and to what extent functional trait distributions, dispersion and redundancy can be mapped using *in-situ* and remotely sensed data (methods illustrated in Fig.1c). Specifically for the third aim, we evaluate the potential of integrating *in-situ* measurements, high-resolution airborne and satellite-based (Sentinel-2) multispectral reflectance images and LiDAR laser scanning, together with environmental variables to assess plant functional traits and infer functional dispersion and redundancy at the ecosystem level.

**Fig.1.**
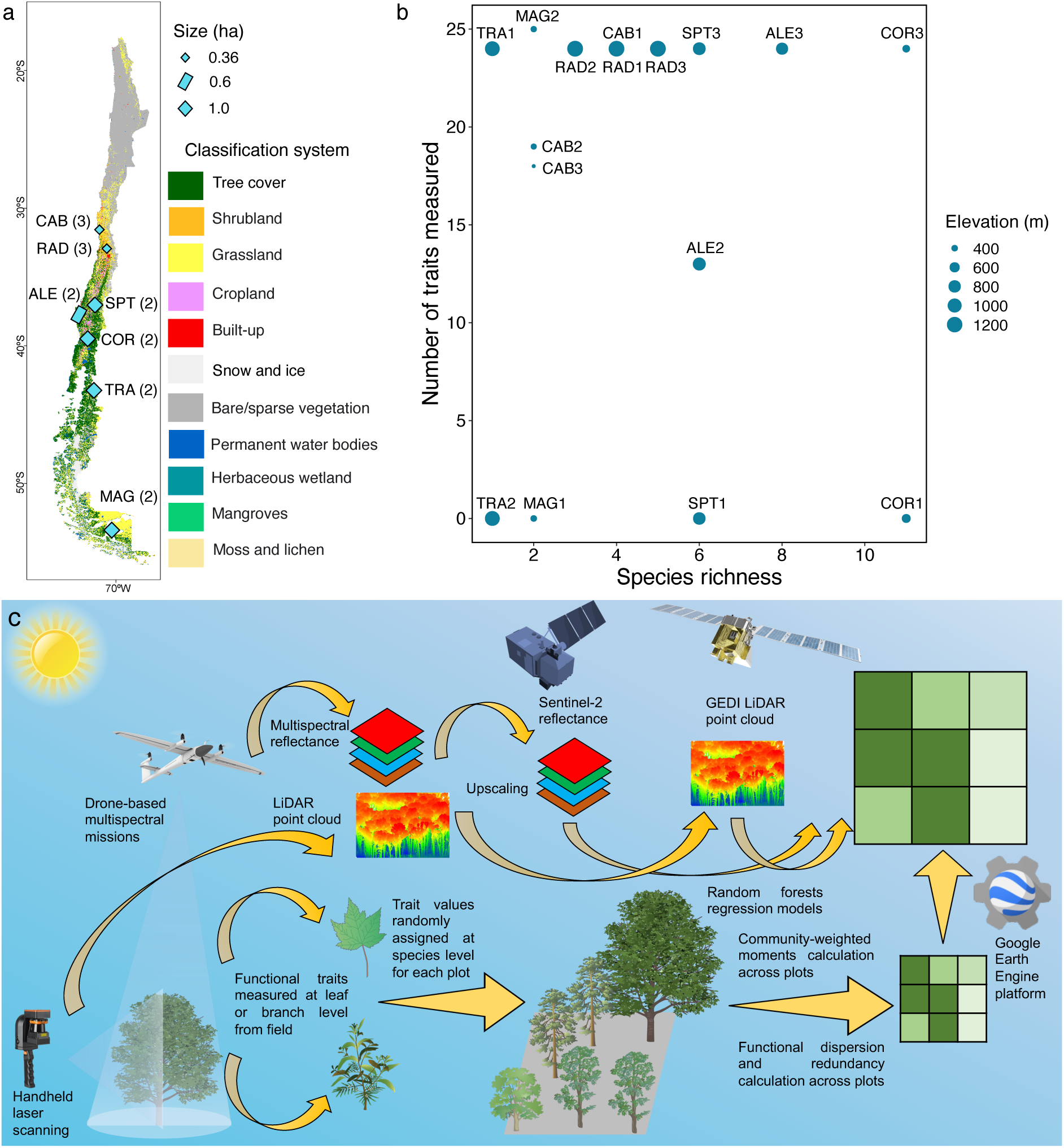
|Mapping plant functional composition across temperate forests in Chile. (a) Geographic locations of seven sites across a large latitudinal gradient. The location of all these plots is displayed as cyan rectangles with black boundaries, and the number of plots in each site is presented in the bracket after the abbreviation of each site name. Note that all the dots are magnified for visualisation purposes. The background map is based on the WorldCover v200 land use and land cover product; (b) Detailed information on species richness and number of traits measured in the field for each plot. The size of each dot indicates the elevation of the plot. Specifically, there is no trait measurement in plots COR1, MAG1, SPT1, and TRA2, but the species identified in these plots are covered in other plots already; (c) Mapping plant functional traits and assessing functional dispersion and redundancy with remotely sensed data using trait-based methods. This process integrates field-collected trait data with advanced remote sensing technologies. In the field, we gathered plant functional traits, multispectral images and LiDAR point clouds of studied forest plots. Community-weighted means (CWM) and community-weighted variances (CWV) were calculated, representing trait averages and variability, and functional dispersion and redundancy were derived from these traits at plot level. We used Random forests regression models to establish the link between multispectral, LiDAR and functional traits using the above measurements. To extend these insights from plot to landscape level, Sentinel-2 optical imagery and Global Ecosystem Dynamics Investigation (GEDI) LiDAR data were utilised, with the Random forests regression model facilitating the upscaling process, enabling a comprehensive evaluation of plant functional traits and dispersion and redundancy across larger spatial scales.

## Results

### The synergy between the environment and functional traits

We tested the importance of each input variable (VarImp) from multisource remotely sensed data in predicting the two community-weighted moments (mean: CWM; variance: CWV) of the leaf and stem functional traits selected (Extended Data Figs.1 to 20). These functional traits include four types: 1) morphological (Mor) traits: leaf fresh weight (FW), leaf dry weight (DW), leaf area (LA), specific leaf area (SLA), trunk wood density (TWD); 2) leaf nutrients (Nutr): nitrogen (N), phosphorus (P), calcium (Ca), and magnesium (Mg) content in leaves; 3) hydraulic (Hydr) traits: water potential at which 50% and 88% of hydraulic conductivity is lost (P50 and P88) respectively, and minimum water potential (midday water potential at the driest month) (WPmd), and 4) photosynthetic (Pho) traits: temperature at carbon compensation point (TmaxL), temperature of optimum photosynthesis (Topt), breadth of temperature optimum (TspanL), and temperature at which the maximum quantum yield of the photosystem II declines to 50% (T50). We also calculated mean variable importance scores of each of the seven groups of input data, i.e. spectral bands (red, red edge, and near-infrared (NIR)), vegetation indices (normalised difference vegetation index (NDVI), normalised difference red edge index (NDRE), soil-adjusted vegetation index (SAVI), and modified soil-adjusted vegetation index (MSAVI)), plant canopy height, climate (temperature and maximum climatological water deficit (MCWD)), hydrological stress (evapotranspiration (ET), evaporative stress index (ESI), and water use efficiency (WUE)), topography (slope and aspect), and soil (cation exchange capacity (CEC), sand, clay, pH water (pHH_2_O), and soil organic carbon (SOC)), to explore how different variables contribute to functional trait mapping (Fig.2).

**Fig.2.**
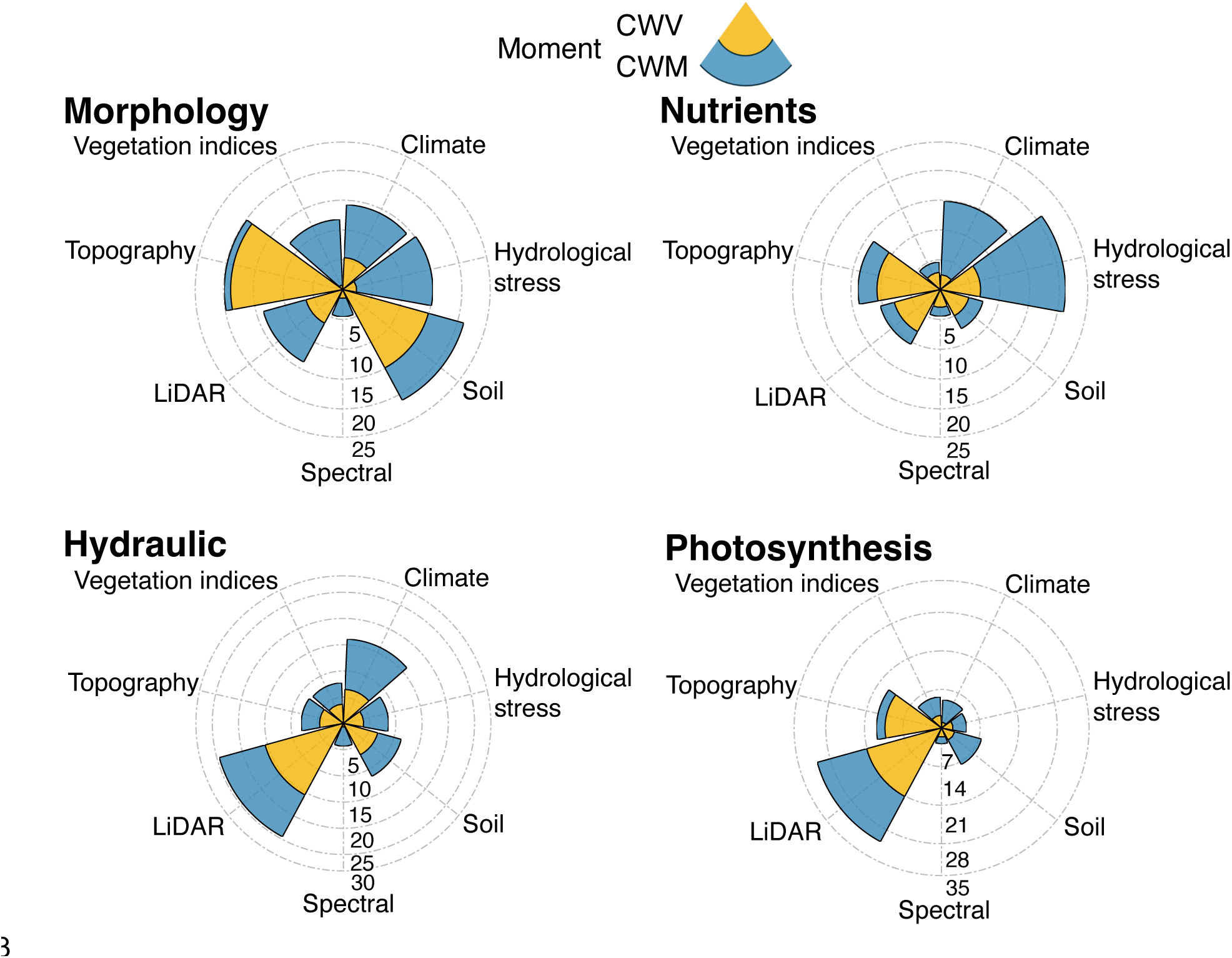
|Importance values of the different input variables for predicting the two community-weighted moments of the four categories of functional traits. Importance values range from 0 to 100. In each of the polar stacked bar charts, the seven types of input data were assigned specific spokes along which stacked bars were drawn, the height of each segment within the stacked bar corresponded to the importance of the variable it represented. Different colours were used to distinguish between the two community-weighted moments (CWM and CWV). Numbers alongside the grey chain axis denote axis scale values ranging from 0 to 100. VI and HS are abbreviations for vegetation indices and hydrological stress, respectively.

For CWM (Fig.2, Extended Data Figs.1 to 4), the top-ranking variable for predicting CWM_Mor_ and CWM_Nutr_ are hydrological stress (mean VarImp=31.99 for CWM_Mor_ and VarImp=36.22 for CWM_Nutr_), suggesting a strong influence of water status on the CWM of morphological traits and nutrients, while soil properties contribute the most in predicting CWM_Hydr_ (mean VarImp=23.32) and CWM_Pho_ (mean VarImp=36.00). In addition, climate follows closely after hydrological stress and soil properties and to predict CWM_Nutr_ (mean VarImp=21.74) and CWM_Hydr_ (mean VarImp=19.20). For CWV (Fig.2, Extended Data Figs.1 to 4), soil properties are identified as the most important variable in the case of morphology, nutrients, and hydraulics (mean VarImp=41.60, 24.97, and 33.99 for CWV_Mor_, CWV_Nutr_, and CWV_Hydr_, respectively). Notably, topography ranks first in predicting CWV_Pho_ (mean VarImp=23.67), and it is also the second most important predictor for CWV_Mor_ (mean VarImp=28.65) and CWV_Nutr_ (mean VarImp=21.23). To gain more detailed understandings of how the input data influence each specific trait, we established relationships between each single input band and the two community-weighted moments of each functional trait (Extended Data Figs.21 to 52 and Tables 2 to 9) using Pearson correlation analysis. Notably, climate and hydrological stress display significance in predicting functional traits, e.g. for for predicting CWM of TWD, WPmd, and Topt, and CWV of TWD and TmaxL, *P*<0.05 are found across MCWD and temperature; and for predicting CWM of SLA, leaf phosphorus content, TspanL, and T50, and CWV of TWD and WPmd *P*<0.05 are found across ET, ESI, and WUE. Additionally, vegetation indices are significant predictors (*P*<0.05) of all hydraulic traits except when using SAVI to predict WPmd.

## Morphology

### Drivers of functional dispersion and redundancy across the latitudinal gradient

We tested the environmental drivers of functional dispersion (FDis) and redundancy (FRed) (Fig.3 and Extended Data Figs.53 to 56) and calculated variable importance scores across the four functional trait categories (morphology, nutrients, hydraulics, and photosynthesis) for the same seven groups of input data mentioned in the subsection above. Overall, hydrological stress and soil properties emerge as the predominant drivers of both FDis and FRed across all trait groups, with notable mean VarImp of 27.97 and 21.28 for FDis, and 28.13 and 22.88 for FRed, respectively. Moreover, hydrological stress stands out as the only type of variable whose importance scores were consistently high for driving both FDis and FRed in all trait groups, while plant canopy height contributed little to predicting FRed of morphology, nutrients, and photosynthesis.

**Fig.3.**
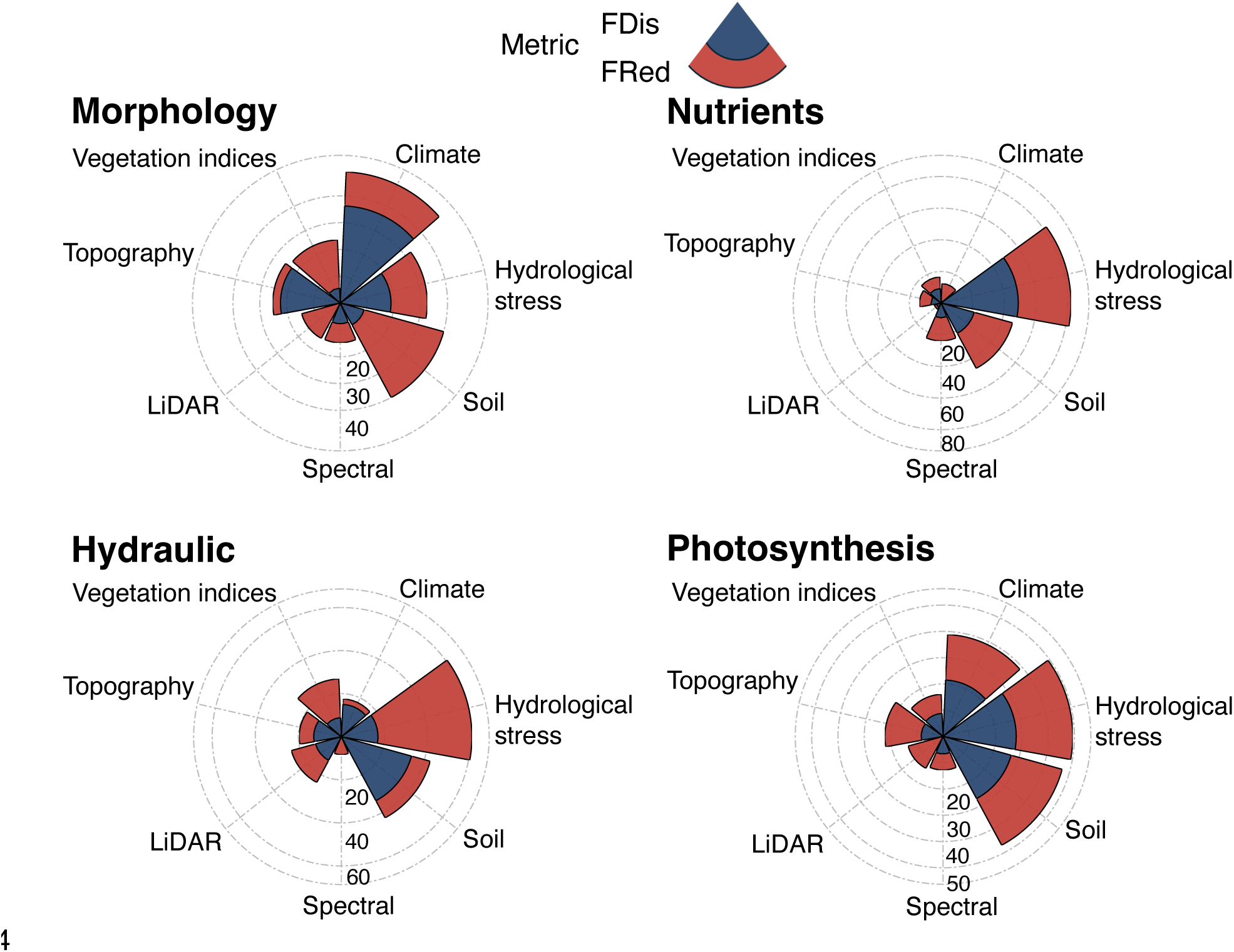
|Variable importance of the input variables for predicting functional dispersion and redundancy corresponding to the four categories of functional traits. Importance values range from 0 to 100. In each of the polar stacked bar charts, the seven types of input data were assigned specific spokes along which stacked bars were drawn. The height of each segment within the stacked bar corresponded to the importance of the variable it represented. Different colours were used to distinguish between the two functional diversity metrics. Numbers alongside the grey chain axis denote axis scale values. VI and HS are abbreviations for vegetation indices and hydrological stress, respectively.

Specifically, soil properties (VarImp=30.81 for FDis) and climate (VarImp=36.27 for FRed) emerge as the primary drivers in terms of morphology. For leaf nutrients, the importance of hydrological stress (VarImp=33.29 for FDis and VarImp=48.76 for FRed) and soil properties (VarImp=25.15 for FDis and VarImp=21.58 for FRed) is prominent. When considering hydraulic traits, hydrological stress (VarImp=43.60) and soil properties (VarImp=33.75) are found as the most important drivers of FDis and FRed, respectively. Lastly, photosynthetic FDis and FRed are notably driven by hydrological stress (VarImp=21.50 for FDis and VarImp=27.84 for FRed) and soil properties (VarImp=20.15 for FDis and VarImp=26.87 for FRed). Additionally, climate also demonstrates high importance in driving FDis (VarImp=17.32) and FRed (VarImp=21.48) of photosynthesis as it ranks third closely to soil properties in terms of both the metrics. Similarly, we also conducted Pearson correlation analysis between each functional dispersion and redundancy and the mean value of each input band, and noteworthy associations are found (Extended Data Figs. 57 to 64 and Tables 10 and 11). Remarkably, slope is significantly (*P*<0.05) associated with all FDis metrics but FDis_Hydr_, while WUE and slope are also significantly (*P*<0.05) associated with all FRed metrics.

### Spatial distribution of functional traits and functional dispersion and redundancy

By leveraging Random forests regression models, we obtained spatial predictions of the two community-weighted moments for each functional trait (Fig.4, all trait maps see Extended Data Figs. 65 to 72) and bivariate maps of functional dispersion and redundancy across the study area (Fig.5). For functional traits, there is a noticeable pattern in the latitudinal distribution between approximately 35°S and 42°S, where CWM and CWV values for almost all functional traits (excluding CWM_P_, CWM_P50_, CWM_P88_, CWM_T50_, CWV_SLA_, CWV_P_, CWV_WPmd_, and CWV_Topt_) are relatively low compared to neighbouring regions.

**Fig.4.**
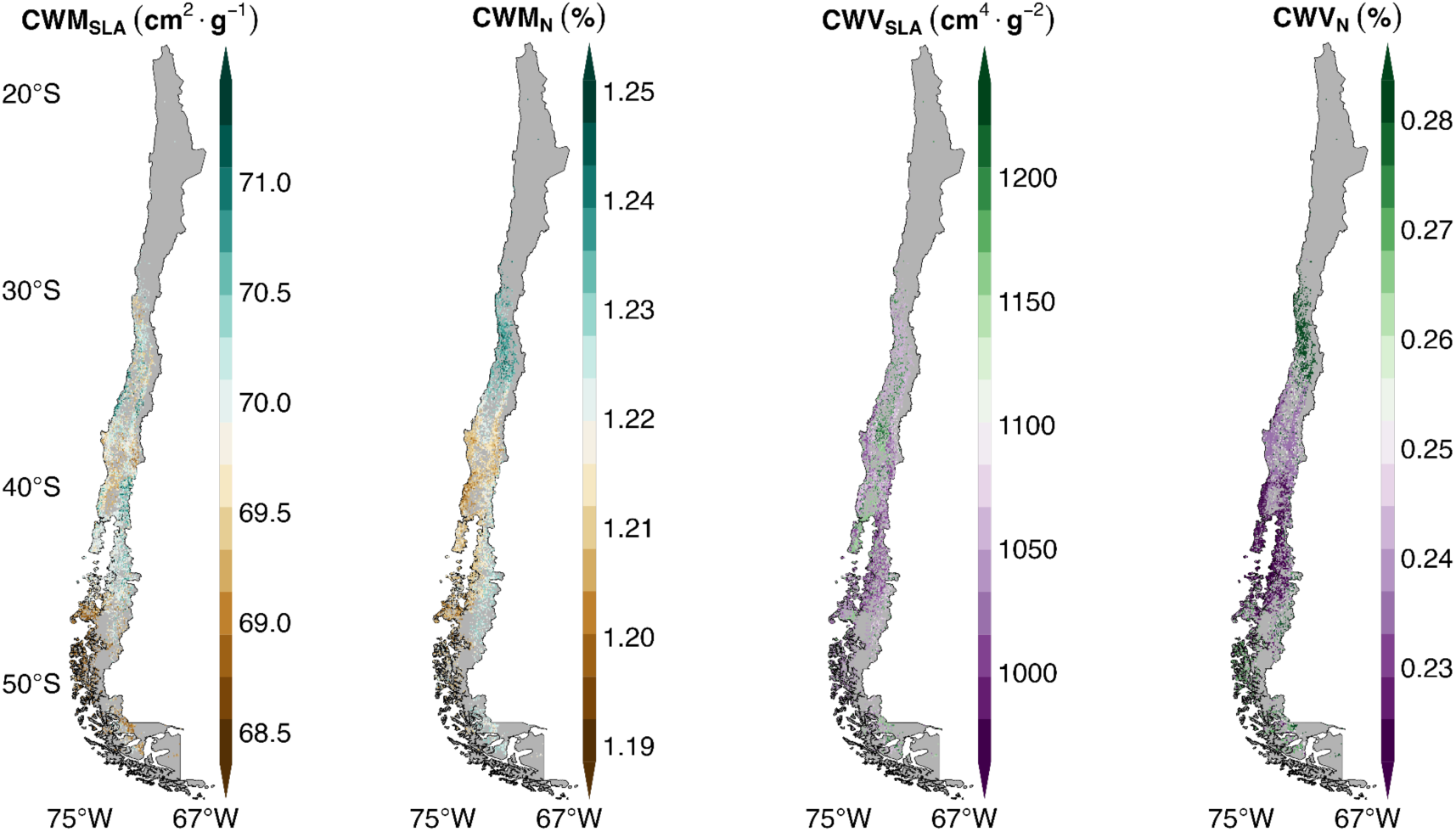
|Distribution maps of functional traits. Geographic variation in CWM and CWV for specific leaf area (SLA) and nitrogen concentration (N), respectively.

**Fig.5.**
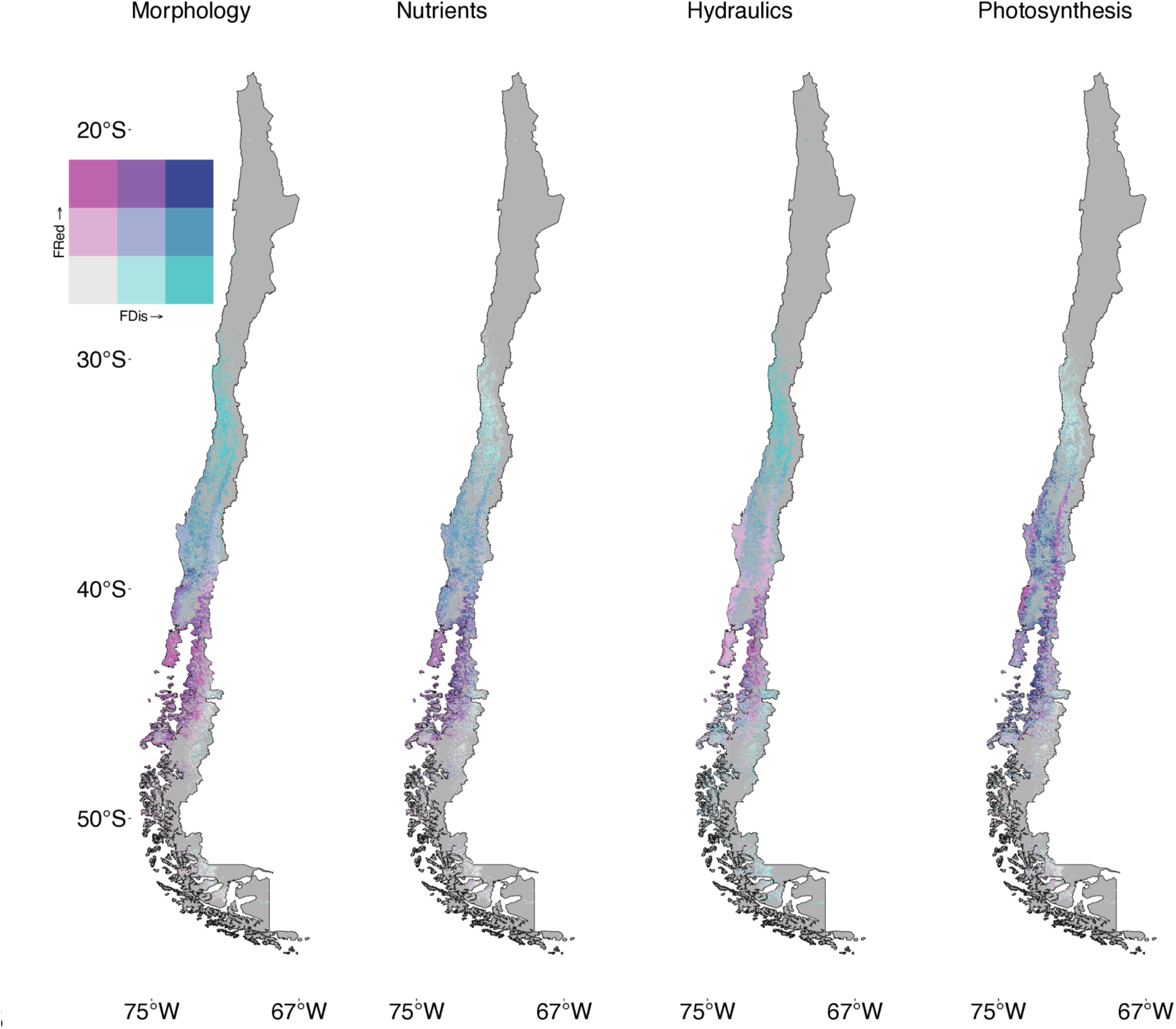
|Bivariate maps of FDis and FRed for morphologic, nutrients, hydraulics, and photosynthetic traits. Each bivariate map provides insights into the distribution of two key aspects of functional diversity within Chilean temperate forest ecosystems. In the legend on the left, the horizontal axis represents the range of FDis values, with “low”, “moderate”, and “high” indicating different levels, the vertical axis represents the range of FRed values, with “low”, “moderate”, and “high” indicating different levels. The square-shaped colour gradient visually conveys the joint distribution of FDis and FRed, where grey in the bottom left corner indicates the minimization of both variables, turquoise in the bottom right corner symbolises the maximisation of FDis while keeping FRed minimised, magenta in the top left corner represents the maximisation of FRed while maintaining FDis minimised, and navy in the top right corner signifies the joint maximisation of both variables.

We found distinct spatial distribution patterns of FDis and FRed across the gradient explored. Notably, we observed a latitudinal gradient wherein both FDis and FRed are high within a narrow band, primarily between latitudes 35°S and 42°S. Within this range, FDis exhibits a range from 0.54 to 2.06 with a mean value of 1.18 (mean values of 1.38, 1.26, 0.84, and 1.25 for FDis_Mor_, FDis_Nutr_, FDis_Hydr_, and FDis_Pho_, respectively), indicating a considerable variability in trait expression among plant communities. Similarly, FRed ranges from 0.24 to 0.63 with a mean of 0.46 (mean values of 0.44, 0.47, 0.48, and 0.46 for FRed_Mor_, FRed_Nutr_, FRed_Hydr_, and FRed_Pho_, respectively), highlighting a moderate degree of functional overlap among species. When delving deeper into subregions, a noteworthy divergence in the dominance of either metric emerges. Specifically, FDis is higher, with the mean value peaking at 1.33 (mean values of 1.65, 1.15, 1.30, and 1.20 for FDis_Mor_, FDis_Nutr_, FDis_Hydr_, and FDis_Pho_, respectively) between latitudes 30°S and 35°S, indicating a heightened variability and divergence of functional traits within plant communities. In contrast, within the latitudinal range of 42°S to 53°S, FRed dominates across the four trait groups with the mean value reaching 0.49 (mean values of 0.48, 0.51 0.53, and 0.46 for FRed_Mor_, FRed_Nutr_, FRed_Hydr_, and FRed_Pho_, respectively), indicating a higher level of functional similarity and potential overlap in resource use and environmental tolerance among species.

In addition, we extracted R^2^ (Extended Data Fig.73 and Tables 12 to 16), mean absolute error (MAE) (Extended Data Tables 12 to 16), and root mean square error (RMSE) (Extended Data Tables 12 to 16) to evaluate model performance (see Methods). In general, the mean R^2^ values are 0.61 and 0.44 for CWM and CWV, respectively. Specifically, predicting morphological and hydraulic traits based on Random forests regression models obtains a mean and R^2^=0.63 for CWM_Mor_ and R^2^=0.76 for CWM_Hydr_, and demonstrates suboptimal efficacy when applied to the mapping of CWM_Nutr_ (mean R^2^=0.53) and CWM_Pho_ (mean R^2^=0.54). In terms of mapping CWV, Random forests regression models achieve R2 of 0.50, 0.31, 0.46, and 0.47 for predicting CWV_Mor_, CWV_Nutr_, CWV_Hydr_, and CWV_Pho_, respectively. The observed mean R^2^ values from Random forests regression models suggest strong and moderate predictability in explaining the central tendency and variation of ecosystem characteristics by functional traits, respectively. The higher R^2^ values for CWM in comparison to CWV illustrate that the models demonstrate a relatively stronger ability to predict the central tendency of the trait distributions than the variance. Moreover, Random forests regression models in the context of assessing FDis and FRed demonstrate equivalent performance compared to mapping functional traits. The mean R^2^ for assessing the four groups (morphology, nutrients, hydraulics, and photosynthesis) of FDis and FRed is 0.44, 0.36, 0.61, and 0.60, respectively. The mean R^2^ values for assessing FDis and FRed are 0.48 and 0.52, respectively.

## Discussion

### Variation of plant functional traits across latitudinal and environmental gradients

The intermediate area includes the Mediterranean-Temperate transition zone^61^ (35.5°–39.5°S), as well as a rainy temperate climate zone south of 40°S (up to 48°S), characterised by a temperate climate with dry summers and warm temperatures^62,63^. The observed pattern of low CWM and CWV values for functional traits underscores the unique ecological dynamics of this region. Despite the overall favourable climatic conditions, including cold to moderate temperatures and ample moisture^64^, the reduced trait variability within plant communities suggests a distinct ecological regime characterised by ecological homogenization^65^. Forests in this area are characteristic of the temperate type with mostly evergreen species^66^, and the dominance of species with similar functional traits in this area may be attributed to the convergence of environmental factors, such as soil properties^67^ and topographic features^68,69^, which limits the dominance of more diverse functional strategies among plant species. Furthermore, more moderate temperatures, compared to the north and south of this area, and the absence of a pronounced dry season, especially toward the south, may contribute to the stabilisation of plant communities^70,71^ and the reduction in trait variability^72,73^ by minimising the influence of seasonal water availability^74^ on species composition and functional trait distributions.

In central Chile (spanning from approximately 30°S to 35°S), several functional traits such as FW, DW, TWD, P50, and P88 consistently exhibit both high CWM and CWV values, likely driven by Mediterranean climates present in this region. Considering that this pattern mainly appears in morphological and hydraulic traits that are paramount for species survival and reproduction in Mediterranean climates with prolonged drought, high temperatures, and concentrated rainfall in winters^75,76^, high CWM indicates prevalent trait values optimised for these climatic conditions^41,77,78^. Small, thick leaves and deep root systems, which enhance water retention and uptake efficiency are characteristic of this type of climate^79,80^. Concurrently, high trait variance suggests species’ abilities to finely tune these traits to specific microhabitats^81^ or respond to localised variations in resource availability^82^, contributing to the community’s ecological diversity and resilience^83,84^ within the Mediterranean ecosystems.

Conversely, the observed increase in both CWM and CWV values for most traits, reflects a shift in ecological dynamics within a cold oceanic (temperate/subpolar) climate regime in the southern region of Chile, spanning from approximately 42°S to 53°S. This higher trait variability suggests an ecological setting characterised by the influence of climatic factors associated with colder temperatures and maritime influences^85–87^. The dominance of traits with higher CWM values indicates a commonness of functional strategies adapted to the colder environmental conditions^27,78,88^ typical of this southernmost region. Traits related to temperature responses, such as TmaxL and Topt, exhibited notably higher CWM values, indicating the persistence of certain functional strategies adapted to the cooler temperatures^89–91^ in this region. Additionally, the dominance of cold-adapted species and the presence of diverse ecological niches shaped by maritime influences contribute to the higher trait variability observed in the southern region of Chile^92^.

### A model for understanding plant functional diversity and redundancy across temperate forests in South America

FDis and FRed help us to better understand the community-level functioning of plant species within the ecosystems. The spatial patterns of these metrics across Chilean forests offer a synthetical understanding of the ecological roles and interactions among plant species, providing essential information for adaptive responses to changing environmental conditions. The integration of *in-situ* functional traits and multi-source remotely sensed data not only provides a robust foundation for comprehensively mapping community functional compositions but also enables a holistic interpretation of plant functional diversity (here represented by FDis), functional redundancy (Fig.4) and ecosystem dynamics. The generated trait maps, coupled with the derived FDis and FRed, offer an overall view of the ecological dynamics across temperate forests in South America.

We reveal a combination of both high FDis and FRed emerging from dense tree cover and high species diversity in the intermediate region of Chile, spanning between latitudes ∼35°S and ∼42°S. This region is a biodiversity hotspot^6,93^, where favourable climatic conditions and varied habitats facilitate the coexistence of a diverse array of plant species with distinct functional traits^94,95^. In light of the transition from the Mediterranean climate in the north to the more temperate rainy climate in the south, the observed high FDis signifies the presence of a wide range of functional strategies among plant communities^96^, while the concurrent high FRed suggests a robustness in ecosystem functioning and resilience to ongoing environmental changes and human disturbances^13,97^. This balance between FDis and FRed suggests a high-resilience system, well-adjusted to a mild climate with ample moisture^98,99^. However, considering this region’s status as a highly threatened area in which forests have experienced extensive conversion to exotic forest plantations^17,100–102^, the high FDis and FRed emphasises the importance of preserving the ecological integrity of this region.

The high FDis observed in the central regions of Chile (∼30°S and ∼35°S), characterised by Mediterranean climate with long dry and hot summers, demonstrates the remarkable adaptability of the regional flora to the harsh environmental conditions^103^. This extensive variability in plant functional traits reflects the diverse strategies adopted by species to thrive in water-limited environments^104^. Such adaptations likely contribute to the high taxonomic diversity observed in this region, as plant species have evolved specialised traits (particularly plant structural traits, i.e. morphological traits like SLA^105,106^, and hydraulic traits like P50^106^ to optimise resource acquisition and utilisation^107^ and constrain plant gas exchange under drought conditions^108^. The findings highlight the ecological significance of South American semi-arid ecosystems in general, and Mediterranean ones in particular, as hotspots of biodiversity and contribute to the overall resilience of the continent’s flora.

In addition, our findings carry implications for understanding the ecological dynamics and biodiversity conservation across the continent, particularly in light of recent environmental events such as extreme fire weather driven by climate change and El Niño in 2017^109^ and 2023^110,111^ and climate-induced large-scale forest browning driven by the exceptionally severe megadrought since 2010^112,113^. The identification of regions with high FDis underscores the critical role of functional diversity in shaping ecosystem resilience in the face of environmental stressors^114,115^. These regions, which are already challenged by arid conditions, are further susceptible to the exacerbating effects of climate oscillations. The extreme fire weather events that occurred in this area in 2017 and 2023 sounded the alarm for human beings and reminded us of the vulnerability of these ecosystems to climate variability and the urgent need for effective conservation and management strategies. In this context, our research provides insights into the mechanisms driving ecosystem functioning and species coexistence in diverse habitats across latitudinal gradients in South America, including those impacted by extreme climatic events. Our findings highlight the importance of considering functional diversity and redundancy metrics alongside traditional measures of species richness and abundance in conservation planning, particularly in regions prone to climate extremes.

Across the region spanning from 42°S to 53°S, our results reveal a distinctive ecological setting characterised by high FRed, Antarctic climate, low temperatures, limited species diversity, and a unique resilience to the effects of climate change. This region stands out as a stark contrast to its northern counterparts, with its harsher climatic conditions contributing to a landscape dominated by few plant species. The observed high FRed suggests a prevalence of similar functional traits among the species present^116^, indicating convergent adaptation to the challenging environmental conditions. Despite the relatively low species diversity, the ecosystem in southern Chile demonstrates a remarkable capacity to thrive under the impacts of climate change, owing to the hardiness and resilience of its constituent species. The harsh climatic conditions create a hostile environment that only the most adaptable and resilient species can inhabit. Consequently, while southern Chile may appear less biodiverse compared to other regions, its ecosystems play a crucial role as refugia for species adapted to cold climates and serve as repositories of genetic diversity^117,118^. However, this resilience does not completely overcome the fragility of the ecosystem, as the combination of harsh environmental conditions and limited number of species makes it particularly susceptible to disturbances driven by climate^119^, pest and disease outbreaks^120,121^ and invasive species introductions. Thus, the conservation and sustainable management of this unique ecosystem in southern South America are of paramount importance, not only for safeguarding biodiversity but also for maintaining derived benefits to people and ensuring the resilience of the region’s fragile ecosystems to ongoing environmental challenges. Efforts like establishing new protected areas such as well-designed and fairly planned national parks and wildlife reserves or expanding existing ones^122^, rehabilitating degraded habitats through reforestation are in urgent need here.

### Mapping trait distributions and functional dispersion and redundancy using *in-situ* and remotely sensed data

The integration of plot-level *in-situ* plant functional traits including distinct trait categories and both multispectral and LiDAR remotely sensed data equipped with both drone- and satellite-based sensors provided not only a comprehensive understanding of trait distributions and variances, but also holistic insights into functional diversity and redundancy (inferred by FDis and FRed in this study) across temperate forests in South America. Using the drone-level data, we obtained detailed information on forest functional composition, while we used the satellite images to upscale results to a larger scale and enabled mapping functional traits and FDis and FRed spatially continuously and explicitly.

Sentinel-2 satellite multispectral images have been proven effective in mapping functional traits^56,123^ and assessing functional diversity and redundancy^55,57^ in diverse environments. Similarly, our study demonstrates the viability of Sentinel-2 data in assessing functional composition across temperate forests of South America. Vegetation indices generated from spectral bands rank second and obtain high importance values for CWM_Pho_ (mean VarImp=19.07), which is consistent with the idea that spectral indices are precise predictors of plant photosynthetic traits^124^. In addition, the LiDAR-derived plant canopy height variable contributes greatly to mapping hydraulic and photosynthetic traits (Fig.2) and inferring morphology- and hydraulics-related FDis (Fig.3) revealing the significance of forest vertical structure in shaping forest composition^125,126^.

Hydrological stress and soil properties perform as important variables for mapping CWM_Mor_, which aligns with the ecological understanding that hydrological stress, such as ET and WUE, primary measures of water availability, and soil properties, such as texture and organic carbon content, major determinants of soil fertility, can crucially affect plant growth^127^. In addition, the prominence of climate variables in determining CWM of nutrient and hydraulic traits suggests a strong ecological link between climate conditions and nutrient uptake and utilisation and water stress, respectively. Temperature, MCWD, and other climatic factors can influence variation in leaf nutrient content^128–130^ and water availability^131,132^, impacting the community’s ability to maintain nutrient enrichment and hydraulic functions under changing environmental conditions. Furthermore, topography contributes the most in mapping CWV_Pho_ and emerges as the second important factor in assessing CWV_Mor_, CWV_Nutr_, and FRed_Mor_, emphasising that the spatial variability in slope and aspect devotes importantly to the distribution of functional components across the landscape through water runoff and soil erosion^133,134^. Different aspects and slopes may create diverse microenvironments with varying soil fertility and water accumulation^135^, introducing microenvironmental gradients^136^ and impacting factors such as soil moisture, temperature, and light availability^137^. The observed importance of topography in morphological FRed indicates that these micro-environmental gradients likely filter functional traits and drive ecosystem resilience^138,139^. Steeper slopes may result in variations in soil depth and composition and aspect determines the amount of sunlight an area receives and thus affects temperature and moisture levels, leading to different species assemblages and morphological trait distributions^140^, which consequently influences FRed.

## Conclusions

Our approach provides an opportunity for mapping plant functional traits and assessing plant functional dispersion and redundancy explicitly and continuously across space and time. We identify a transitional zone, the intermediate latitudinal range between approximately 35°S and 42°S across forests in Chile, which may represent a critical threshold where potentially the most and least resilient forests to environmental change may converge due to the coexistence of high functional dispersion and redundancy. The co-occurrence of diverse functional traits and redundant functional groups suggests an ecosystem that possesses both the capacity to adapt to changing conditions and the resilience to withstand disturbances. In the southern region (approximately 42°S to 53°S), forests exhibit high functional redundancy and low functional dispersion, indicating stability but potential vulnerability to novel environmental challenges^141,142^. Conservation efforts such as ecosystem restoration, implementation of controlled burns and other fire management techniques, and establishment of green infrastructure elements should focus on maintaining stability while enhancing functional diversity to increase forest resilience. Conversely, in the central area (approximately 30°S to 35°S), high functional dispersion but low functional redundancy suggests adaptability to environmental changes but vulnerability to species loss. Conservation strategies should prioritise maintaining species richness to ensure functional redundancy and ecosystem stability. Lastly, the intermediate zone (approximately 35°S to 42°S) displays both high functional dispersion and redundancy, indicating being potentially the most resilient to environmental change. However, with reduced trait variability and increased environmental homogenization in this transitional zone, conservation efforts could target restoration, habitat connectivity enhancement, and community engagement to mitigate degradation and enhance ecosystem functioning.

## Methods

### Field measurements

We conducted measurements of *in-situ* census from 8,104 individuals with diameter at breast height (DBH) ≥ 5 cm at species level across 16 plots in seven sites along a wide latitudinal and elevational gradient (from 327.34 to 1,251.71 m) that cover 29 dominant tree species and two main forest types (evergreen and deciduous), namely Las Cabras (CAB), Radal Siete Tazas (RAD), San Pablo de Tregua (SPT), Alerce Costero (ALE), Correntoso (COR), Trapananda National Reserve (TRA), and Magallanes National Reserve (MAG) in Chile (see Fig.1a, b) from 10 February 2020 to 25 January 2022. To get a holistic view of community dynamics and ecosystem functioning and acquire a comprehensive assessment of functional diversity and redundancy within temperate forests in South America, we collected and measured a diverse set of foliar and canopy functional traits from different functional groups (Extended Data Tables 17 to 20; Extended Data Fig.74) including eight morphological traits (leaf fresh weight, leaf dry weight, leaf area, specific leaf area, leaf mass per area, leaf dry matter content, trunk wood density, and branch wood density), six nutrients (i.e. leaf chemical traits) (calcium, potassium, magnesium, nitrogen, and phosphorus content in leaves, and the ratio of leaf nitrogen content and leaf phosphorus content), five hydraulic traits (water potential at which 50% and 88% of hydraulic conductivity is lost (P50 and P88) respectively, minimum water potential, and safety margin P50 and P88), and five photosynthetic traits (temperature at carbon compensation point, temperature of optimum photosynthesis, photosynthesis rate at optimum temperature, breadth of temperature optimum, and temperature at which the maximum quantum yield of the photosystem II declines to 50%) from 770 trees, to gain insights into the physical structure of plants, nutrient cycling and ecosystem productivity, water transport strategies, and the efficiency of energy capture and utilisation, respectively.

### Identification of trait-trait correlations

We tested correlations between all functional traits (Extended Data Fig.75) and proposed the mean absolute within-class correlation index (MACI) that is defined as follows to identify dominant functional traits.

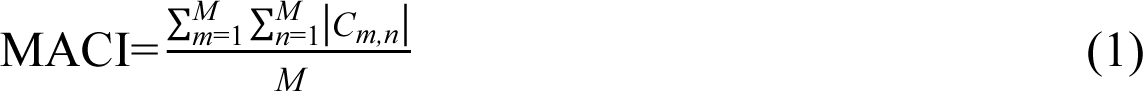

where *M* is the total number of functional traits in the same class, and in an *M*×*M* correlation matrix, *m* and *n* indicate the ordinal numbers of the functional traits, and *C_m,n_* stands for the correlation value between the *m*th and the *n*th functional traits.

By setting the threshold as 0.9, functional traits whose MACI values are greater than 0.9 were excluded because we assumed they highly correlated with other functional traits in the same category.

### Calculating functional trait composition

To take full advantage of the census and trait data of all individuals, we proposed a new method to pre-process the metadata. Firstly, for species in which traits of part of individuals were measured, we assigned trait values to individuals whose actual trait values were not measured randomly at the species level for every trait in every plot, then the arithmetic mean trait values calculated from all the individuals above were assigned to the left species for which only census was collected. Eventually, we ensured that all individuals participated in the following analyses. To better understand community dynamics, assembly processes, ecosystem functioning, and responses to environmental changes, we calculated the functional composition of each trait by quantifying the first two central moments—namely, the mean (first) and variance (second)—of trait distributions using the basal area of each individual as the weighting factor at community level^143^ to characterise the trait structure across biological communities, or a region, or other scales of interest^144,145^ considering they offer different facets to summarise the trait information. Specifically, these two community-weighted moments are defined as follows:

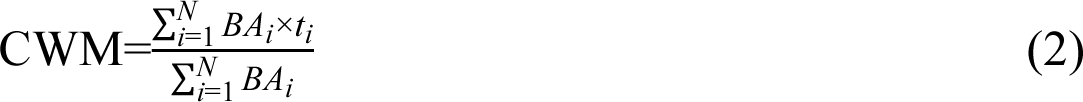

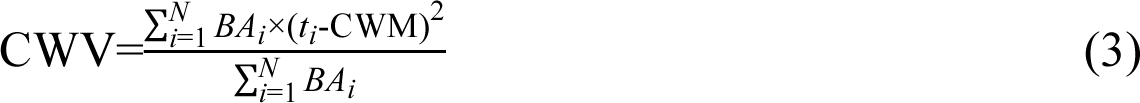

where CWM and CWV are abbreviations for community-weighted mean and community-weighted variance, *N* is the total number of tree individuals in a community, which, in this study’s case, a plot, *BA_i_* and *t_i_* denote the basal area and trait value of the *i*th individual.

To enhance our understanding of community ecological dynamics and the functional roles of species, we also generated the global functional trait space across all these plots (Extended Data Fig.76) and functional trait space for each plot (Extended Data Figs.77 to 92). In detail, functional trait spaces within individual plots help quantify how species within the same plot are functionally distinct or similar and provide information on how species respond to specific environmental gradients within those plots, and examining the global functional trait space allows for the assessment of the overall functional diversity and redundancy across multiple plots and for the identification of general patterns of trait variation along broader environmental gradients.

### Calculating functional dispersion and redundancy

Existing studies have developed multiple indices to measure functional diversity, various of which decomposed functional diversity into three main components^42,146–148^. Recently, more indices characterising other aspects of functional diversity have been proposed. Functional dispersion (FDis)^32^, defined as the mean distance in multidimensional trait space of individual species to the centroid of all species, to quantify the effective number of functionally distinct species for a given level of species dispersion and measure the spread and variability of species within the trait space. We calculated FDis using the “dbFD” function of the R “FD” package^149^:

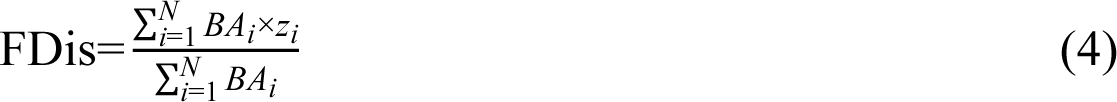

where *BA_i_* is the basal area of species *i* in a plot, and *z_i_* stands for the distance of species *i* in a plot to the weighted centroid of the *N* individual species in the trait space.

In addition, functional redundancy (FRed) measures the extent to which species within a community share similar functional roles and helps assess the resilience and stability of ecosystems by indicating whether there are backup species that can compensate for the loss of others in terms of functional roles^150^. To calculate FRed, we employed the “uniqueness” function of the R “adiv” package^151^. Specifically, in a community composed of *N* species:

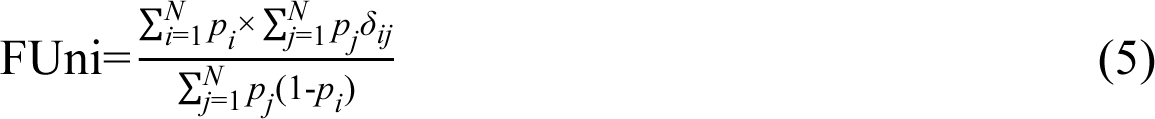

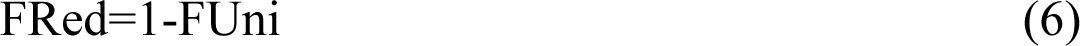

where FUni is functional uniqueness that refers to the distinctiveness of species with similar traits measured at community level, *p_i_* with 0<*p_i_*≤1 and 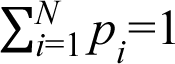 represents the relative abundance of a given species *i*, and 𝛿_*ij*_ denotes the pairwise functional dissimilarity between species *i* and *j* (𝛿_*ij*_ = 𝛿_*ji*_ and 𝛿_*ji*_ = *0*).

In summary, using FDis and FRed provides a more holistic understanding of functional diversity and redundancy; they complement each other in capturing different aspects of community structure and functioning. In detail, high dispersion with low redundancy may indicate a community with diverse strategies, while low dispersion with high redundancy may suggest a more functionally homogeneous community^152^. Hence, we use FDis and FRed as proxies to assess functional diversity and redundancy, respectively.

### Airborne remotely sensed data

We implemented drone missions to obtain high-resolution multispectral images using the MicaSense Altum multispectral camera across 11 plots. We also collected LiDAR data using the ZEB1 3D handheld laser scanner from GeoSLAM across 14 plots. For the plots without multispectral or LiDAR data, we obtained SuperDove multispectral (containing either five or eight bands) satellite images from Planet (https://www.planet.com/) and also extracted information from the global forest canopy height (GFCH) product^153^ generated by combining forest structure measurements from the GEDI LiDAR system with time-series data from Landsat images (Extended Data Tables 21 and 22 for details).

In terms of multispectral images, we kept the blue, green, red, red edge, and NIR bands due to their consistent presence in both drone images and SuperDove satellite images. We also extracted additional four vegetation indices based on these five original spectral bands to assess functional traits and functional diversity and redundancy (Extended Data Table 23). As for LiDAR data, we generated plant height from the LiDAR point cloud.

### Environmental variables

#### Climatic covariates

Climatic variables are critical for understanding plant adaptability, responses to climate change, and ecosystem functioning^154–156^. They provide insights into the physiological mechanisms of plant species, influence species distribution and range limits, and help predict future ecosystem services and stability and explain why certain functional traits are key in ecosystems. This explanation is crucial for estimating how changes in climate might influence ecosystem functioning and stability and informs strategies for mitigating the impact of climate change on ecosystems and biodiversity. Therefore, we gained three long-term monthly climatic variables, namely temperature, vapour pressure deficit (VPD), and climatological water deficit (CWD), using the TerraClimate dataset^157^ at a spatial resolution of 1/24 degrees (∼4-km) from January 1983 to December 2022 (https://www.climatologylab.org/terraclimate.html). Furthermore, we calculated the maximum climatological water deficit (MCWD), which is defined as the most negative value of CWD, to describe the accumulated water stress that occurs across a dry season^158^:

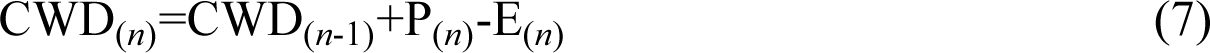

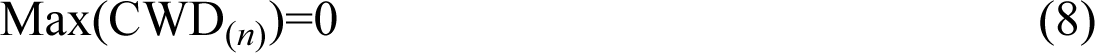

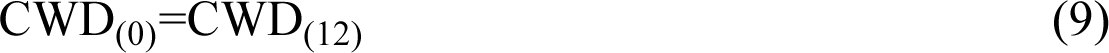

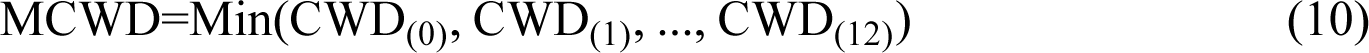

where *n* (*n*=1, 2, …, 12) is the index of a month over a calendar year, P and E are precipitation (mm/month) and evapotranspiration (mm/month). We set CWD_(6)_=0 as June is normally the wettest month in Chile^159^ by assuming the soil is saturated and applied this calculation to the mean annual cycle of precipitation^158^. To facilitate understanding and modelling, we calculated the additive inverse of MCWD so that positive MCWD values indicate increases in water stress^160^.

#### Soil conditions

Soil properties, encompassing parameters such as chemical soil (cation exchange capacity at pH 7 (CEC, in mmol(c)/kg), pH water (pHH_2_O, pH×10), and soil organic carbon (SOC, in dg/kg)) and physical soil (clay and sand, in g/kg), play a pivotal role in mapping functional traits and assessing functional diversity and redundancy. CEC helps understand nutrient availability, while soil texture determines water-holding capacity and root penetration, influencing the suitability of different plant species. pHH_2_O reveals soil acidity or alkalinity, aiding in plant selection, while SOC informs about stress conditions. Additionally, soil properties impact microbial communities, plant-soil feedback, and ecosystem functions, making soil data an indispensable tool for plant functional traits^161,162^ and ensuring the enhancement of functional diversity^163^ and redundancy^164^. Consequently, we gathered all the soil information mentioned above from SoilGrids (https://soilgrids.org/) with a spatial resolution of 250 m^165^ and calculated mean values of these five metrics from the top 30 cm.

#### Topography

Topological data, with their key elements of slope and aspect, exerts a profound influence on the mapping of functional traits and the assessment of functional diversity and redundancy. Slope and aspect create microclimatic variations, influencing temperature and moisture gradients, which in turn impact the distribution of plant species with diverse functional traits. These variations underpin the adaptation of plants to specific ecological niches, leading to a heterogeneous landscape that supports a wide array of functional traits. Topography’s importance extends to its role in succession patterns, species interactions, conservation efforts, and sustainable land management, making it an essential factor in understanding and preserving the functional diversity of ecosystems. So we got the highest-resolution digital elevation model of the earth^166^ from the shuttle radar topography mission launched on 11 February 2000 by the National Aeronautics and Space Administration at 30 m (https://cmr.earthdata.nasa.gov/search/concepts/C1000000240-LPDAAC_ECS.html) and derived slope and aspect from the original digital elevation model.

#### Hydrological stress

As a key component of the global water cycle, hydrological stress is central to ecosystem functions^167^, of which, evapotranspiration (ET) serves as a key indicator of water availability and an essential variable that links ecosystem functions, surface water cycle, carbon and energy cycle, and climate feedback^168,169^, crucial for understanding the water requirements of diverse plant species; the Evaporative Stress Index (ESI) aids in monitoring drought conditions, providing insights into plant stress and resilience^170^; and water use efficiency (WUE). These data contribute to predicting the impact of environmental change on plant communities and enable our understanding of how plants adapt to and thrive under various hydrological stress conditions, ultimately supporting the resilience and functional diversity of ecosystems. We selected the ECOsystem Spaceborne Thermal Radiometer Experiment on Space Station (ECOSTRESS) launched to the International Space Station on 29 June 2018 by the National Aeronautics and Space Administration^171,172^ and extracted ET, ESI, and WUE as indicators of hydrological stress. Specifically, we accessed 4383, 3917, and 4095 tiles for ET, ESI, and WUE, respectively, through AppEEARS (https://appeears.earthdatacloud.nasa.gov/) from 01 November 2019 to 31 January 2023.

### Satellite remotely sensed data

We utilised Sentinel-2 high-resolution and multispectral data launched by the European Space Agency as part of the Copernicus programme^173^ and the GFCH product mentioned above for predicting functional traits and functional diversity and redundancy across the whole forest area in Chile. The Sentinel-2 mission comprises a constellation of two identical satellites, Sentinel-2A and Sentinel-2B, operating synergistically to capture imagery of the Earth’s surface. These satellites are equipped with a sophisticated multispectral instrument, capturing data across 13 spectral bands, ranging from visible to infrared wavelengths. This extensive spectral coverage allows for detailed analysis of vegetation health, land cover changes, and water quality, making Sentinel-2 images invaluable for applications in ecology, forestry, and climate studies. The open-access nature of the data facilitates global scientific research, enabling a deeper understanding of the Earth’s dynamic and ever-changing features.

### Functional traits mapping and functional dispersion and redundancy assessment at plot level

We calculated the first two central moments for all the remotely sensed data, climatic covariates, soil properties, topographical variables, and hydrological stress metrics mentioned above across each plot. For input band *x*:

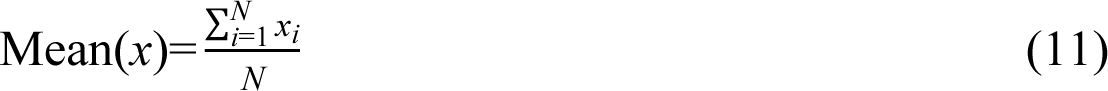

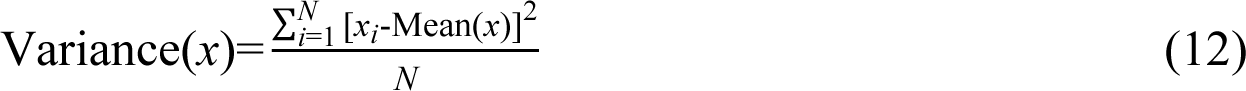

where *x_i_* is the *i*th value of the input band *x*, *N* is the total number of data points.

For each of the central moment of the data, they were merged together and divided into seven categories, namely spectral bands (blue, green, red, NIR, and red edge), vegetation indices (NDVI, NDRE, SAVI, and MSAVI), structure (height), climate (temperature, VPD, and MCWD), hydrological stress (ET, ESI, and WUE), topography (slope and aspect), and soil (CEC, clay, sand, pHH_2_O, and SOC). To simplify the model and avoid overfitting, we also tested the correlation between all variable’s mean values (Extended Data Fig.93) and applied the MACI to each of them, and dropped those whose MACI values were greater than 0.9. Eventually, the first two central moments of red, red edge, NIR, NDVI, NDRE, SAVI, MSAVI, height, MCWD, temperature, ET, ESI, WUE, slope, aspect, CEC, clay, sand, pHH_2_O, and SOC were selected as predictors of the Random forests regression^174^ model to predict the pairwise two community-weighted moments for each of the selected functional traits, and we used the mean value of these predictors to assess FDis and FRed. To get the optimal values of the two key parameters of Random forests regression, number of trees (ntree) ranging from 5 to 100 with a step of 5 and the number of variables to be randomly selected at each split (mtry) ranging from 1 (minimum number of input features) to 20 (maximum number of input features), we allocated 70% of the data for training purposes and reserved the remaining 30% for model testing, and we calculated the R^2^, mean absolute error (MAE, defined as below), and root mean square error (RMSE, defined as below) to comprehensively evaluate the performances of all Random forests regression models and tuned the model on the basis of leave-one-out cross-validation (LOO-CV) using the “caret” package in R^175^. The optimal model is determined as the one with minimum RMSE value.

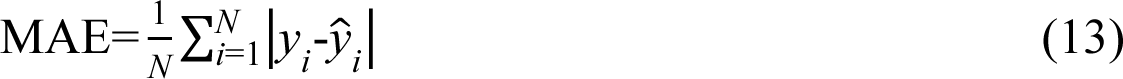

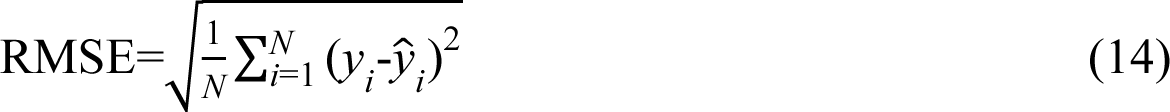

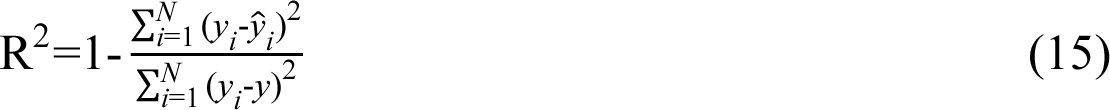

where *y_i_* and *ŷ_i_* represent the *i*th observed and predicted value, respectively, and *N* indicates the total number of observations.

### Scaling up from plots to temperate forest ecosystems

We identified forest areas in Chile by generating a forest mask that is defined as a combination of tree cover and shrubland from the WorldCover v200 land use and land cover product (https://worldcover2021.esa.int/) at 10 m released by the European Space Agency^176^. Then we gathered Sentinel-2 multispectral satellite remotely sensed images from 2019 to 2023, and we conducted pre-processings and applied all the optimal parameters gained from above to the Random forests regression model by leveraging the Google Earth Engine (https://earthengine.google.com/) cloud computing platform^177^ to predict functional traits and functional diversity and redundancy in forests across Chile (Fig.1c).

## Acknowledgements

X.D. received the Pay It Forward Scholarship (by China Oxford Scholarship Fund, Oxford) and the New Blackfriars Scholarship (by Blackfriars Hall, Oxford). The data collection for this project was funded by the Natural Environment Research Council (NERC) ARBOLES project (Grant: NE/S011811/1). J.A.G. is funded by the NERC (Grant: NE/T011084/1) and the Oxford University John Fell Fund (10667). S.D. is funded by an Oxford Martin School Fellowship. Y.M. is supported by the Frank Jackson Foundation.

## Author contributions

X.D. and J.A.G. conceived the idea and designed the framework for the manuscript. X.D., D.E.C., and J.A.G. developed and executed the methodology. Field plots and traits data were installed and collected by D.E.C., R.U.J., J.A.G and Y.M. Results were generated by X.D., with visualisation by X.D., H.Z., W.S.M., and A.R. X.D. wrote the first draft of the manuscript, with critical feedback from D.E.C., R.U.J., A.R., H.Z., D.G., S.D., Y.M., and J.A.G.

## Notes

### Competing Interest Statement

The authors have declared no competing interest.

