## Supplementary for "Higher functional resilience of temperate forests at intermediate latitudes of a large latitudinal gradient in South America"

Xiongjie Deng *et al*.

Extended data

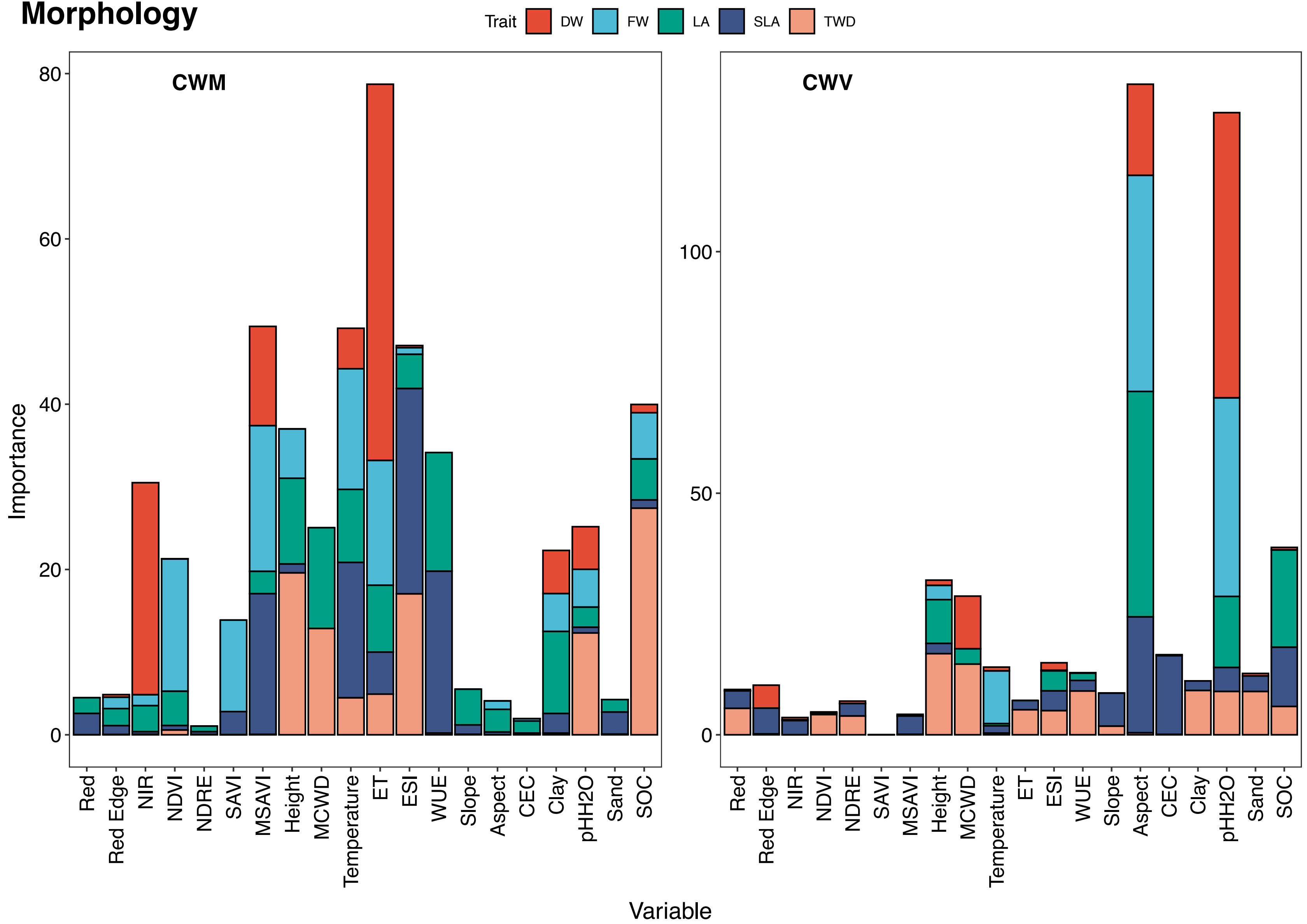


**Fig.1|The contribution of each variable in predicting morphological traits.** Variable importance of all variables inputted into Random forests regression models for predicting the two community-weighted moments of morphological traits. Each panel contains two stacked bar charts, representing the variable importance of all variables in predicting CWM and CWV of functional traits, respectively. All stacked bar charts are arranged in the order of spectral bands, vegetation indices, plant canopy height, climatic covariates, hydrological stress, topography, and soil conditions. Each stacked bar denotes the importance of an input variable for predicting functional traits with different segments corresponding to distinct functional traits. Colours are used to distinguish between different functional traits, refer to the legend for colour-key associations. See Extended Data Figs. 5 to 9 for more details about variable importance for predicting each morphological trait individually.

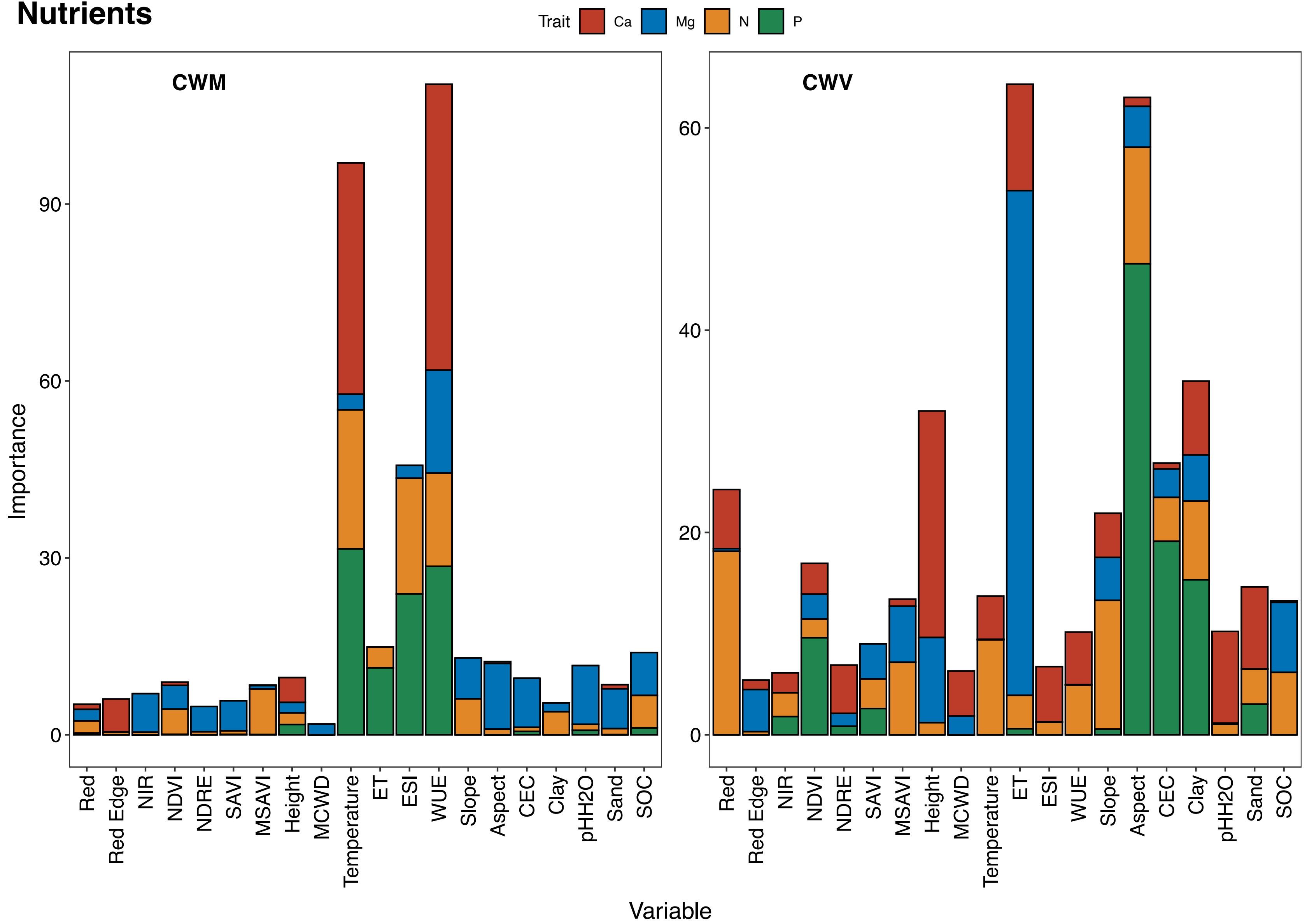


**Fig.2|The contribution of each variable in predicting nutrient traits.** Variable importance of all variables inputted into Random forests regression models for predicting the two community-weighted moments of nutrient traits. Each panel contains two stacked bar charts, representing the variable importance of all variables in predicting CWM and CWV of functional traits, respectively. All stacked bar charts are arranged in the order of spectral bands, vegetation indices, plant canopy height, climatic covariates, hydrological stress, topography, and soil conditions. Each stacked bar denotes the importance of an input variable for predicting functional traits with different segments corresponding to distinct functional traits. Colours are used to distinguish between different functional traits, refer to the legend for colour-key associations. See Extended Data Figs. 10 to 13 for more details about variable importance for predicting each nutrient trait individually.

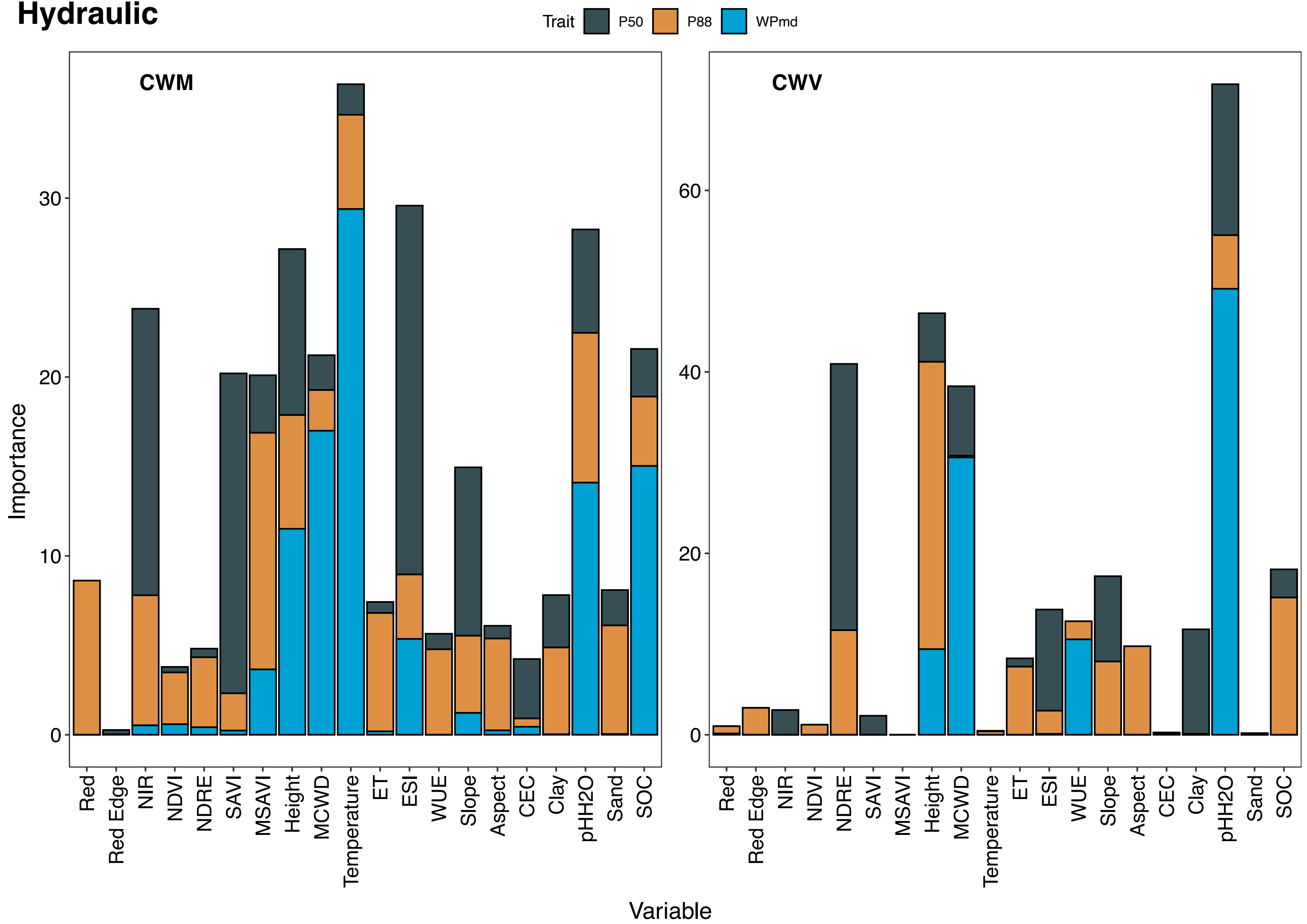


**Fig.3|The contribution of each variable in predicting hydraulic traits.** Variable importance of all variables inputted into Random forests regression models for predicting the two community-weighted moments of hydraulic traits. Each panel contains two stacked bar charts, representing the variable importance of all variables in predicting CWM and CWV of functional traits, respectively. All stacked bar charts are arranged in the order of spectral bands, vegetation indices, plant canopy height, climatic covariates, hydrological stress, topography, and soil conditions. Each stacked bar denotes the importance of an input variable for predicting functional traits with different segments corresponding to distinct functional traits. Colours are used to distinguish between different functional traits, refer to the legend for colour-key associations. See Extended Data Figs. 14 to 16 for more details about variable importance for predicting each hydraulic trait individually.

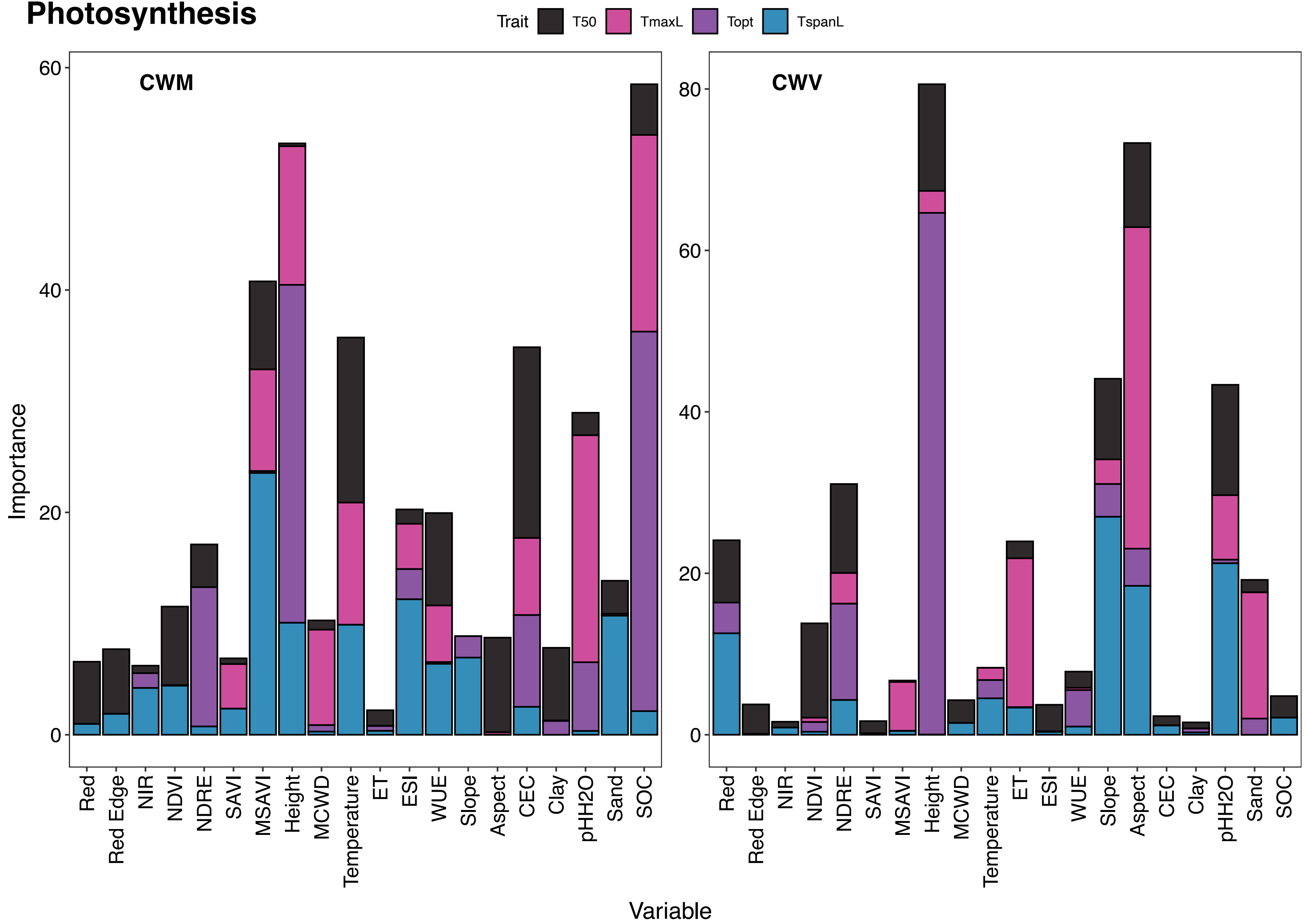


**Fig.4|The contribution of each variable in predicting photosynthetic traits.** Variable importance of all variables inputted into Random forests regression models for predicting the two community-weighted moments of photosynthetic traits. Each panel contains two stacked bar charts, representing the variable importance of all variables in predicting CWM and CWV of functional traits, respectively. All stacked bar charts are arranged in the order of spectral bands, vegetation indices, plant canopy height, climatic covariates, hydrological stress, topography, and soil conditions. Each stacked bar denotes the importance of an input variable for predicting functional traits with different segments corresponding to distinct functional traits. Colours are used to distinguish between different functional traits, refer to the legend for colour-key associations. See Extended Data Figs. 17 to 20 for more details about variable importance for predicting each photosynthetic trait individually.

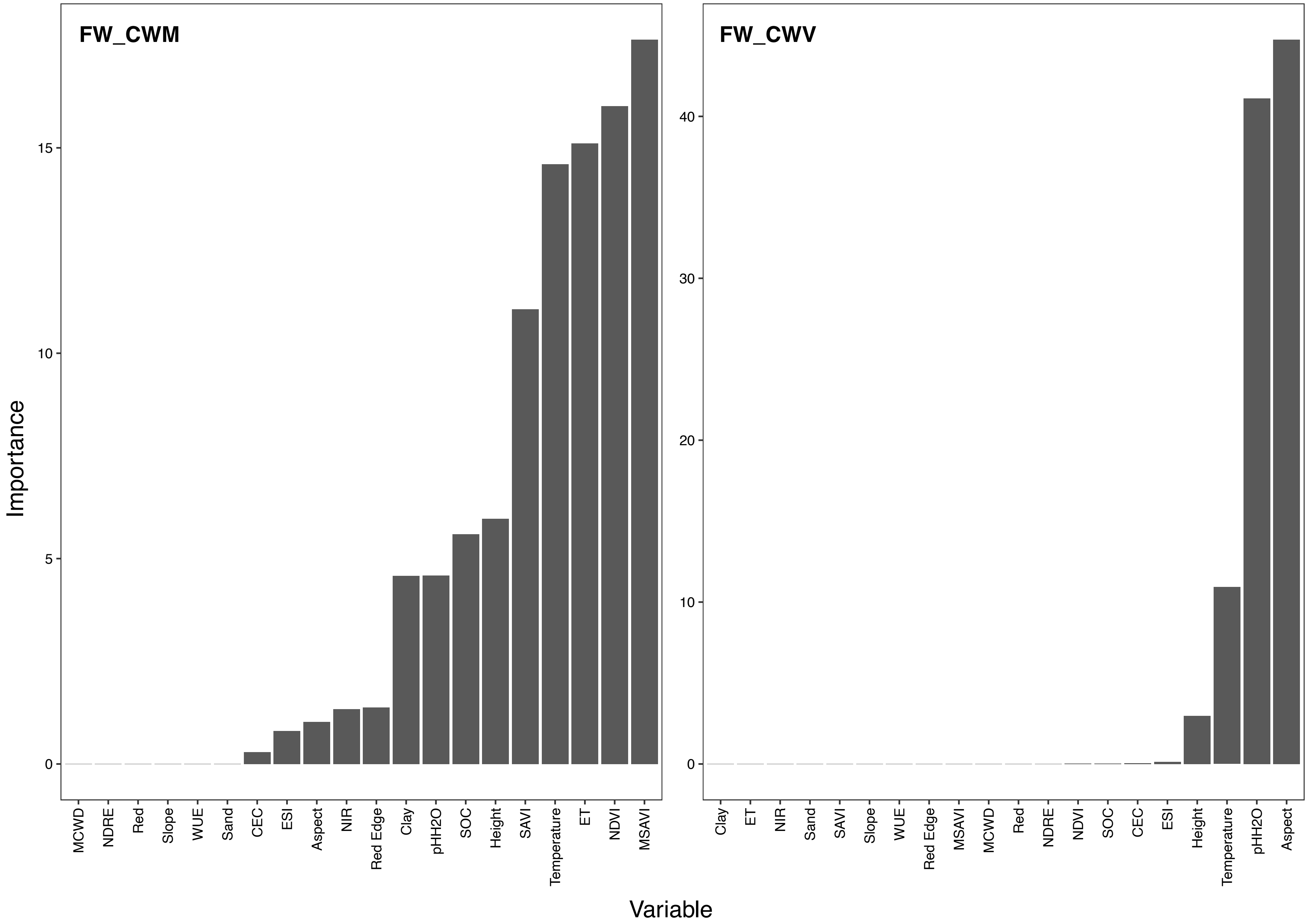


**Fig.5|Variable importance of each input band for predicting FW.**

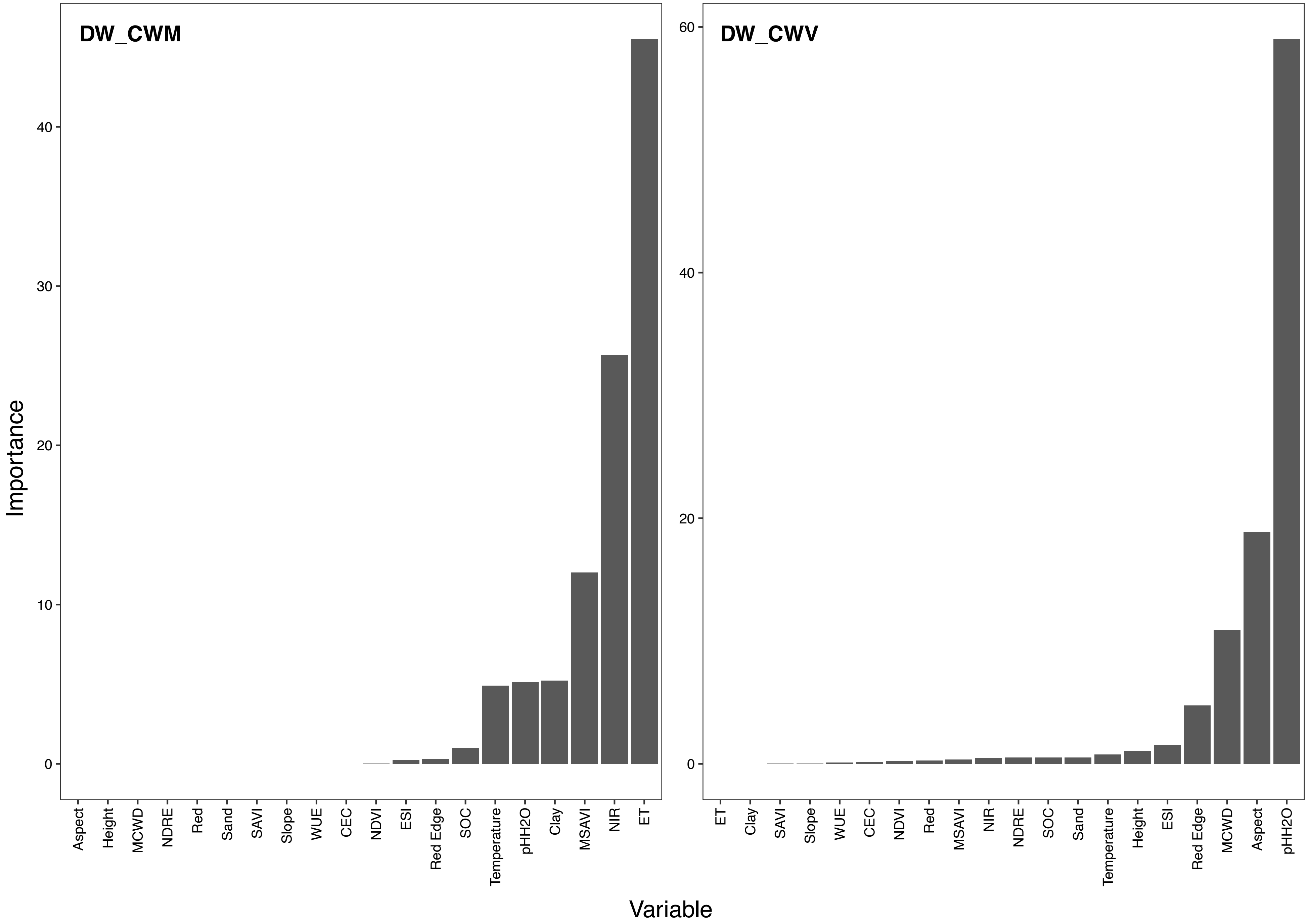


**Fig.6|Variable importance of each input band for predicting DW.**

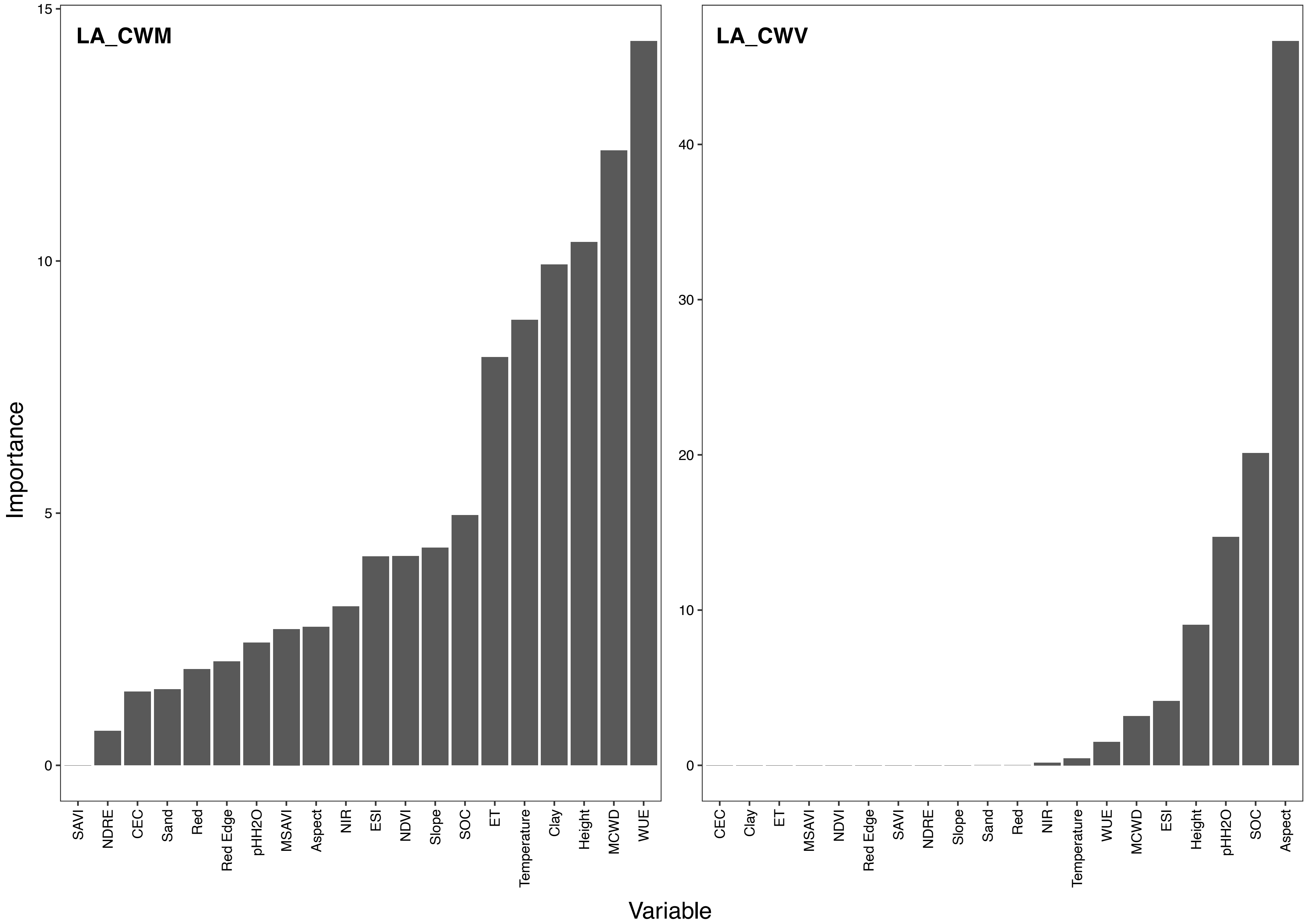


**Fig.7|Variable importance of each input band for predicting LA.**

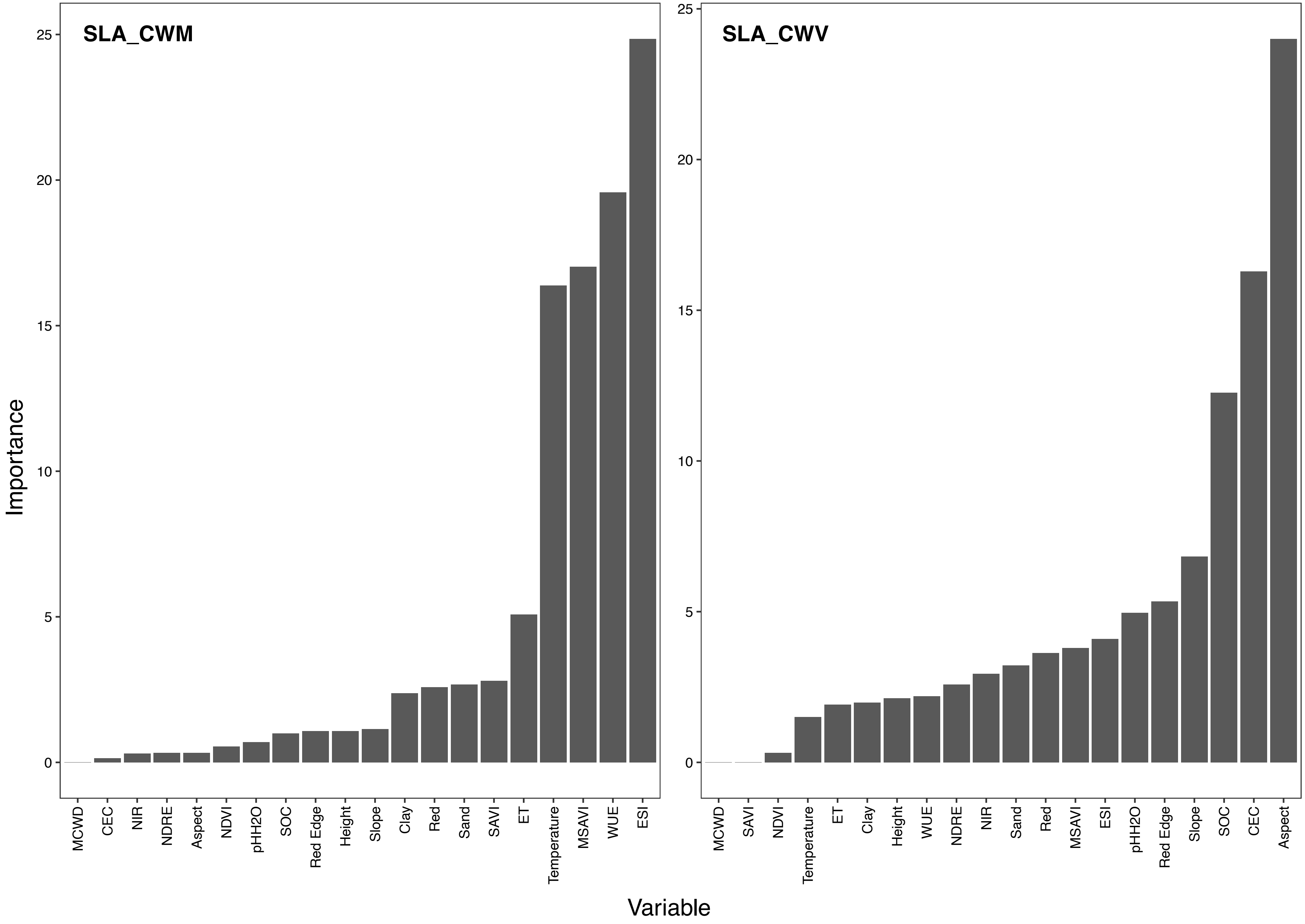


**Fig.8|Variable importance of each input band for predicting SLA.**

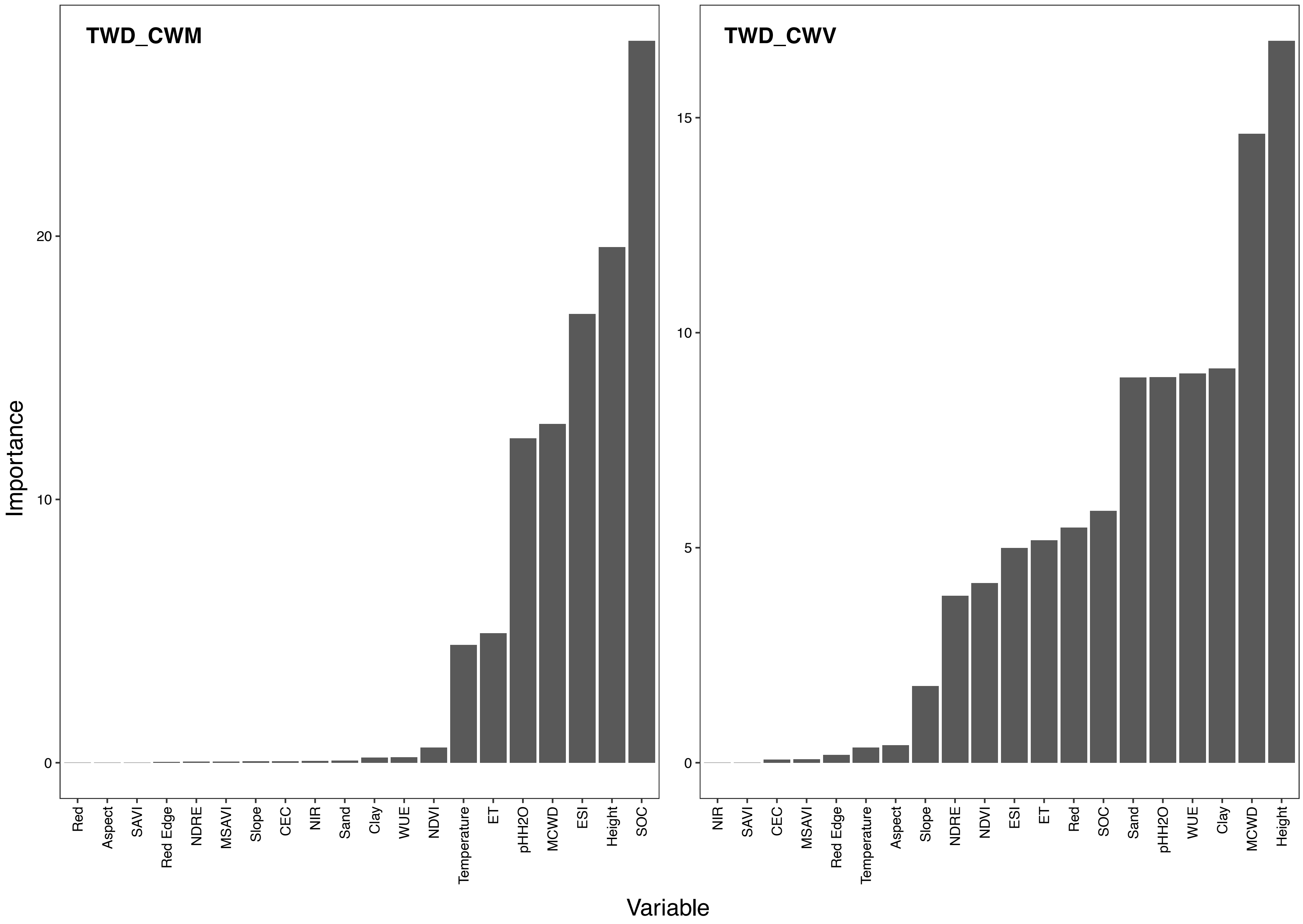


**Fig.9|Variable importance of each input band for predicting TWD.**

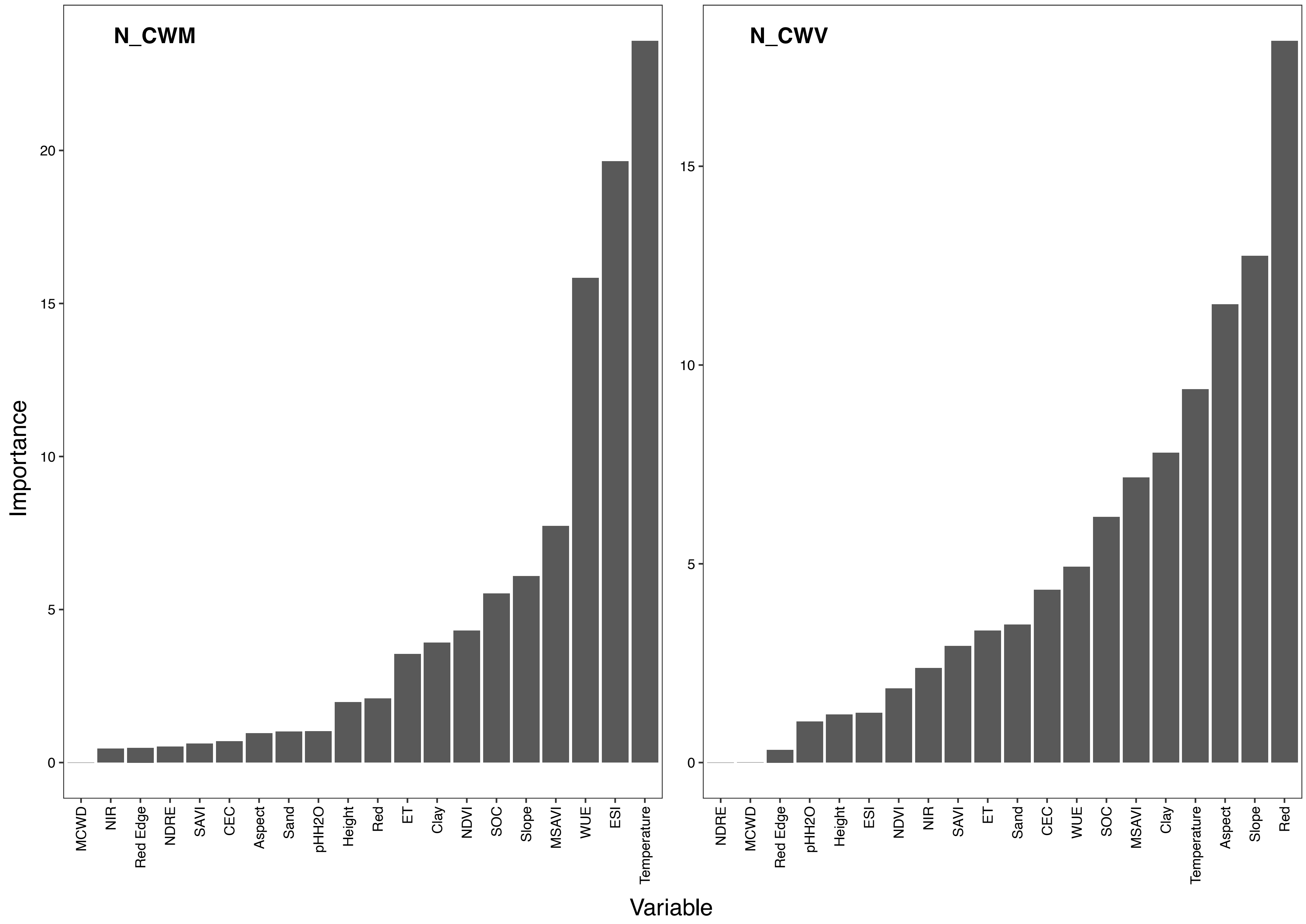


**Fig.10|Variable importance of each input band for predicting N.**

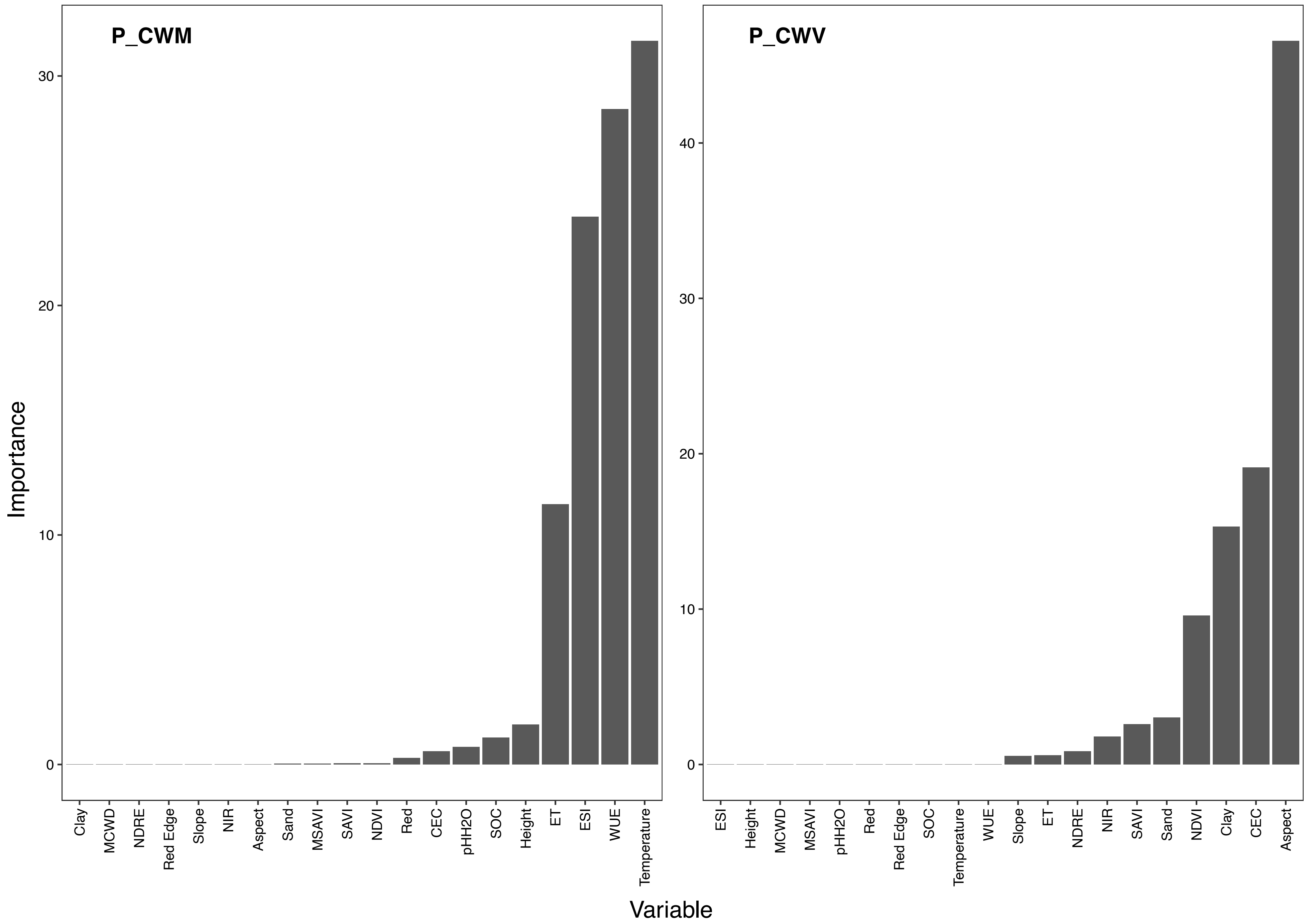


**Fig.11|Variable importance of each input band for predicting P.**

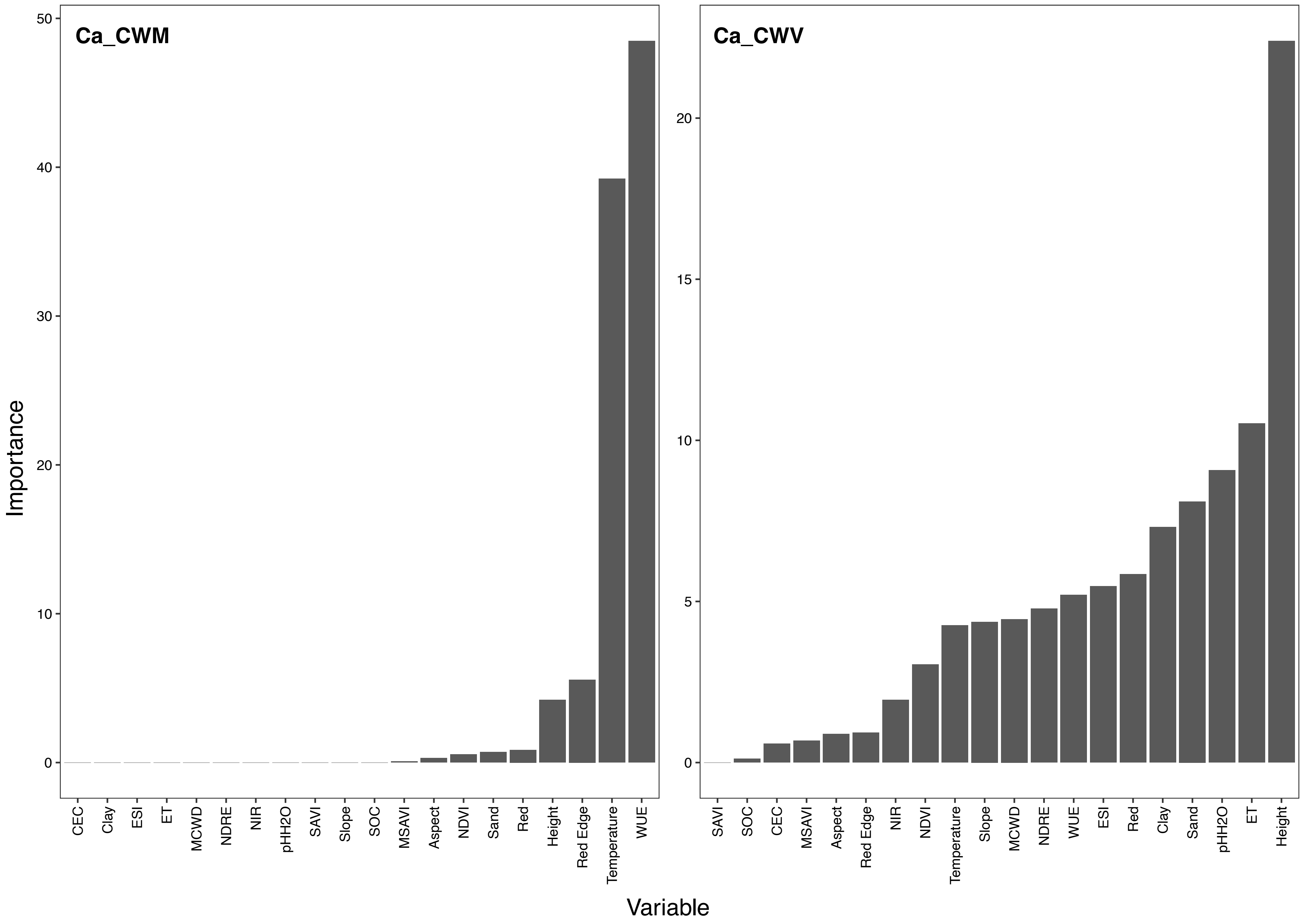


**Fig.12|Variable importance of each input band for predicting Ca.**

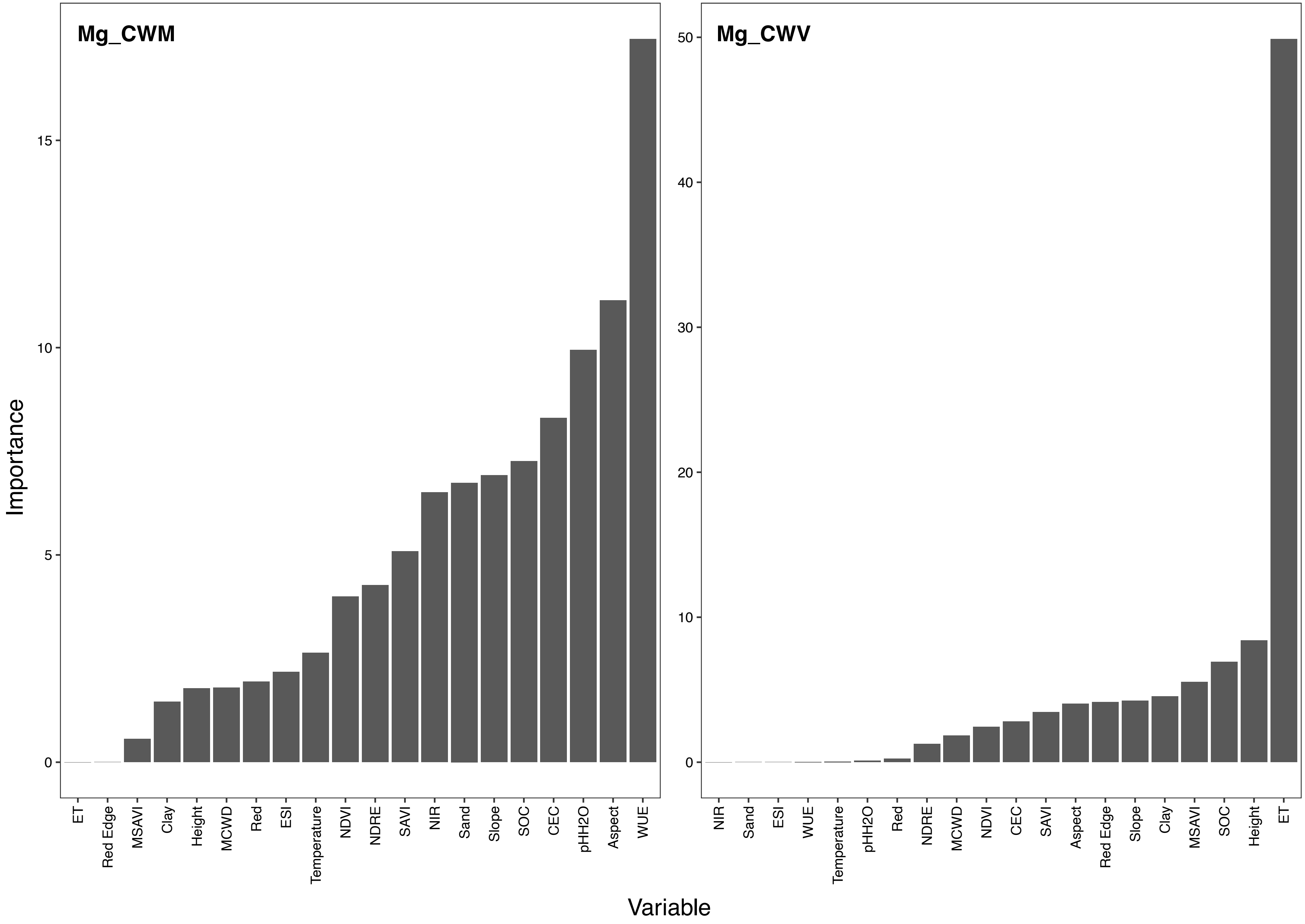


**Fig.13|Variable importance of each input band for predicting Mg.**

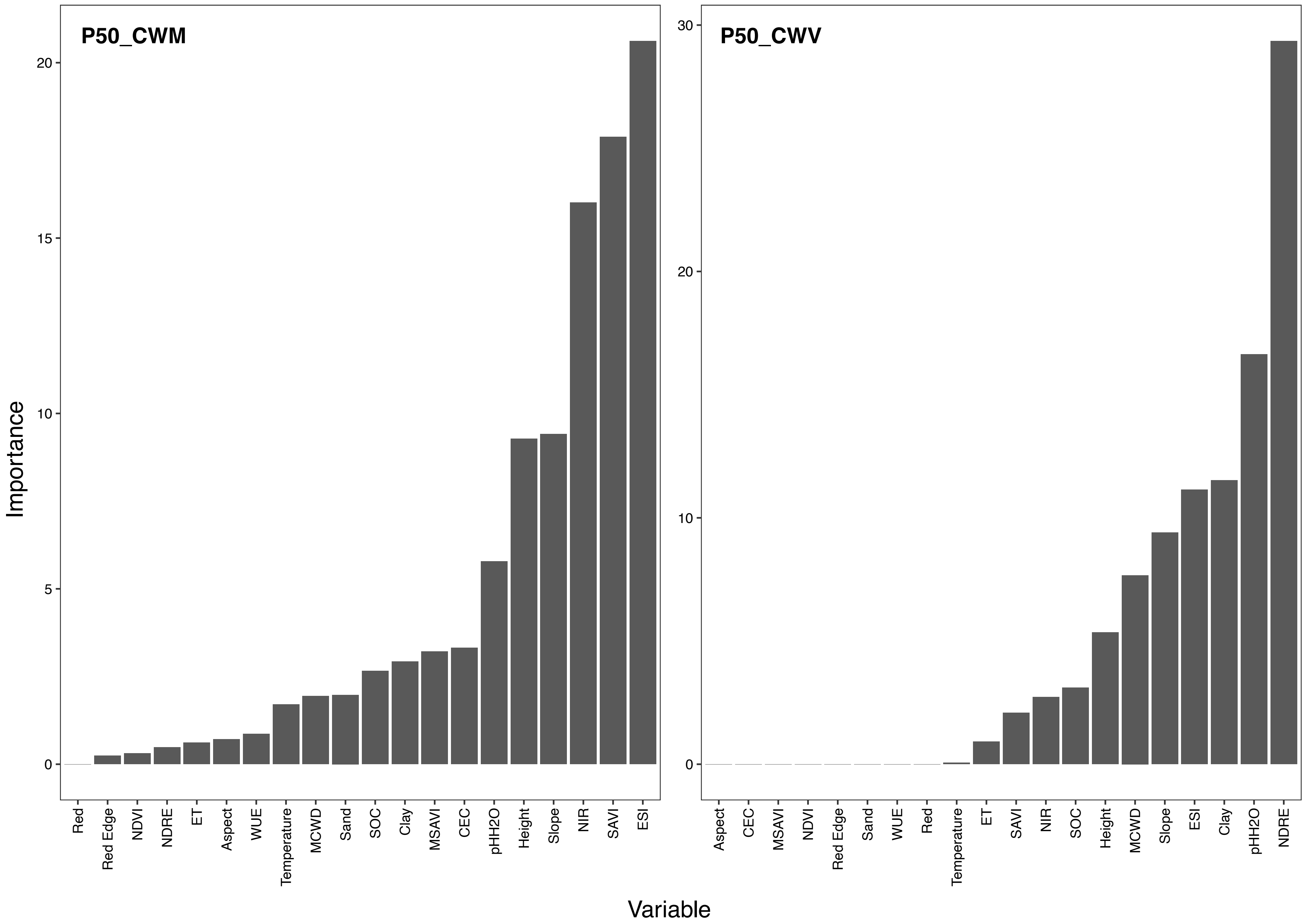


**Fig.14|Variable importance of each input band for predicting P50.**

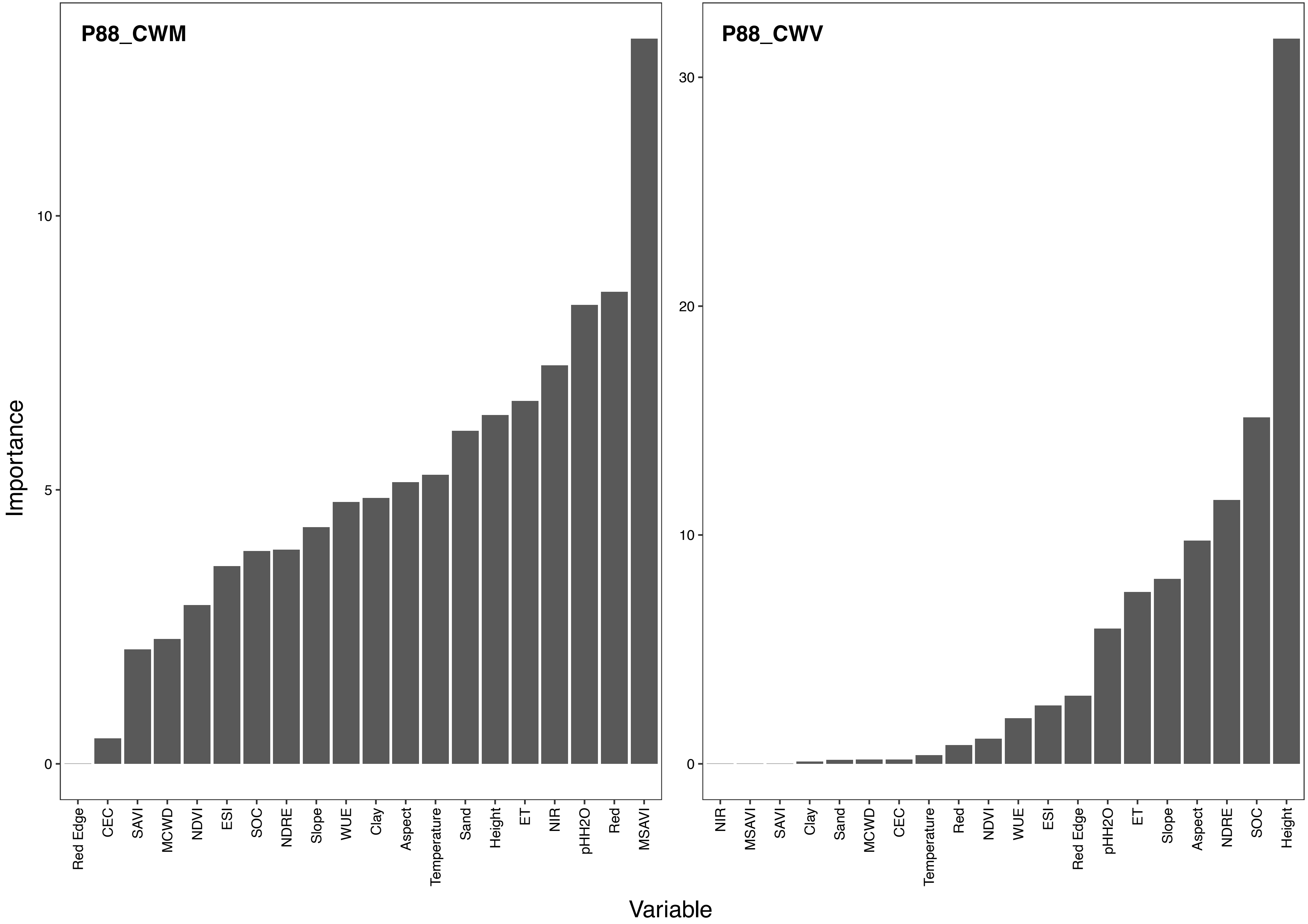


**Fig.15|Variable importance of each input band for predicting P88.**

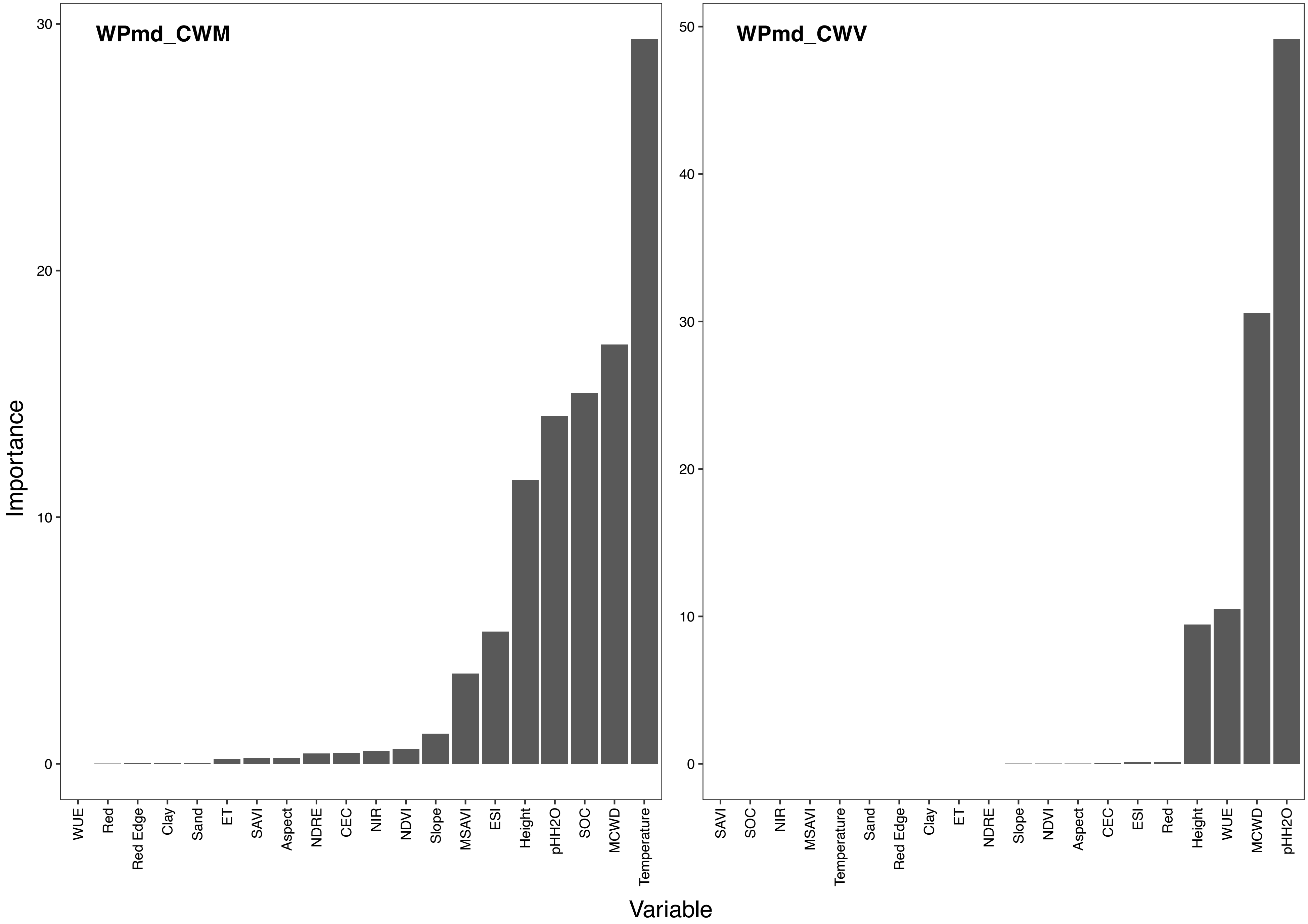


**Fig.16|Variable importance of each input band for predicting WPmd.**

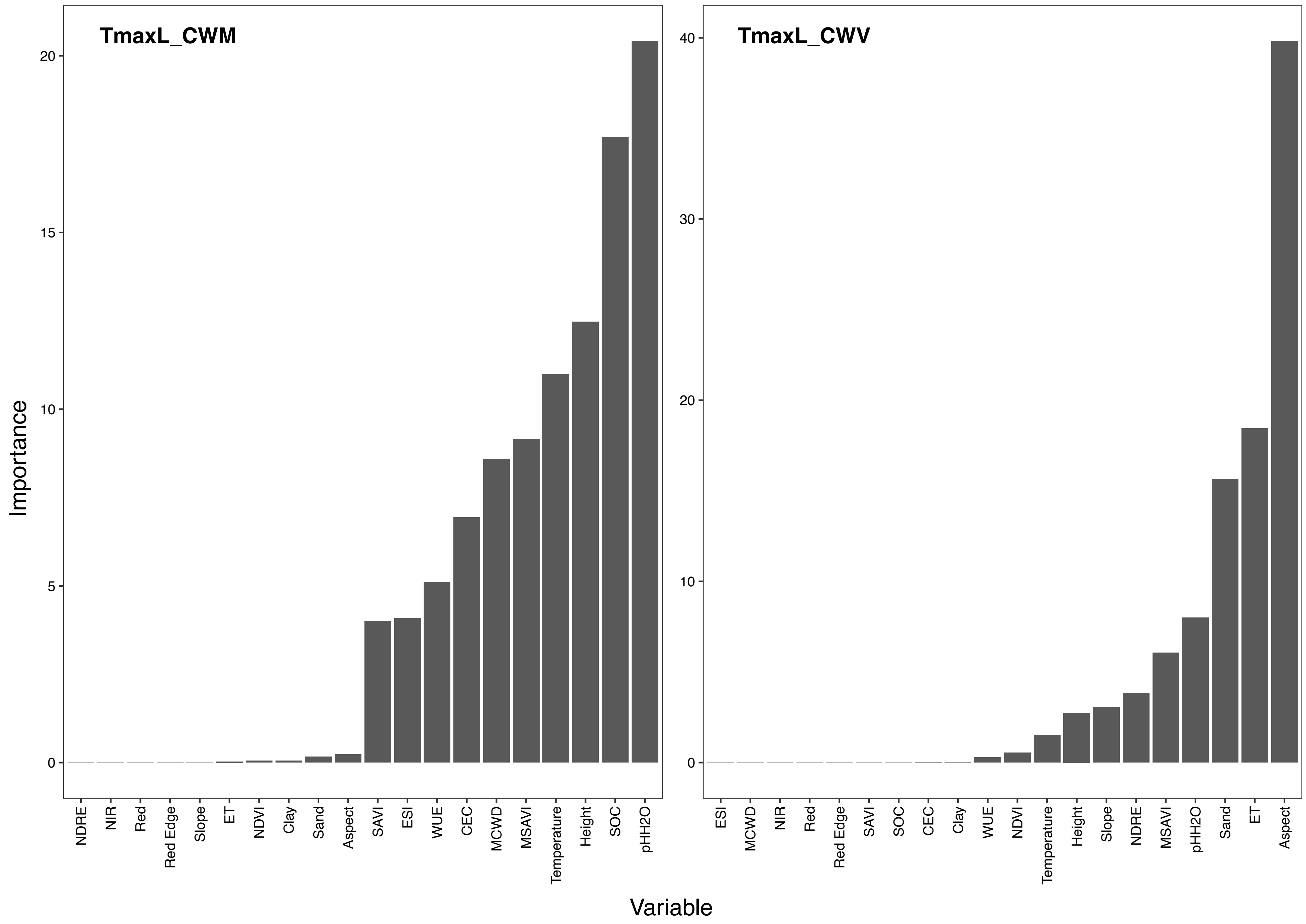


**Fig17|Variable importance of each input band for predicting TmaxL.**

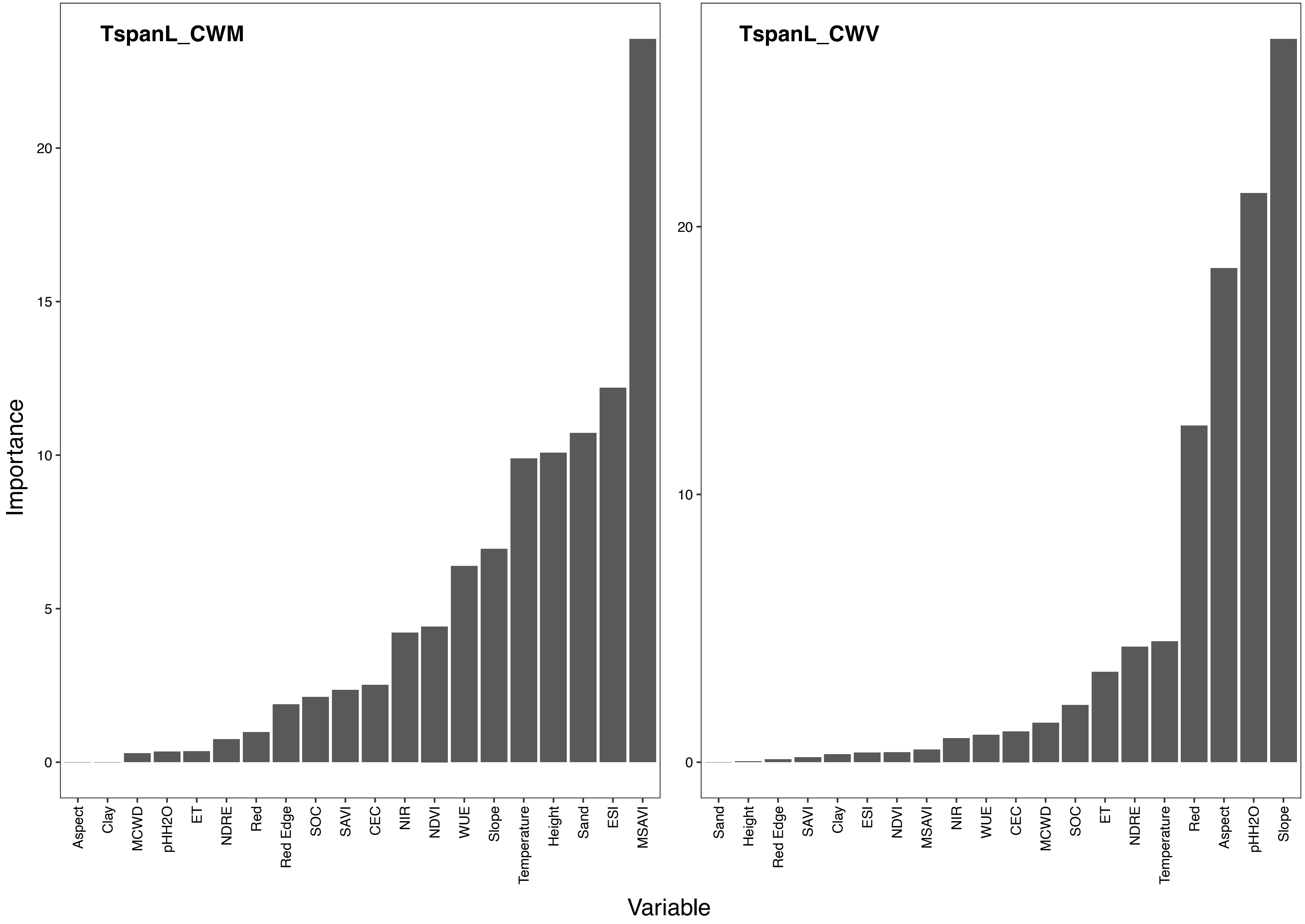


**Fig.18|Variable importance of each input band for predicting TspanL.**

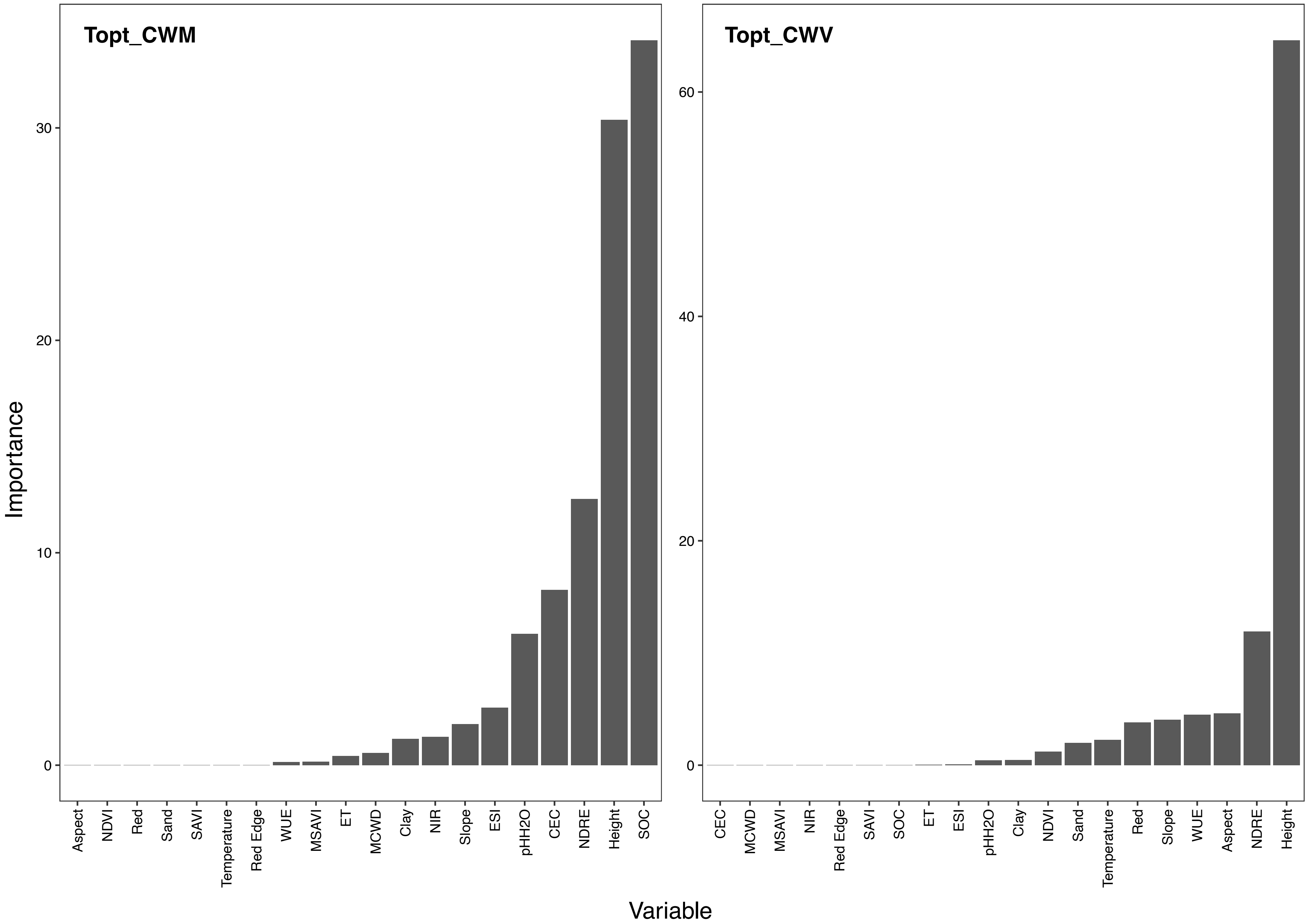


**Fig.19|Variable importance of each input band for predicting Topt.**

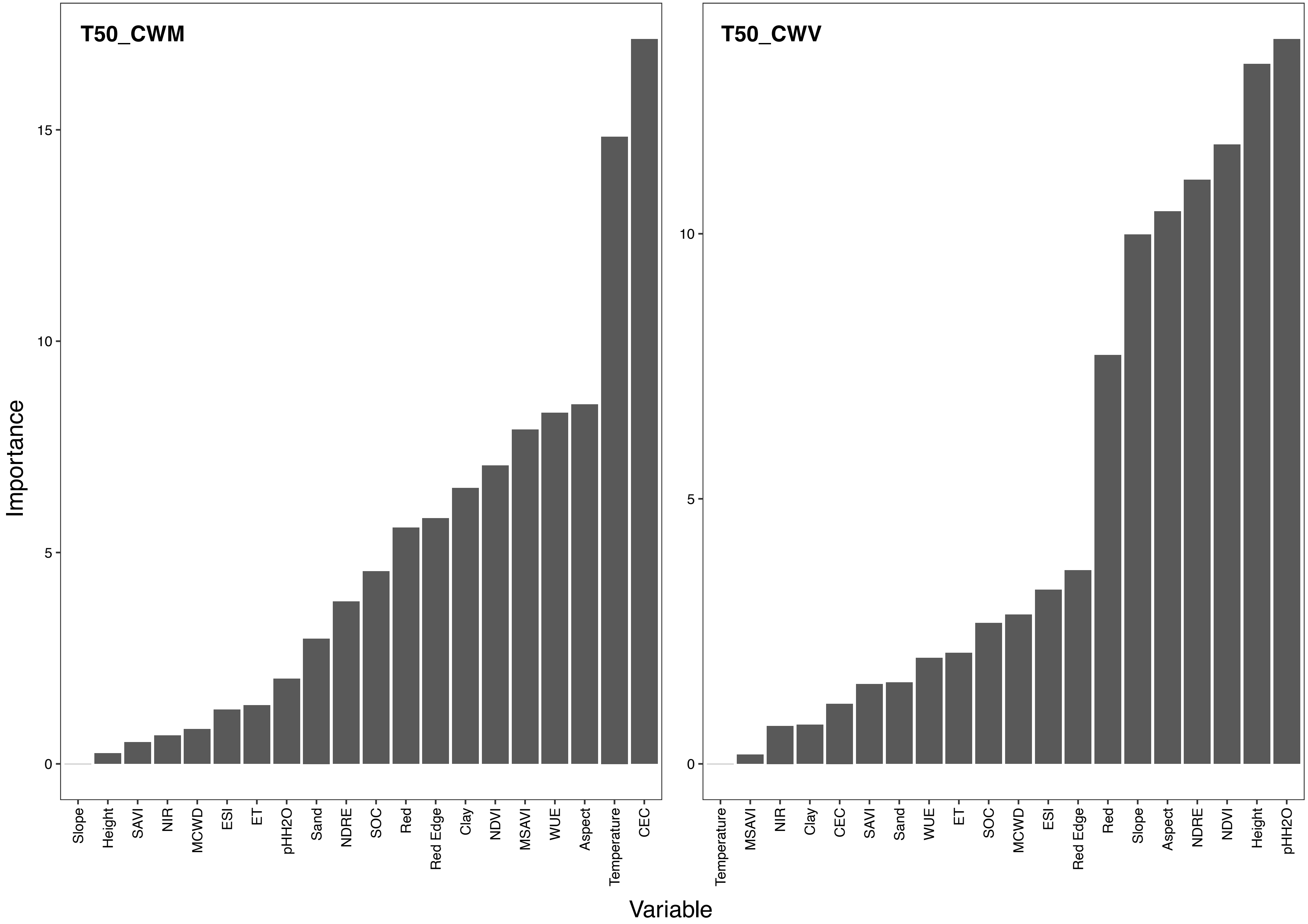


**Fig.20|Variable importance of each input band for predicting T50.**

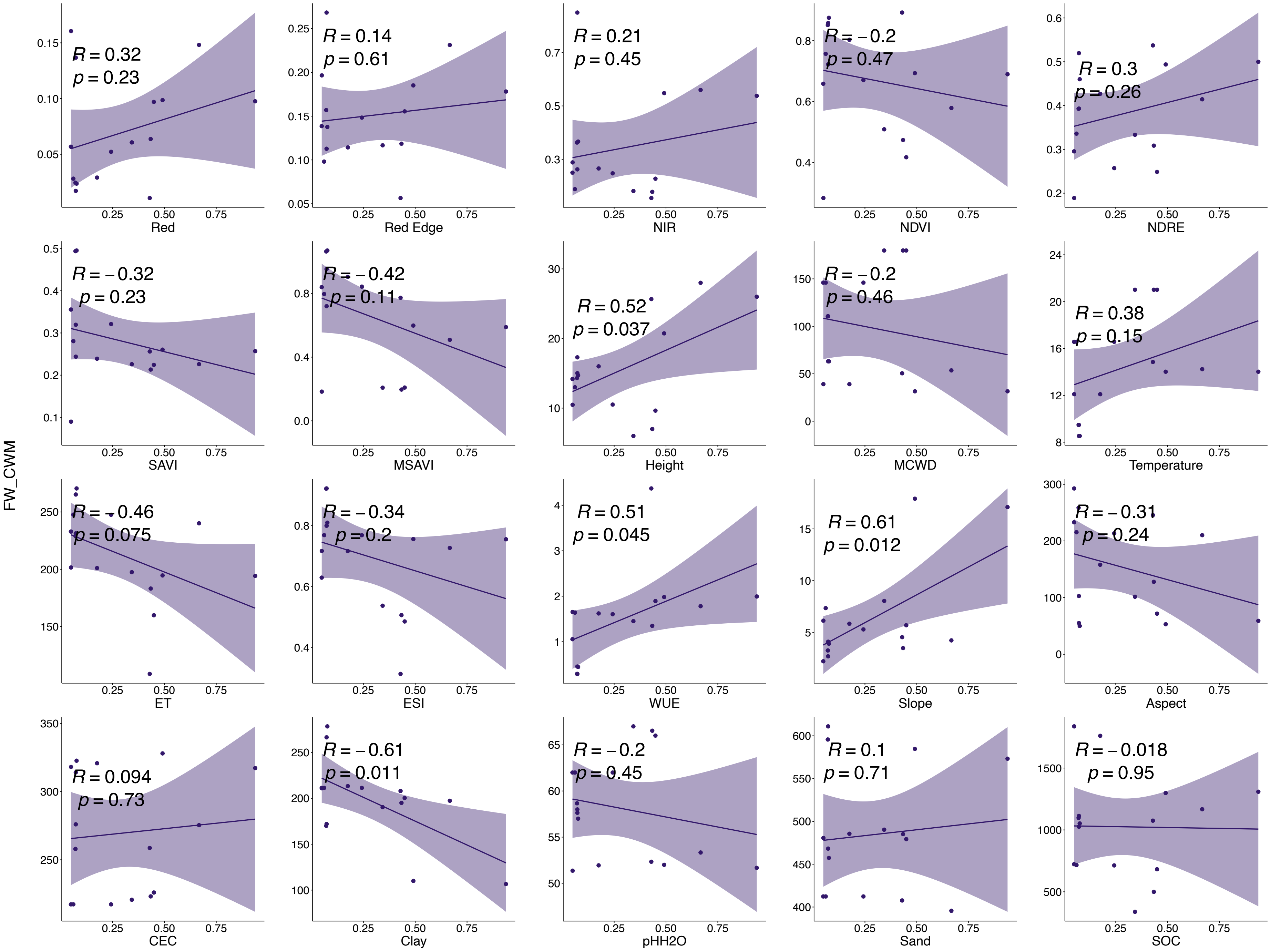


**Fig.21|Relationships between CWM of FW and mean values of each input band.**

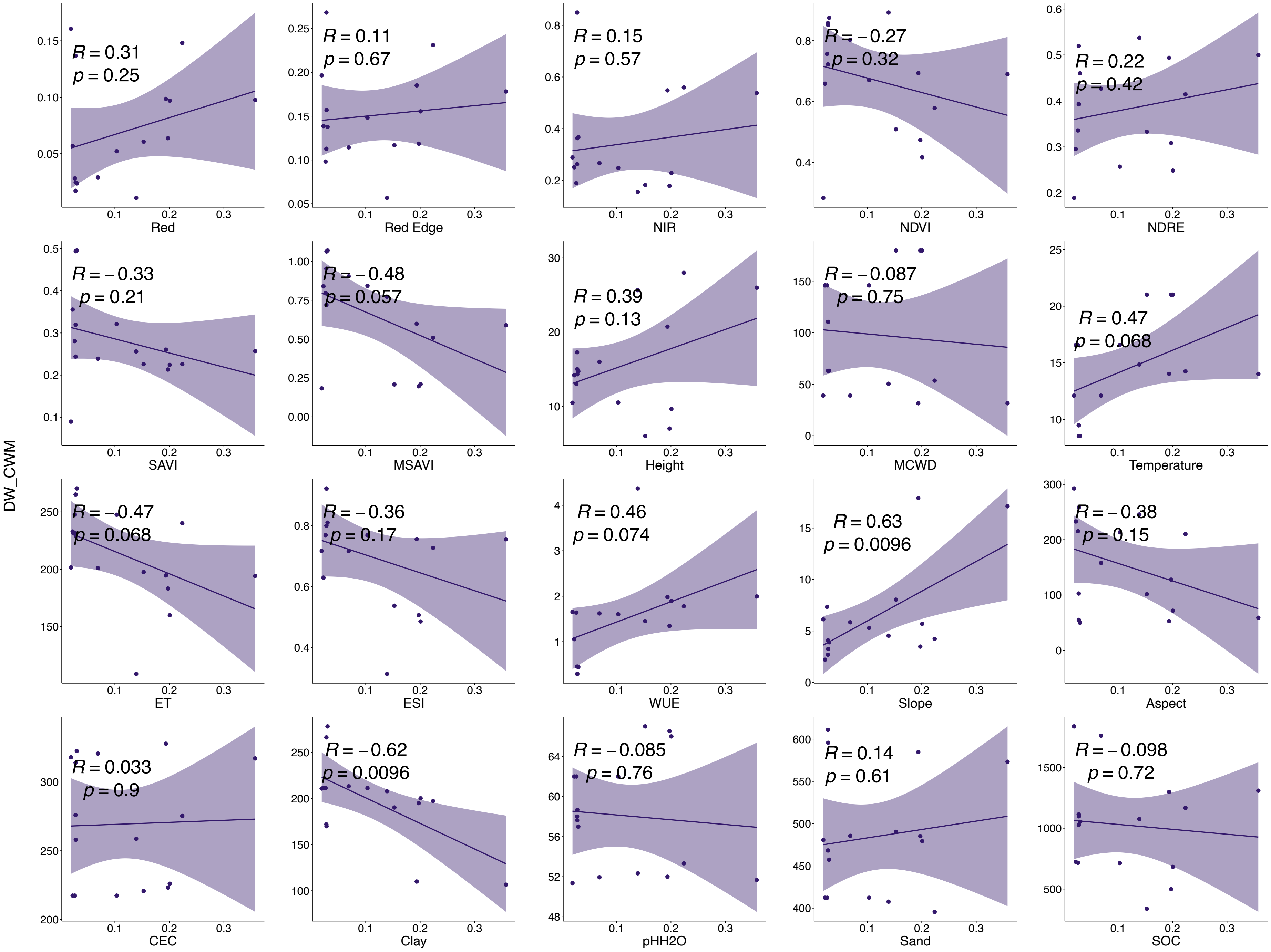


**Fig.22|Relationships between CWM of DW and mean values of each input band.**

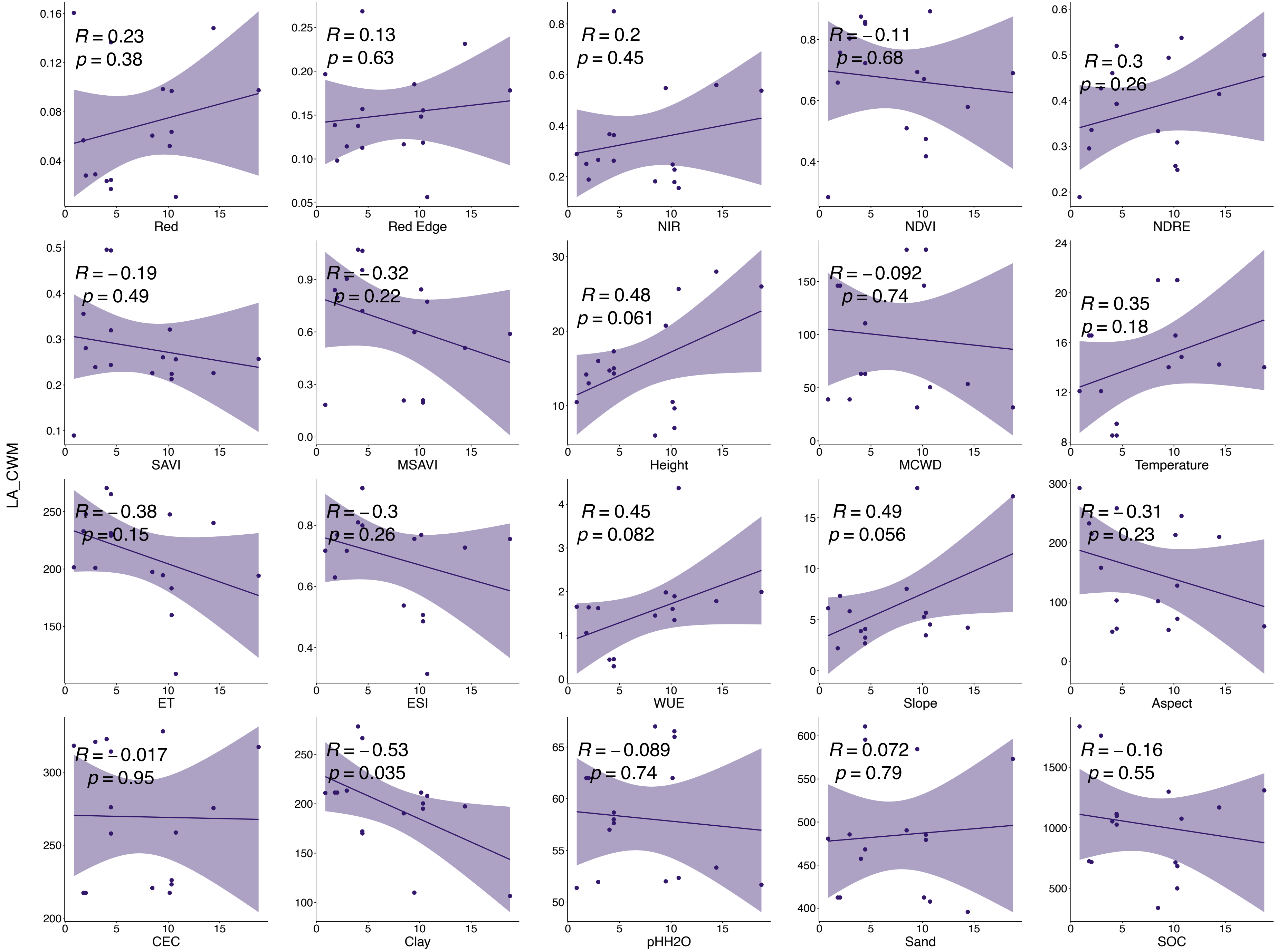


**Fig.23|Relationships between CWM of LA and mean values of each input band.**

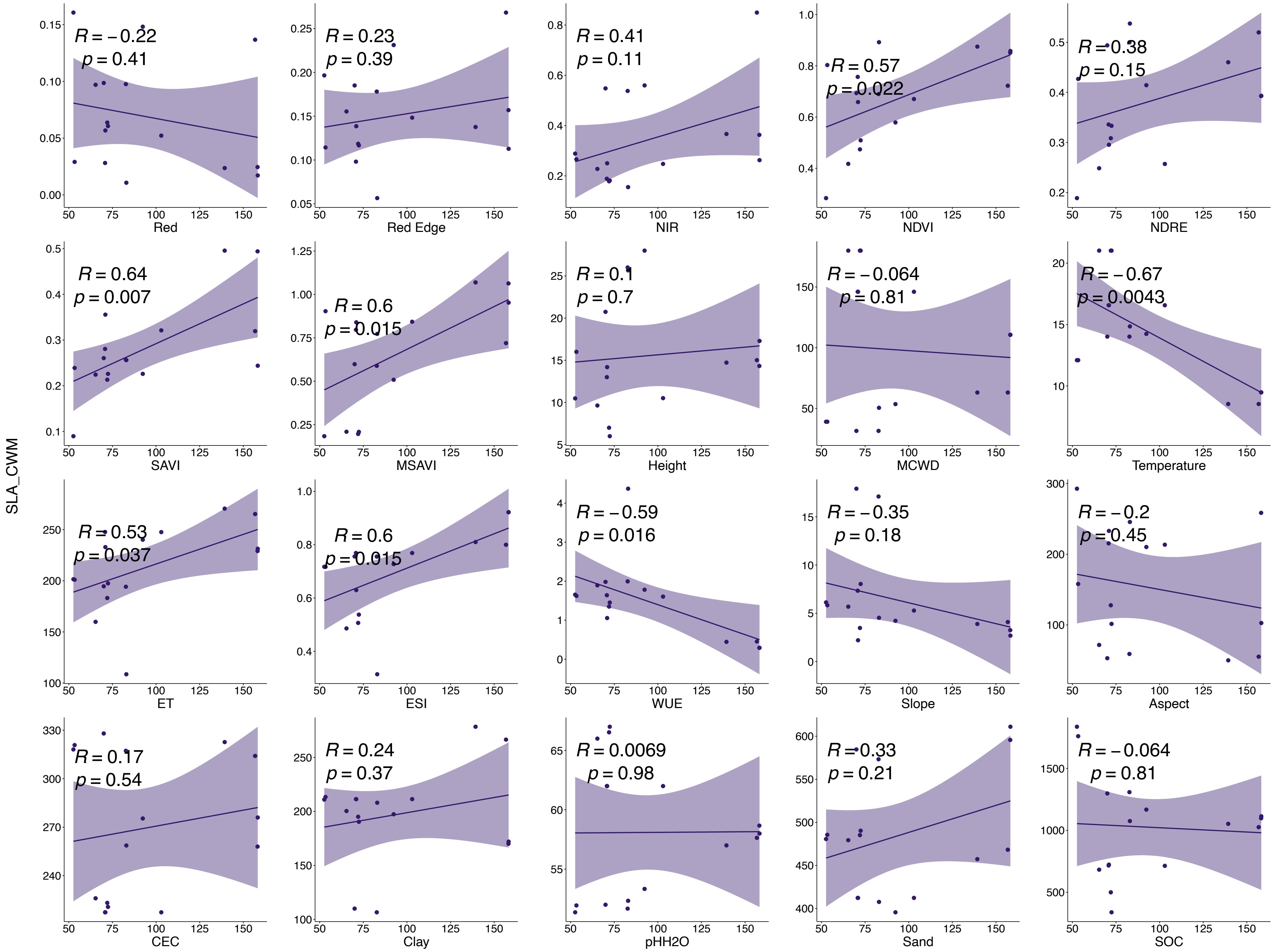


**Fig.24|Relationships between CWM of SLA and mean values of each input band.**

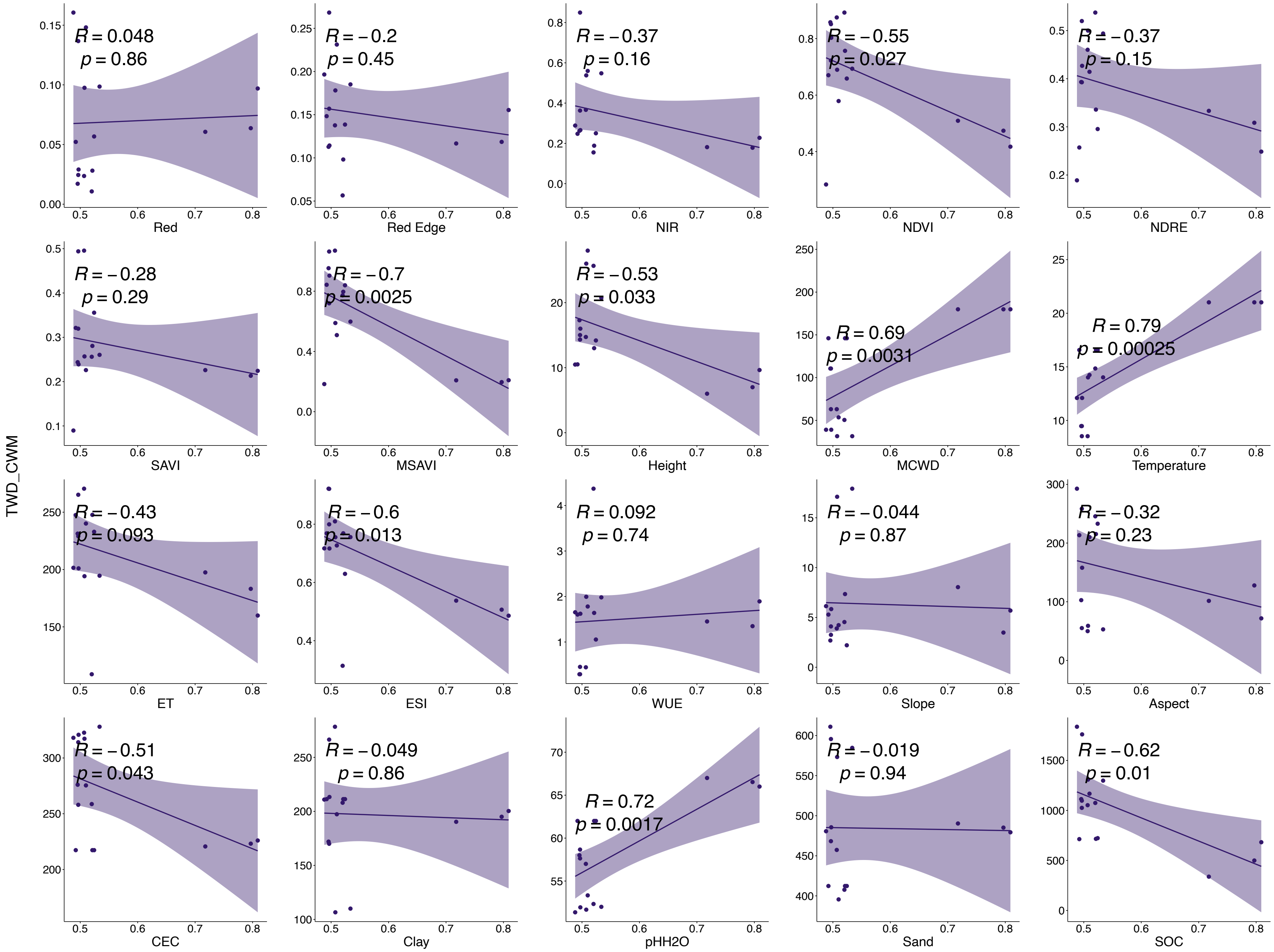


**Fig.25|Relationships between CWM of TWD and mean values of each input band.**

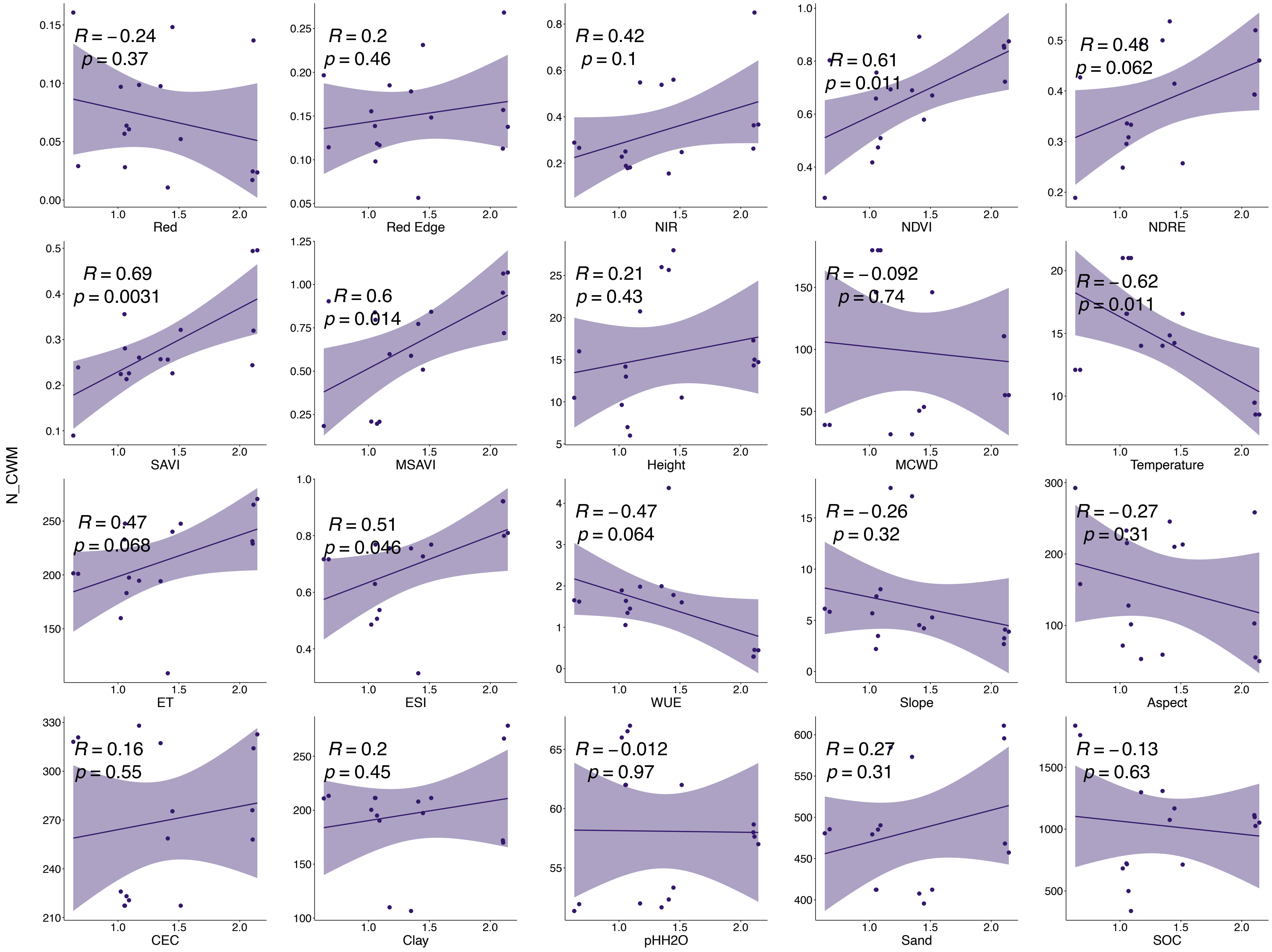


**Fig.26|Relationships between CWM of N and mean values of each input band.**

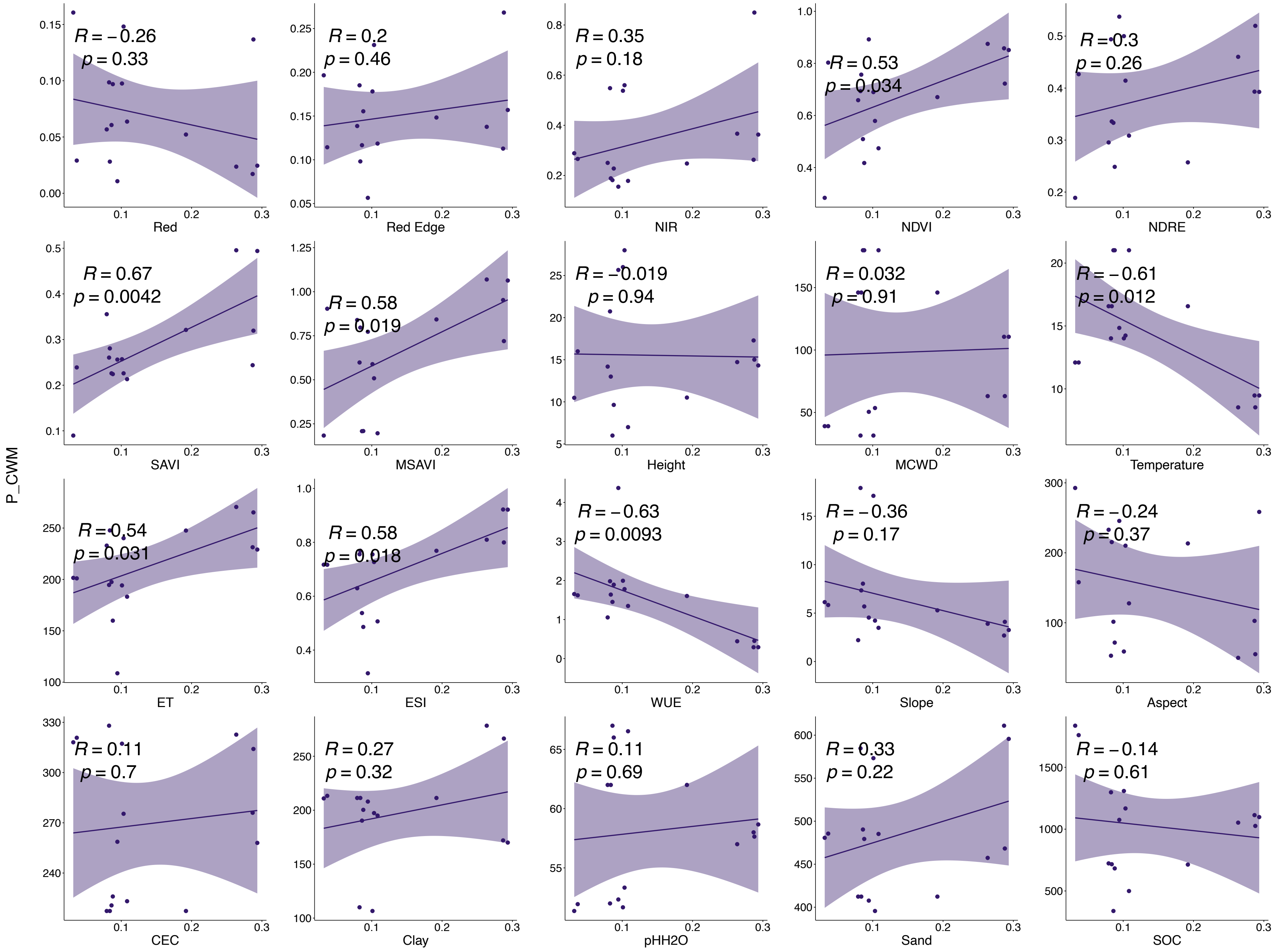


**Fig.27|Relationships between CWM of P and mean values of each input band.**

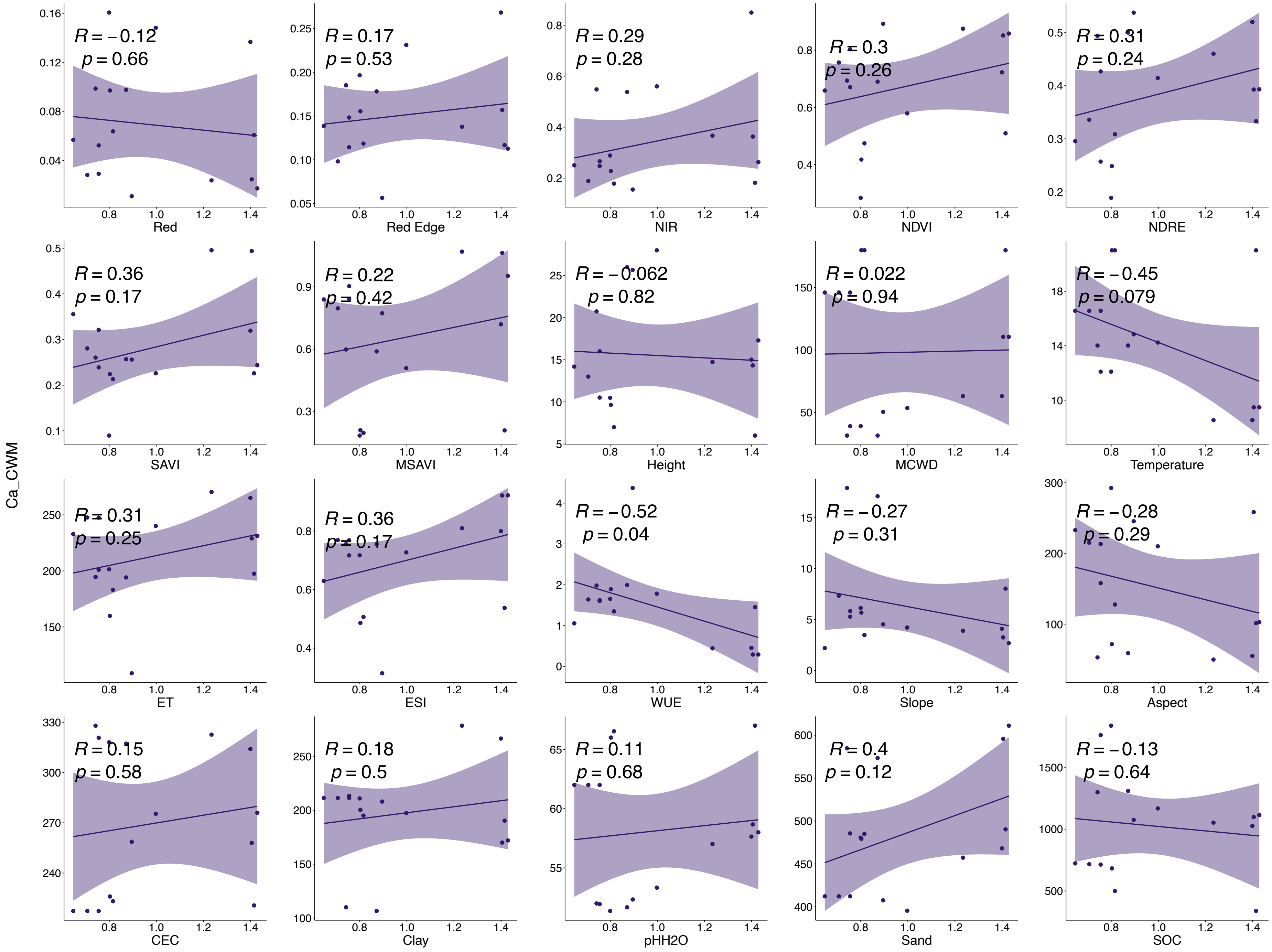


**Fig.28|Relationships between CWM of Ca and mean values of each input band.**

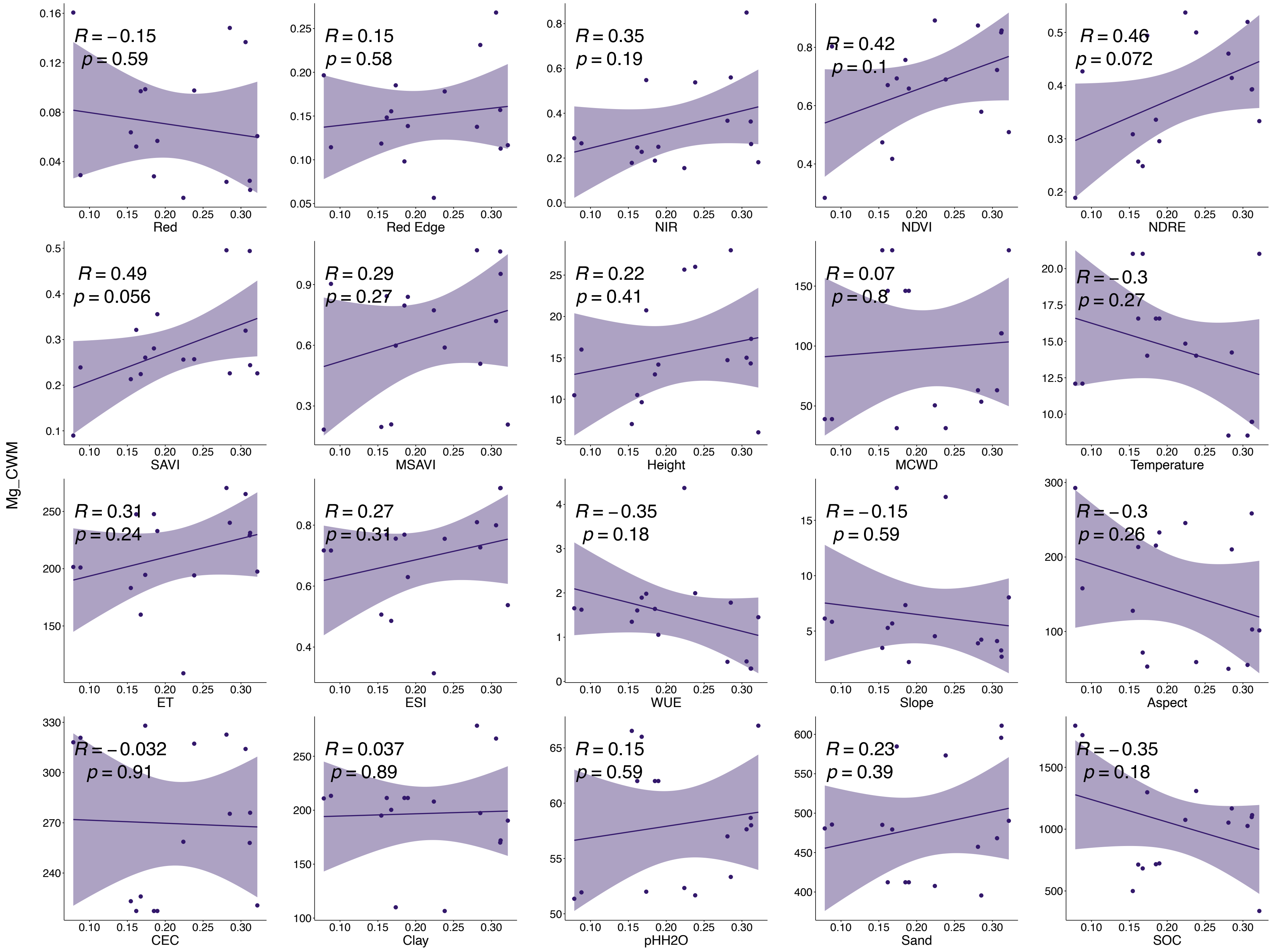


**Fig.29|Relationships between CWM of Mg and mean values of each input band.**

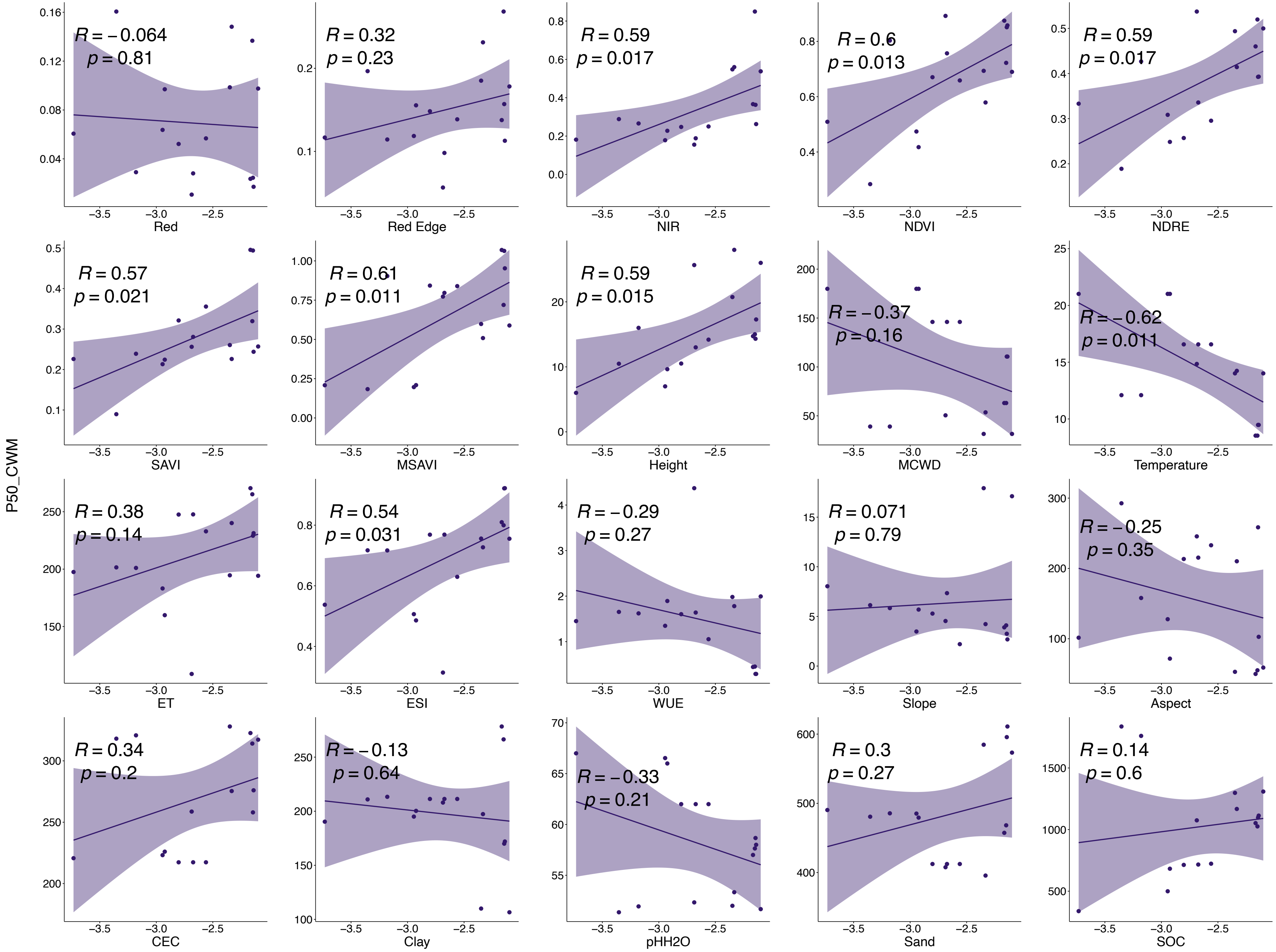


**Fig.30|Relationships between CWM of P50 and mean values of each input band.**

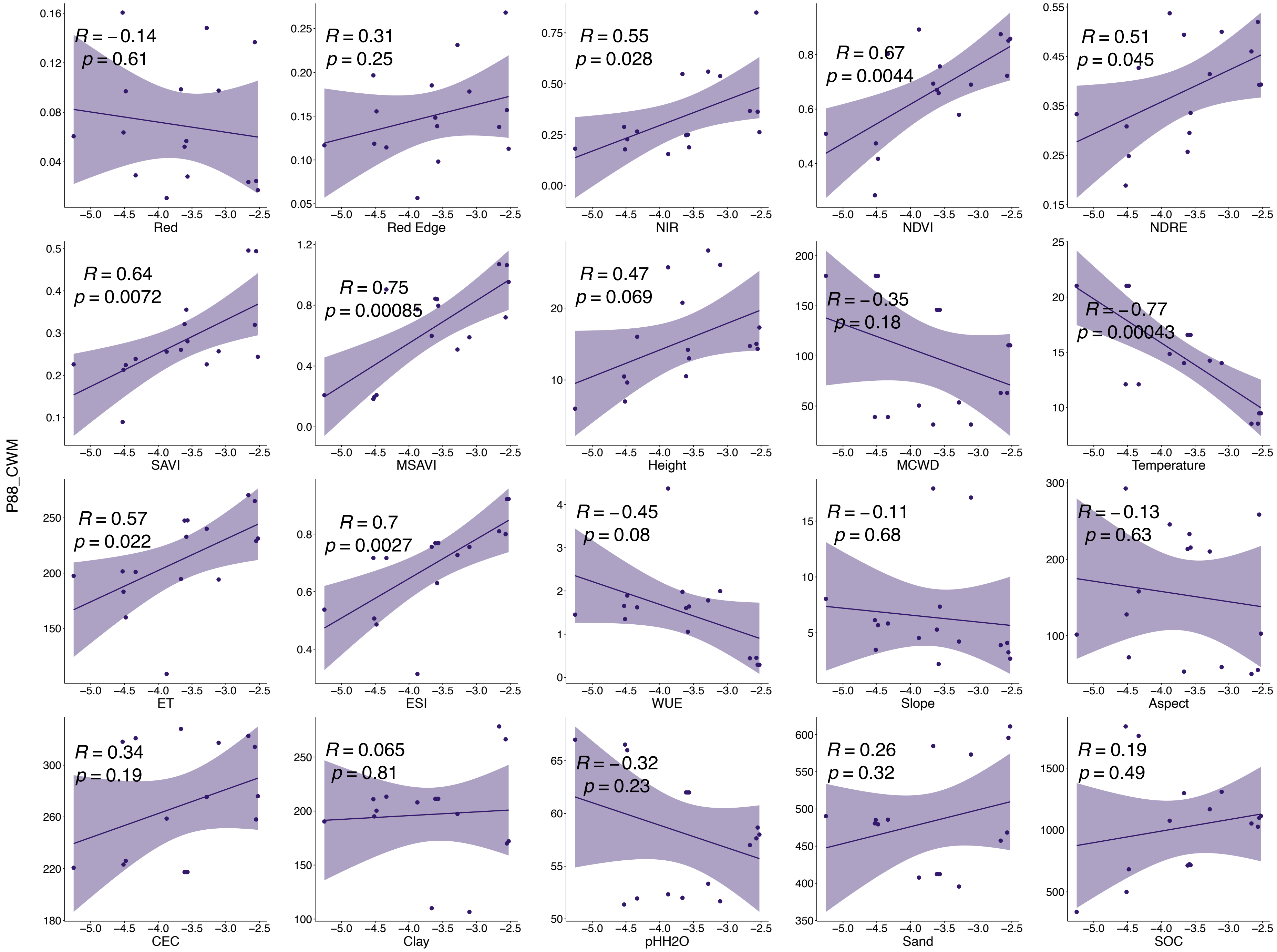


**Fig.31|Relationships between CWM of P88 and mean values of each input band.**

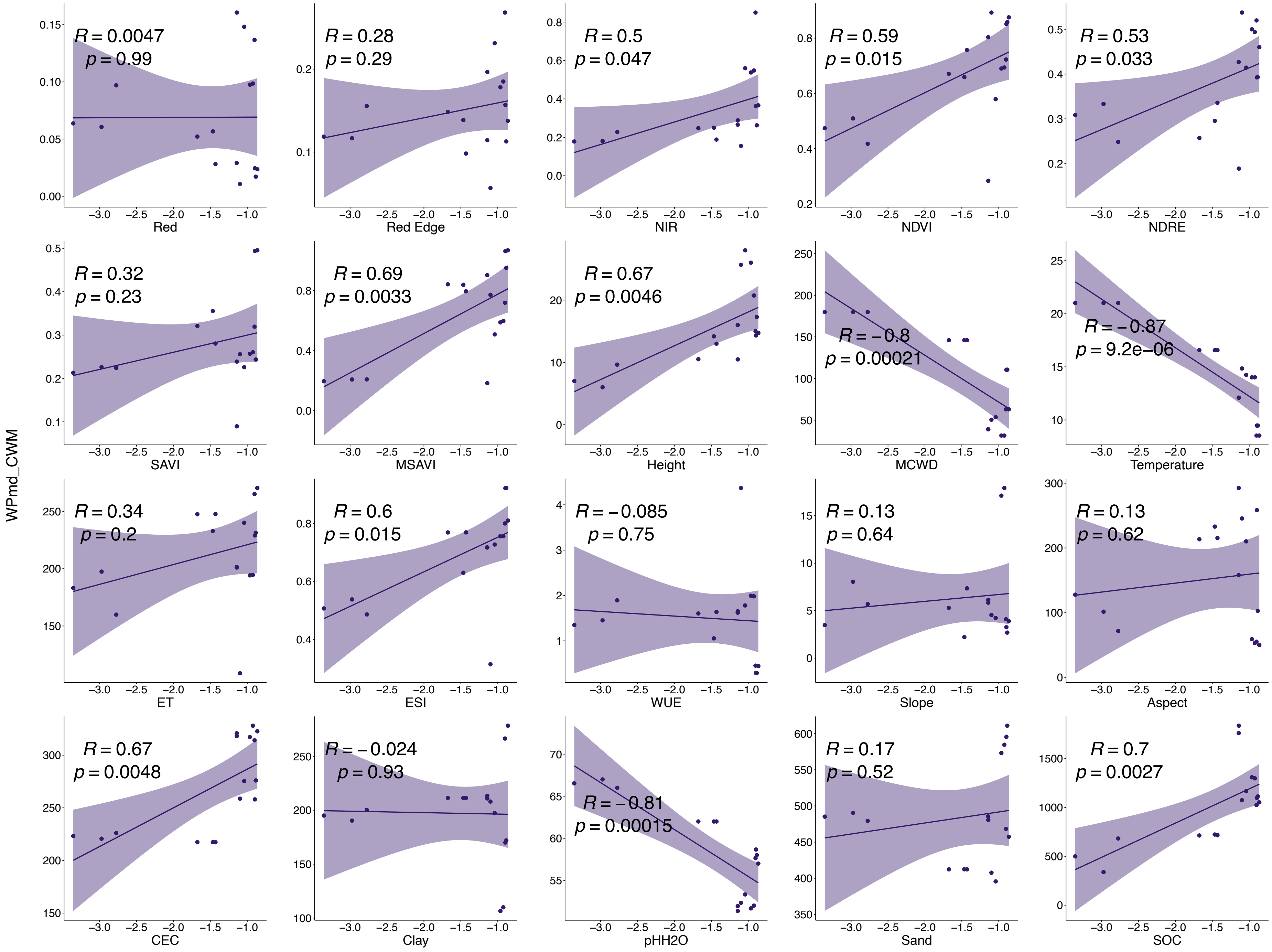


**Fig.32|Relationships between CWM of WPmd and mean values of each input band.**

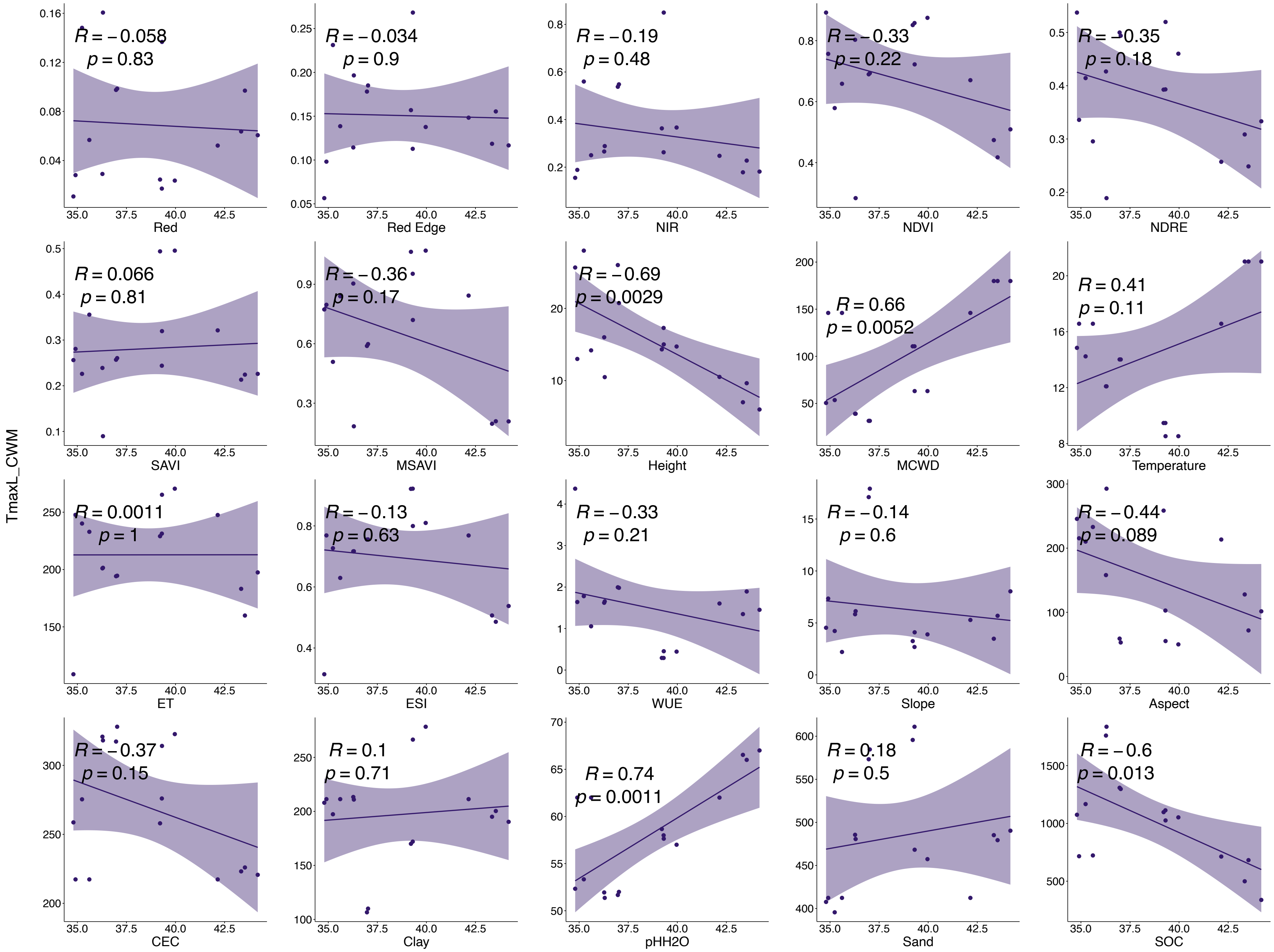


**Fig.33|Relationships between CWM of TmaxL and mean values of each input band.**

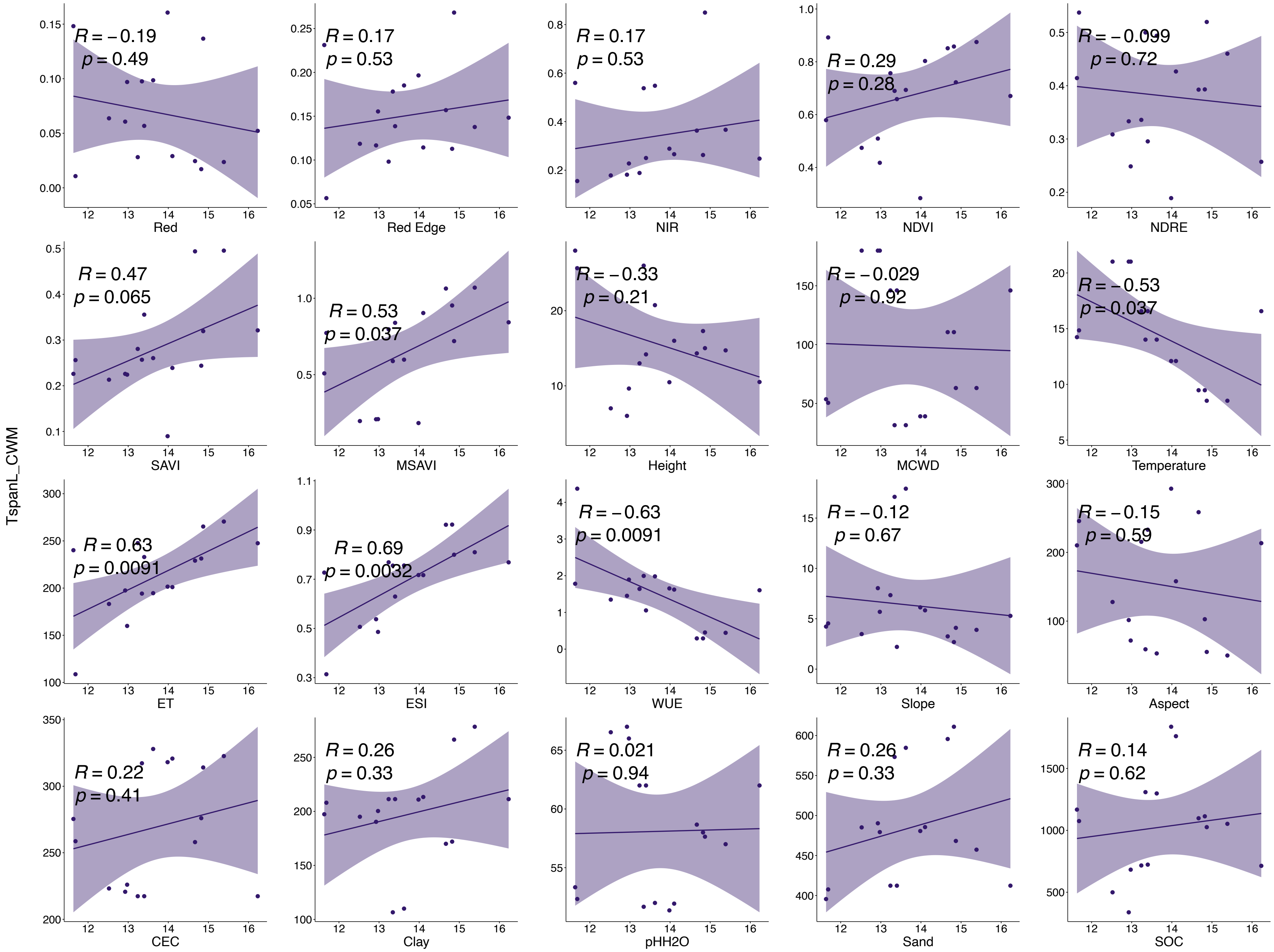


**Fig.34|Relationships between CWM of TspanL and mean values of each input band.**

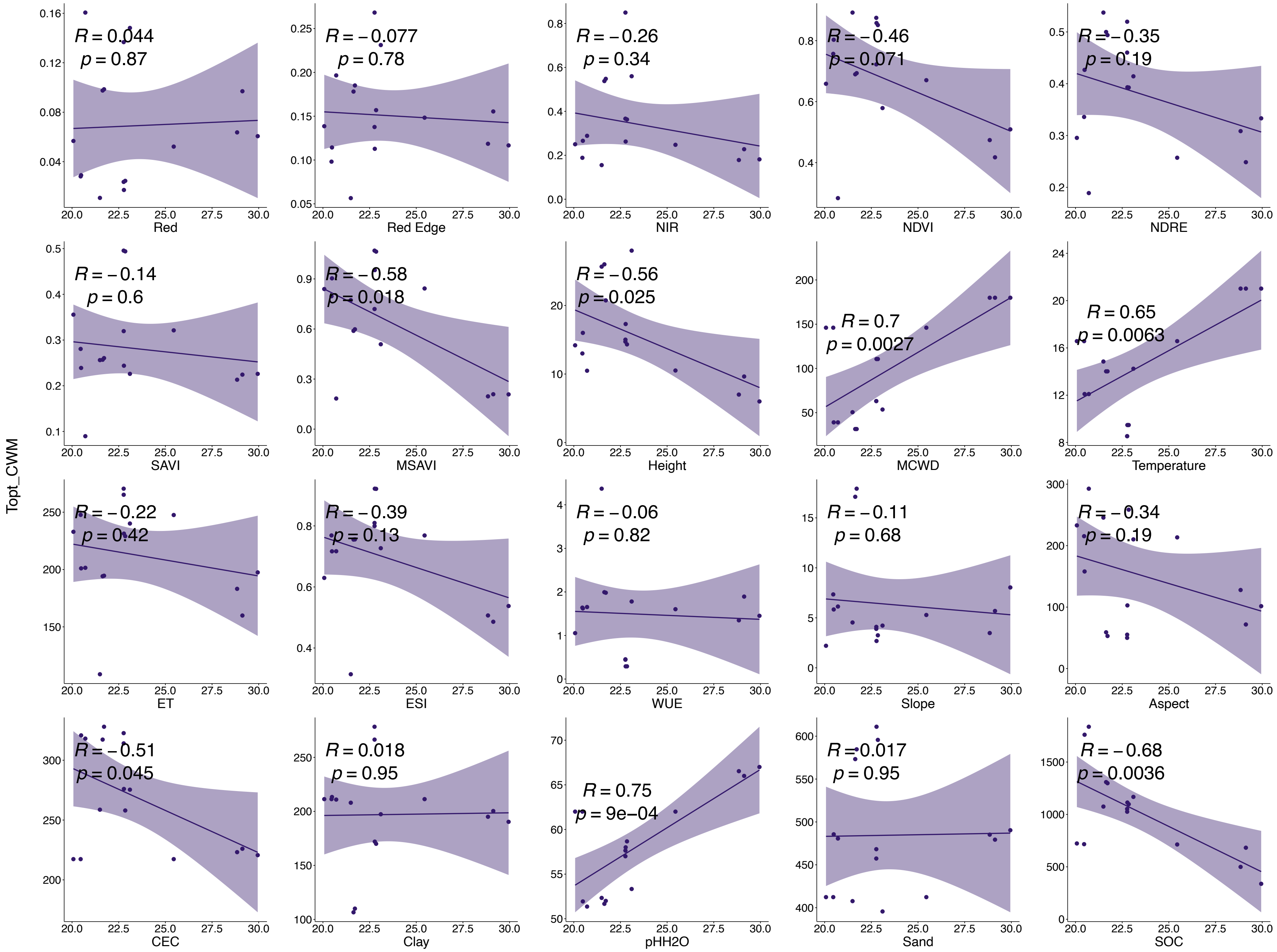


**Fig.35|Relationships between CWM of Topt and mean values of each input band.**

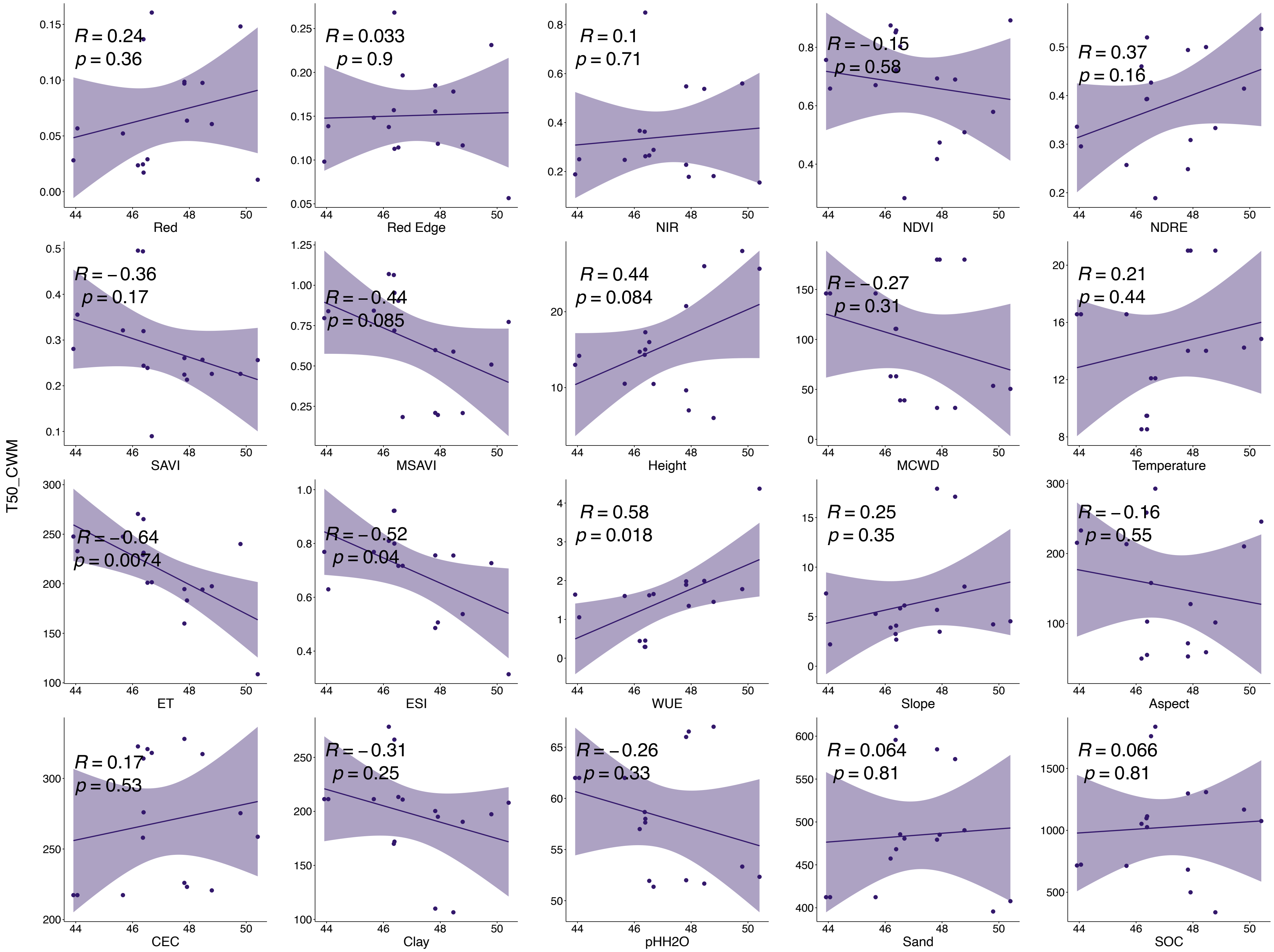


**Fig.36|Relationships between CWM of T50 and mean values of each input band.**

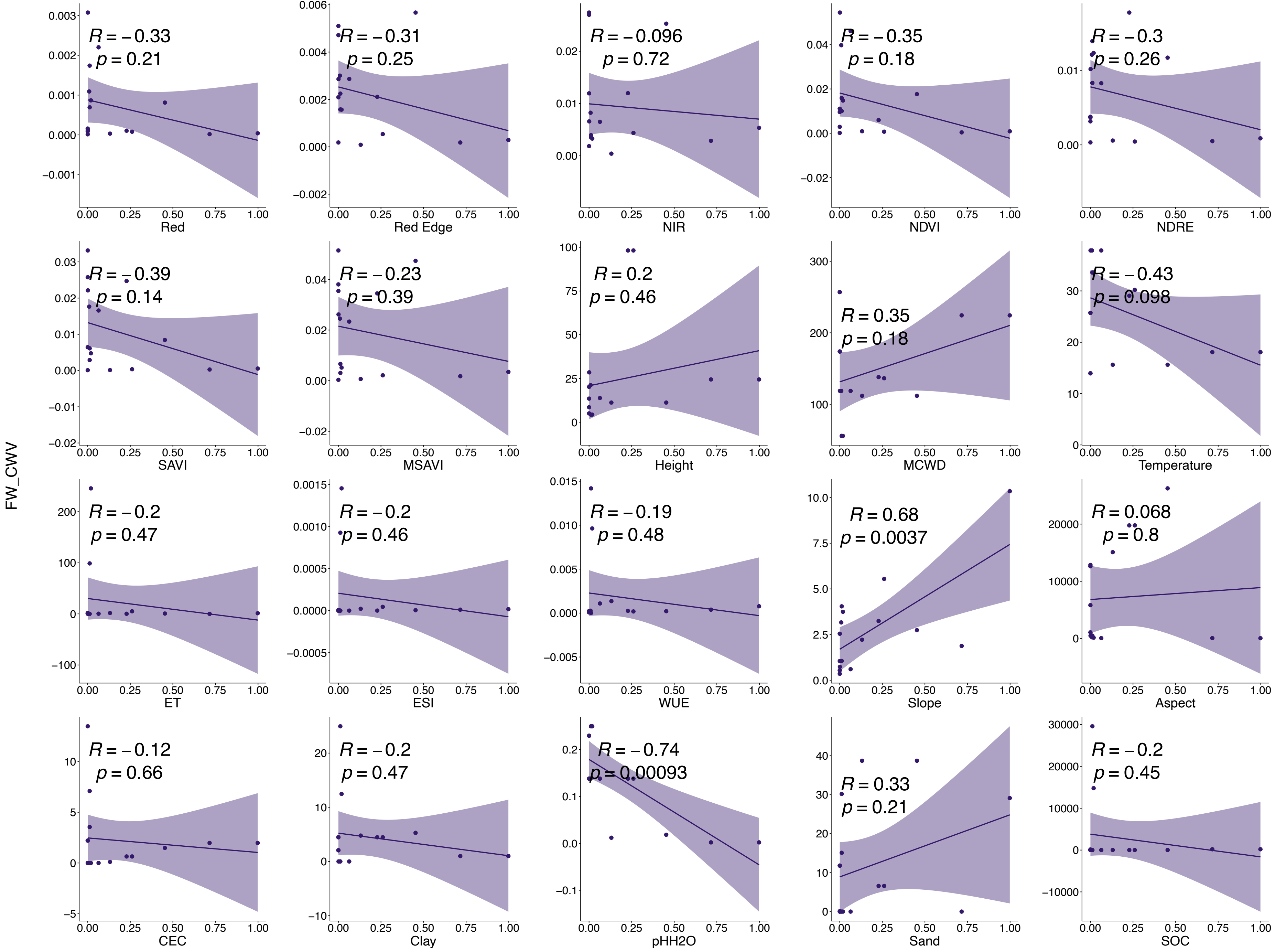


**Fig.37|Relationships between CWV of FW and variance values of each input band.**

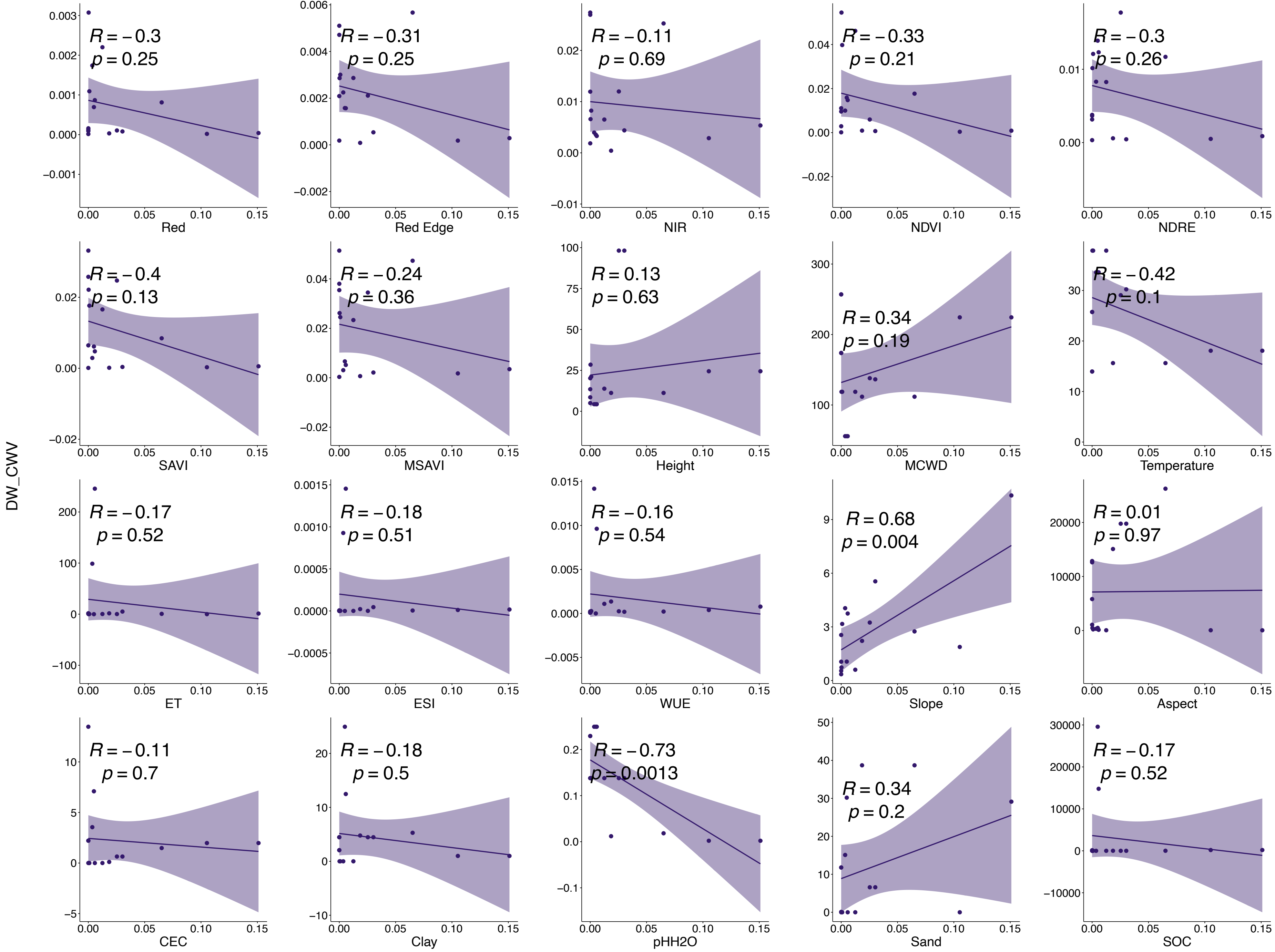


**Fig.38|Relationships between CWV of DW and variance values of each input band.**

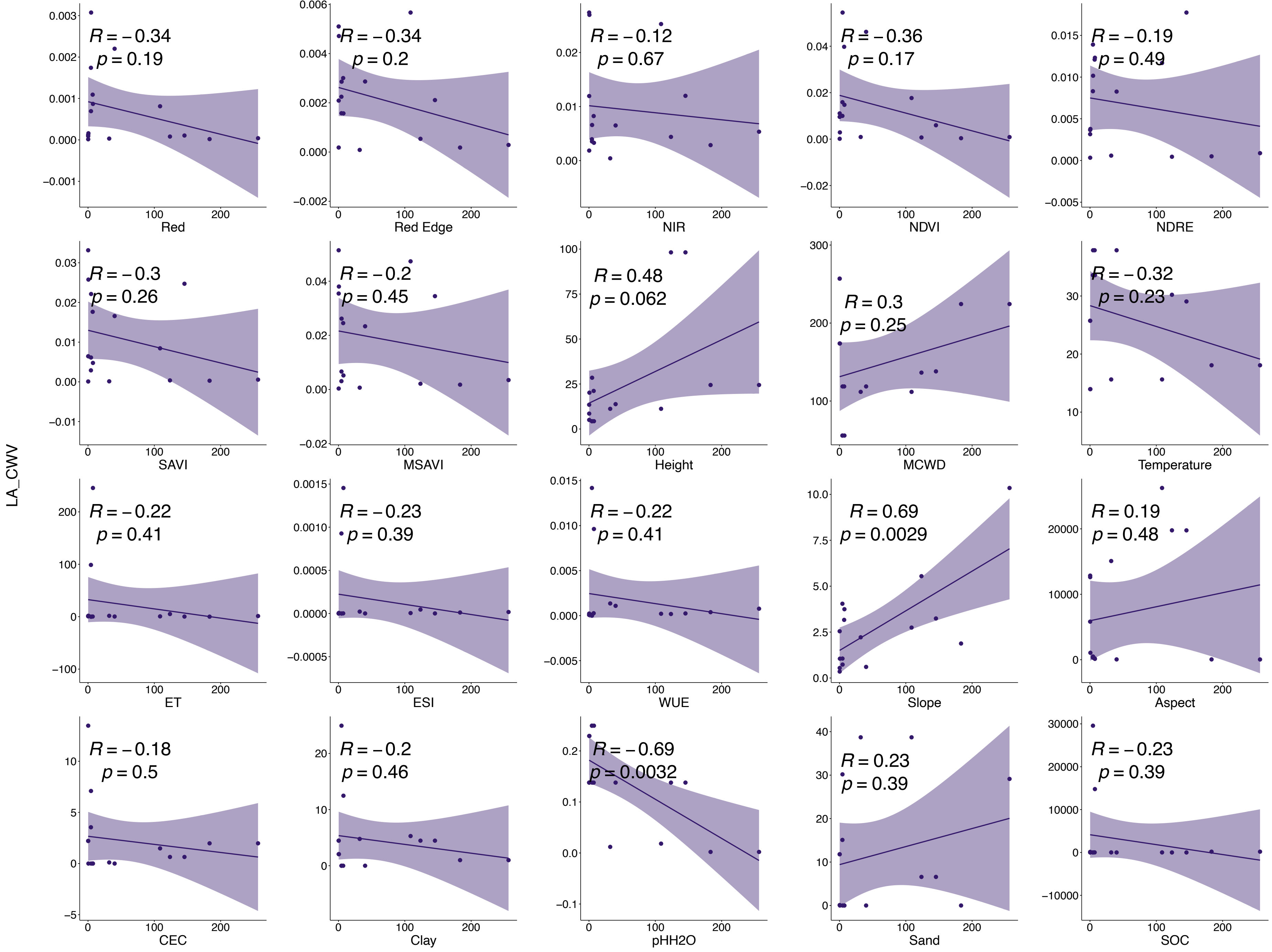


**Fig.39|Relationships between CWV of LA and variance values of each input band.**

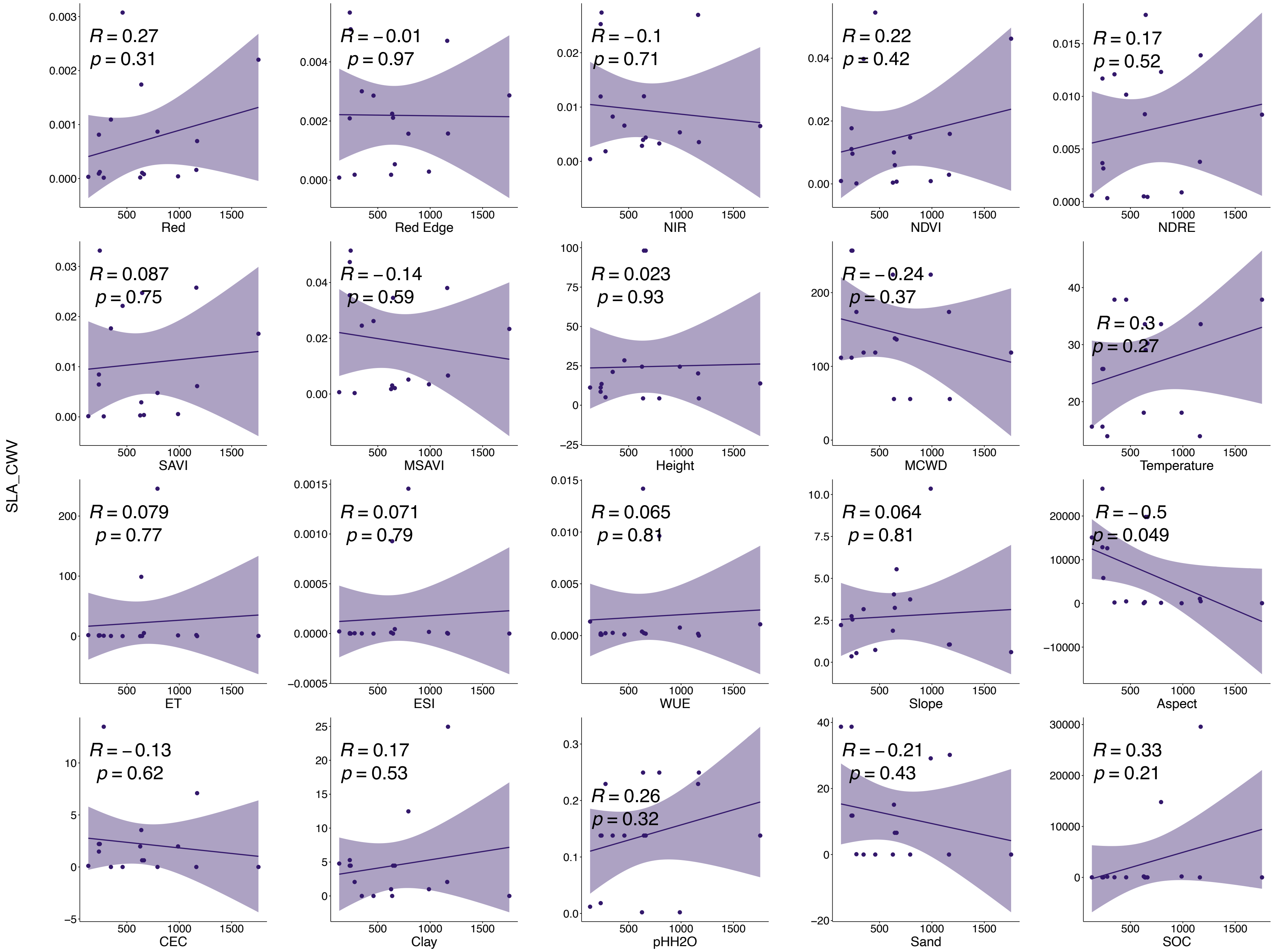


**Fig.40|Relationships between CWV of SLA and variance values of each input band.**

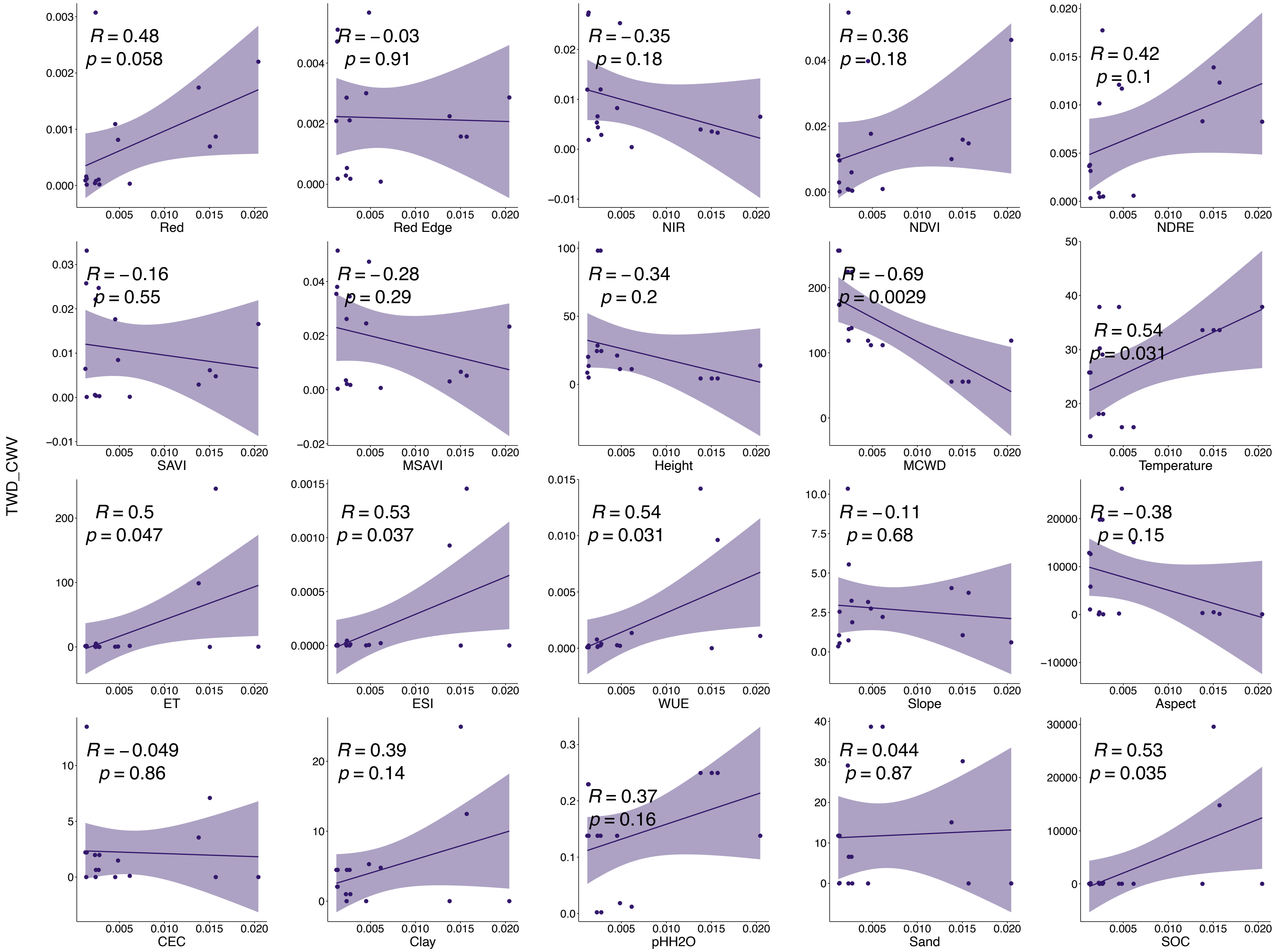


**Fig.41|Relationships between CWV of TWD and variance values of each input band.**

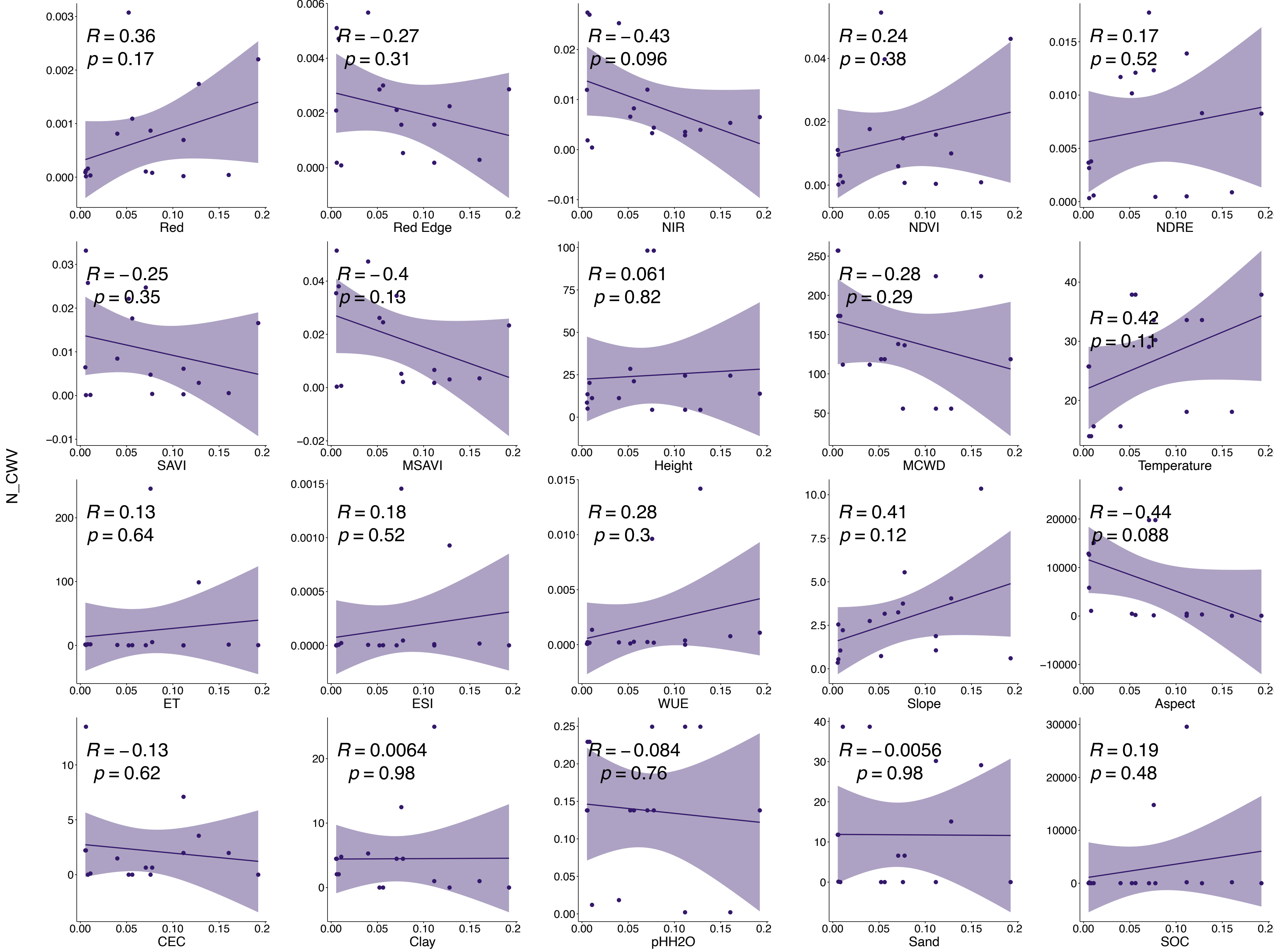


**Fig.42|Relationships between CWV of N and variance values of each input band.**

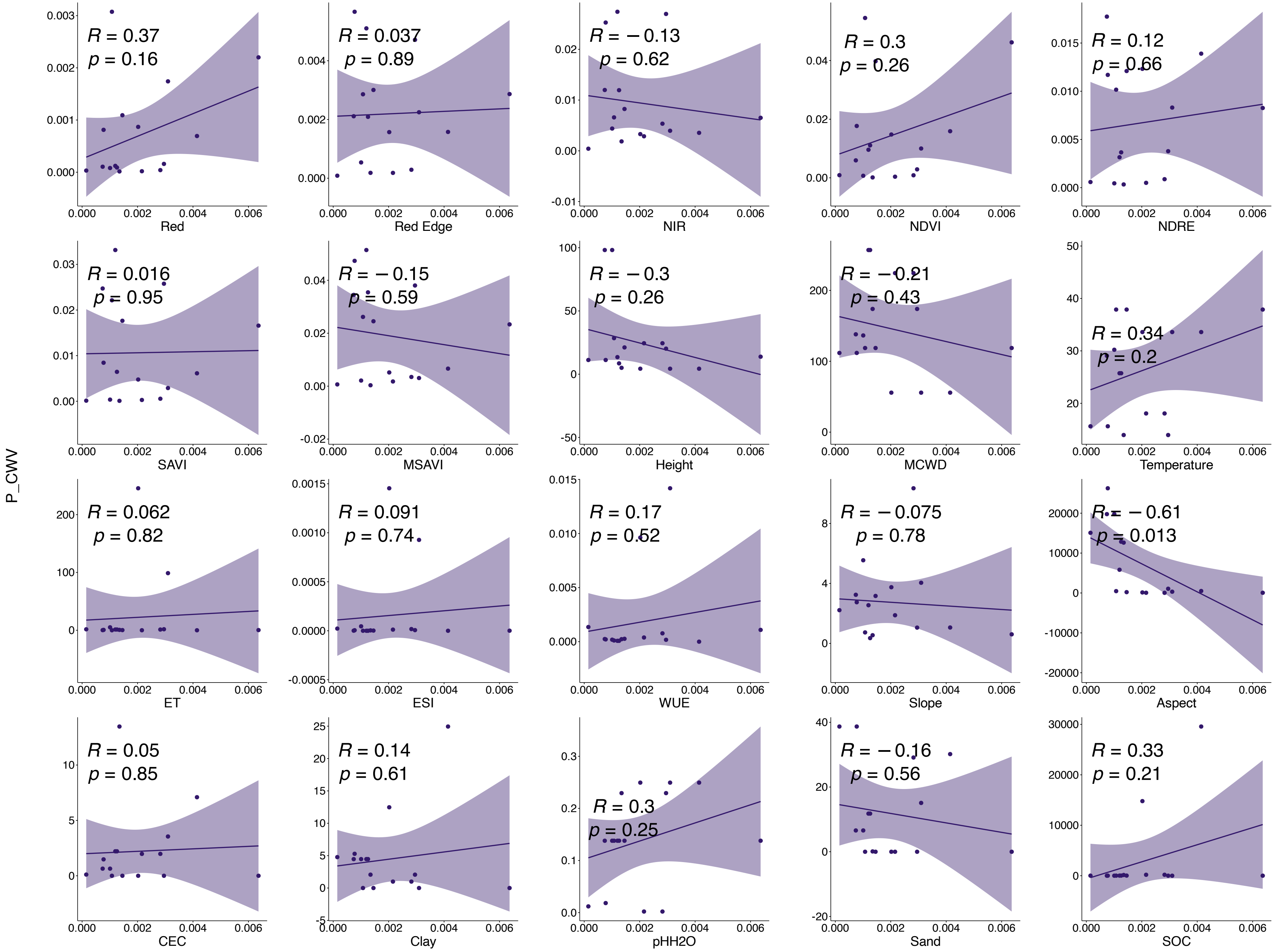


**Fig.43|Relationships between CWV of P and variance values of each input band.**

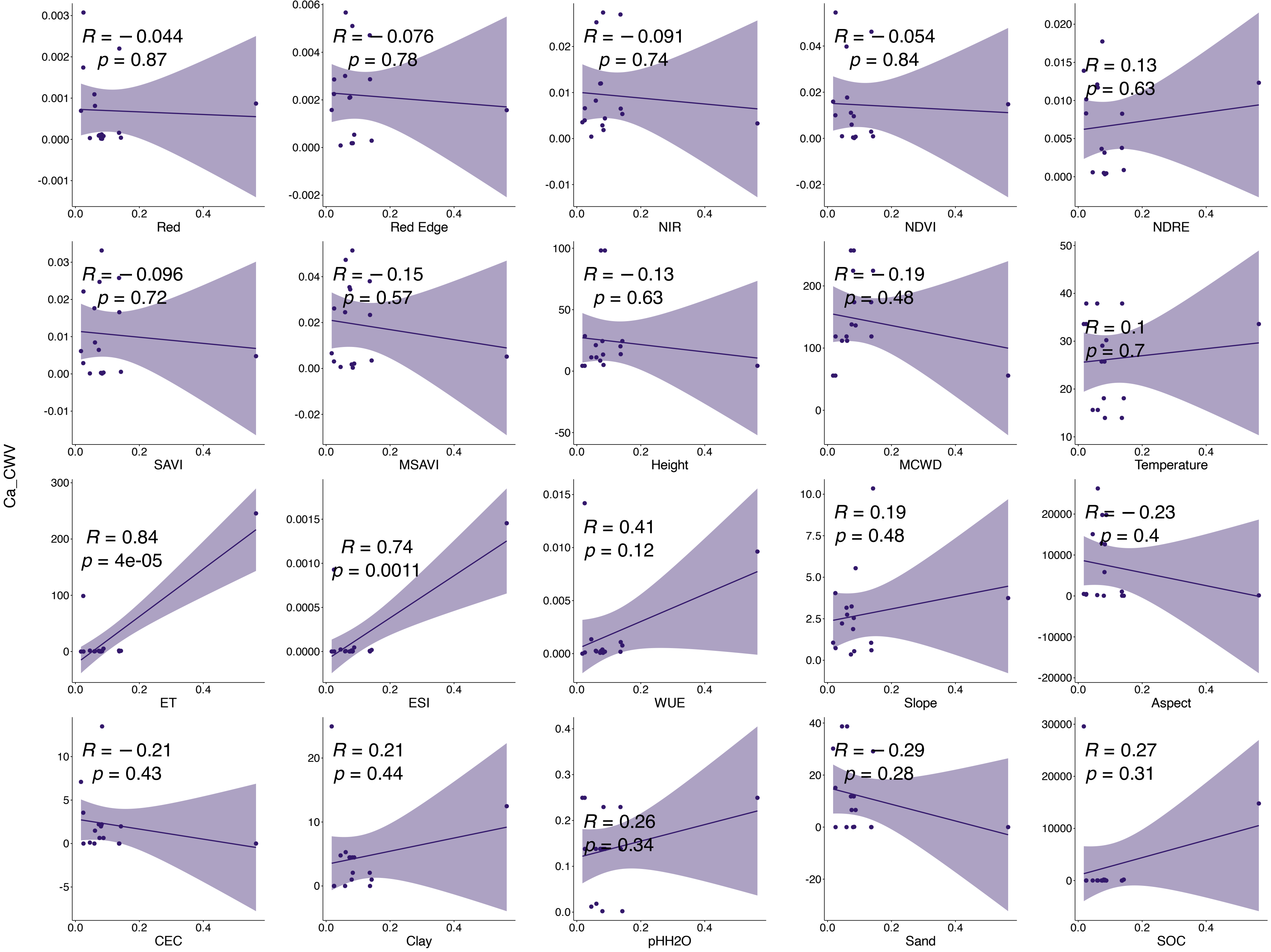


**Fig.44|Relationships between CWV of Ca and variance values of each input band.**

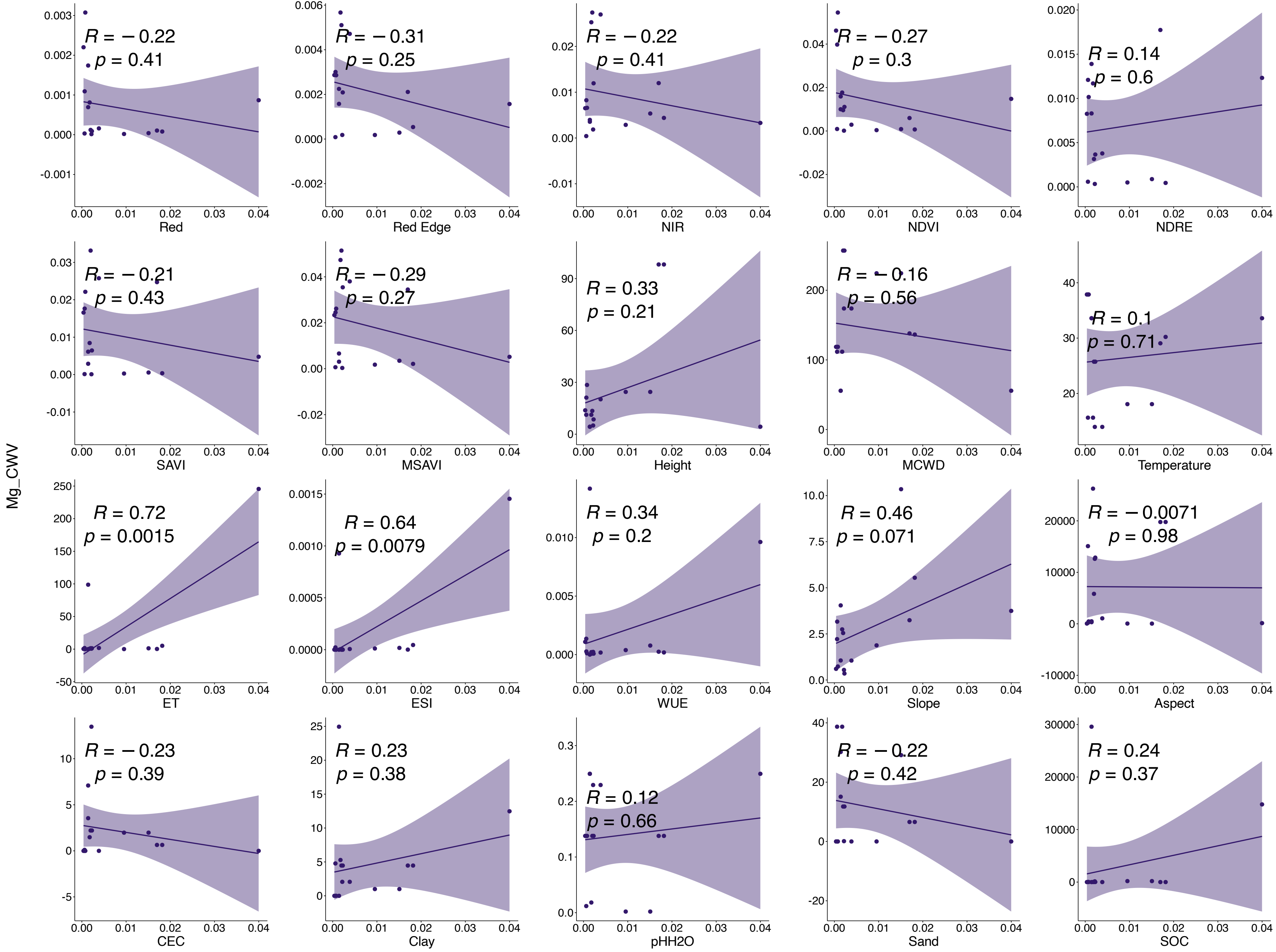


**Fig.45|Relationships between CWV of Mg and variance values of each input band.**

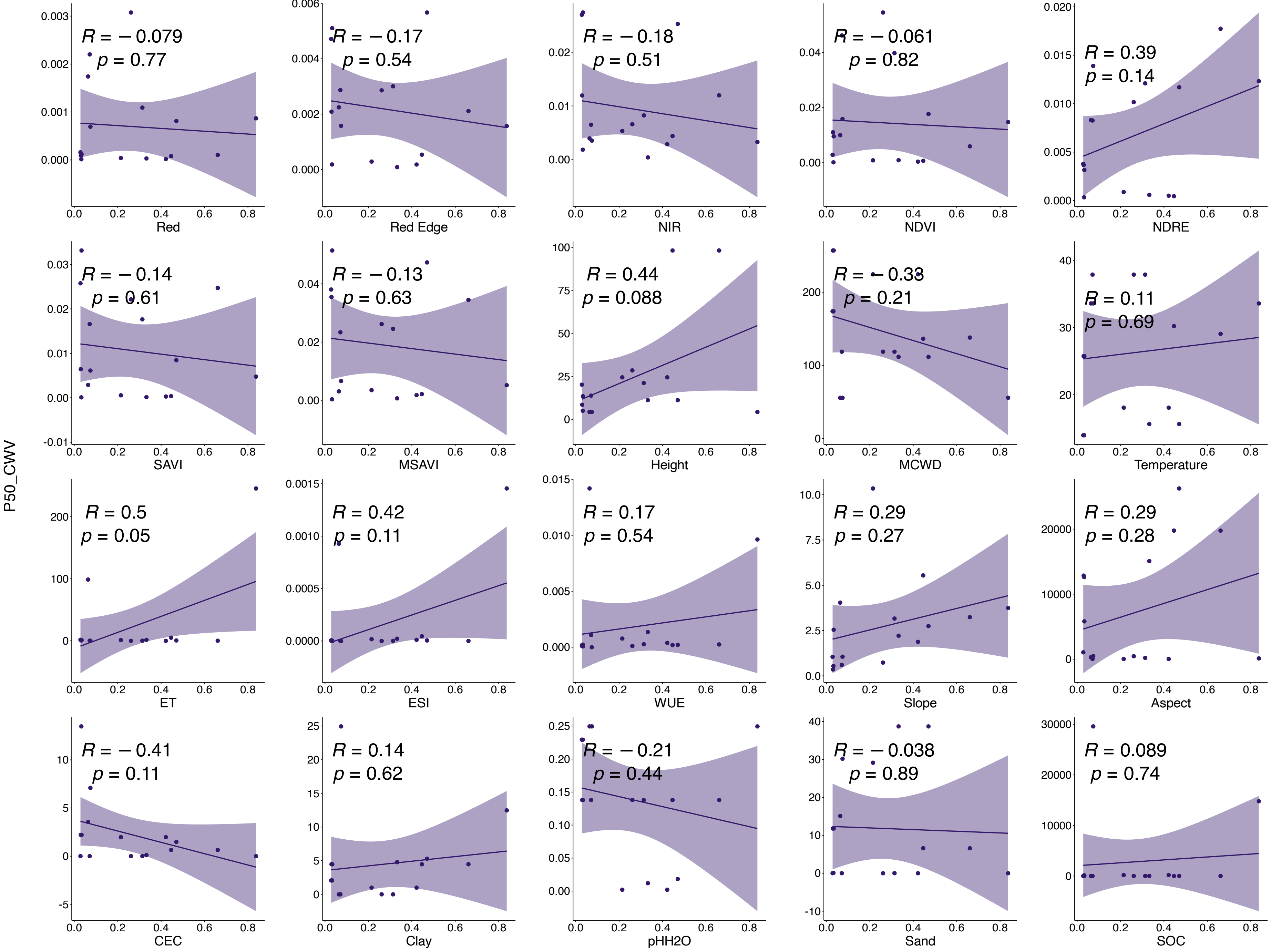


**Fig.46|Relationships between CWV of P50 and variance values of each input band.**

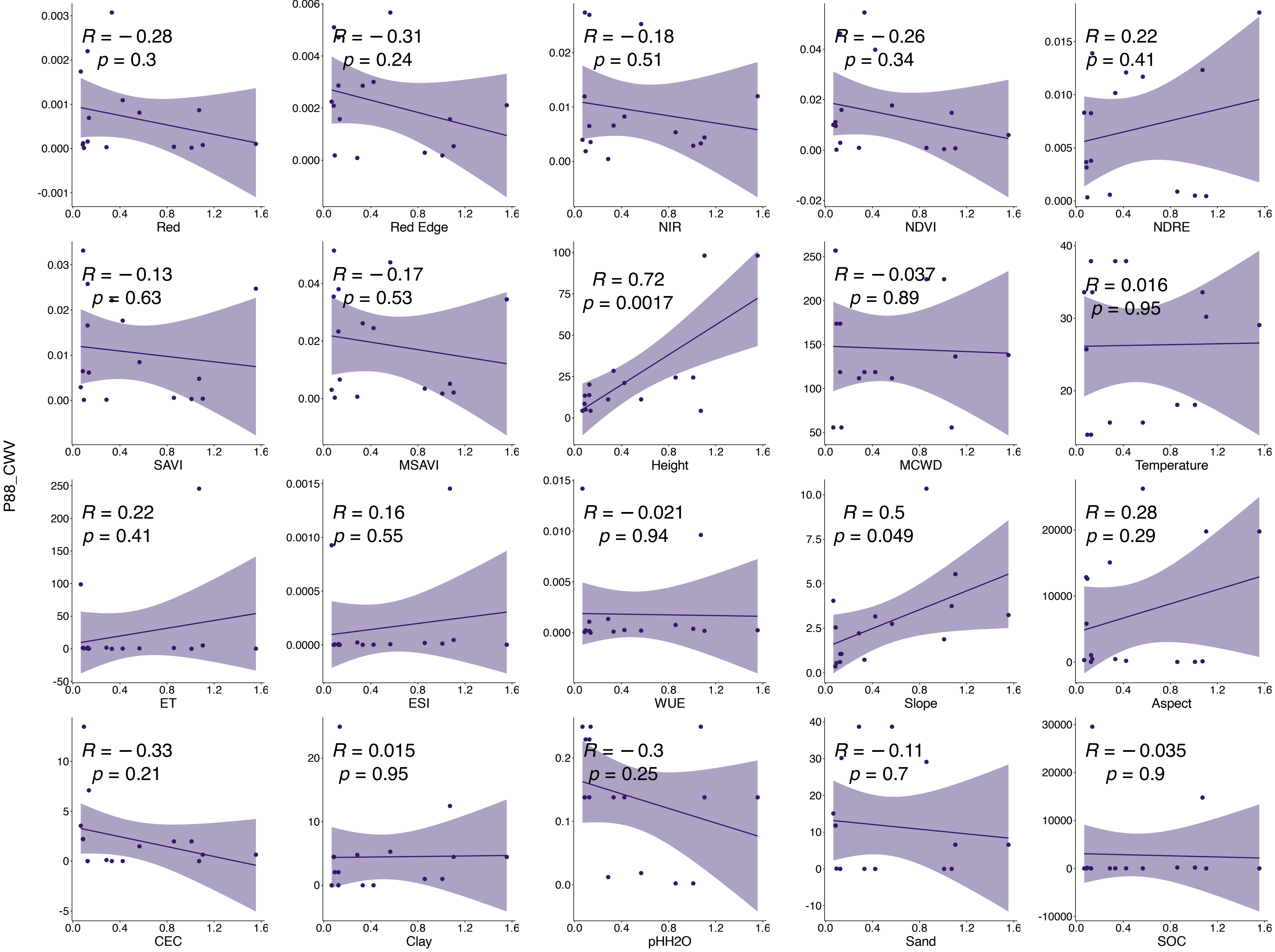


**Fig.47|Relationships between CWV of P88 and variance values of each input band.**

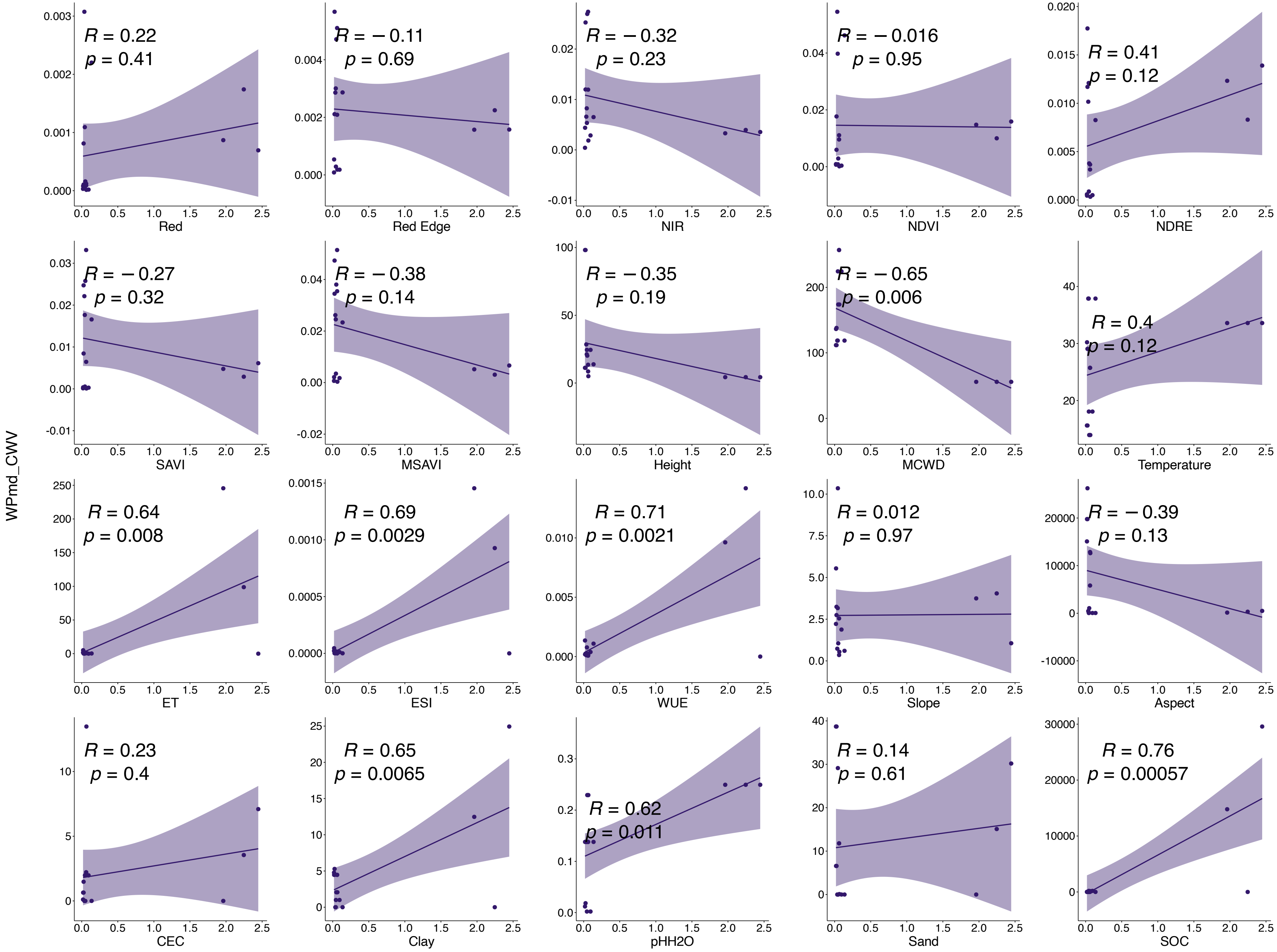


**Fig.48|Relationships between CWV of WPmd and variance values of each input band.**

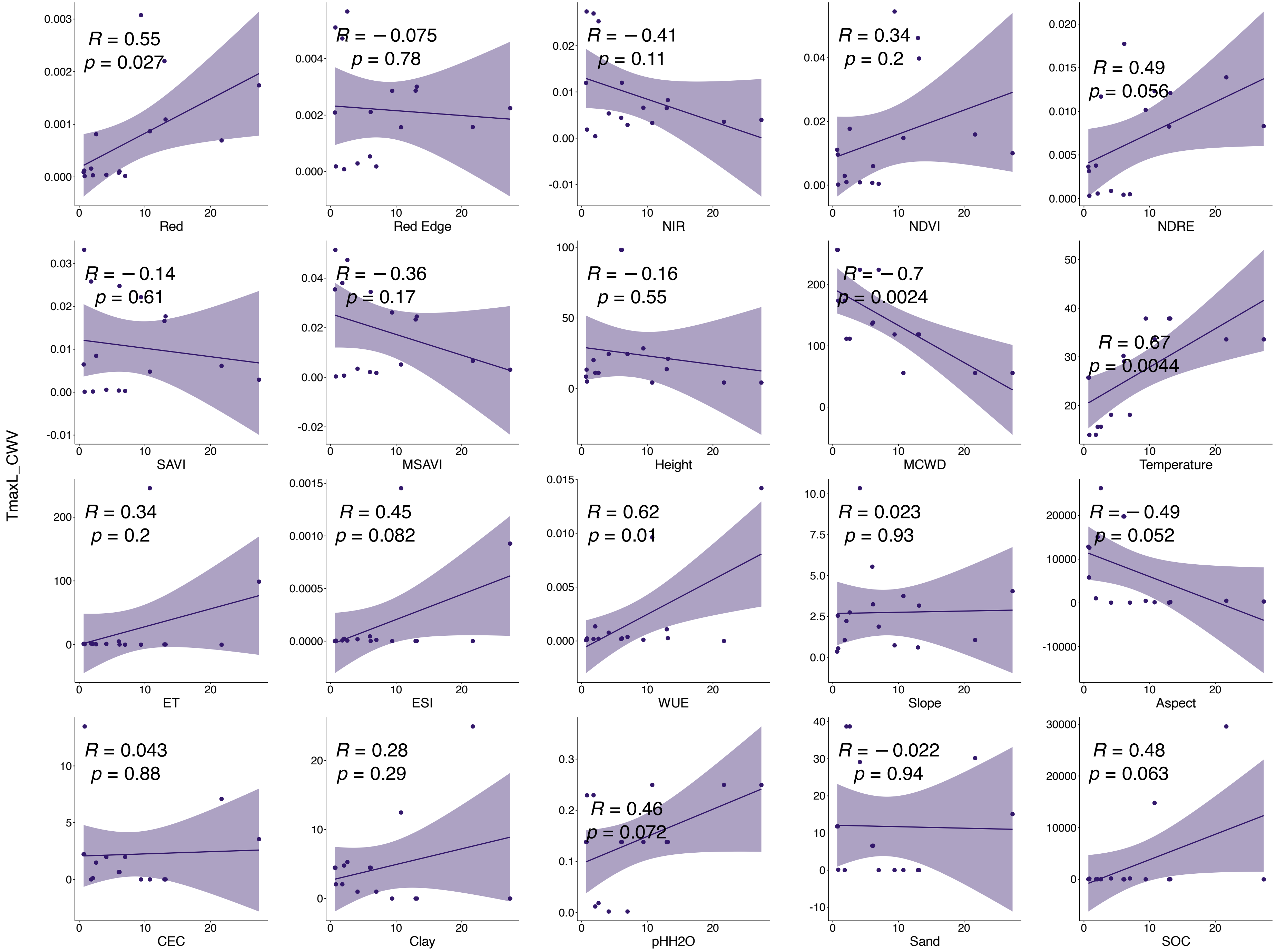


**Fig.49|Relationships between CWV of TmaxL and variance values of each input band.**

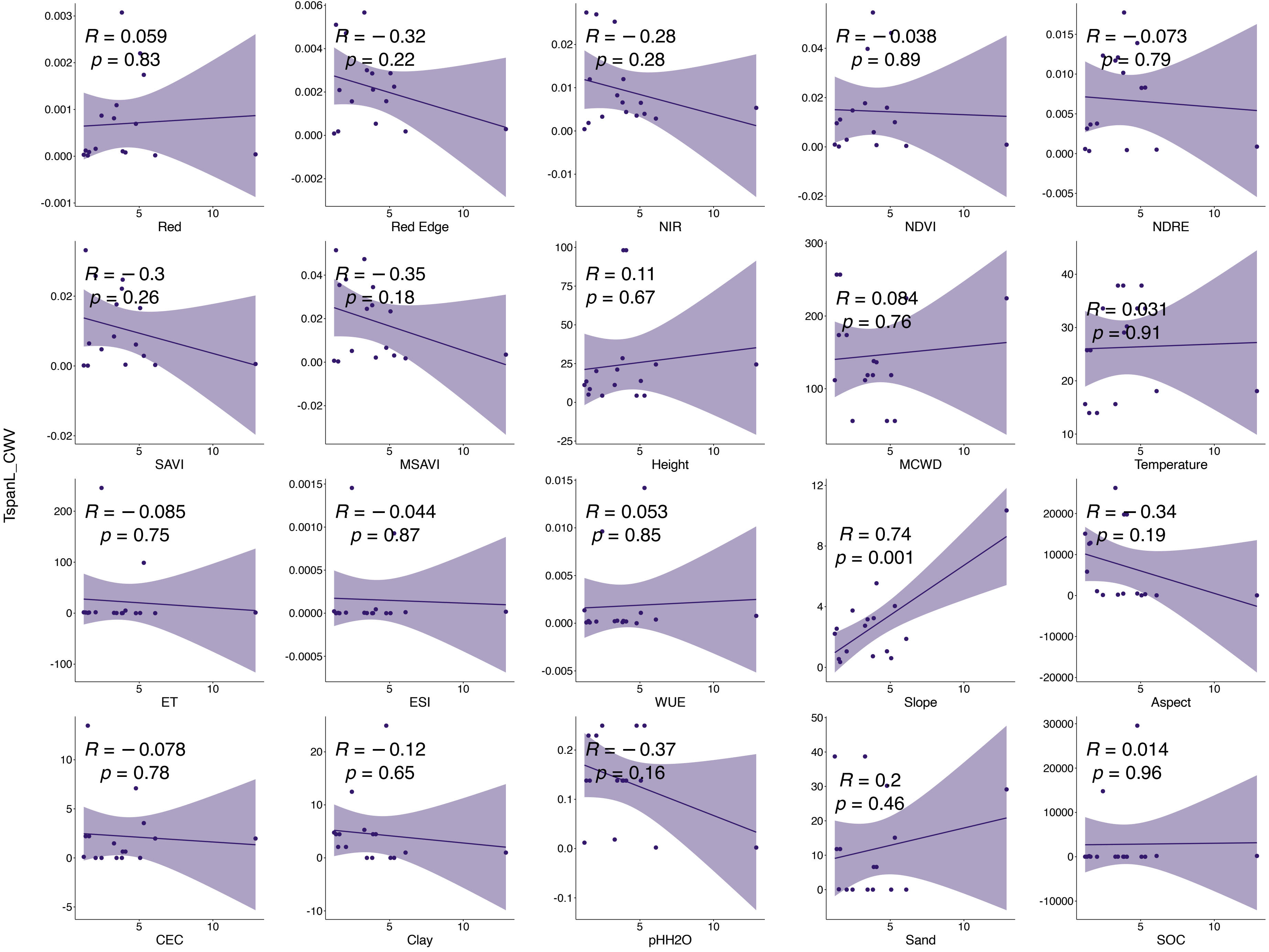


**Fig.50|Relationships between CWV of TspanL and variance values of each input band.**

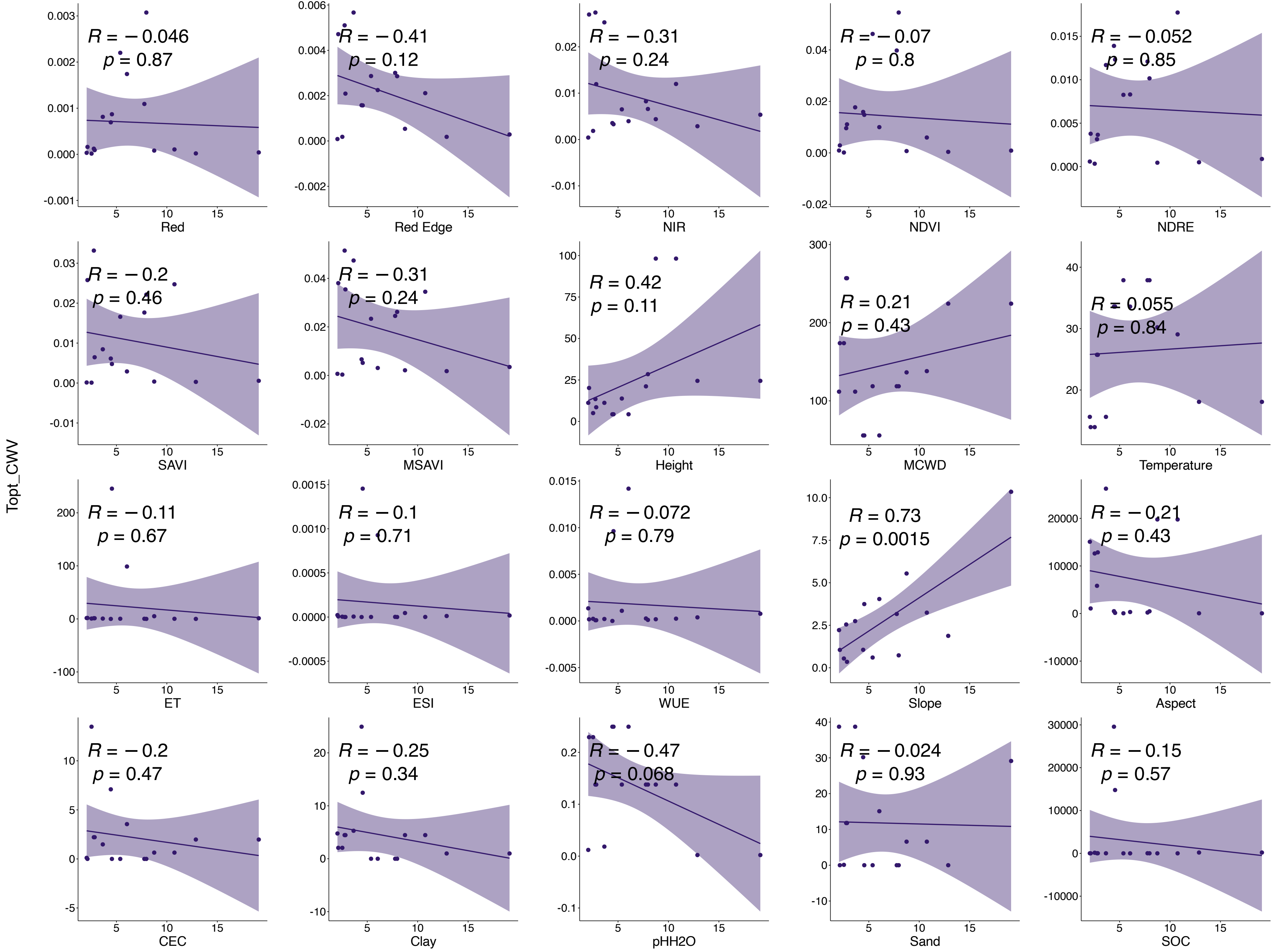


**Fig.51|Relationships between CWV of Topt and variance values of each input band.**

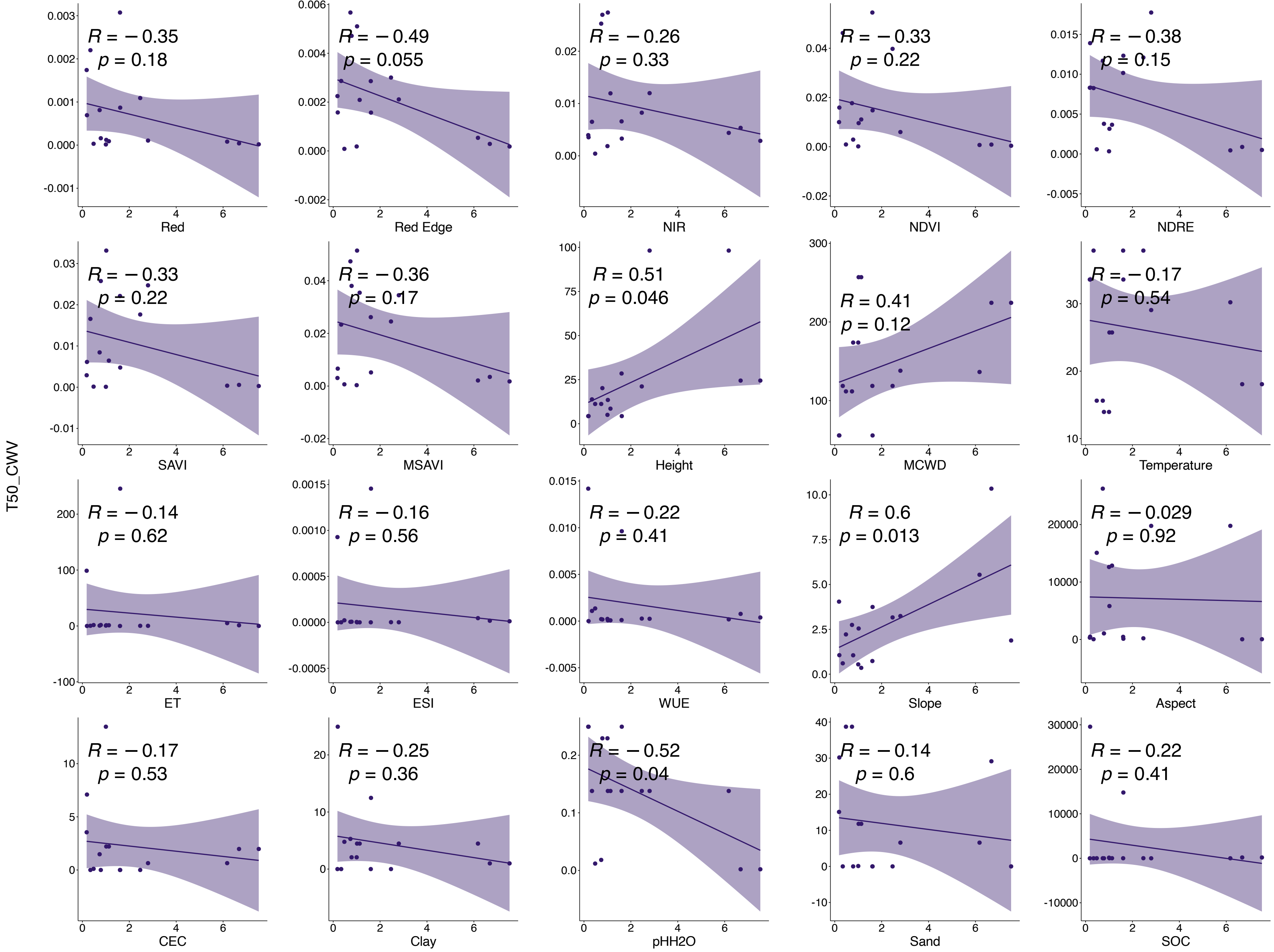


**Fig.52|Relationships between CWV of T50 and variance values of each input band.**

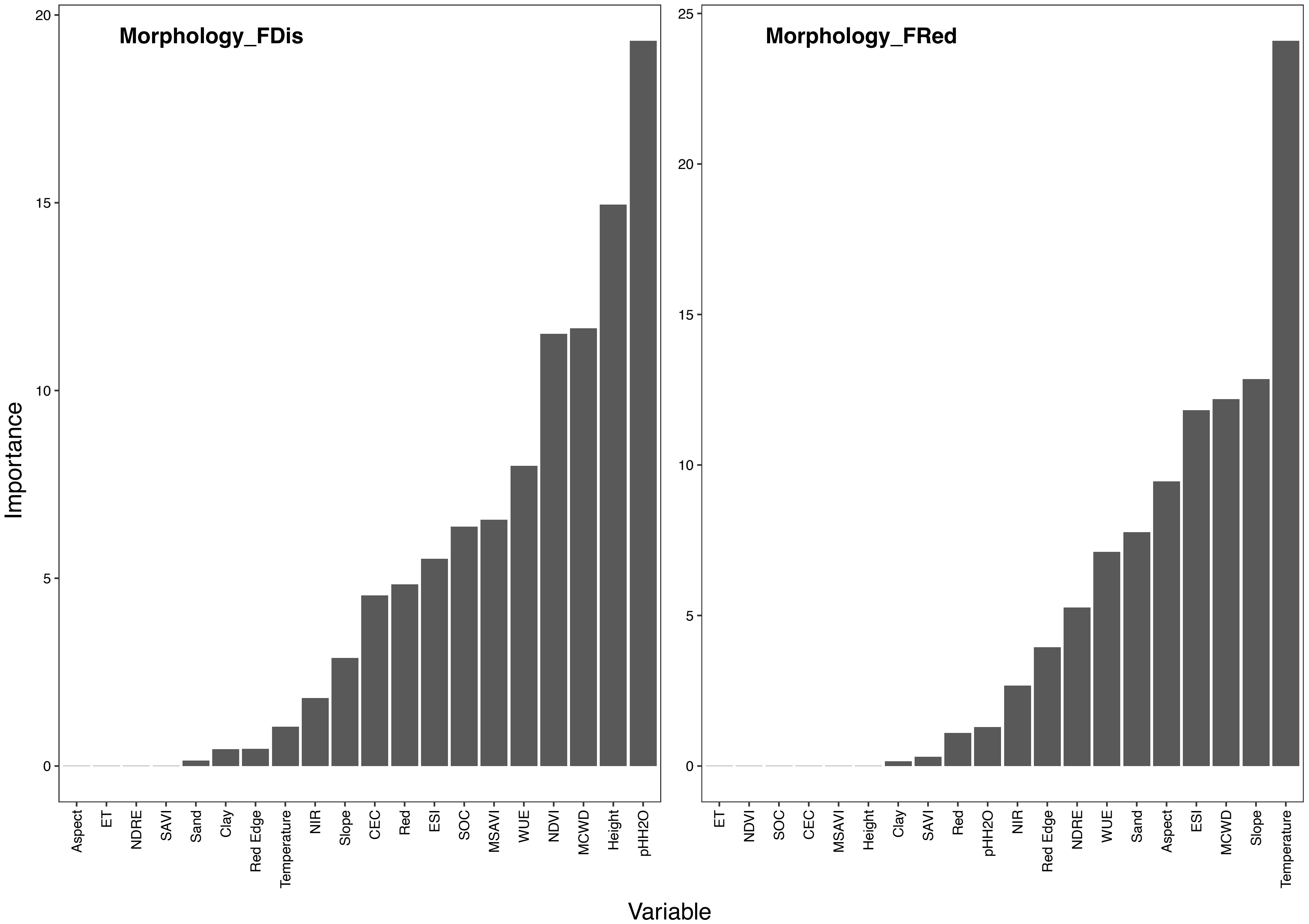


**Fig.53|Drivers of morphological functional dispersion and redundancy.** Importance of all variables in Random forests regression models in predicting morphological FDis (left) and FRed (right).

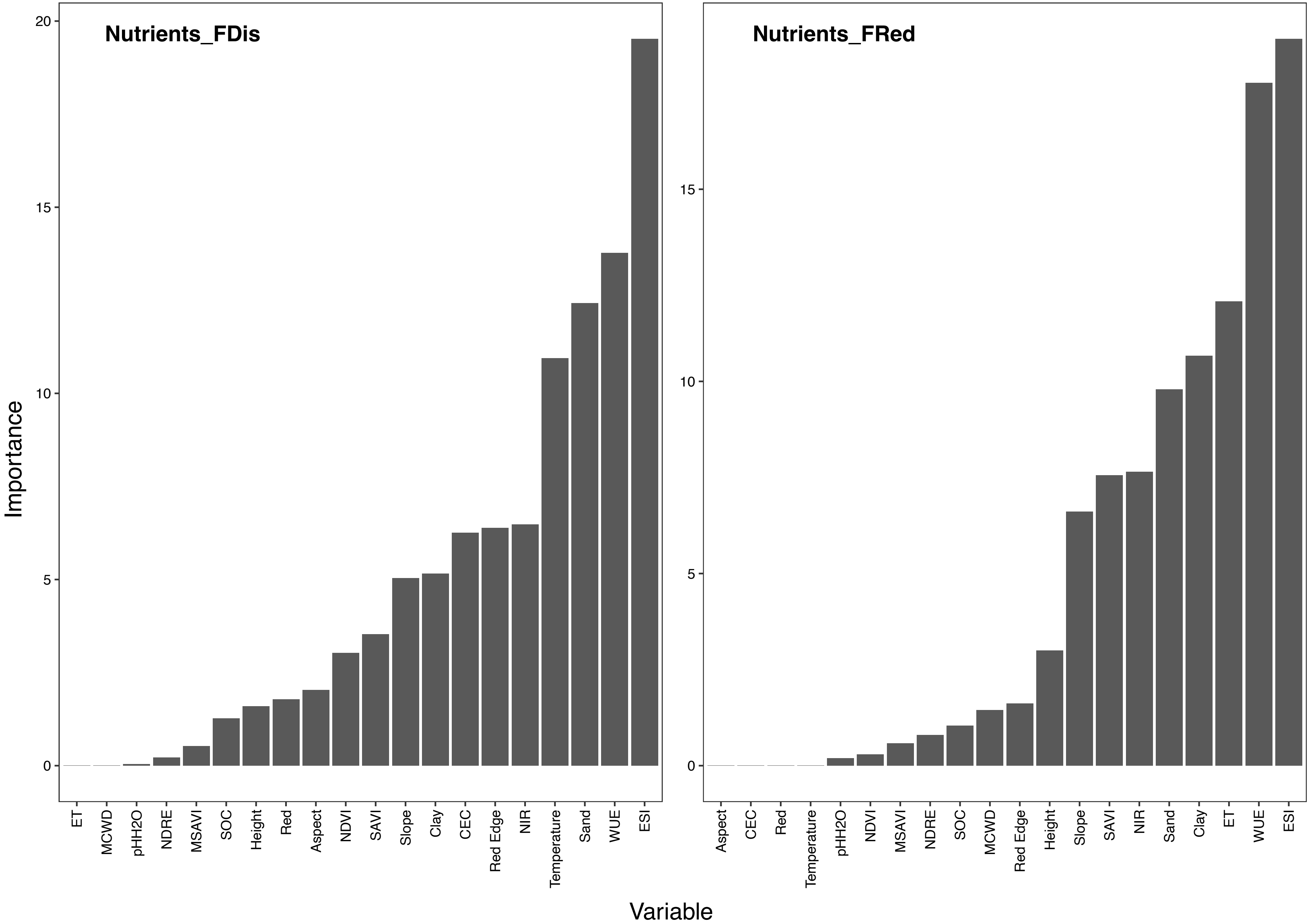


**Fig.54|Drivers of nutrient functional dispersion and redundancy.** Importance of all variables in Random forests regression models in predicting nutrient FDis (left) and FRed (right).

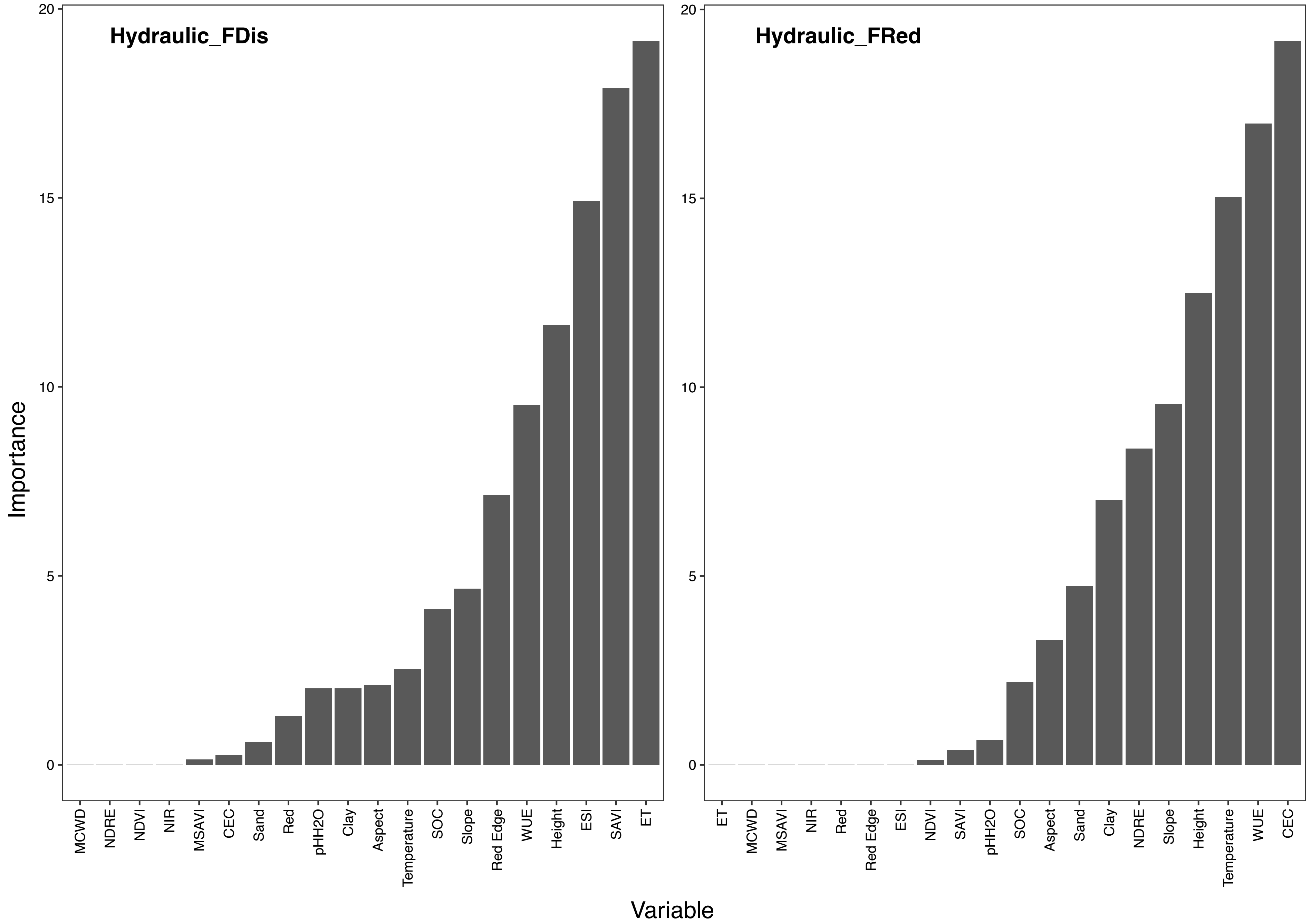


**Fig.55|Drivers of hydraulic functional dispersion and redundancy.** Importance of all variables in Random forests regression models in predicting hydraulic FDis (left) and FRed (right).

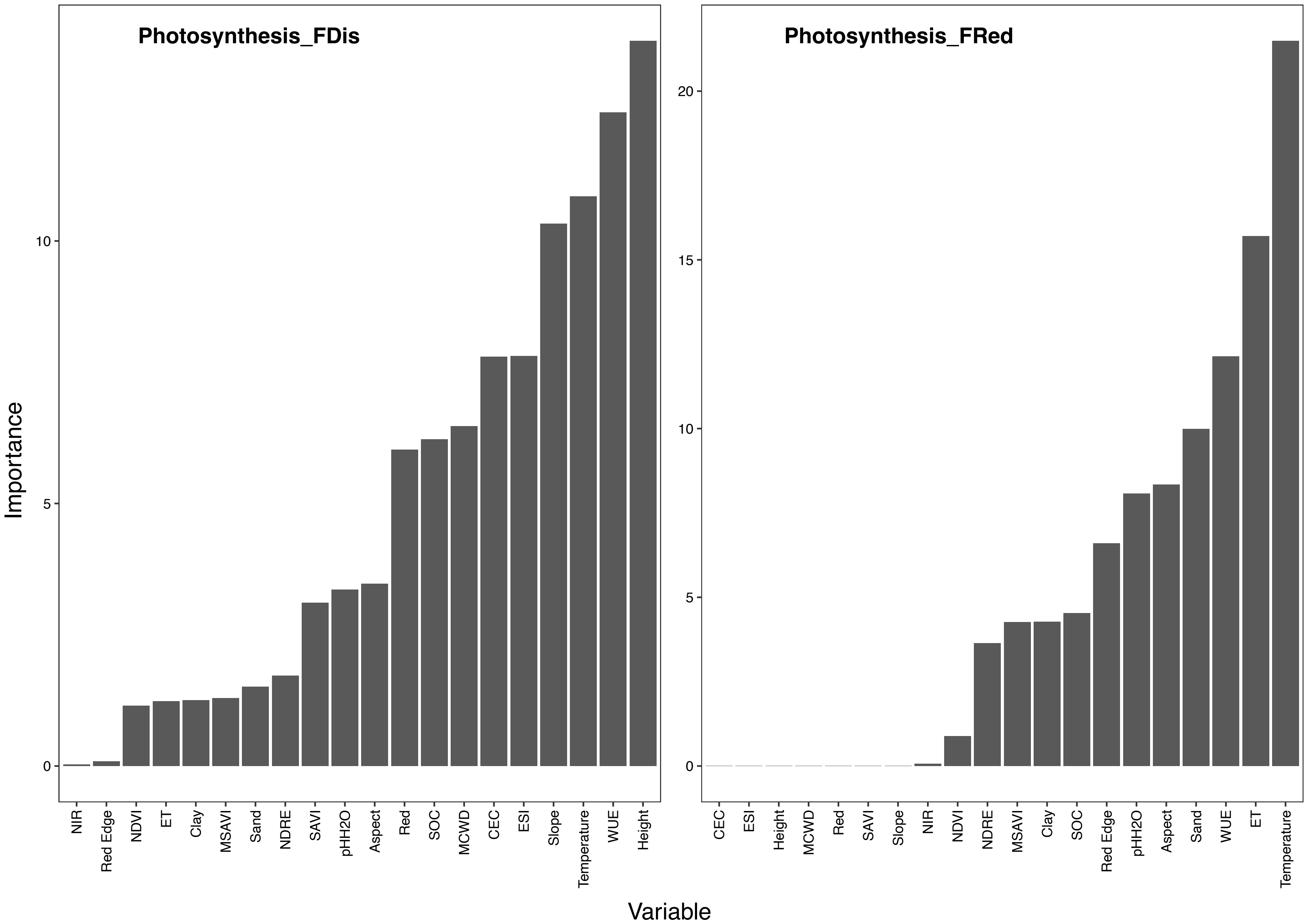


**Fig.56|Drivers of photosynthetic functional dispersion and redundancy.** Importance of all variables in Random forests regression models in predicting photosynthetic FDis (left) and FRed (right).

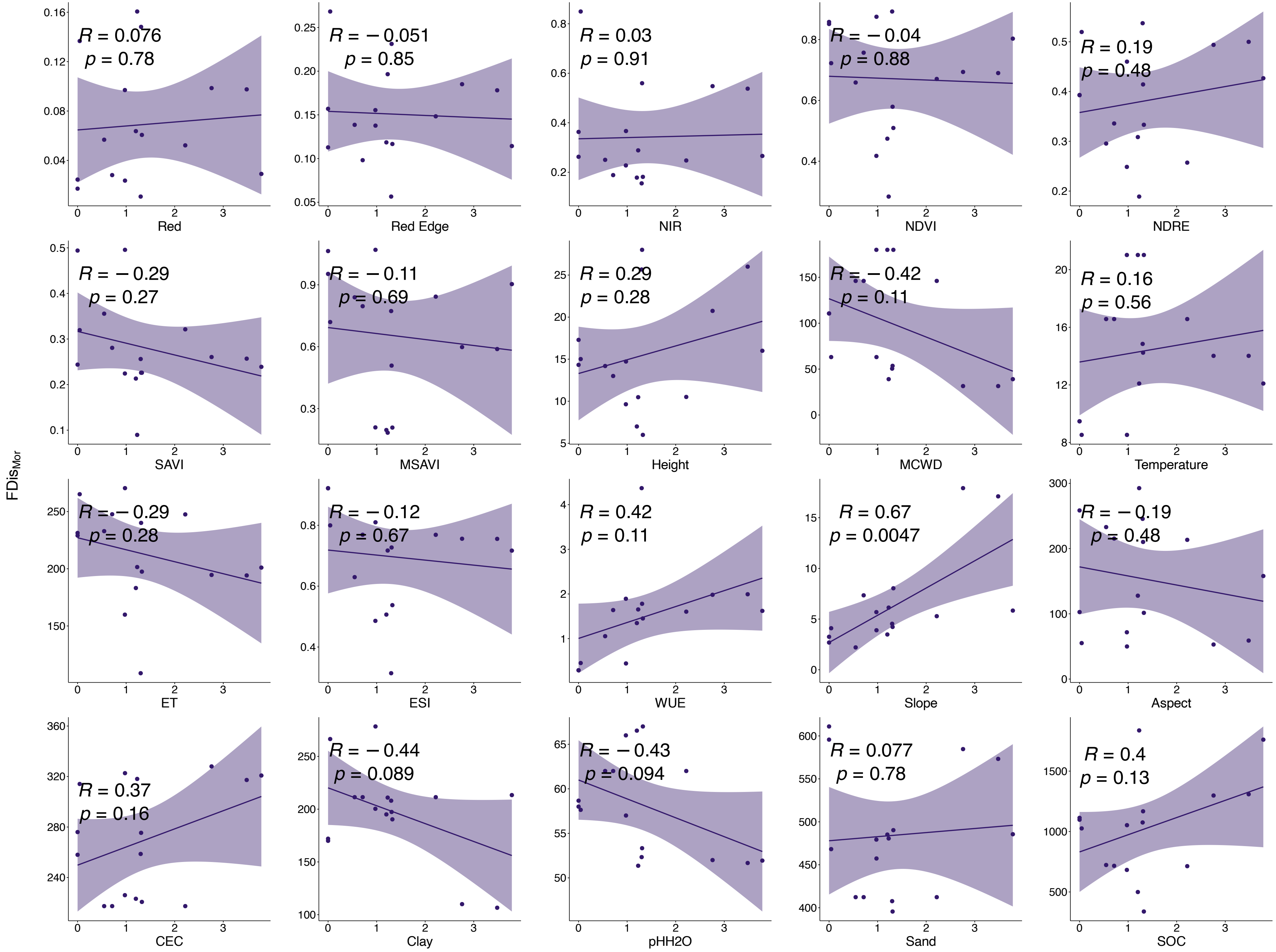


**Fig.57|Relationships between $FDis_{Morphology}$ and mean values of each input band.**

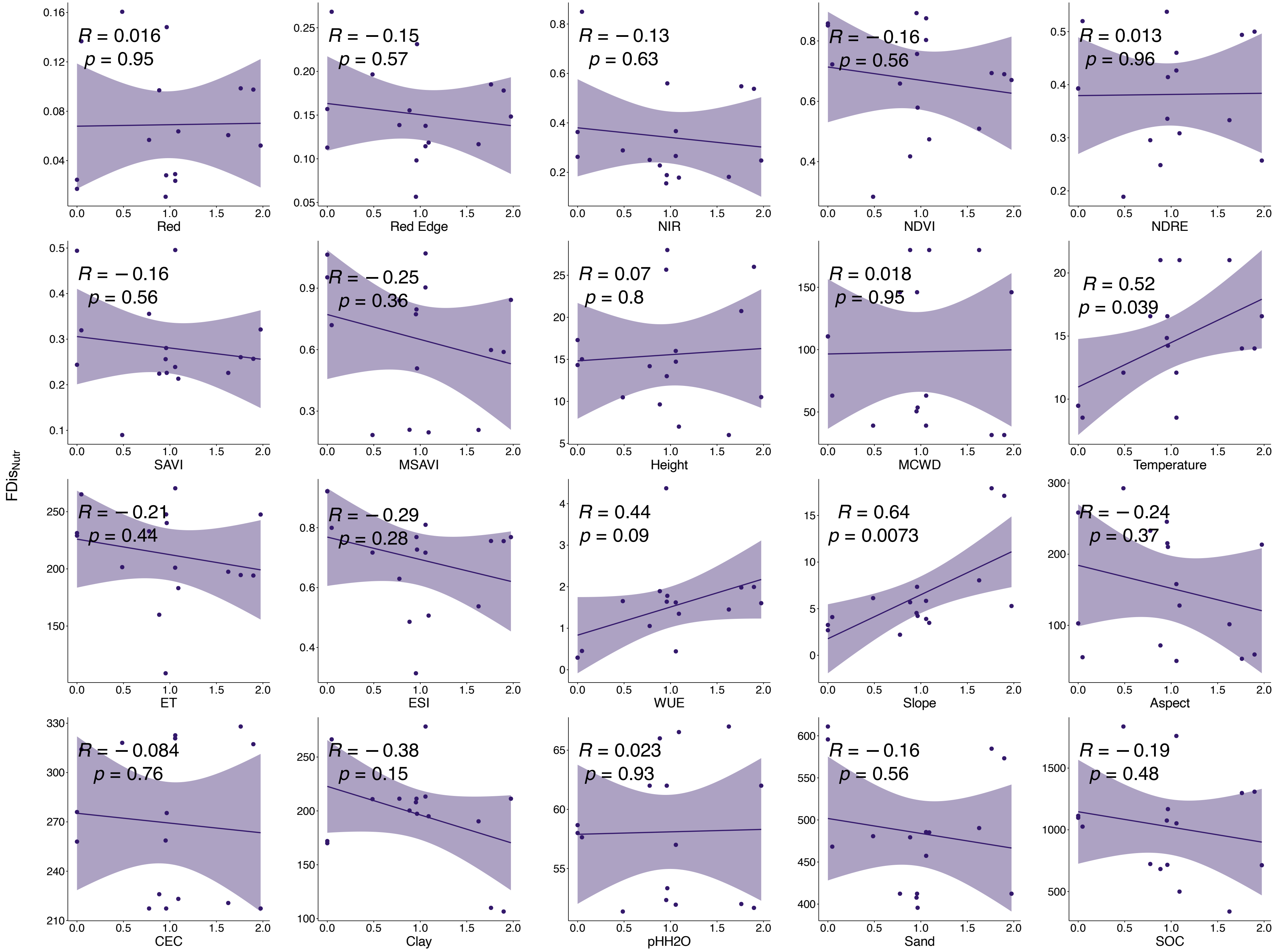


**Fig.58|Relationships between FDis$_{\text{Nutrients}}$ and mean values of each input band.**

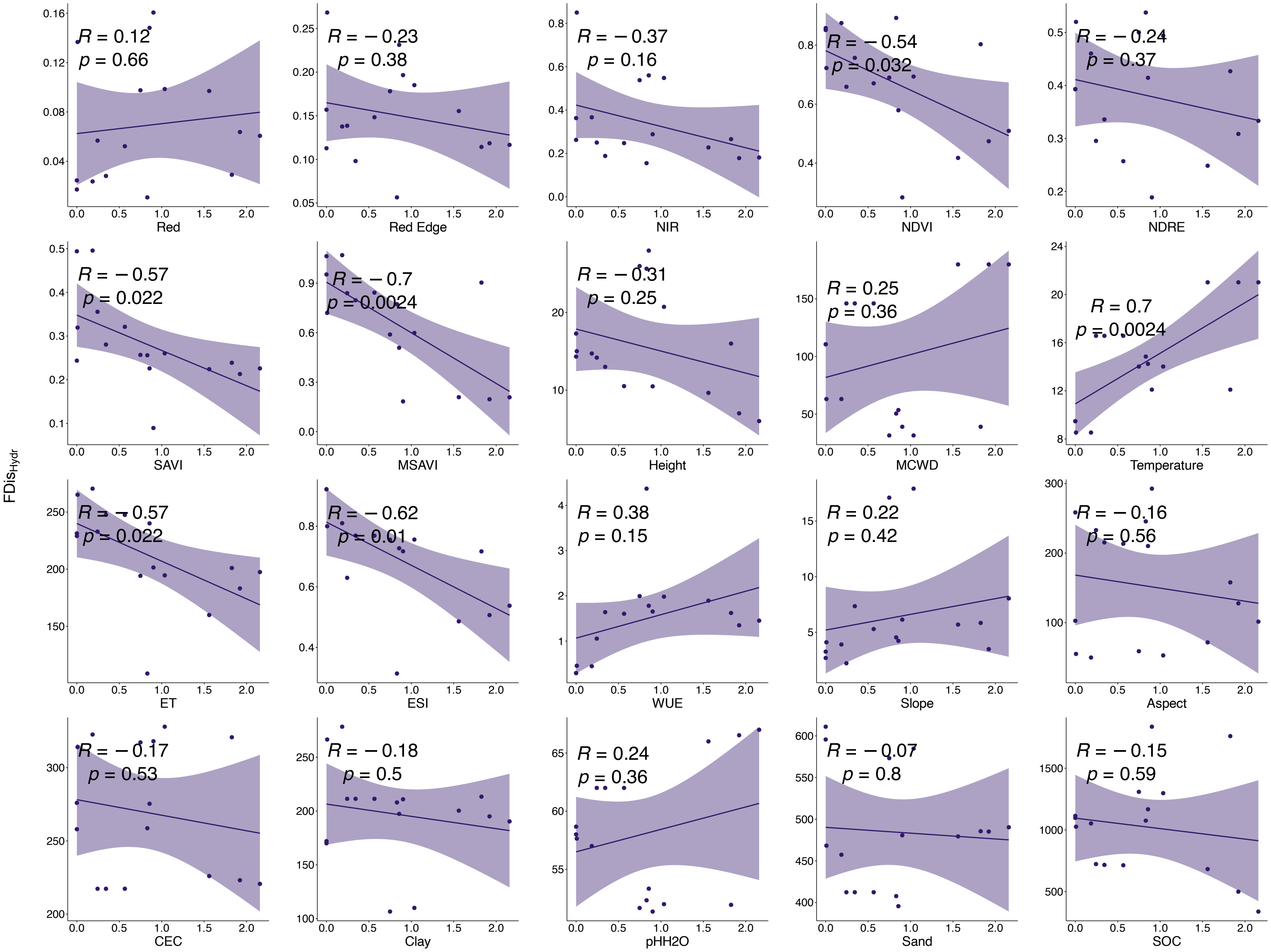


**Fig.59|Relationships between $FDis_{Hydraulic}$ and mean values of each input band.**

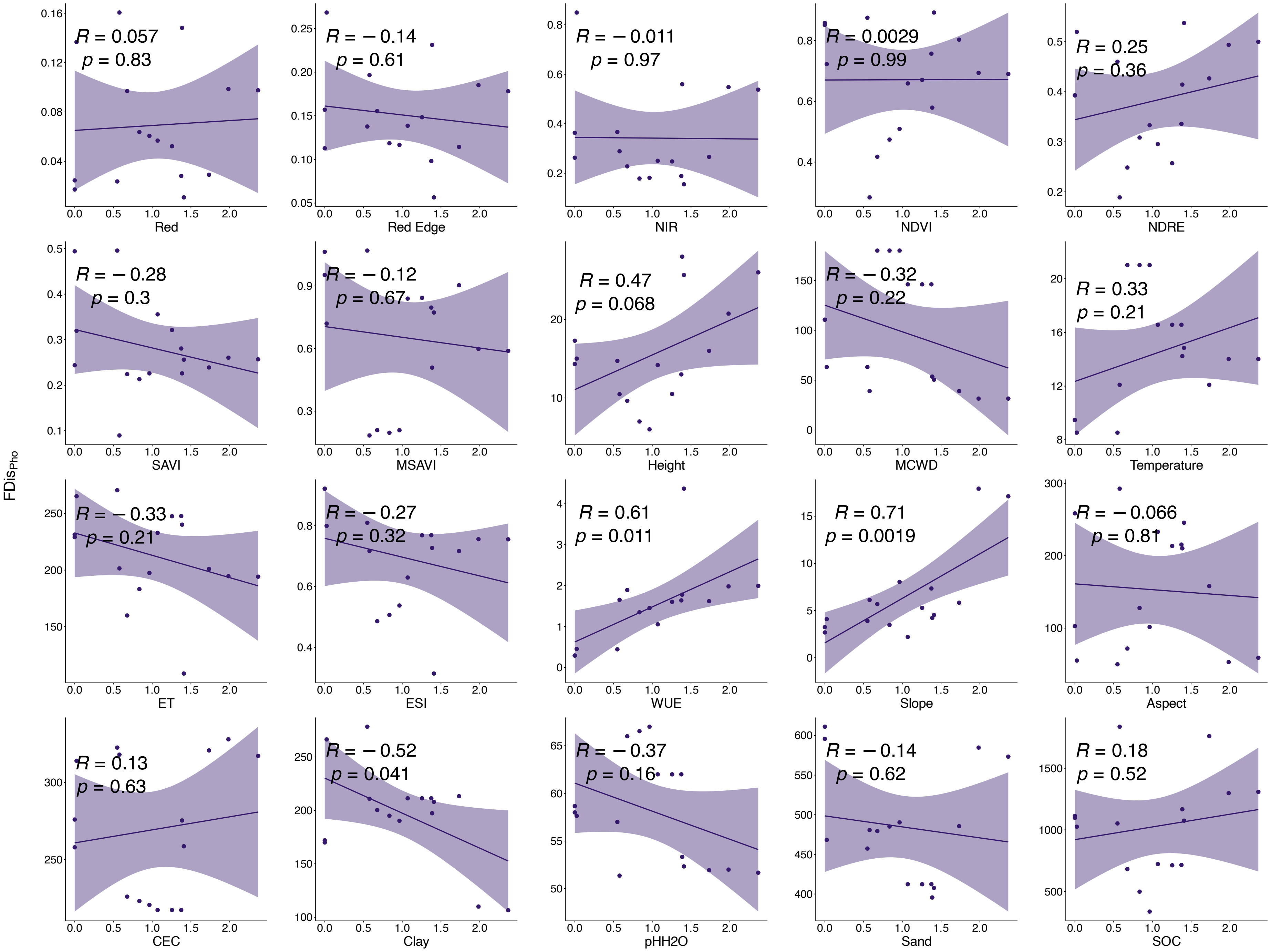


**Fig.60|Relationships between $FDis_{Photosynthesis}$ and mean values of each input band.**

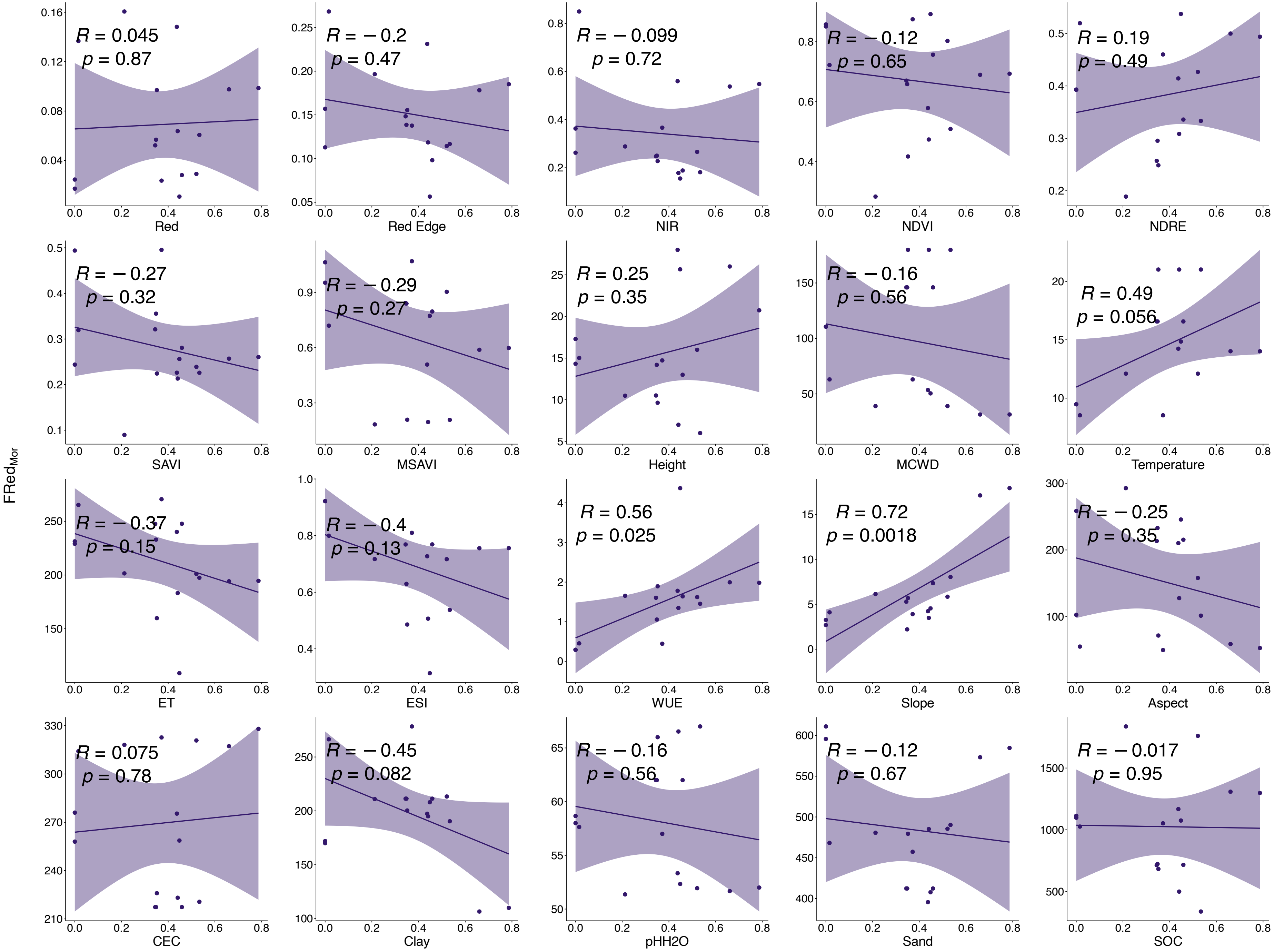


**Fig.61|Relationships between $FRed_{Morphology}$ and mean values of each input band.**

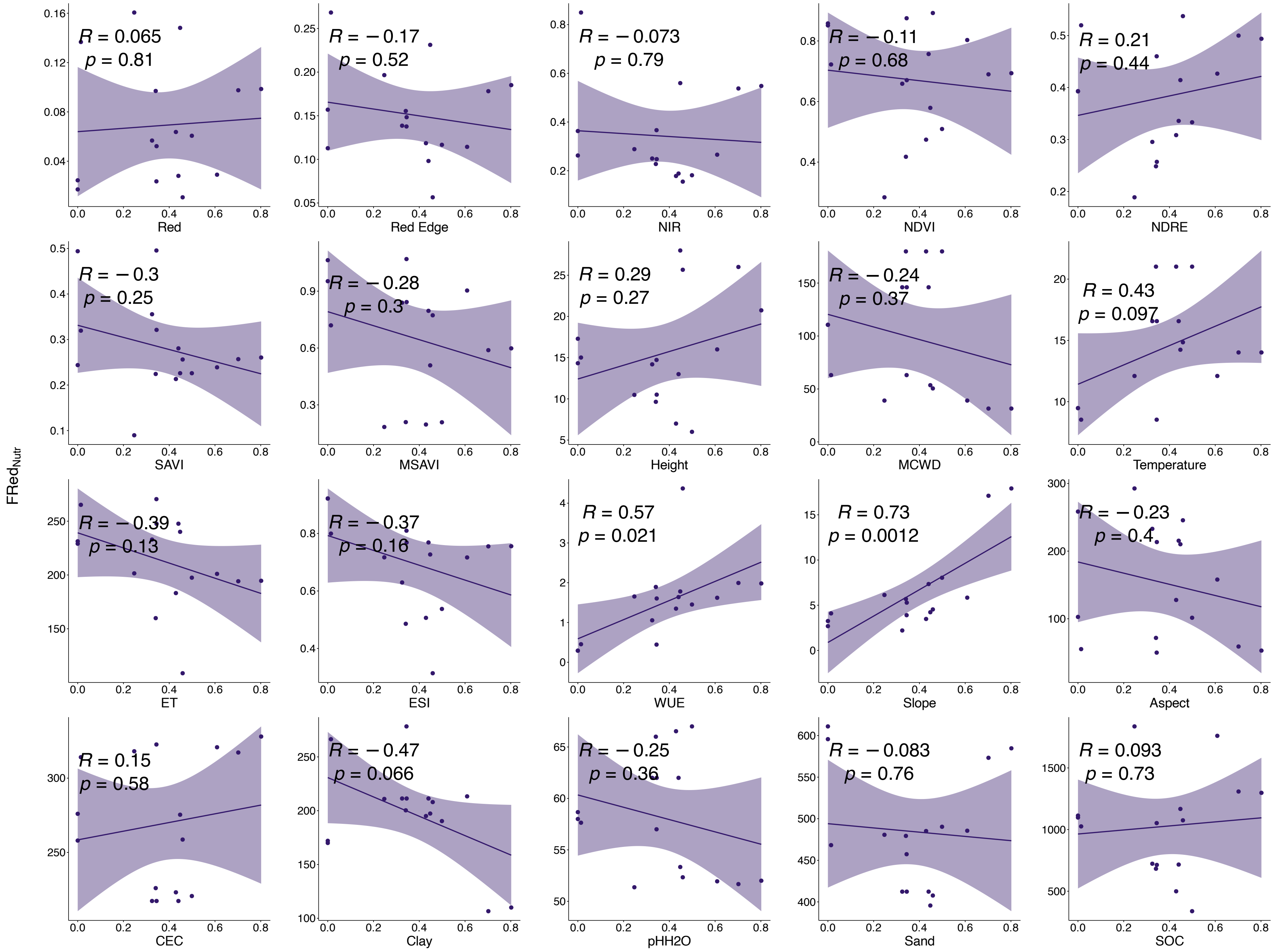


**Fig.62|Relationships between $FRed_{Nutrients}$ and mean values of each input band.**

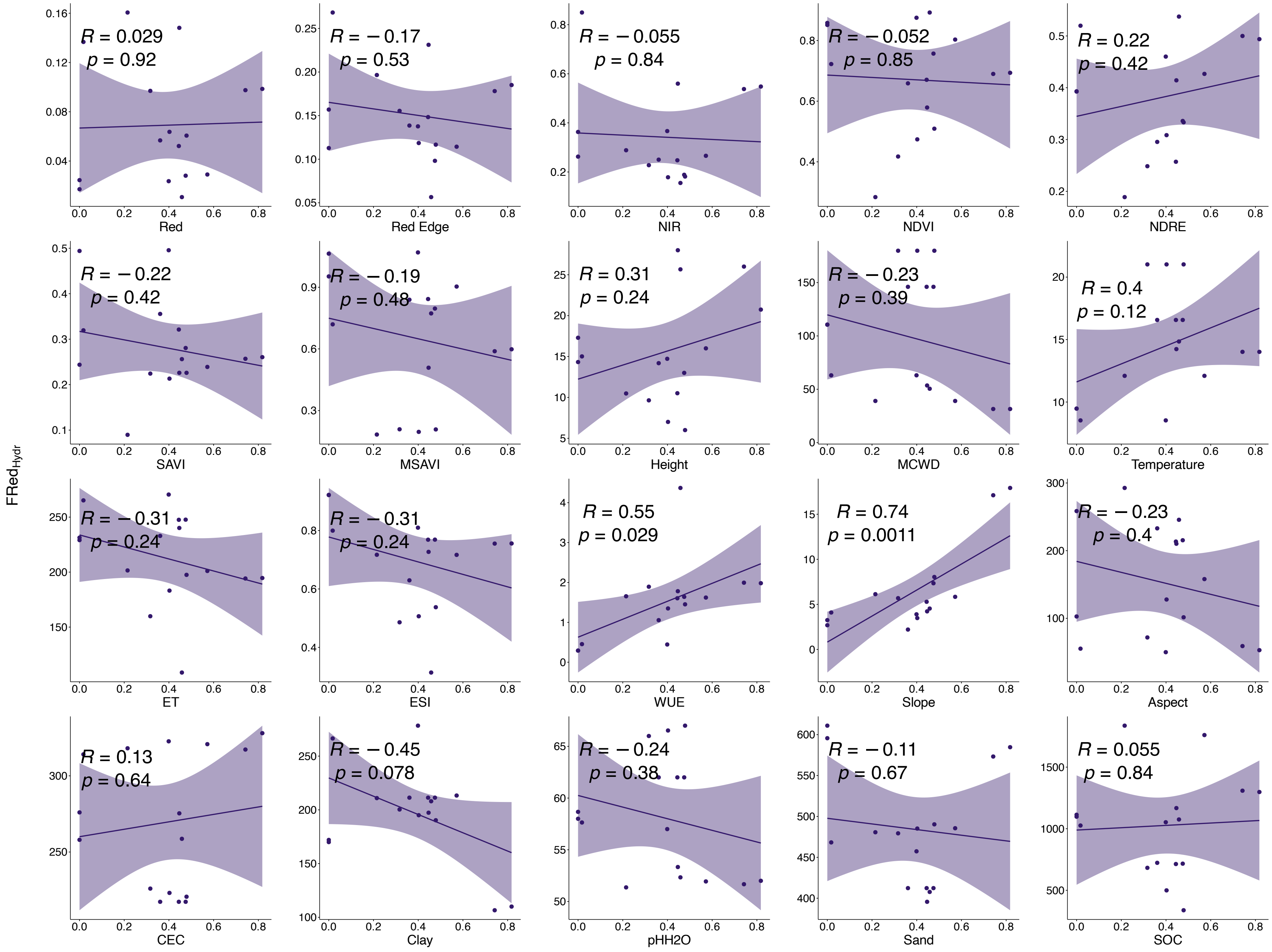


**Fig.63|Relationships between $FRed_{Hydraulic}$ and mean values of each input band.**

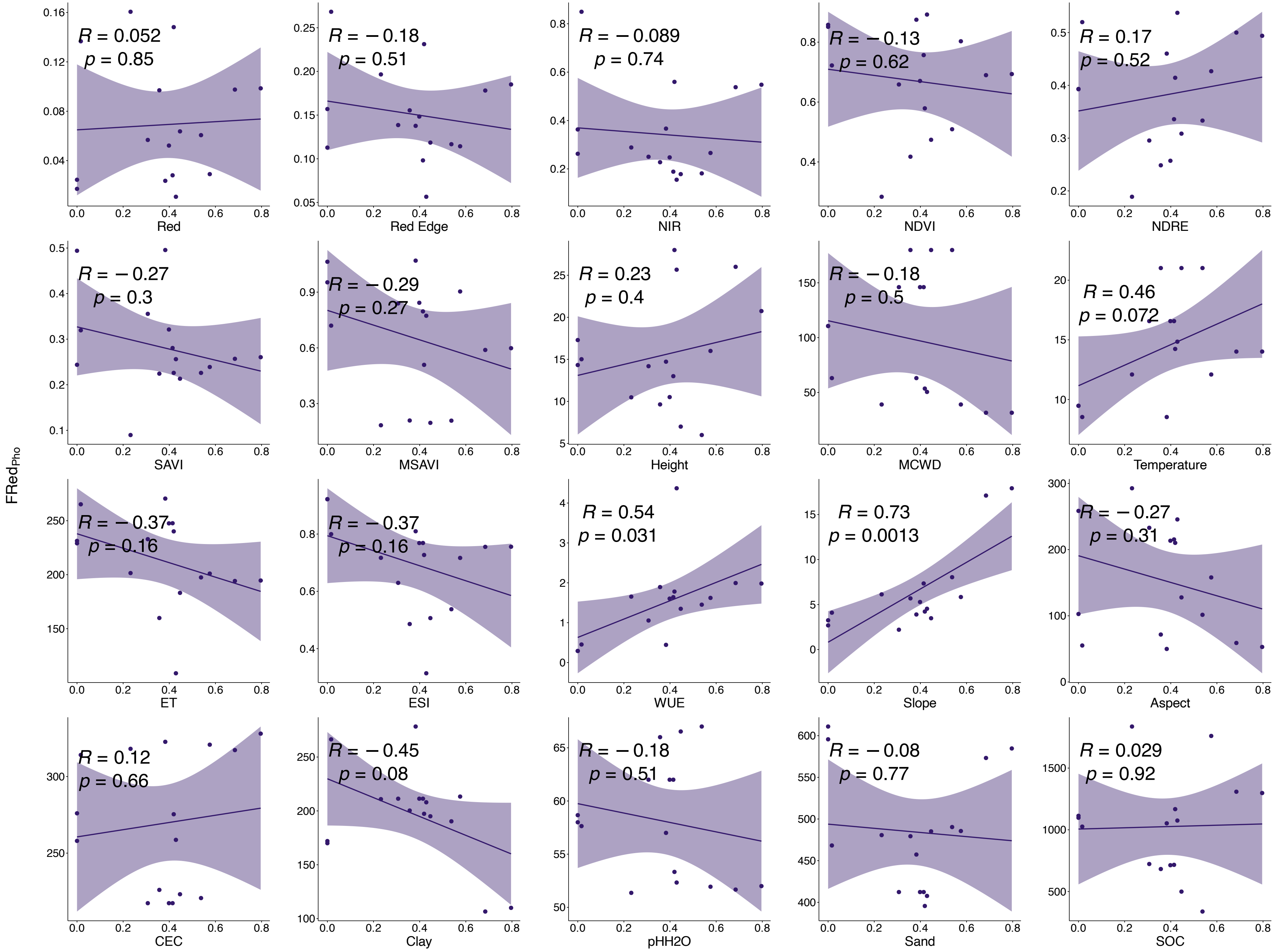


**Fig.64|Relationships between $FRed_{Photosynthesis}$ and mean values of each input band.**

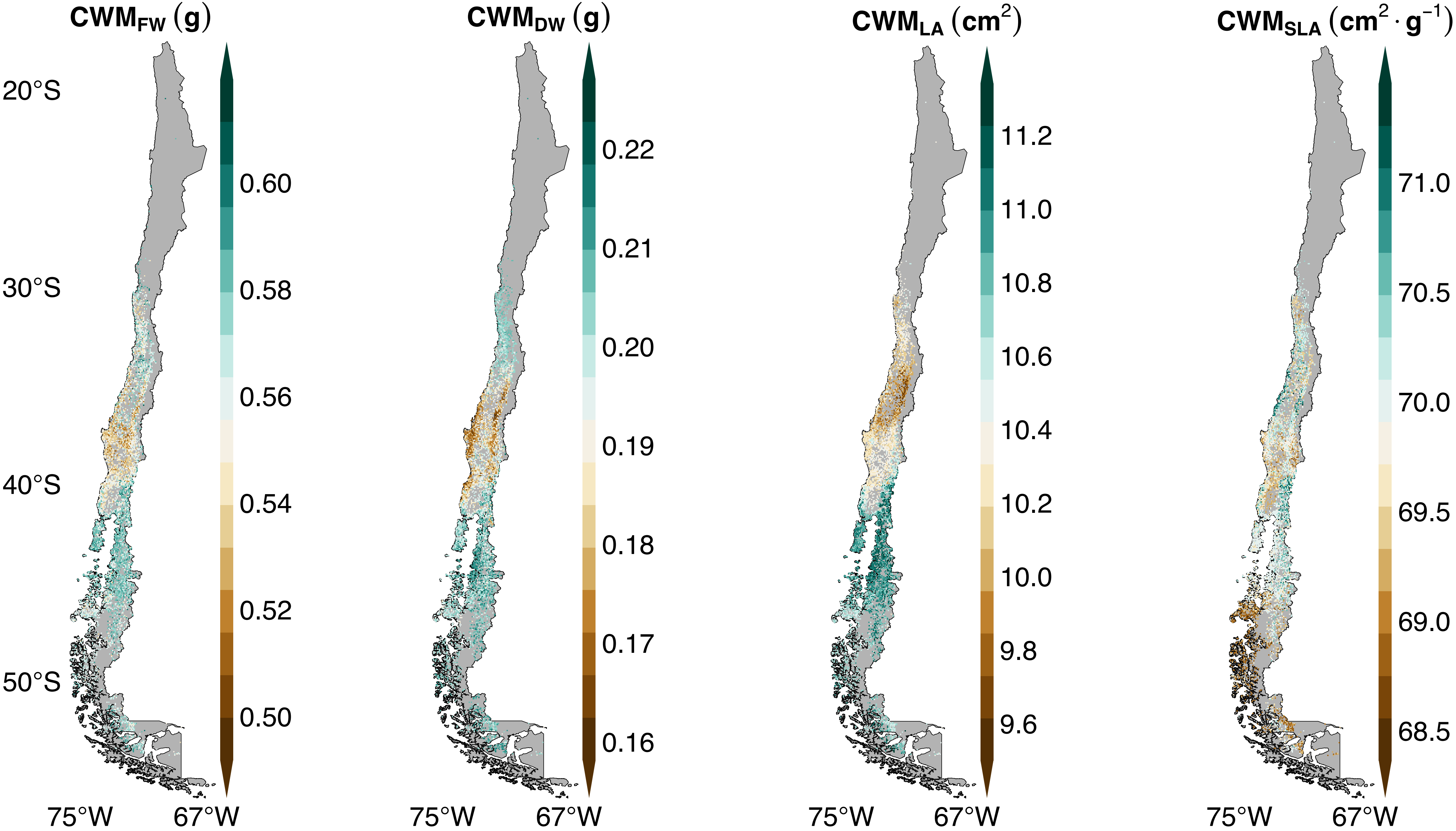


**Fig.65|Distribution maps of CWM of FW, DW, LA, and SLA.**

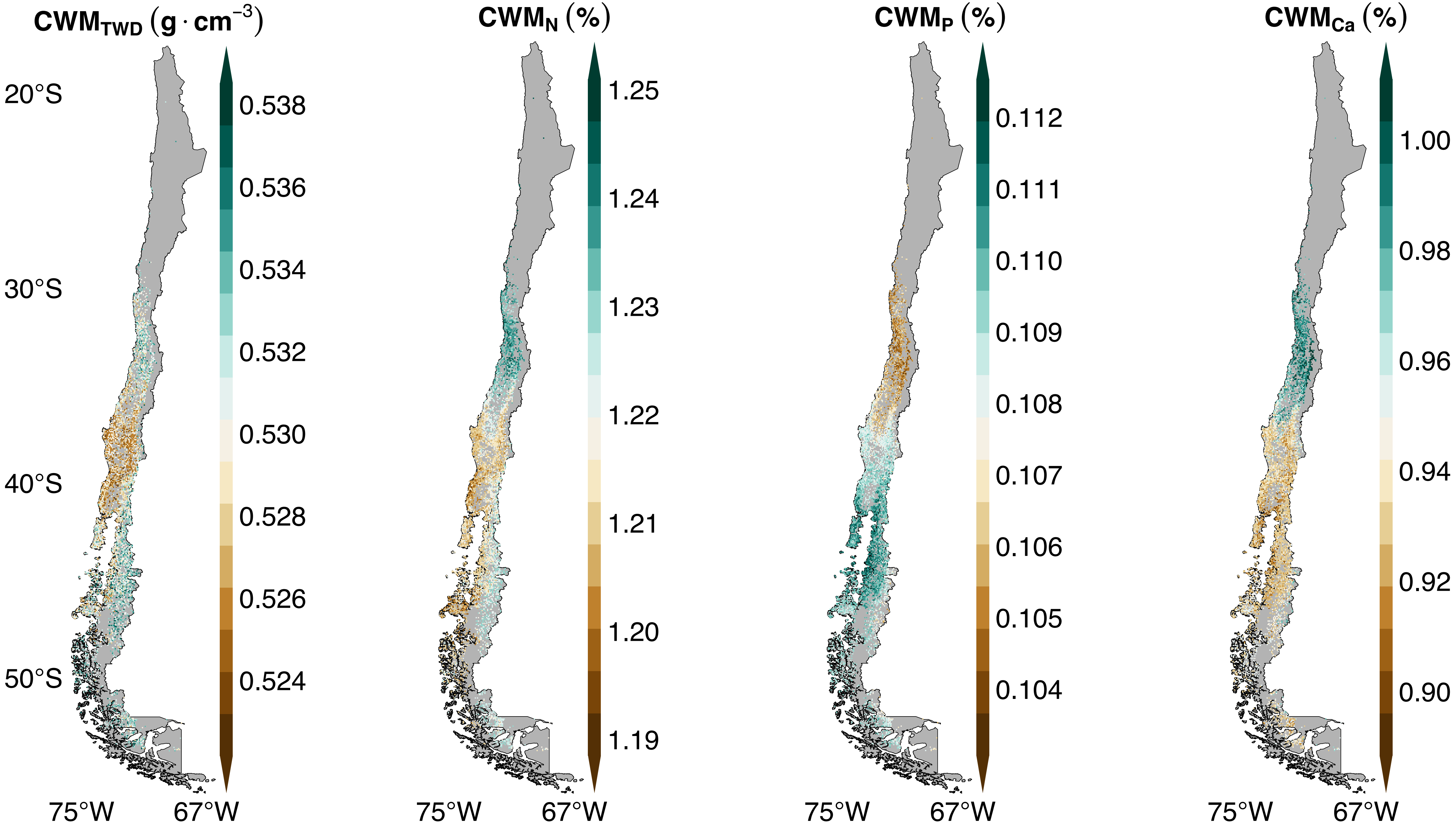


**Fig.66|Distribution maps of CWM of TWD, N, P, and Ca.**

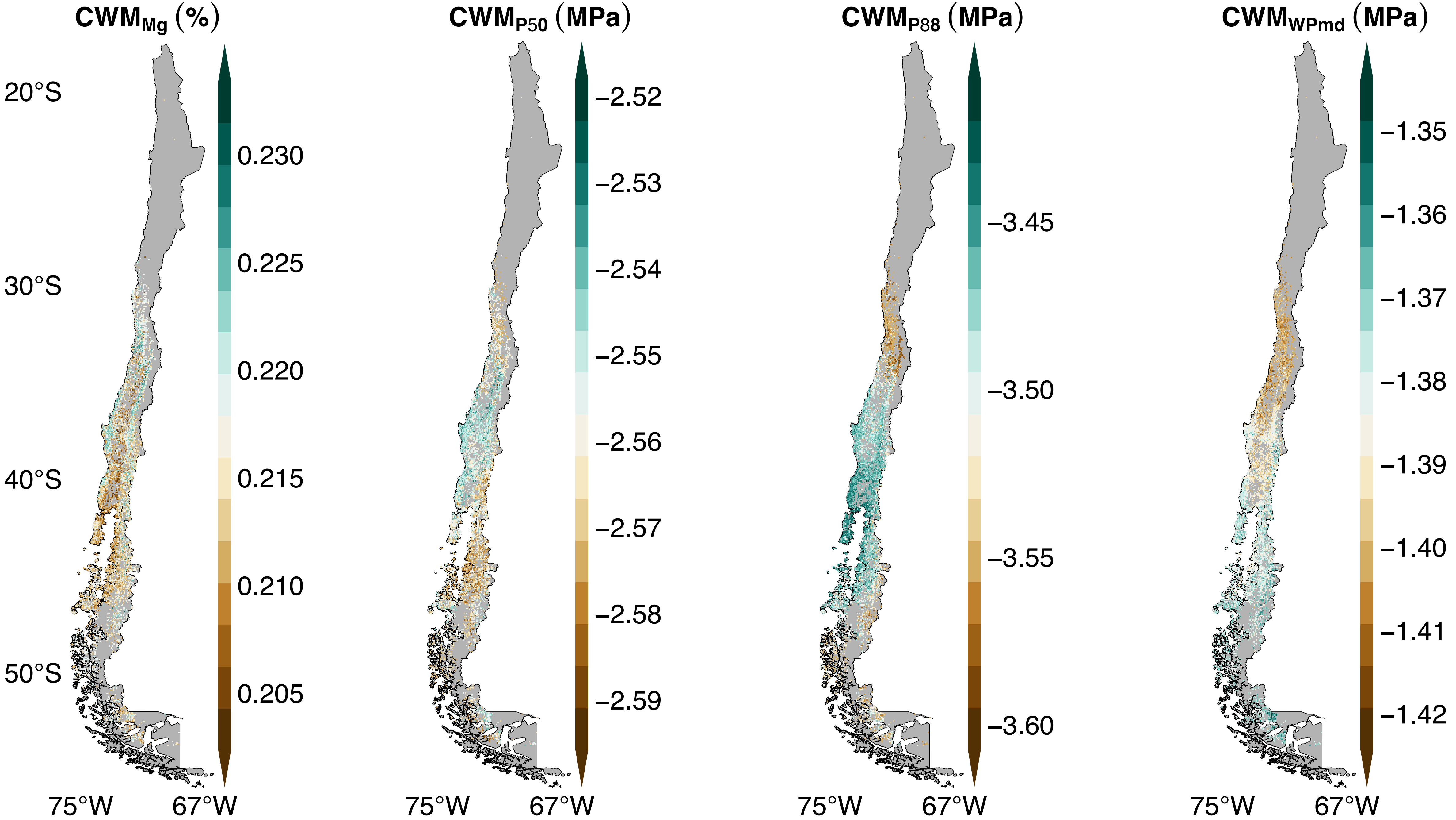


**Fig.67|Distribution maps of CWM of Mg, P50, P88, and WPmd.**

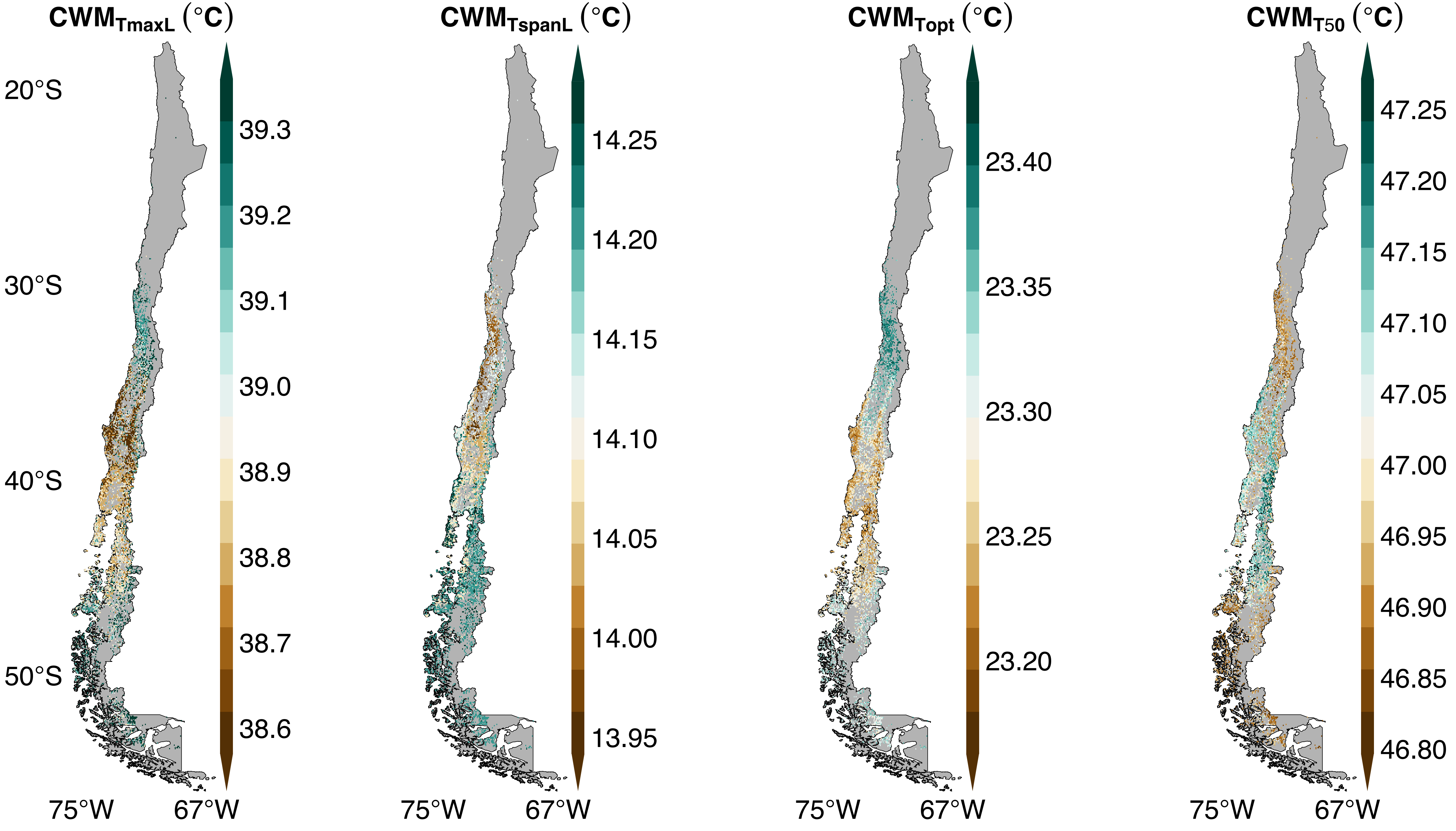


**Fig.68|Distribution maps of CWM of TmaxL, TspanL, Topt, and T50.**

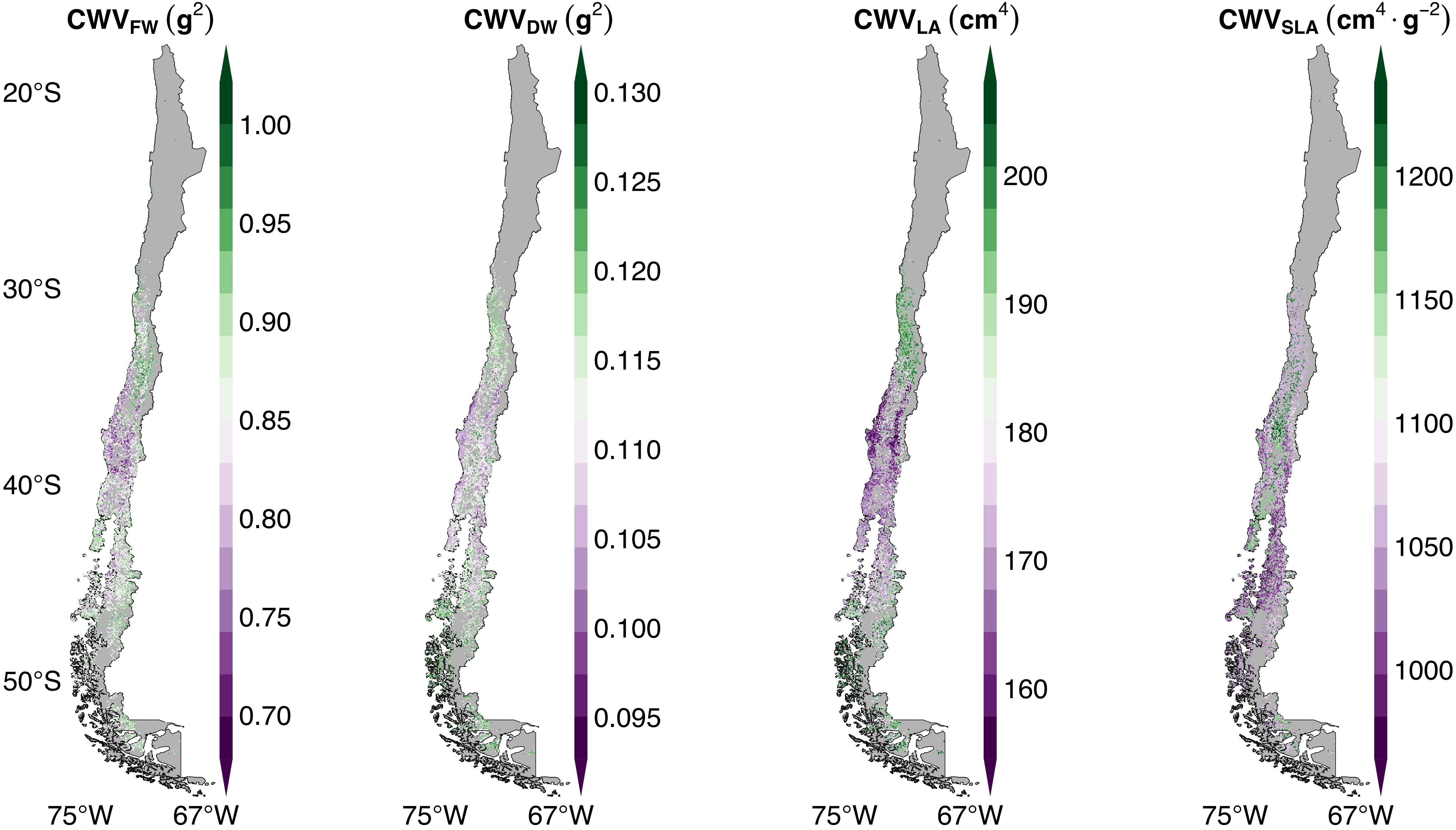


**Fig.69|Distribution maps of CWV of FW, DW, LA, and SLA.**

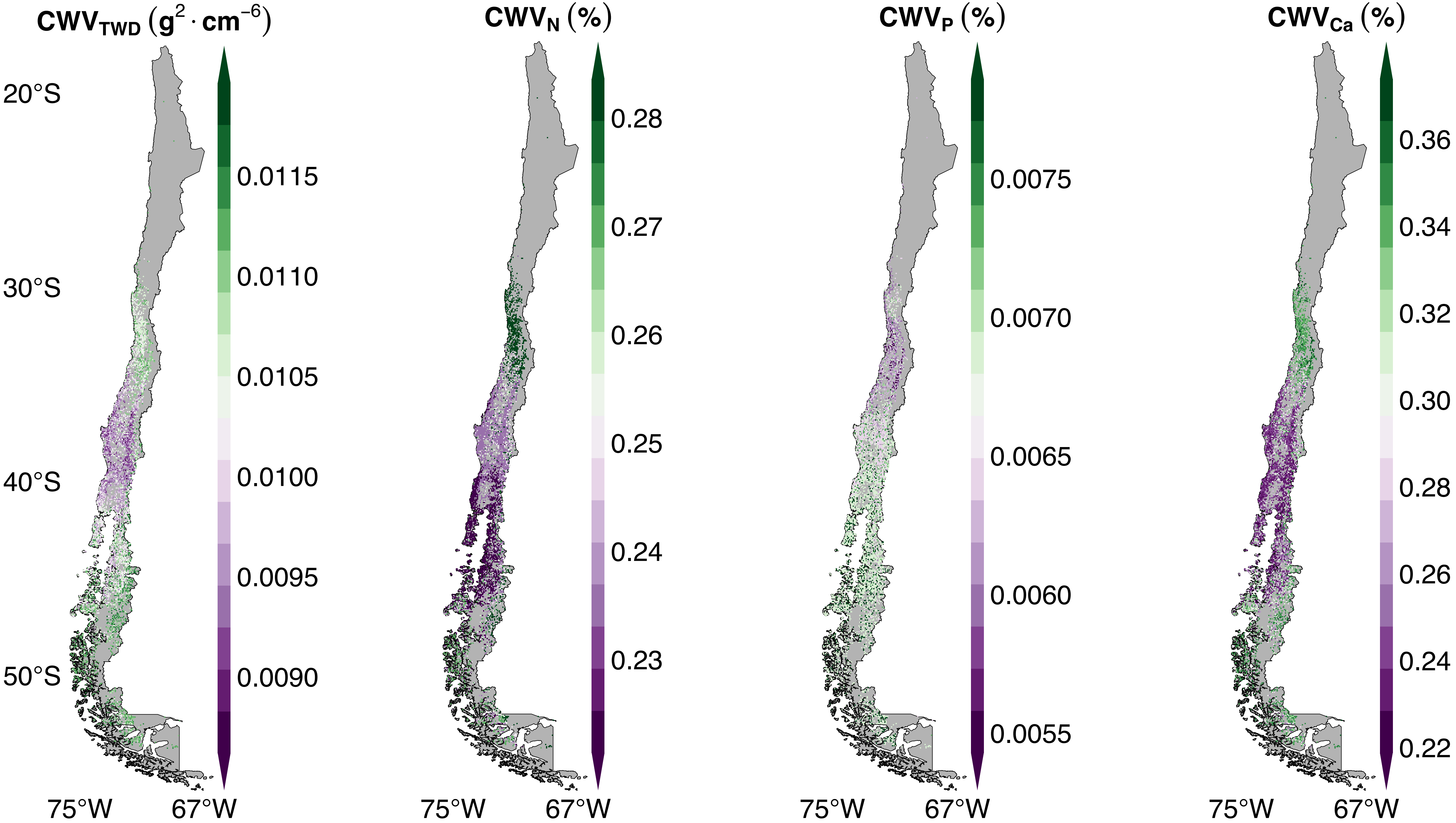


**Fig.70|Distribution maps of CWV of TWD, N, P, and Ca.**

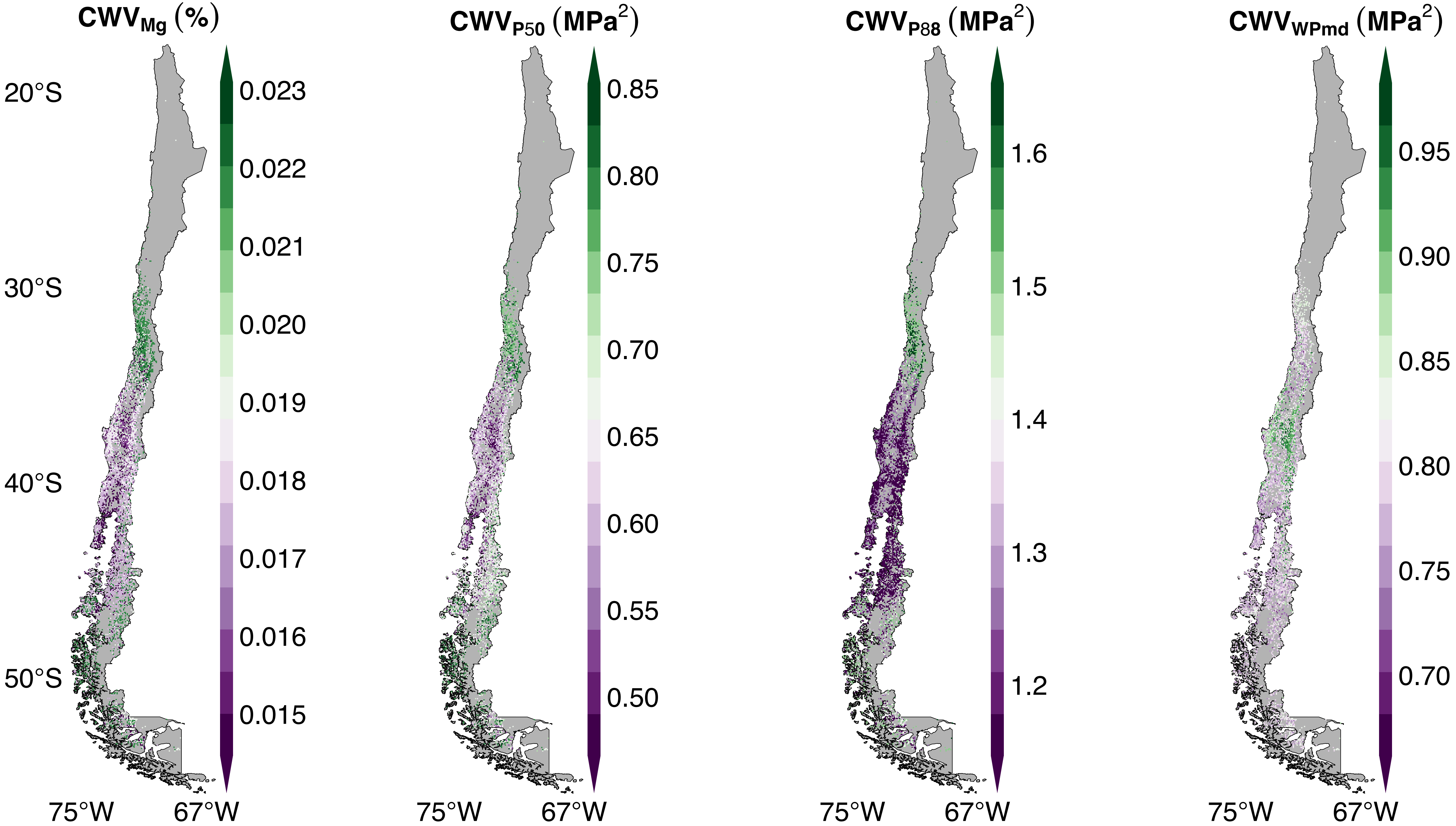


**Fig.71|Distribution maps of CWV of Mg, P50, P88, and WPmd.**

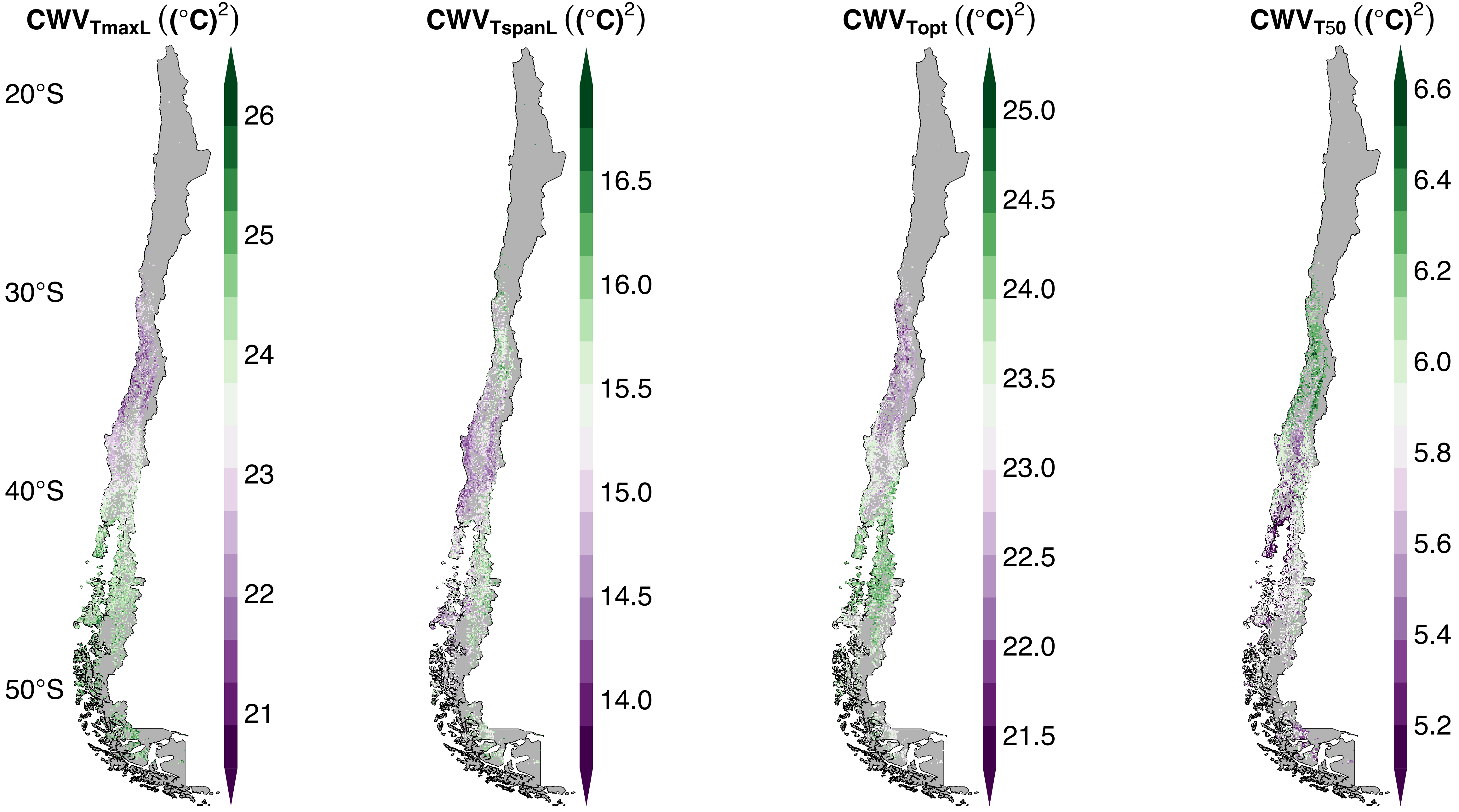


**Fig.72|Distribution maps of CWV of TmaxL, TspanL, Topt, and T50.**

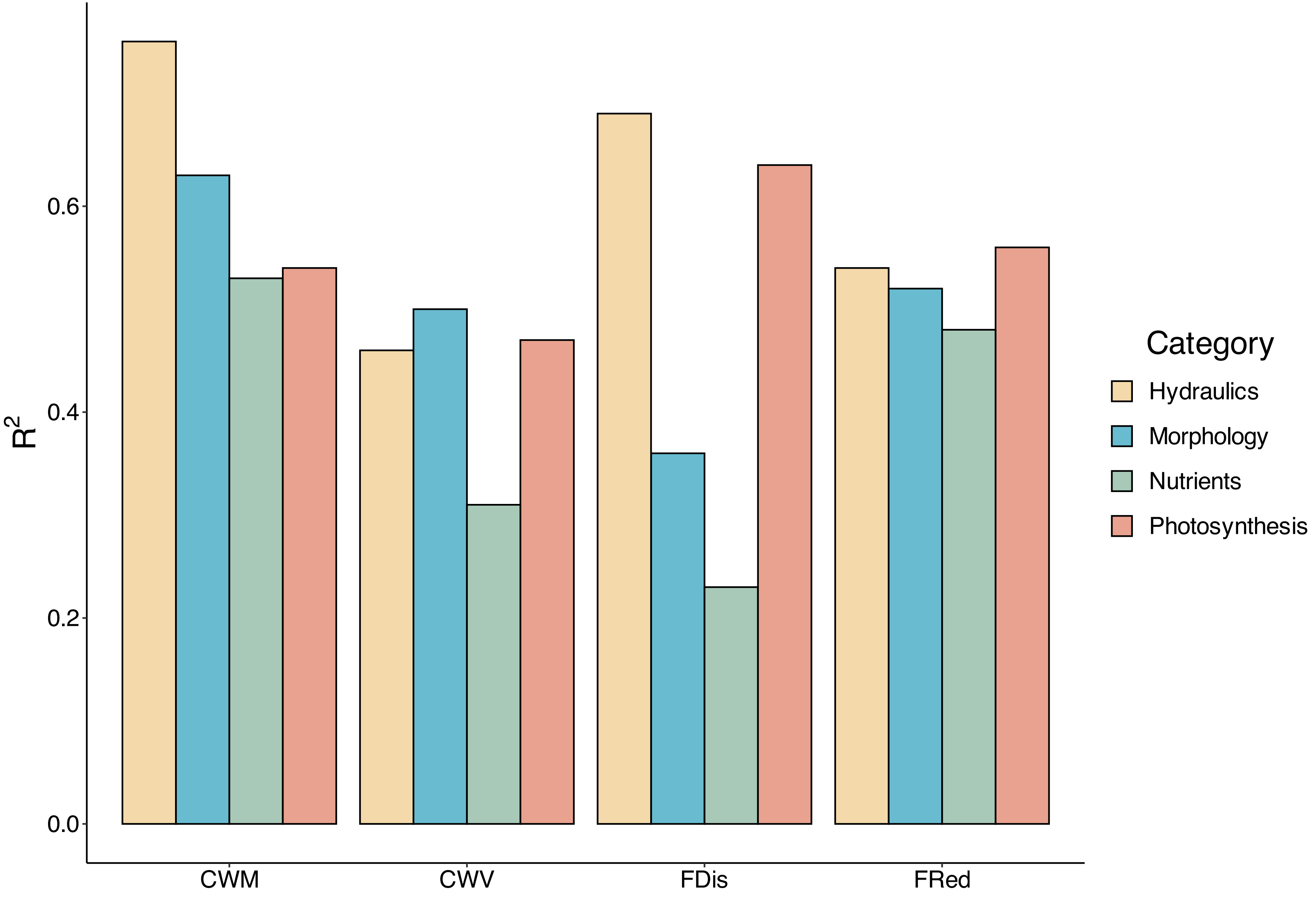


**Fig.73|Performance of Random forests regression models for predicting functional traits and functional dispersion and redundancy.** Bar plots of $R^2$ from models for assessing the two community-weighted moments of functinoal traits and functional dispersion and redundancy. Higher $R^2$ values (closer to 1) are generally desirable as they indicate a more accurate and reliable fit of the model to the observed data (all the input bands in this particular case), while a lower $R^2$ value (closer to 0) implies that the model does not effectively explain the variability in the dependent variable (a functional trait or functional dispersion/redundancy). Four different colours were applied to distinguish the four categories of functional traits.

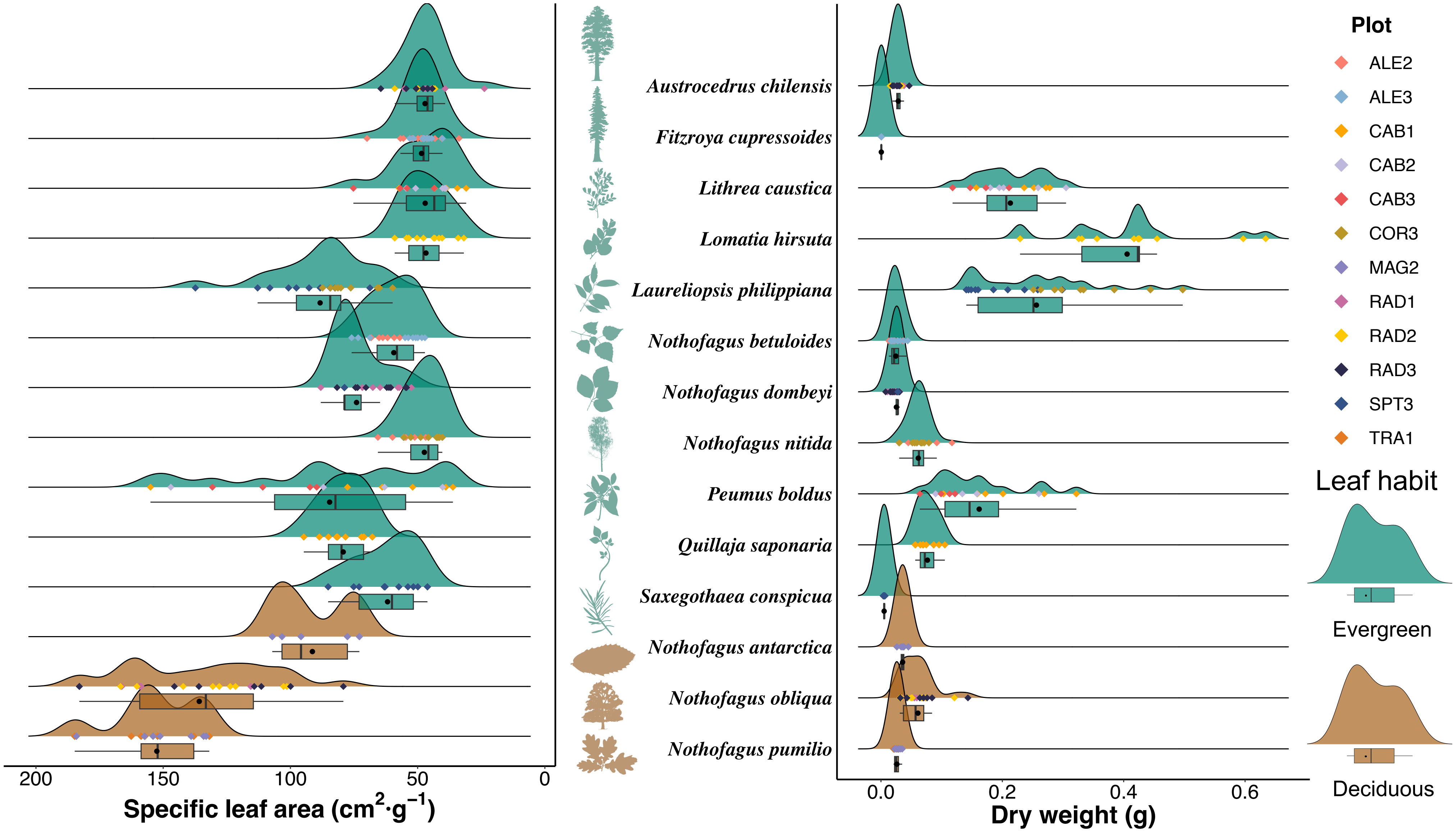


**Fig.74|Distributions of functional traits collected in the field across all plots in Chile.** Here we selected specific leaf area and leaf dry weight as examples. In the presented visualisation, a ridge plot integrates with its corresponding box plot situated below, offering an overall description of a trait distribution pattern of a certain species indicated by the middle italics and plant icons. The ridge plot affords insights into the shape and characteristics of the distribution and overall distributional trends, while the box plot offers trait statistical information and a grasp of central statistical tendencies and the presence of extreme values of the trait. The ridge plot employs kernel density estimation to represent the continuous distribution of a trait, revealing notable peaks and troughs, while the accompanying box plot consisting of a rectangular "box" and two whiskers indicates the distribution of the trait values. In a certain box, the black dot is the mean trait value, the three black vertical lines from the left to the right denote the first (25%) quartile, median, and the third (75%) quartile of a trait, and the two black horizontal whiskers extend to minimum (leftward) and maximum (rightward) trait values. Different colours of the areas under the ridge plot curves and the boxes are applied to differentiate the two main leaf habits across all plots in Chile. And the rhombuses with four right angles between the ridge plot and the box plot indicate the trait values of the species in different plots represented by different colours. Icon credit: The silhouettes were downloaded from PhyloPic ([https://www.phylopic.org/](https://www.phylopic.org/)) under Public Domain licences, X. Deng changed their colours to distinguish their corresponding habits.

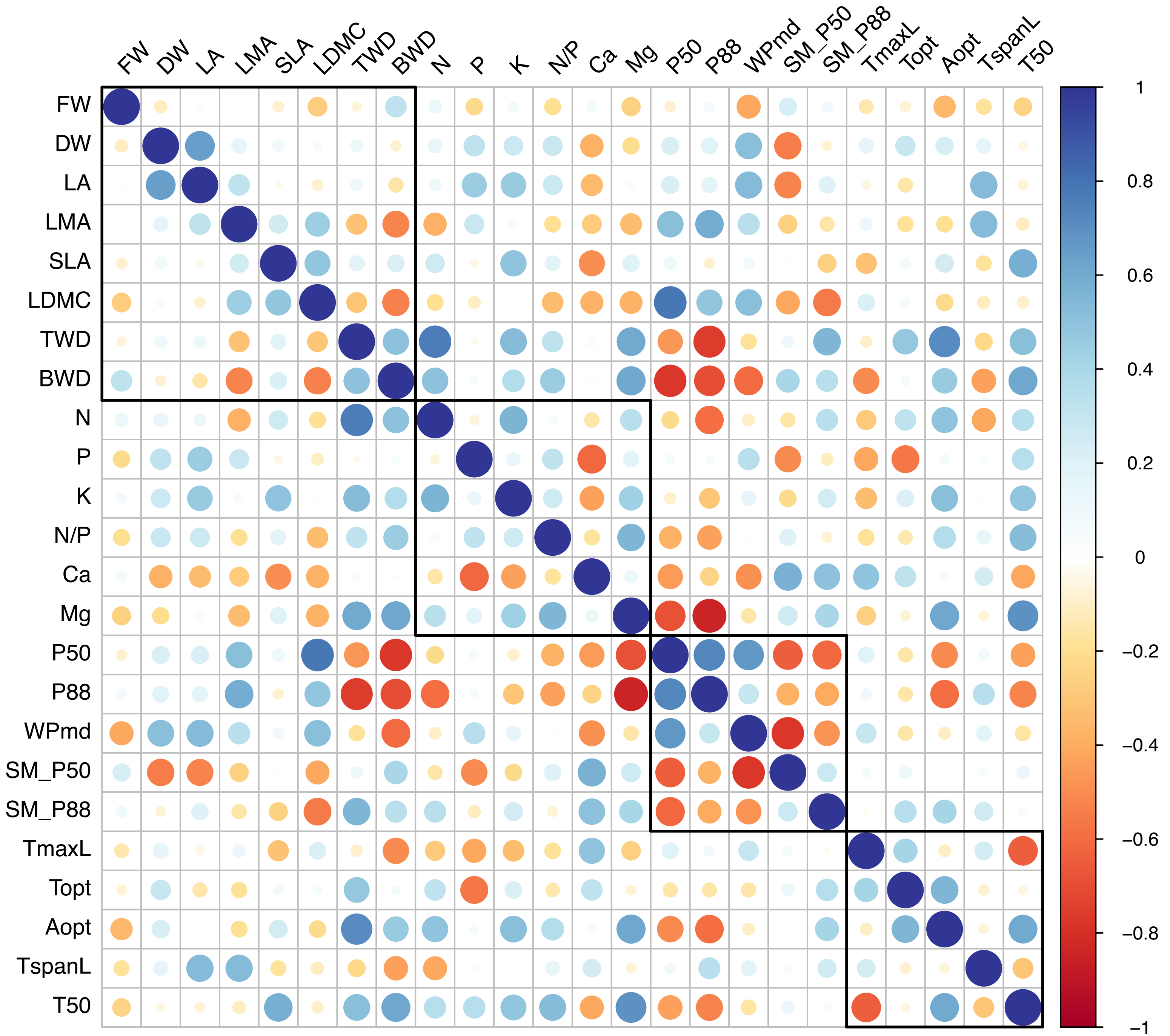


**Fig.75|Trait within-class correlation.**

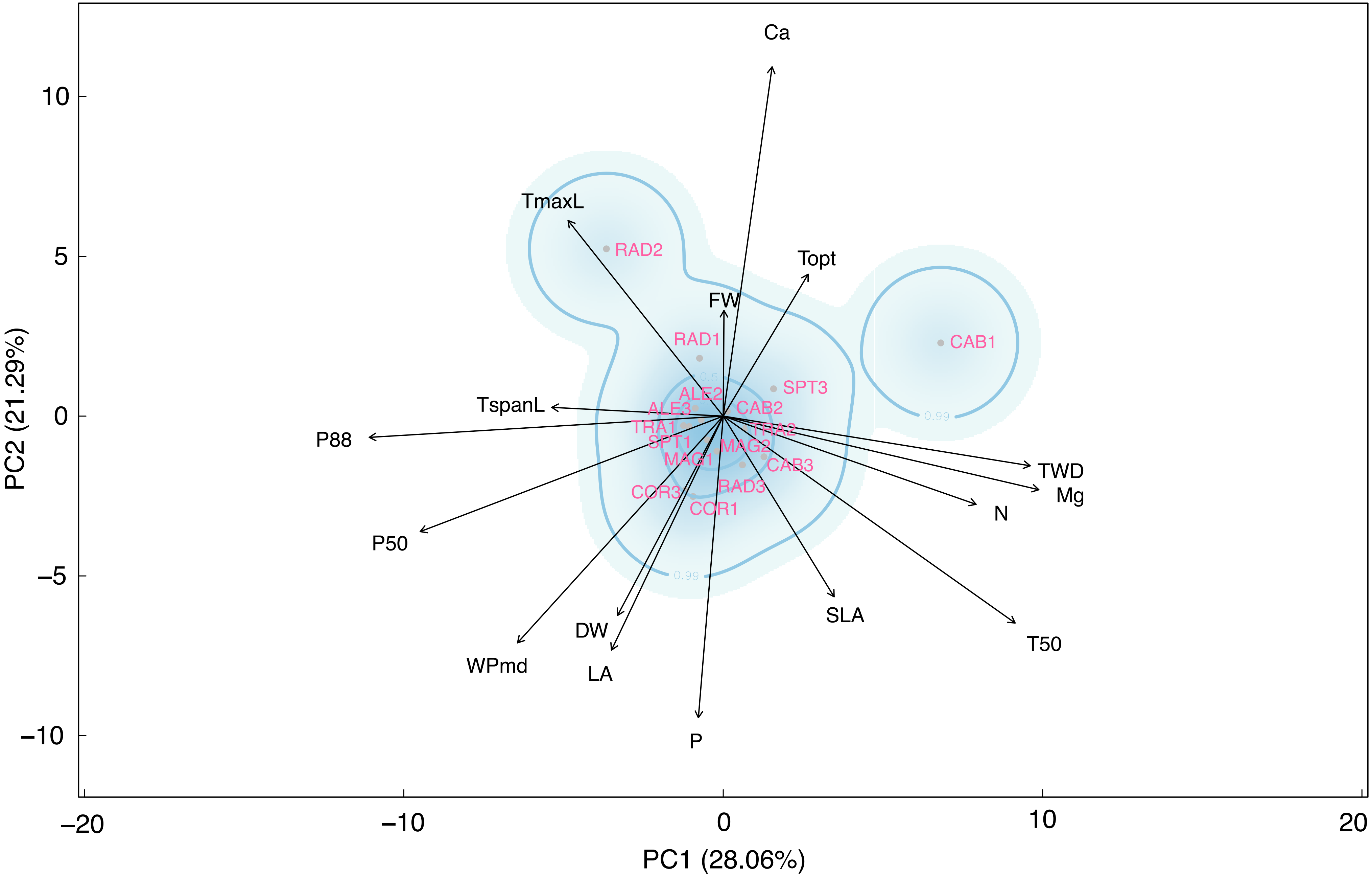


**Fig.76|Global functional trait space produced from PCA of CWM of 16 selected functional traits across all plots in Chile**. Grey dots represent distributions of these plots in the trait space with their corresponding plot IDs near them presenting in pink labels, contour boundaries and colour gradients stand for density of plots in the space, black arrows illustrate direction and weights of each trait along PC1 and PC2.

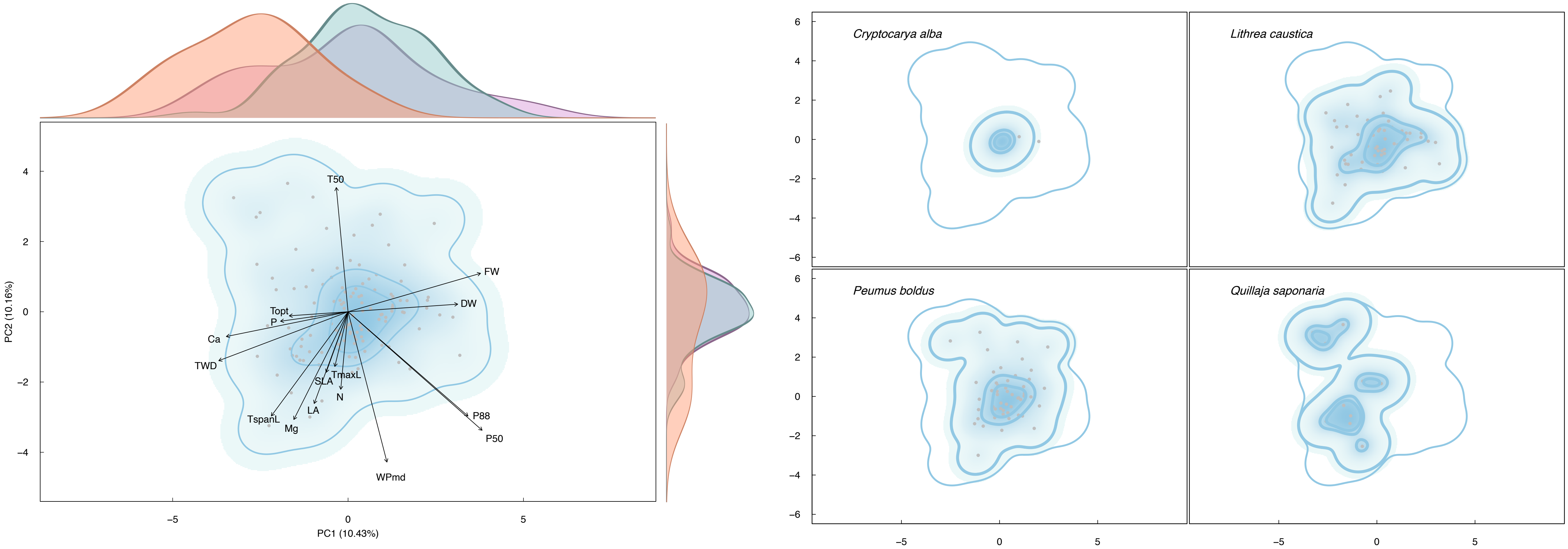


**Fig.77|Functional trait space (CAB1).**

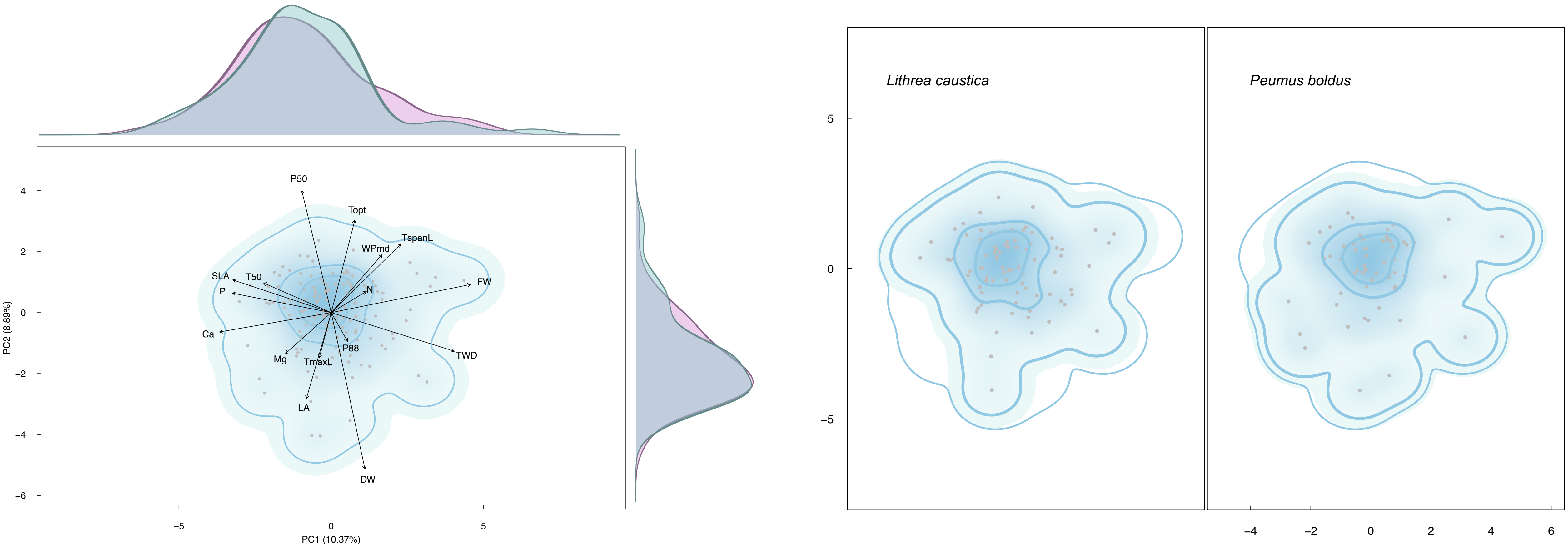


**Fig.78|Functional trait space (CAB2).**

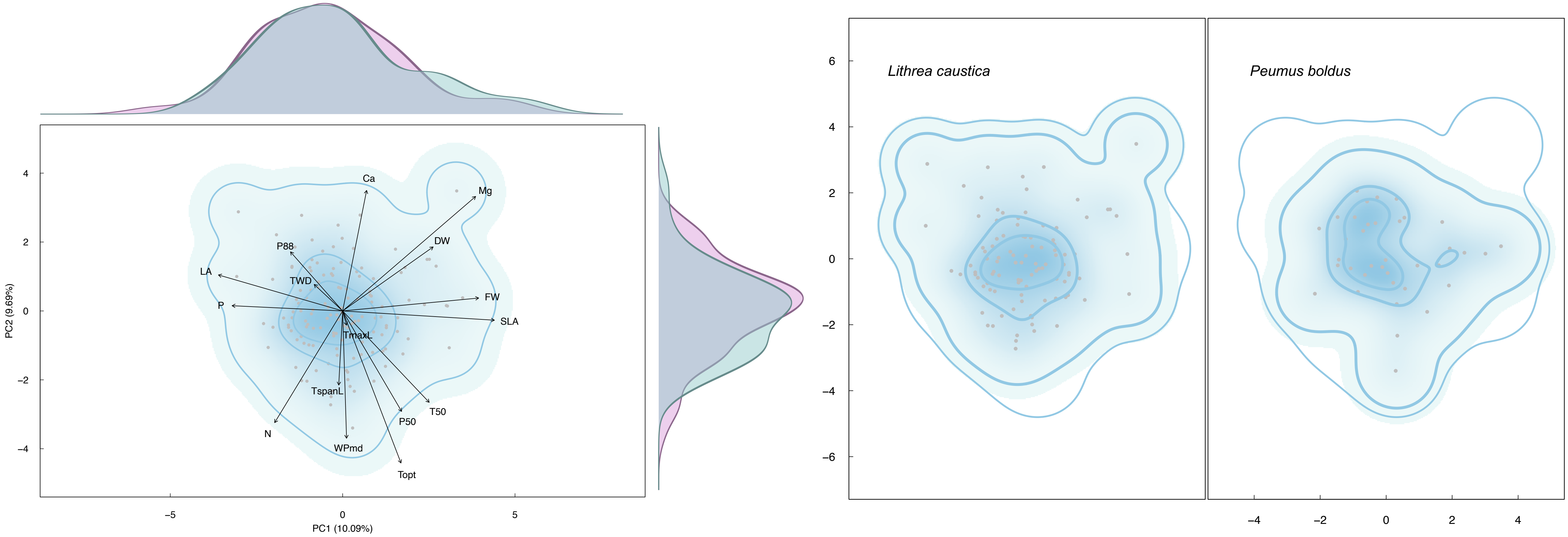


**Fig.79|Functional trait space (CAB3).**

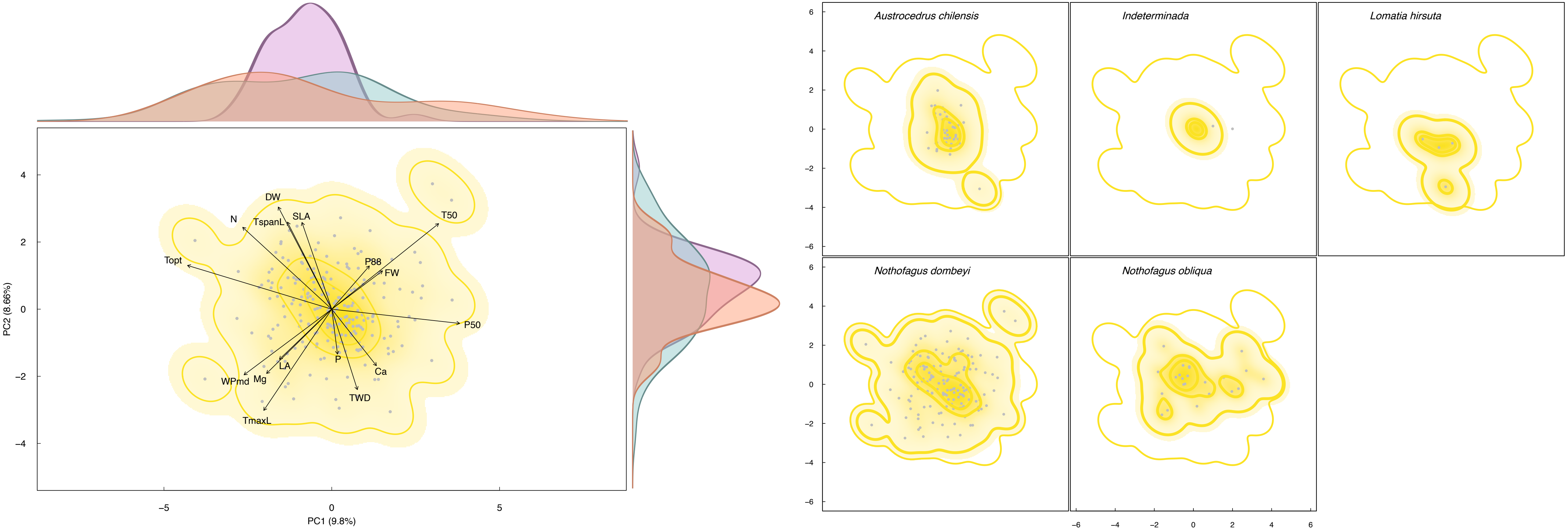


**Fig.80|Functional trait space (RAD3).**

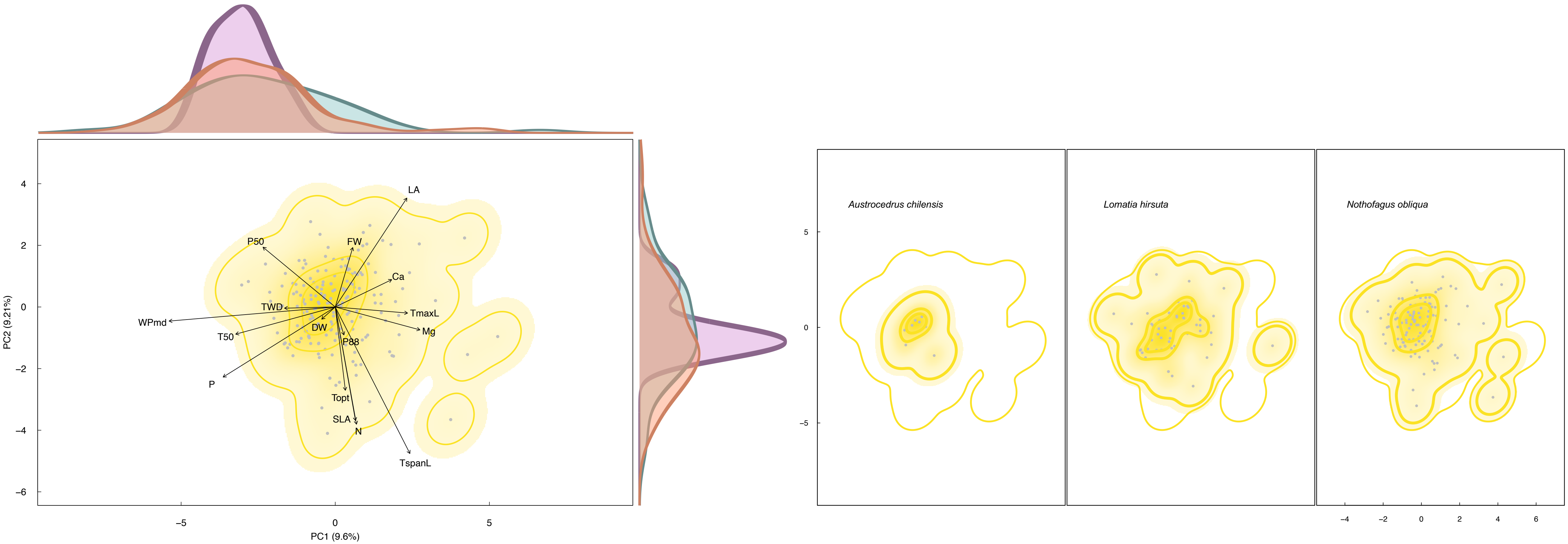


**Fig.81|Functional trait space (RAD2).**

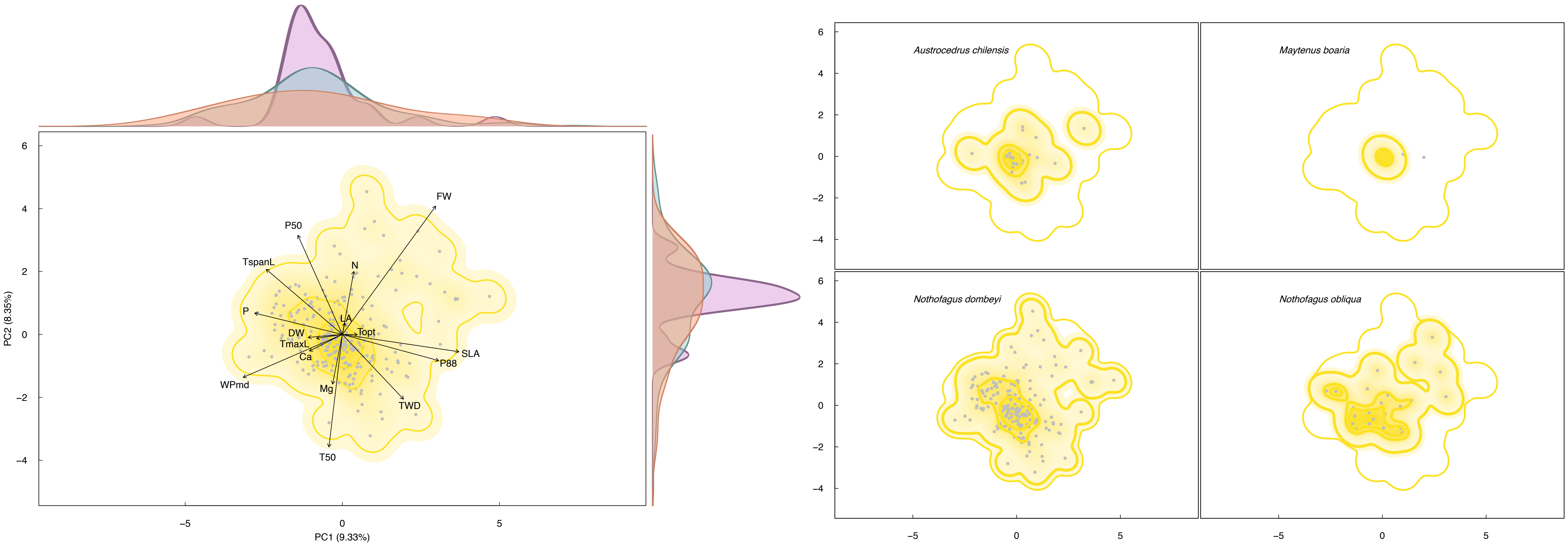


**Fig.82|Functional trait space (RAD1).**

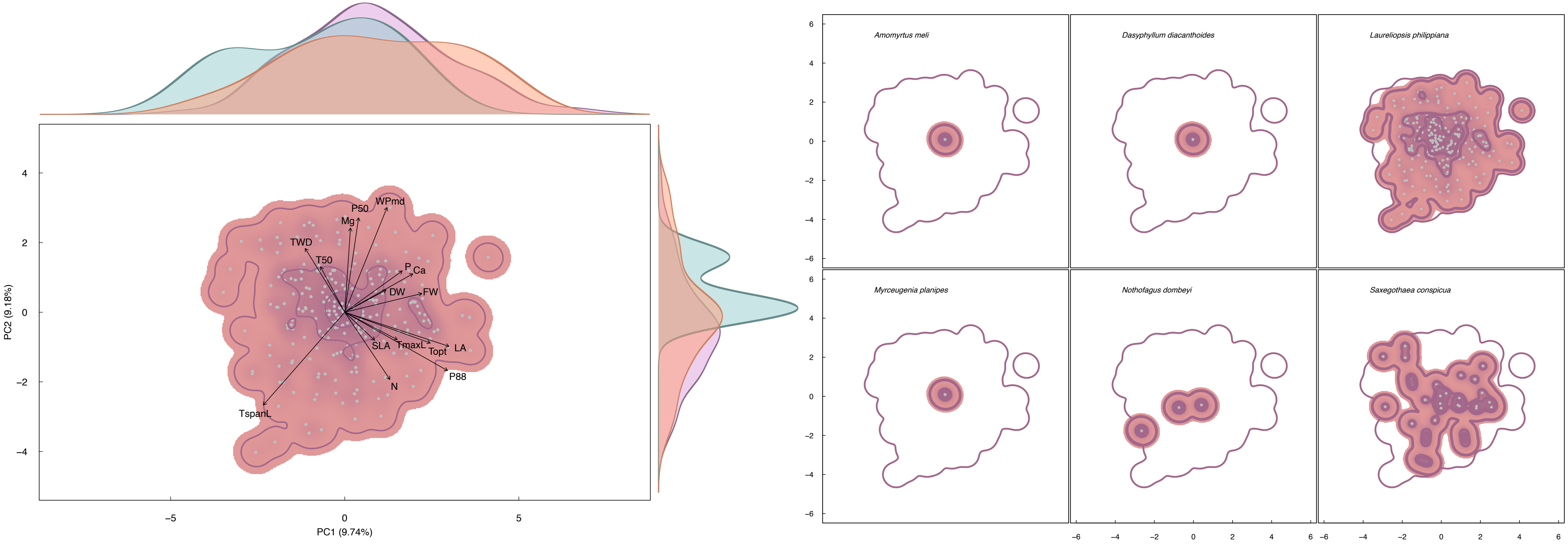


**Fig.83|Functional trait space (SPT1).**

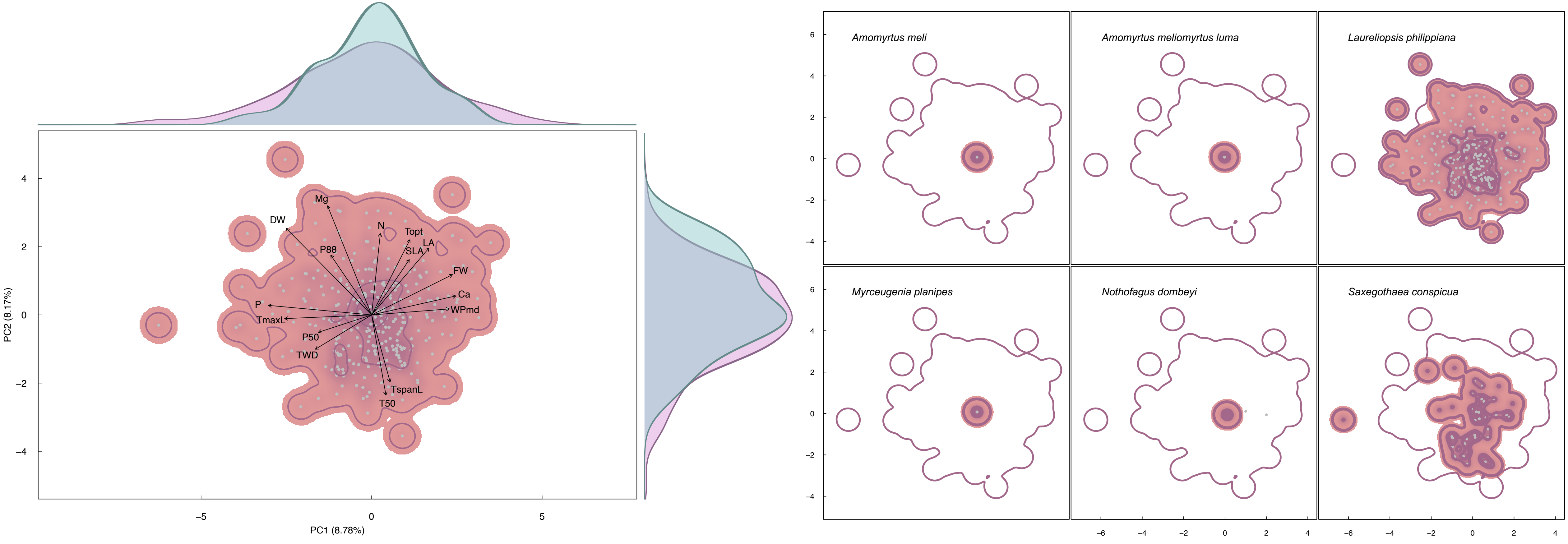


**Fig.84|Functional trait space (SPT3).**

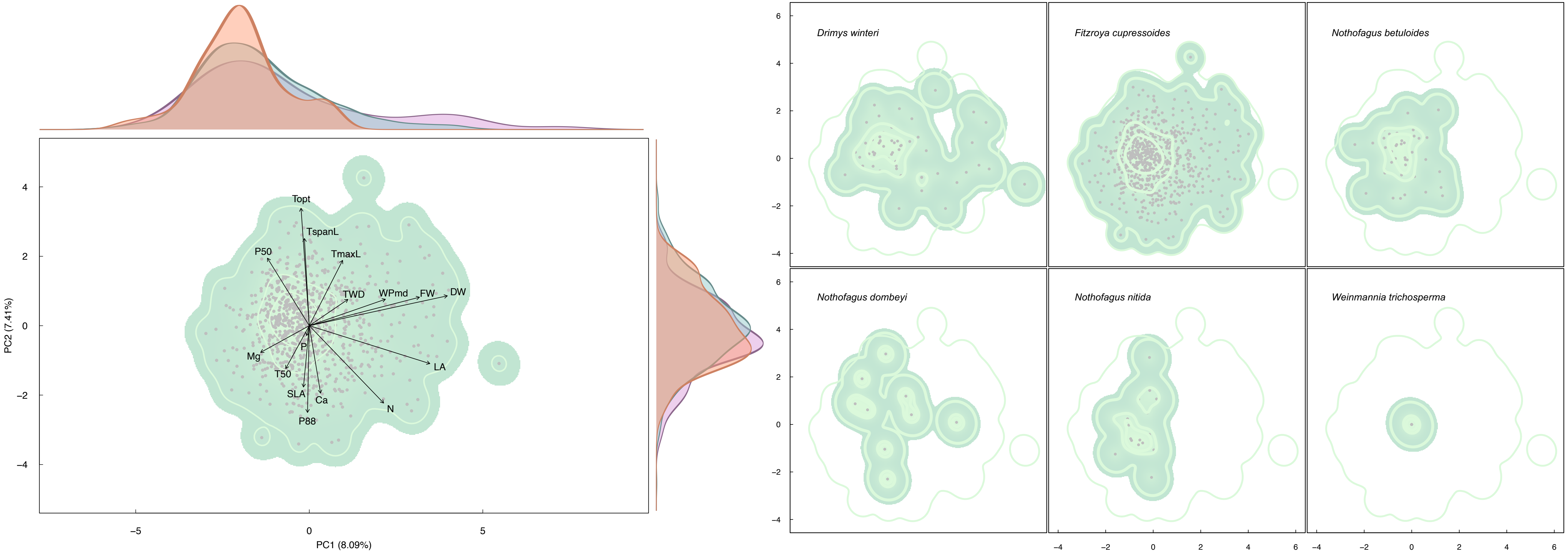


**Fig.85|Functional trait space (ALE2).**

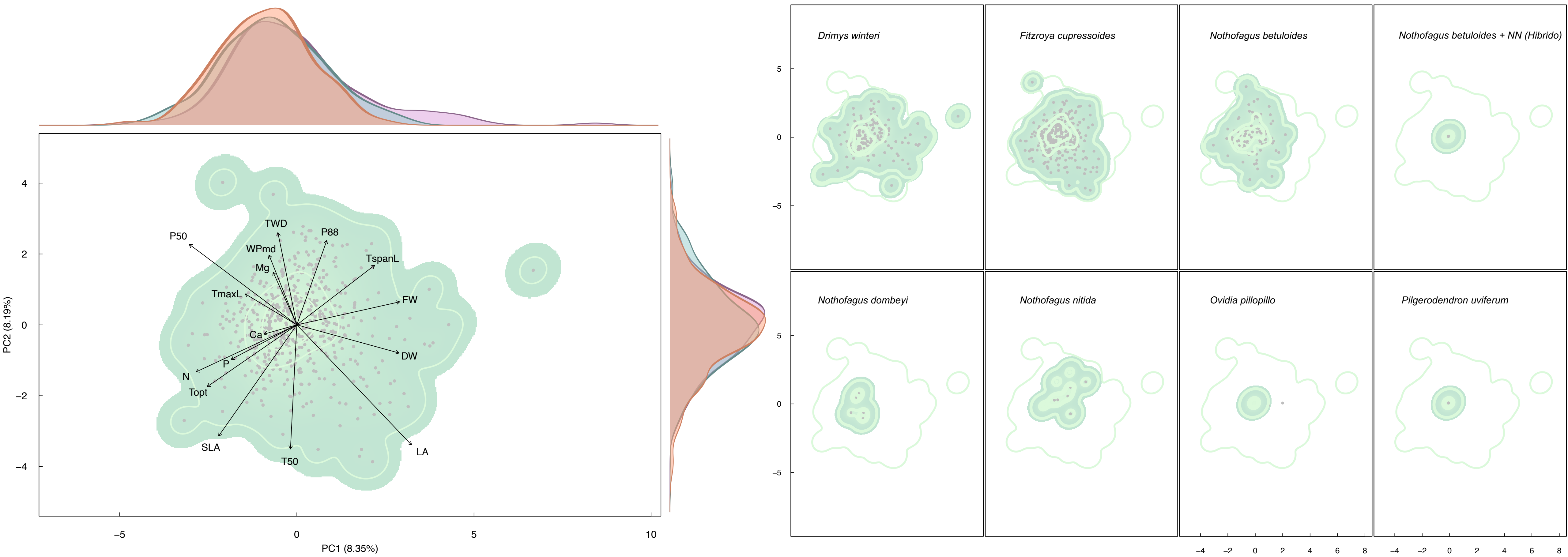


**Fig.86|Functional trait space (ALE3).**

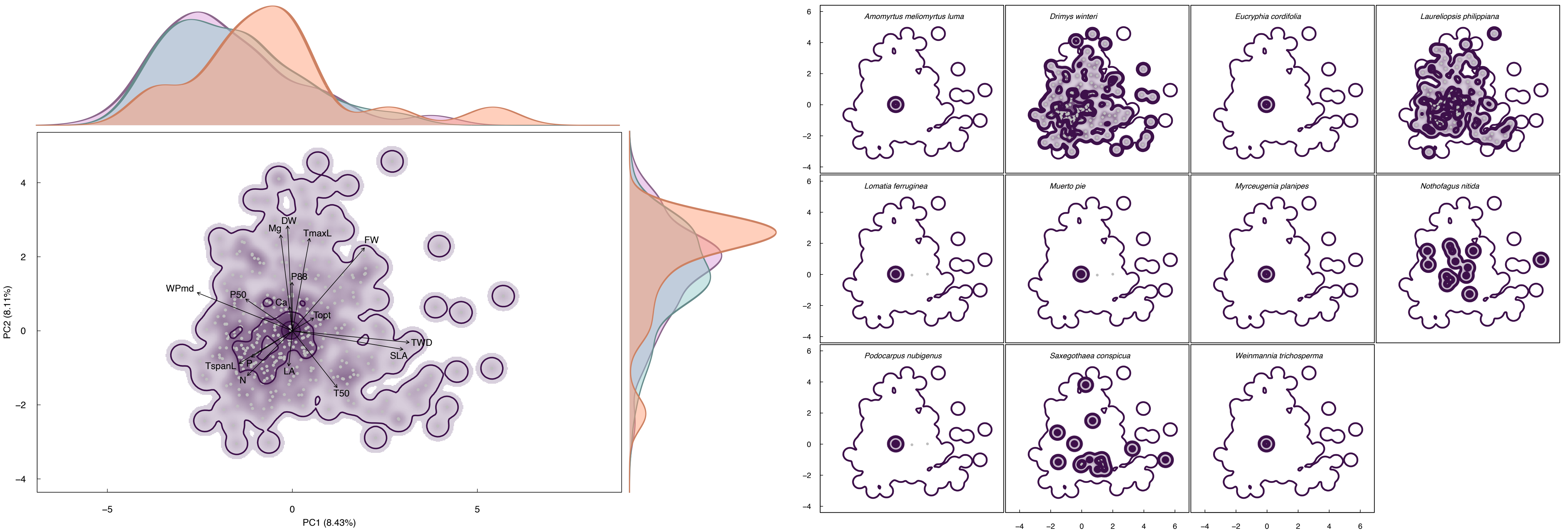


**Fig.87|Functional trait space (COR1).**

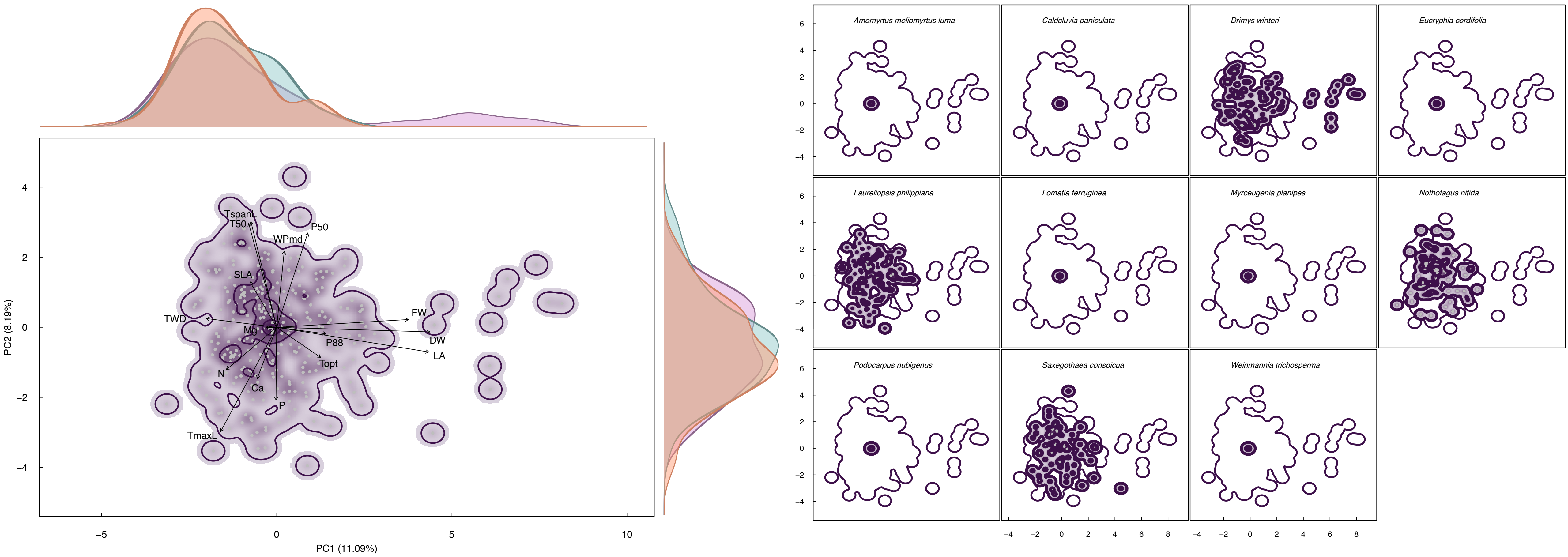


**Fig.88|Functional trait space (COR3).**

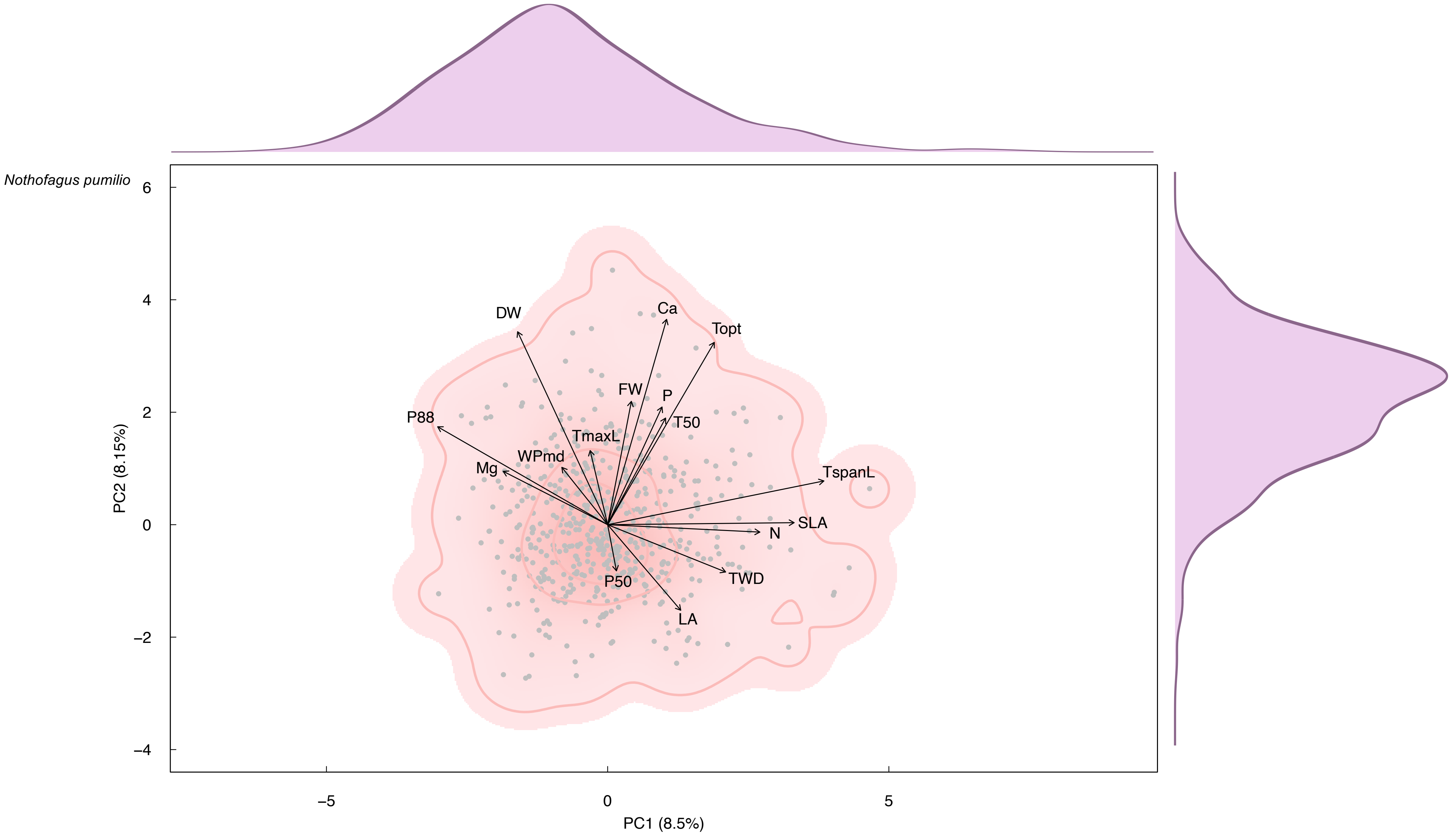


**Fig.89|Functional trait space (TRA1).**

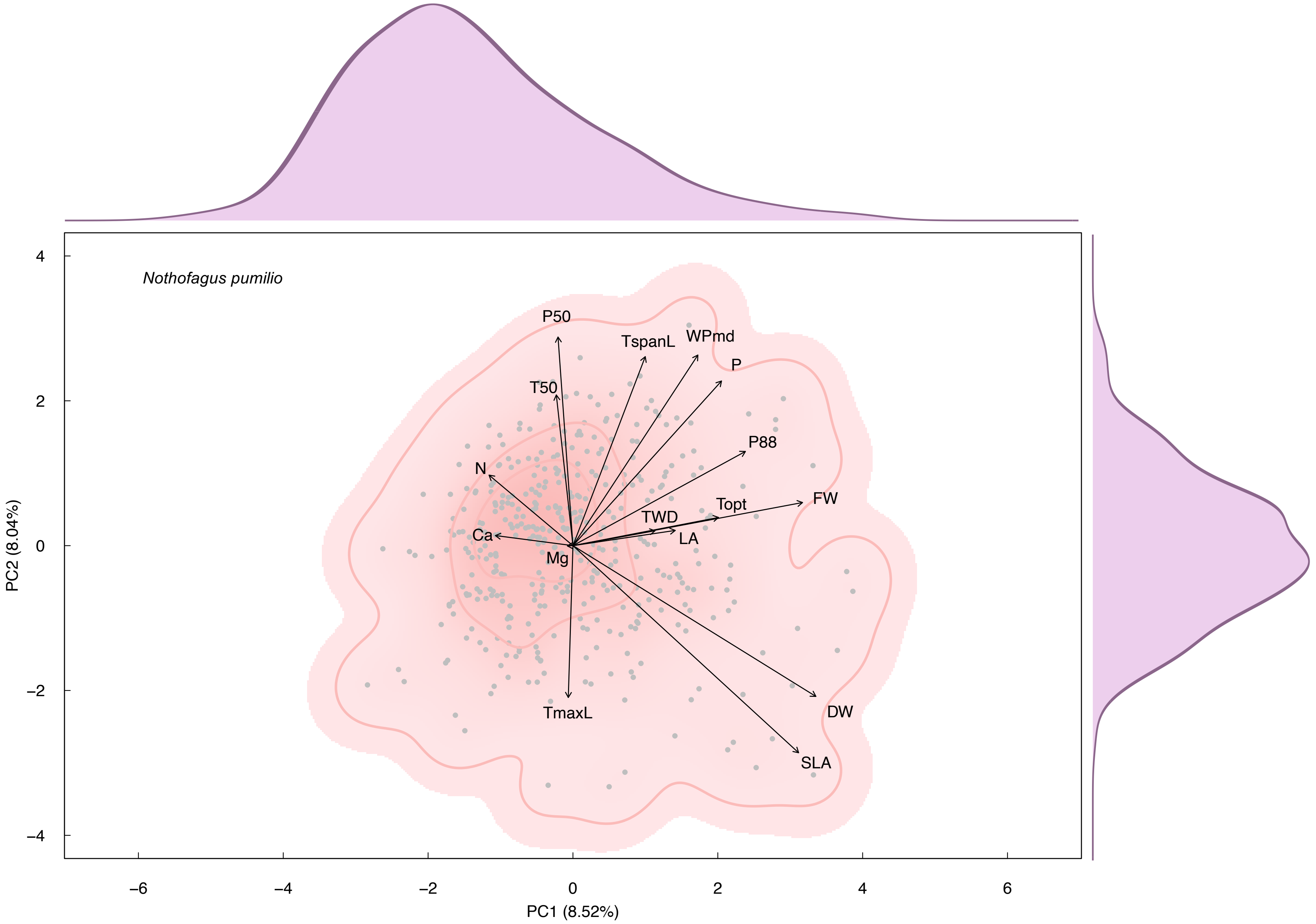


**Fig.90|Functional trait space (TRA2).**

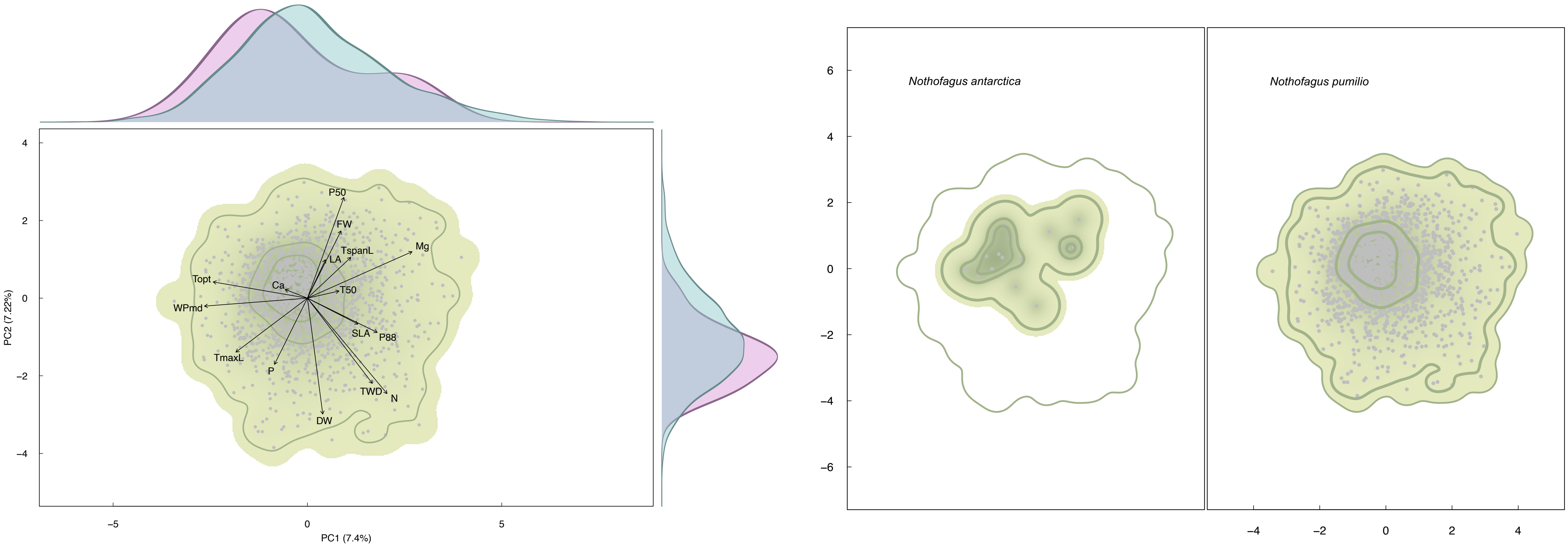


**Fig.91|Functional trait space (MAG1).**

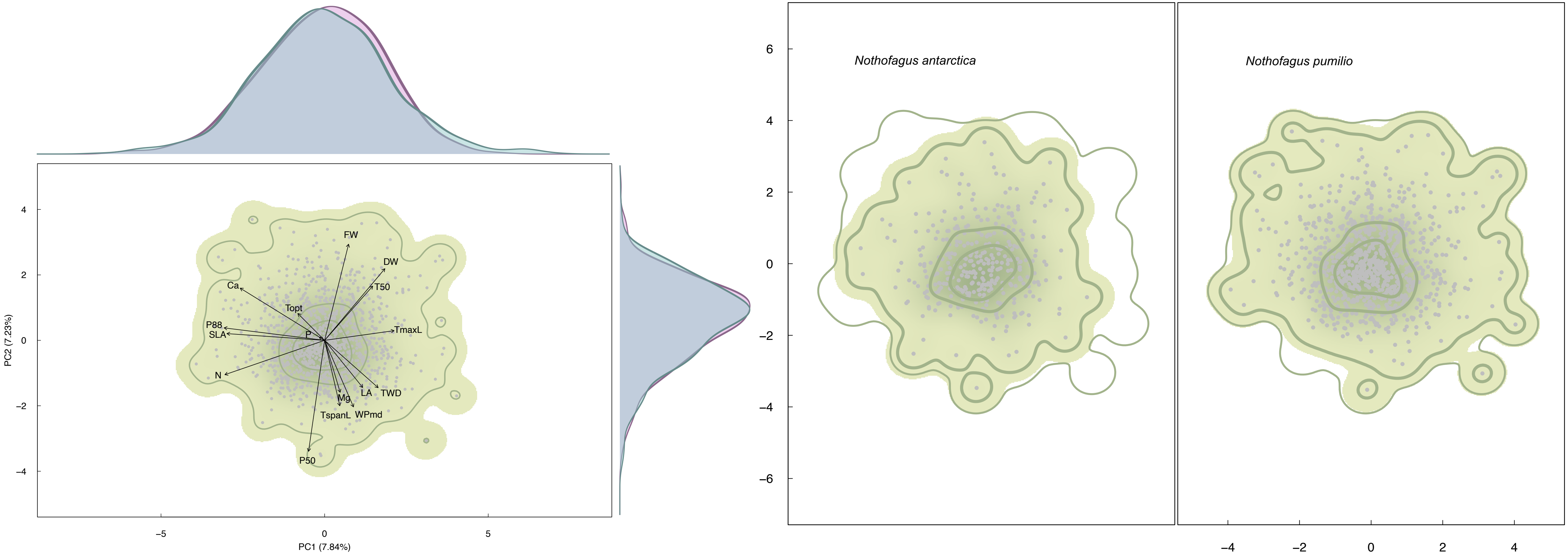


**Fig.92|Functional trait space (MAG2).**

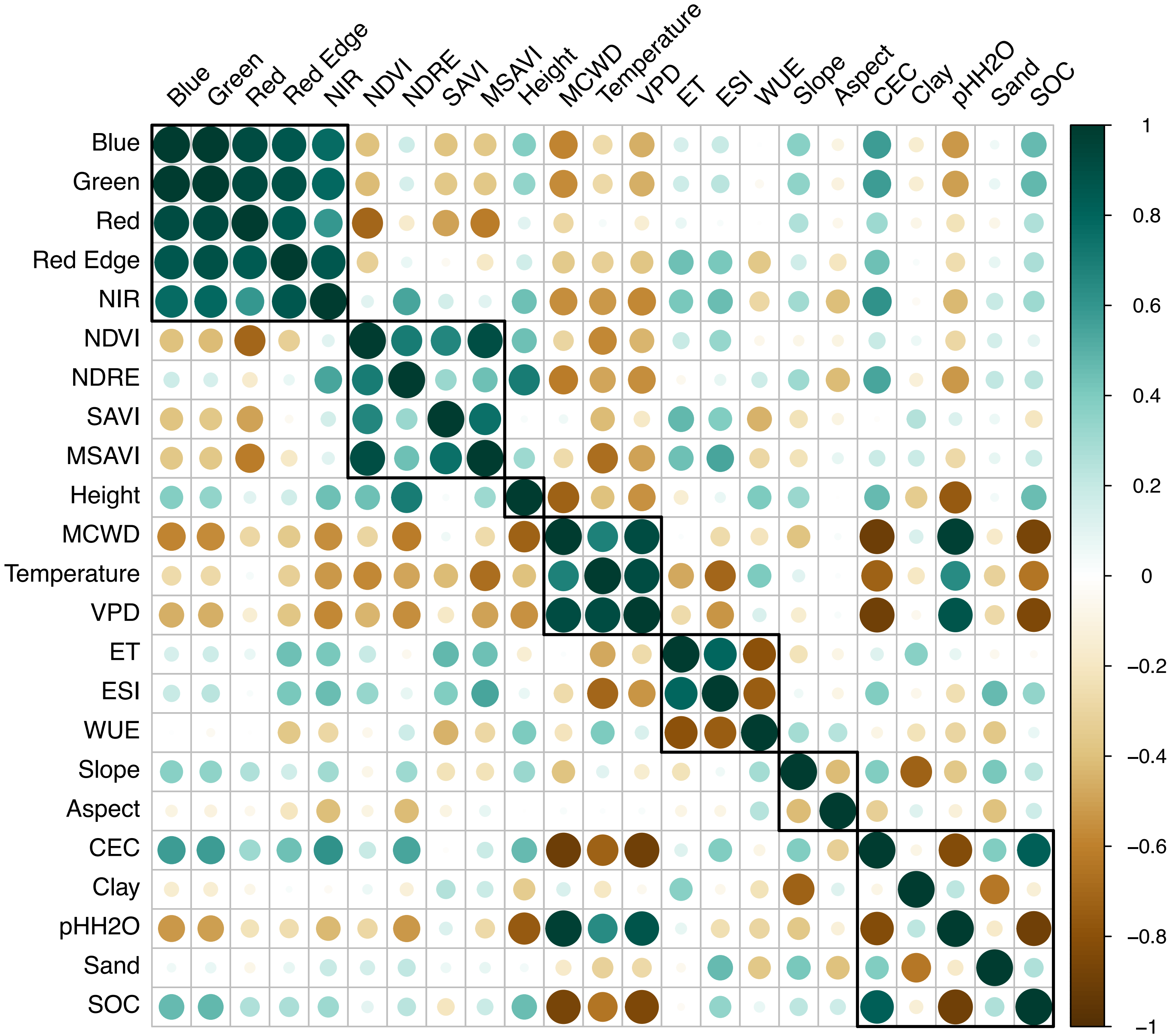


**Fig.93|Input band within-class correlation.**

**Table 1.** Model performance for mapping the two community-weighted moments of morphological traits. The bold numbers indicate the highest $R^2$ values.

| Site | Plot ID | Longitude (˚) | Latitude (˚) | Elevation (m) | Size (ha) |
|---|---|---|---|---|---|
| Las Cabras | CAB1 | -71.192 | -34.211 | 388.45 | 0.36 |
| Las Cabras | CAB2 | -71.193 | -34.212 | 386.41 | 0.36 |
| Las Cabras | CAB3 | -71.193 | -34.213 | 358.7 | 0.36 |
| Radal 7 tazas | RAD1 | -70.97 | -35.48 | 1233.53 | 0.36 |
| Radal 7 tazas | RAD2 | -70.969 | -35.479 | 1245.03 | 0.36 |
| Radal 7 tazas | RAD3 | -70.969 | -35.478 | 1244.595 | 0.36 |
| San Pablo de Tregua | SPT1 | -72.074 | -39.598 | 876.293 | 1 |
| San Pablo de Tregua | SPT3 | -72.098 | -39.604 | 774 | 1 |
| Alerce Costero | ALE2 | -73.445 | -40.171 | 820 | 0.6 |
| Alerce Costero | ALE3 | -73.443 | -40.173 | 841.586 | 0.6 |
| Correntoso | COR1 | -72.65 | -41.521 | 513.025 | 1 |
| Correntoso | COR3 | -72.646 | -41.522 | 442 | 1 |
| Trapananda National Reserve | TRA1 | -71.762 | -45.338 | 1102 | 1 |
| Trapananda National Reserve | TRA2 | -71.764 | -45.34 | 1108.75 | 1 |
| Magallanes National Reserve | MAG1 | -71.032 | -53.142 | 401.376 | 1 |
| Magallanes National Reserve | MAG2 | -71.026 | -53.143 | 392.472 | 1 |

**Table 2.** Correlation coefficients (R) and P-values (P) between each input band and CWM of morphological traits. P-values smaller than 0.05 are bold.

| Band \ Trait | FW | | DW | | LA | | SLA | | TWD | |
|---|---|---|---|---|---|---|---|---|---|---|
| | R | P | R | P | R | P | R | P | R | P |
| Red | 0.3170 | 0.2316 | 0.3067 | 0.2478 | 0.2342 | 0.3826 | -0.2212 | 0.4104 | 0.0480 | 0.8600 |
| Red Edge | 0.1377 | 0.6111 | 0.1147 | 0.6724 | 0.1296 | 0.6324 | 0.2321 | 0.3871 | -0.2019 | 0.4534 |
| NIR | 0.2053 | 0.4456 | 0.1544 | 0.5681 | 0.2043 | 0.4480 | 0.4108 | 0.1140 | -0.3726 | 0.1552 |
| NDVI | -0.1963 | 0.4663 | -0.2681 | 0.3155 | -0.1122 | 0.6790 | 0.5667 | **0.0221** | -0.5517 | **0.0267** |
| NDRE | 0.3008 | 0.2576 | 0.2190 | 0.4152 | 0.2988 | 0.2609 | 0.3752 | 0.1522 | -0.3740 | 0.1535 |
| SAVI | -0.3165 | 0.2324 | -0.3299 | 0.2121 | -0.1856 | 0.4913 | 0.6447 | **0.0070** | -0.2810 | 0.2918 |
| MSAVI | -0.4151 | 0.1099 | -0.4844 | 0.0573 | -0.3230 | 0.2224 | 0.5969 | **0.0146** | -0.7001 | **0.0025** |
| Height | 0.5247 | **0.0369** | 0.3932 | 0.1319 | 0.4788 | 0.0606 | 0.1034 | 0.7030 | -0.5346 | **0.0329** |
| MCWD | -0.1976 | 0.4633 | -0.0872 | 0.7482 | -0.0915 | 0.7361 | -0.0636 | 0.8149 | 0.6895 | **0.0031** |
| Temperature | 0.3790 | 0.1477 | 0.4679 | 0.0676 | 0.3550 | 0.1773 | -0.6729 | **0.0043** | 0.7922 | **0.0003** |
| ET | -0.4569 | 0.0752 | -0.4680 | 0.0675 | -0.3804 | 0.1461 | 0.5256 | **0.0365** | -0.4345 | 0.0926 |
| ESI | -0.3360 | 0.2033 | -0.3600 | 0.1708 | -0.2974 | 0.2633 | 0.5956 | **0.0149** | -0.6032 | **0.0134** |
| WUE | 0.5066 | **0.0452** | 0.4587 | 0.0739 | 0.4476 | 0.0821 | -0.5926 | **0.0156** | 0.0918 | 0.7352 |
| Slope | 0.6125 | **0.0117** | 0.6252 | **0.0096** | 0.4869 | 0.0558 | -0.3520 | 0.1811 | -0.0436 | 0.8727 |
| Aspect | -0.3139 | 0.2364 | -0.3759 | 0.1513 | -0.3149 | 0.2349 | -0.2013 | 0.4547 | -0.3197 | 0.2273 |
| CEC | 0.0938 | 0.7297 | 0.0331 | 0.9032 | -0.0169 | 0.9505 | 0.1656 | 0.5400 | -0.5116 | **0.0428** |
| Clay | -0.6144 | **0.0113** | -0.6250 | **0.0096** | -0.5292 | **0.0351** | 0.2382 | 0.3744 | -0.0486 | 0.8582 |
| $pHH_2O$ | -0.2020 | 0.4532 | -0.0846 | 0.7555 | -0.0891 | 0.7427 | 0.0069 | 0.9799 | 0.7195 | **0.0017** |
| Sand | 0.1012 | 0.7093 | 0.1393 | 0.6068 | 0.0716 | 0.7922 | 0.3312 | 0.2102 | -0.0188 | 0.9448 |
| SOC | -0.0183 | 0.9465 | -0.0983 | 0.7173 | -0.1602 | 0.5534 | -0.0639 | 0.8142 | -0.6205 | **0.0103** |

**Table 3.** Correlation coefficients (R) and P-values (P) between each input band and CWM of nutrient traits. P-values smaller than 0.05 are bold.

| Band \ Trait | N | | P | | Ca | | Mg | |
|---|---|---|---|---|---|---|---|---|
| | R | P | R | P | R | P | R | P |
| Red | -0.2426 | 0.3653 | -0.2589 | 0.3330 | -0.1195 | 0.6593 | -0.1473 | 0.5861 |
| Red Edge | 0.1971 | 0.4644 | 0.1977 | 0.4629 | 0.1677 | 0.5347 | 0.1478 | 0.5848 |
| NIR | 0.4222 | 0.1033 | 0.3531 | 0.1797 | 0.2889 | 0.2778 | 0.3475 | 0.1872 |
| NDVI | 0.6137 | **0.0114** | 0.5318 | **0.0340** | 0.3021 | 0.2554 | 0.4205 | 0.1049 |
| NDRE | 0.4759 | 0.0624 | 0.2967 | 0.2645 | 0.3131 | 0.2377 | 0.4610 | 0.0723 |
| SAVI | 0.6907 | **0.0031** | 0.6741 | **0.0042** | 0.3625 | 0.1677 | 0.4869 | 0.0558 |
| MSAVI | 0.5983 | **0.0143** | 0.5786 | **0.0189** | 0.2173 | 0.4188 | 0.2912 | 0.2739 |
| Height | 0.2129 | 0.4285 | -0.0192 | 0.9437 | -0.0619 | 0.8200 | 0.2194 | 0.4143 |
| MCWD | -0.0915 | 0.7361 | 0.0324 | 0.9053 | 0.0215 | 0.9369 | 0.0702 | 0.7961 |
| Temperature | -0.6166 | **0.0110** | -0.6079 | **0.0125** | -0.4521 | 0.0787 | -0.2964 | 0.2650 |
| ET | 0.4672 | 0.0681 | 0.5389 | **0.0312** | 0.3077 | 0.2463 | 0.3132 | 0.2376 |
| ESI | 0.5050 | **0.0460** | 0.5826 | **0.0179** | 0.3634 | 0.1666 | 0.2716 | 0.3089 |
| WUE | -0.4734 | 0.0640 | -0.6271 | **0.0093** | -0.5168 | **0.0404** | -0.3519 | 0.1813 |
| Slope | -0.2647 | 0.3218 | -0.3597 | 0.1712 | -0.2727 | 0.3068 | -0.1453 | 0.5914 |
| Aspect | -0.2718 | 0.3084 | -0.2403 | 0.3699 | -0.2847 | 0.2853 | -0.3015 | 0.2565 |
| CEC | 0.1600 | 0.5538 | 0.1054 | 0.6976 | 0.1493 | 0.5811 | -0.0321 | 0.9060 |
| Clay | 0.2030 | 0.4507 | 0.2682 | 0.3152 | 0.1826 | 0.4986 | 0.0373 | 0.8908 |
| $pHH_2O$ | -0.0117 | 0.9656 | 0.1096 | 0.6862 | 0.1107 | 0.6831 | 0.1470 | 0.5870 |
| Sand | 0.2720 | 0.3082 | 0.3251 | 0.2192 | 0.4027 | 0.1220 | 0.2318 | 0.3877 |
| SOC | -0.1289 | 0.6342 | -0.1387 | 0.6085 | -0.1272 | 0.6387 | -0.3514 | 0.1820 |

**Table 4.** Correlation coefficients (R) and P-values (P) between each input band and CWM of hydraulic traits. P-values smaller than 0.05 are bold.

| Trait / Band | P50 | | P88 | | WPmd | |
|---|---|---|---|---|---|---|
| | R | P | R | P | R | P |
| Red | -0.0643 | 0.8131 | -0.1394 | 0.6066 | 0.0047 | 0.9863 |
| Red Edge | 0.3180 | 0.2301 | 0.3081 | 0.2457 | 0.2792 | 0.2949 |
| NIR | 0.5853 | **0.0172** | 0.5493 | 0.0275 | 0.5033 | **0.0469** |
| NDVI | 0.6040 | **0.0132** | 0.6714 | **0.0044** | 0.5943 | **0.0152** |
| NDRE | 0.5859 | **0.0171** | 0.5069 | **0.0451** | 0.5345 | **0.0329** |
| SAVI | 0.5696 | **0.0213** | 0.6427 | **0.0072** | 0.3176 | 0.2307 |
| MSAVI | 0.6140 | **0.0114** | 0.7483 | **0.0009** | 0.6875 | **0.0033** |
| Height | 0.5934 | **0.0154** | 0.4651 | 0.0695 | 0.6692 | **0.0046** |
| MCWD | -0.3695 | 0.1590 | -0.3535 | 0.1792 | -0.7982 | **0.0002** |
| Temperature | -0.6160 | **0.0111** | -0.7747 | **0.0004** | -0.8749 | **0.0000** |
| ET | 0.3849 | 0.1410 | 0.5665 | **0.0221** | 0.3400 | 0.1975 |
| ESI | 0.5397 | **0.0310** | 0.6972 | **0.0027** | 0.5964 | **0.0148** |
| WUE | -0.2908 | 0.2745 | -0.4502 | 0.0802 | -0.0850 | 0.7544 |
| Slope | 0.0714 | 0.7928 | -0.1113 | 0.6817 | 0.1270 | 0.6393 |
| Aspect | -0.2510 | 0.3484 | -0.1312 | 0.6281 | 0.1332 | 0.6230 |
| CEC | 0.3407 | 0.1965 | 0.3419 | 0.1949 | 0.6669 | **0.0048** |
| Clay | -0.1263 | 0.6412 | 0.0646 | 0.8121 | -0.0244 | 0.9286 |
| $pHH_2O$ | -0.3317 | 0.2094 | -0.3154 | 0.2341 | -0.8076 | **0.0002** |
| Sand | 0.2963 | 0.2651 | 0.2638 | 0.3235 | 0.1740 | 0.5194 |
| SOC | 0.1439 | 0.5950 | 0.1882 | 0.4852 | 0.6962 | **0.0027** |

**Table 5.** Correlation coefficients (R) and P-values (P) between each input band and CWM of photosynthetic traits. P-values smaller than 0.05 are bold.

| Band \ Trait | TmaxL | | TspanL | | Topt | | T50 | |
|---|---|---|---|---|---|---|---|---|
| | R | P | R | P | R | P | R | P |
| Red | -0.0583 | 0.8302 | -0.1877 | 0.4863 | 0.0438 | 0.8721 | 0.2432 | 0.3642 |
| Red Edge | -0.0339 | 0.9007 | 0.1707 | 0.5275 | -0.0768 | 0.7775 | 0.0331 | 0.9033 |
| NIR | -0.1884 | 0.4848 | 0.1709 | 0.5269 | -0.2572 | 0.3363 | 0.1015 | 0.7082 |
| NDVI | -0.3270 | 0.2164 | 0.2861 | 0.2827 | -0.4629 | 0.0710 | -0.1509 | 0.5769 |
| NDRE | -0.3509 | 0.1826 | -0.0986 | 0.7163 | -0.3484 | 0.1860 | 0.3720 | 0.1560 |
| SAVI | 0.0655 | 0.8094 | 0.4717 | 0.0651 | -0.1411 | 0.6021 | -0.3620 | 0.1682 |
| MSAVI | -0.3600 | 0.1709 | 0.5256 | **0.0366** | -0.5809 | **0.0183** | -0.4439 | 0.0850 |
| Height | -0.6929 | **0.0029** | -0.3330 | 0.2075 | -0.5574 | **0.0249** | 0.4447 | 0.0843 |
| MCWD | 0.6625 | **0.0052** | -0.0289 | 0.9153 | 0.6961 | **0.0027** | -0.2701 | 0.3117 |
| Temperature | 0.4147 | 0.1102 | -0.5256 | **0.0365** | 0.6514 | **0.0063** | 0.2066 | 0.4426 |
| ET | 0.0011 | 0.9968 | 0.6286 | **0.0091** | -0.2160 | 0.4217 | -0.6415 | **0.0074** |
| ESI | -0.1315 | 0.6273 | 0.6878 | **0.0032** | -0.3947 | 0.1302 | -0.5169 | **0.0403** |
| WUE | -0.3292 | 0.2132 | -0.6283 | **0.0091** | -0.0605 | 0.8239 | 0.5817 | **0.0181** |
| Slope | -0.1413 | 0.6018 | -0.1151 | 0.6713 | -0.1113 | 0.6816 | 0.2500 | 0.3504 |
| Aspect | -0.4384 | 0.0894 | -0.1455 | 0.5908 | -0.3446 | 0.1911 | -0.1630 | 0.5463 |
| CEC | -0.3745 | 0.1529 | 0.2234 | 0.4056 | -0.5069 | **0.0451** | 0.1711 | 0.5264 |
| Clay | 0.1024 | 0.7058 | 0.2600 | 0.3309 | 0.0180 | 0.9472 | -0.3068 | 0.2478 |
| pH$H_2O$ | 0.7377 | **0.0011** | 0.0206 | 0.9395 | 0.7461 | **0.0009** | -0.2613 | 0.3283 |
| Sand | 0.1838 | 0.4957 | 0.2588 | 0.3331 | 0.0173 | 0.9493 | 0.0643 | 0.8129 |
| SOC | -0.6026 | **0.0135** | 0.1360 | 0.6154 | -0.6821 | **0.0036** | 0.0661 | 0.8077 |

**Table 6.** Correlation coefficients (R) and P-values (P) between each input band and CWV of morphological traits. P-values smaller than 0.05 are bold.

| Trait / Band | FW | | DW | | LA | | SLA | | TWD | |
|---|---|---|---|---|---|---|---|---|---|---|
| | R | P | R | P | R | P | R | P | R | P |
| Red | -0.3292 | 0.2131 | -0.3034 | 0.2532 | -0.3447 | 0.1911 | 0.2698 | 0.3122 | 0.4826 | 0.0583 |
| Red Edge | -0.3074 | 0.2468 | -0.3056 | 0.2497 | -0.3381 | 0.2002 | -0.0104 | 0.9694 | -0.0301 | 0.9119 |
| NIR | -0.0959 | 0.7237 | -0.1075 | 0.6918 | -0.1157 | 0.6695 | -0.1000 | 0.7124 | -0.3505 | 0.1833 |
| NDVI | -0.3525 | 0.1806 | -0.3314 | 0.2099 | -0.3569 | 0.1748 | 0.2150 | 0.4238 | 0.3565 | 0.1754 |
| NDRE | -0.2980 | 0.2623 | -0.3020 | 0.2556 | -0.1857 | 0.4912 | 0.1748 | 0.5172 | 0.4211 | 0.1043 |
| SAVI | -0.3877 | 0.1379 | -0.3985 | 0.1263 | -0.3013 | 0.2568 | 0.0871 | 0.7484 | -0.1615 | 0.5501 |
| MSAVI | -0.2285 | 0.3947 | -0.2428 | 0.3649 | -0.2025 | 0.4520 | -0.1440 | 0.5948 | -0.2830 | 0.2882 |
| Height | 0.2008 | 0.4559 | 0.1307 | 0.6296 | 0.4764 | 0.0621 | 0.0228 | 0.9331 | -0.3390 | 0.1990 |
| MCWD | 0.3504 | 0.1833 | 0.3427 | 0.1938 | 0.3047 | 0.2512 | -0.2387 | 0.3732 | -0.6926 | **0.0029** |
| Temperature | -0.4282 | 0.0980 | -0.4214 | 0.1041 | -0.3172 | 0.2313 | 0.2959 | 0.2658 | 0.5391 | **0.0312** |
| ET | -0.1957 | 0.4676 | -0.1721 | 0.5239 | -0.2214 | 0.4099 | 0.0788 | 0.7717 | 0.5022 | **0.0474** |
| ESI | -0.2006 | 0.4563 | -0.1766 | 0.5129 | -0.2286 | 0.3944 | 0.0707 | 0.7947 | 0.5257 | **0.0365** |
| WUE | -0.1894 | 0.4824 | -0.1649 | 0.5418 | -0.2231 | 0.4062 | 0.0654 | 0.8099 | 0.5395 | **0.0310** |
| Slope | 0.6807 | **0.0037** | 0.6765 | **0.0040** | 0.6934 | **0.0029** | 0.0639 | 0.8142 | -0.1108 | 0.6830 |
| Aspect | 0.0685 | 0.8010 | 0.0103 | 0.9699 | 0.1905 | 0.4798 | -0.4993 | **0.0490** | -0.3803 | 0.1462 |
| CEC | -0.1208 | 0.6558 | -0.1063 | 0.6953 | -0.1808 | 0.5029 | -0.1343 | 0.6200 | -0.0488 | 0.8574 |
| Clay | -0.1959 | 0.4671 | -0.1820 | 0.4998 | -0.1999 | 0.4580 | 0.1709 | 0.5269 | 0.3875 | 0.1381 |
| pH$H_2O$ | -0.7449 | **0.0009** | -0.7308 | **0.0013** | -0.6886 | **0.0032** | 0.2643 | 0.3225 | 0.3727 | 0.1551 |
| Sand | 0.3290 | 0.2134 | 0.3371 | 0.2017 | 0.2323 | 0.3867 | -0.2103 | 0.4343 | 0.0438 | 0.8721 |
| SOC | -0.2012 | 0.4549 | -0.1717 | 0.5250 | -0.2322 | 0.3868 | 0.3293 | 0.2130 | 0.5292 | **0.0350** |

**Table 7.** Correlation coefficients (R) and P-values (P) between each input band and CWV of nutrient traits. P-values smaller than 0.05 are bold.

| Band \ Trait | N | | P | | Ca | | Mg | |
|---|---|---|---|---|---|---|---|---|
| | R | P | R | P | R | P | R | P |
| Red | 0.3644 | 0.1653 | 0.3674 | 0.1615 | -0.0443 | 0.8707 | -0.2221 | 0.4084 |
| Red Edge | -0.2691 | 0.3136 | 0.0371 | 0.8916 | -0.0761 | 0.7795 | -0.3064 | 0.2485 |
| NIR | -0.4303 | 0.0962 | -0.1324 | 0.6250 | -0.0909 | 0.7378 | -0.2201 | 0.4127 |
| NDVI | 0.2354 | 0.3801 | 0.3024 | 0.2550 | -0.0536 | 0.8436 | -0.2744 | 0.3037 |
| NDRE | 0.1747 | 0.5176 | 0.1200 | 0.6580 | 0.1303 | 0.6306 | 0.1437 | 0.5954 |
| SAVI | -0.2486 | 0.3532 | 0.0162 | 0.9525 | -0.0963 | 0.7229 | -0.2113 | 0.4320 |
| MSAVI | -0.3978 | 0.1270 | -0.1456 | 0.5904 | -0.1549 | 0.5669 | -0.2925 | 0.2716 |
| Height | 0.0608 | 0.8231 | -0.3000 | 0.2589 | -0.1295 | 0.6328 | 0.3298 | 0.2122 |
| MCWD | -0.2799 | 0.2938 | -0.2126 | 0.4291 | -0.1911 | 0.4784 | -0.1583 | 0.5582 |
| Temperature | 0.4175 | 0.1076 | 0.3353 | 0.2043 | 0.1030 | 0.7041 | 0.1015 | 0.7084 |
| ET | 0.1259 | 0.6423 | 0.0621 | 0.8193 | 0.8439 | **0.0000** | 0.7233 | **0.0015** |
| ESI | 0.1754 | 0.5159 | 0.0914 | 0.7363 | 0.7379 | **0.0011** | 0.6371 | **0.0079** |
| WUE | 0.2782 | 0.2968 | 0.1750 | 0.5169 | 0.4094 | 0.1153 | 0.3377 | 0.2009 |
| Slope | 0.4073 | 0.1174 | -0.0750 | 0.7826 | 0.1915 | 0.4775 | 0.4622 | 0.0715 |
| Aspect | -0.4407 | 0.0875 | -0.6064 | **0.0128** | -0.2250 | 0.4020 | -0.0071 | 0.9792 |
| CEC | -0.1341 | 0.6206 | 0.0497 | 0.8549 | -0.2106 | 0.4338 | -0.2313 | 0.3887 |
| Clay | 0.0064 | 0.9814 | 0.1386 | 0.6086 | 0.2090 | 0.4372 | 0.2332 | 0.3847 |
| $pHH_2O$ | -0.0840 | 0.7571 | 0.3027 | 0.2544 | 0.2576 | 0.3354 | 0.1175 | 0.6648 |
| Sand | -0.0056 | 0.9835 | -0.1583 | 0.5582 | -0.2852 | 0.2843 | -0.2171 | 0.4194 |
| SOC | 0.1916 | 0.4772 | 0.3300 | 0.2120 | 0.2694 | 0.3130 | 0.2401 | 0.3703 |

**Table 8.** Correlation coefficients (R) and P-values (P) between each input band and CWV of hydraulic traits. P-values smaller than 0.05 are bold.

| Band \ Trait | P50 | | P88 | | WPmd | |
|---|---|---|---|---|---|---|
| | R | P | R | P | R | P |
| Red | -0.0793 | 0.7703 | -0.2753 | 0.3021 | 0.2236 | 0.4052 |
| Red Edge | -0.1651 | 0.5411 | -0.3117 | 0.2399 | -0.1081 | 0.6904 |
| NIR | -0.1768 | 0.5125 | -0.1770 | 0.5120 | -0.3184 | 0.2294 |
| NDVI | -0.0609 | 0.8227 | -0.2567 | 0.3373 | -0.0159 | 0.9535 |
| NDRE | 0.3891 | 0.1363 | 0.2197 | 0.4135 | 0.4080 | 0.1167 |
| SAVI | -0.1392 | 0.6073 | -0.1288 | 0.6345 | -0.2664 | 0.3185 |
| MSAVI | -0.1295 | 0.6326 | -0.1696 | 0.5299 | -0.3819 | 0.1444 |
| Height | 0.4406 | 0.0876 | 0.7197 | **0.0017** | -0.3457 | 0.1897 |
| MCWD | -0.3311 | 0.2104 | -0.0368 | 0.8924 | -0.6544 | **0.0060** |
| Temperature | 0.1072 | 0.6929 | 0.0155 | 0.9545 | 0.4004 | 0.1243 |
| ET | 0.4982 | **0.0495** | 0.2202 | 0.4126 | 0.6370 | **0.0080** |
| ESI | 0.4178 | 0.1073 | 0.1604 | 0.5528 | 0.6936 | **0.0029** |
| WUE | 0.1668 | 0.5369 | -0.0207 | 0.9393 | 0.7087 | **0.0021** |
| Slope | 0.2933 | 0.2703 | 0.4998 | **0.0487** | 0.0116 | 0.9659 |
| Aspect | 0.2904 | 0.2752 | 0.2827 | 0.2888 | -0.3907 | 0.1345 |
| CEC | -0.4143 | 0.1106 | -0.3338 | 0.2064 | 0.2279 | 0.3959 |
| Clay | 0.1352 | 0.6176 | 0.0154 | 0.9550 | 0.6494 | **0.0065** |
| $pHH_2O$ | -0.2099 | 0.4353 | -0.3043 | 0.2518 | 0.6153 | **0.0112** |
| Sand | -0.0379 | 0.8891 | -0.1053 | 0.6980 | 0.1368 | 0.6134 |
| SOC | 0.0888 | 0.7437 | -0.0349 | 0.8980 | 0.7639 | **0.0006** |

**Table 9.** Correlation coefficients (R) and P-values (P) between each input band and CWV of photosynthetic traits. P-values smaller than 0.05 are bold.

| Trait / Band | TmaxL | | TspanL | | Topt | | T50 | |
|---|---|---|---|---|---|---|---|---|
| | R | P | R | P | R | P | R | P |
| Red | 0.5512 | **0.0269** | 0.0593 | 0.8274 | -0.0457 | 0.8665 | -0.3521 | 0.1810 |
| Red Edge | -0.0750 | 0.7825 | -0.3219 | 0.2240 | -0.4092 | 0.1155 | -0.4880 | 0.0551 |
| NIR | -0.4112 | 0.1136 | -0.2850 | 0.2847 | -0.3111 | 0.2409 | -0.2601 | 0.3306 |
| NDVI | 0.3398 | 0.1979 | -0.0379 | 0.8891 | -0.0705 | 0.7954 | -0.3251 | 0.2192 |
| NDRE | 0.4873 | 0.0556 | -0.0728 | 0.7888 | -0.0521 | 0.8481 | -0.3800 | 0.1465 |
| SAVI | -0.1390 | 0.6078 | -0.2981 | 0.2621 | -0.1985 | 0.4611 | -0.3251 | 0.2192 |
| MSAVI | -0.3585 | 0.1727 | -0.3514 | 0.1821 | -0.3116 | 0.2401 | -0.3572 | 0.1744 |
| Height | -0.1600 | 0.5540 | 0.1143 | 0.6733 | 0.4190 | 0.1062 | 0.5059 | **0.0456** |
| MCWD | -0.7017 | **0.0024** | 0.0840 | 0.7571 | 0.2104 | 0.4341 | 0.4050 | 0.1196 |
| Temperature | 0.6715 | **0.0044** | 0.0314 | 0.9080 | 0.0555 | 0.8383 | -0.1653 | 0.5407 |
| ET | 0.3393 | 0.1985 | -0.0849 | 0.7545 | -0.1145 | 0.6729 | -0.1358 | 0.6160 |
| ESI | 0.4477 | 0.0820 | -0.0439 | 0.8717 | -0.1015 | 0.7083 | -0.1585 | 0.5577 |
| WUE | 0.6195 | **0.0105** | 0.0531 | 0.8450 | -0.0723 | 0.7901 | -0.2205 | 0.4119 |
| Slope | 0.0235 | 0.9313 | 0.7408 | **0.0010** | 0.7260 | **0.0015** | 0.6033 | **0.0134** |
| Aspect | -0.4932 | 0.0522 | -0.3423 | 0.1943 | -0.2111 | 0.4325 | -0.0286 | 0.9163 |
| CEC | 0.0425 | 0.8757 | -0.0776 | 0.7751 | -0.1951 | 0.4691 | -0.1676 | 0.5349 |
| Clay | 0.2805 | 0.2926 | -0.1222 | 0.6520 | -0.2543 | 0.3418 | -0.2458 | 0.3588 |
| pH$H_2O$ | 0.4609 | 0.0724 | -0.3681 | 0.1607 | -0.4673 | 0.0680 | -0.5186 | **0.0396** |
| Sand | -0.0218 | 0.9362 | 0.1980 | 0.4623 | -0.0242 | 0.9291 | -0.1427 | 0.5981 |
| SOC | 0.4750 | 0.0630 | 0.0138 | 0.9597 | -0.1527 | 0.5724 | -0.2231 | 0.4062 |

**Table 10.** Correlation coefficients (R) and P-values (P) between each input band and the four categories of functional dispersion. P-values smaller than 0.05 are bold.

| Metric / Band | $FDis_{Mor}$ | | $FDis_{Nutr}$ | | $FDis_{Hydr}$ | | $FDis_{Pho}$ | |
|---|---|---|---|---|---|---|---|---|
| | R | P | R | P | R | P | R | P |
| Red | 0.0761 | 0.7793 | 0.0156 | 0.9544 | 0.1175 | 0.6648 | 0.0572 | 0.8334 |
| Red Edge | -0.0515 | 0.8498 | -0.1540 | 0.5690 | -0.2337 | 0.3837 | -0.1368 | 0.6135 |
| NIR | 0.0296 | 0.9135 | -0.1302 | 0.6307 | -0.3696 | 0.1588 | -0.0106 | 0.9689 |
| NDVI | -0.0397 | 0.8839 | -0.1560 | 0.5640 | -0.5372 | **0.0319** | 0.0029 | 0.9914 |
| NDRE | 0.1901 | 0.4806 | 0.0132 | 0.9614 | -0.2413 | 0.3679 | 0.2453 | 0.3598 |
| SAVI | -0.2929 | 0.2709 | -0.1557 | 0.5648 | -0.5668 | **0.0221** | -0.2785 | 0.2963 |
| MSAVI | -0.1073 | 0.6924 | -0.2462 | 0.3581 | -0.7036 | **0.0024** | -0.1158 | 0.6693 |
| Height | 0.2855 | 0.2838 | 0.0703 | 0.7958 | -0.3068 | 0.2478 | 0.4670 | 0.0682 |
| MCWD | -0.4170 | 0.1080 | 0.0183 | 0.9463 | 0.2451 | 0.3601 | -0.3234 | 0.2217 |
| Temperature | 0.1566 | 0.5624 | 0.5198 | **0.0390** | 0.7032 | **0.0024** | 0.3290 | 0.2134 |
| ET | -0.2905 | 0.2750 | -0.2062 | 0.4436 | -0.5666 | **0.0221** | -0.3322 | 0.2088 |
| ESI | -0.1165 | 0.6673 | -0.2892 | 0.2773 | -0.6221 | **0.0101** | -0.2652 | 0.3208 |
| WUE | 0.4202 | 0.1051 | 0.4373 | 0.0903 | 0.3785 | 0.1483 | 0.6140 | **0.0114** |
| Slope | 0.6683 | **0.0047** | 0.6425 | **0.0073** | 0.2178 | 0.4178 | 0.7128 | **0.0019** |
| Aspect | -0.1893 | 0.4825 | -0.2403 | 0.3701 | -0.1581 | 0.5586 | -0.0663 | 0.8072 |
| CEC | 0.3680 | 0.1608 | -0.0838 | 0.7576 | -0.1683 | 0.5332 | 0.1315 | 0.6273 |
| Clay | -0.4384 | 0.0894 | -0.3750 | 0.1524 | -0.1830 | 0.4975 | -0.5153 | **0.0411** |
| $pHH_2O$ | -0.4333 | 0.0936 | 0.0233 | 0.9318 | 0.2443 | 0.3618 | -0.3659 | 0.1634 |
| Sand | 0.0766 | 0.7781 | -0.1567 | 0.5622 | -0.0696 | 0.7980 | -0.1358 | 0.6160 |
| SOC | -0.0367 | 0.1262 | -0.1908 | 0.4792 | -0.1473 | 0.5862 | 0.1755 | 0.5155 |

**Table 11.** Correlation coefficients (R) and P-values (P) between each input band and the four categories of functional redundancy. P-values smaller than 0.05 are bold.

| Metric / Band | $FRed_{Mor}$ | | $FRed_{Nutr}$ | | $FRed_{Hydr}$ | | $FRed_{Pho}$ | |
|---|---|---|---|---|---|---|---|---|
| | R | P | R | P | R | P | R | P |
| Red | 0.0446 | 0.8696 | 0.0649 | 0.8113 | 0.0288 | 0.9158 | 0.0517 | 0.8492 |
| Red Edge | -0.1957 | 0.4677 | -0.1737 | 0.5201 | -0.1695 | 0.5302 | -0.1773 | 0.5113 |
| NIR | -0.0990 | 0.7152 | -0.0729 | 0.7885 | -0.0550 | 0.8397 | -0.0892 | 0.7426 |
| NDVI | -0.1248 | 0.6453 | -0.1134 | 0.6759 | -0.0524 | 0.8470 | -0.1329 | 0.6238 |
| NDRE | 0.1852 | 0.4923 | 0.2067 | 0.4425 | 0.2154 | 0.4231 | 0.1748 | 0.5173 |
| SAVI | -0.2661 | 0.3192 | -0.3044 | 0.2517 | -0.2174 | 0.4186 | -0.2744 | 0.3036 |
| MSAVI | -0.2943 | 0.2685 | -0.2757 | 0.3013 | -0.1910 | 0.4786 | -0.2915 | 0.2734 |
| Height | 0.2492 | 0.3519 | 0.2929 | 0.2709 | 0.3086 | 0.2449 | 0.2258 | 0.4004 |
| MCWD | -0.1579 | 0.5593 | -0.2400 | 0.3706 | -0.2318 | 0.3877 | -0.1832 | 0.4971 |
| Temperature | 0.4864 | 0.0561 | 0.4295 | 0.0969 | 0.4009 | 0.1238 | 0.4613 | 0.0721 |
| ET | -0.3746 | 0.1528 | -0.3935 | 0.1315 | -0.3119 | 0.2395 | -0.3693 | 0.1592 |
| ESI | -0.3974 | 0.1275 | -0.3677 | 0.1611 | -0.3097 | 0.2430 | -0.3676 | 0.1613 |
| WUE | 0.5575 | **0.0248** | 0.5720 | **0.0206** | 0.5462 | **0.0286** | 0.5384 | **0.0314** |
| Slope | 0.7172 | **0.0018** | 0.7328 | **0.0012** | 0.7390 | **0.0011** | 0.7306 | **0.0013** |
| Aspect | -0.2505 | 0.3495 | -0.2269 | 0.3982 | -0.2272 | 0.3974 | -0.2724 | 0.3074 |
| CEC | 0.0750 | 0.7825 | 0.1510 | 0.5768 | 0.1286 | 0.6351 | 0.1198 | 0.6585 |
| Clay | -0.4483 | 0.0816 | -0.4697 | 0.0664 | -0.4530 | 0.0780 | -0.4498 | 0.0804 |
| $pHH_2O$ | -0.1572 | 0.5610 | -0.2465 | 0.3573 | -0.2364 | 0.3780 | -0.1796 | 0.5056 |
| Sand | -0.1152 | 0.6709 | -0.0831 | 0.7597 | -0.1145 | 0.6729 | -0.0799 | 0.7686 |
| SOC | 0.1683 | 0.9506 | 0.0926 | 0.7329 | 0.0548 | 0.8402 | 0.0288 | 0.9156 |

**Table 12.** Model performance for mapping the two community-weighted moments of morphological traits. The bold numbers indicate the highest $R^2$ values.

| Trait | FW | | DW | | LA | | SLA | | TWD | |
|---|---|---|---|---|---|---|---|---|---|---|
| Moment | CWM | CWV | CWM | CWV | CWM | CWV | CWM | CWV | CWM | CWV |
| $R^2$ | 0.50 | **0.64** | 0.58 | 0.59 | 0.33 | 0.51 | 0.82 | 0.40 | **0.92** | 0.37 |
| RMSE | 0.18 | 0.19 | 0.07 | 0.03 | 4.04 | 57.25 | 18.71 | 565.68 | 0.05 | 0.00 |
| MAE | 0.14 | 0.14 | 0.05 | 0.02 | 3.14 | 47.72 | 15.48 | 453.34 | 0.04 | 0.00 |

**Table 13.** Model performance for mapping the two community-weighted moments of nutrient traits. The bold numbers indicate the highest $R^2$ values.

| Trait | N | | P | | Ca | | Mg | |
|---|---|---|---|---|---|---|---|---|
| Moment | CWM | CWV | CWM | CWV | CWM | CWV | CWM | CWV |
| $R^2$ | 0.75 | 0.27 | **0.77** | 0.39 | 0.36 | **0.44** | 0.25 | 0.15 |
| RMSE | 0.28 | 0.05 | 0.05 | 0.00 | 0.23 | 0.15 | 0.07 | 0.01 |
| MAE | 0.25 | 0.04 | 0.03 | 0.00 | 0.17 | 0.09 | 0.06 | 0.01 |

**Table 14.** Model performance for mapping the two community-weighted moments of hydraulic traits. The bold numbers indicate the highest $R^2$ values.

| Trait | P50 | | P88 | | WPmd | |
|---|---|---|---|---|---|---|
| Moment | CWM | CWV | CWM | CWV | CWM | CWV |
| $R^2$ | 0.54 | 0.27 | 0.77 | 0.14 | **0.97** | **0.96** |
| RMSE | 0.36 | 0.30 | 0.49 | 0.53 | 0.20 | 0.39 |
| MAE | 0.29 | 0.25 | 0.42 | 0.46 | 0.16 | 0.21 |

**Table 15.** Model performance for mapping the two community-weighted moments of photosynthetic traits. The bold numbers indicate the highest $R^2$ values.

| Trait | TmaxL | | Topt | | TspanL | | T50 | |
|---|---|---|---|---|---|---|---|---|
| Moment | CWM | CWV | CWM | CWV | CWM | CWV | CWM | CWV |
| $R^2$ | 0.57 | 0.54 | **0.68** | **0.77** | 0.34 | 0.18 | 0.57 | 0.38 |
| RMSE | 2.08 | 5.49 | 1.83 | 2.59 | 1.04 | 2.49 | 1.26 | 1.88 |
| MAE | 1.41 | 3.69 | 1.47 | 1.66 | 0.76 | 1.60 | 0.91 | 1.42 |

**Table 16.** Model performance for assessing the four groups of FDis and FRed. The bold numbers indicate the highest $R^2$ values.

| Group | Morphology | | Nutrients | | Hydraulic | | Photosynthesis | |
|---|---|---|---|---|---|---|---|---|
| Metric | FDis | FRed | FDis | FRed | FDis | FRed | FDis | FRed |
| $R^2$ | 0.36 | 0.52 | 0.23 | 0.48 | 0.69 | 0.54 | 0.64 | 0.56 |
| RMSE | 0.91 | 0.16 | 0.55 | 0.17 | 0.41 | 0.18 | 0.44 | 0.16 |
| MAE | 0.69 | 0.12 | 0.45 | 0.13 | 0.32 | 0.15 | 0.33 | 0.13 |

**Table 17.** Description of all morphological traits measured in Chile and the reasons why they were measured in this study.

| Trait | Abbreviation | Unit | Description | Trait selection justification | Reference |
|---|---|---|---|---|---|
| Leaf fresh weight | FW | g | Mass of a fresh leaf | Higher FW suggests robust growth and favourable environmental conditions, while lower values may indicate stress or resource limitations | (Mokhtarpour et al., 2010) |
| Leaf dry weight | DW | g | Mass of a dry leaf | DW assesses the plant's biomass allocation and long-term growth strategies and resource-use efficiency | (Niklas et al., 2007) |
| Specific leaf area | SLA | $cm^2 \cdot g^{-1}$ | A ratio indicating how much leaf area a plant builds with a given amount of leaf biomass | SLA correlated with whole plant growth and reflects the trade-off between resource acquisition and conservation | (Wilson et al., 1999) |
| Leaf area | LA | $cm^2$ | Area of leaves | LA quantifies the surface area available for photosynthesis, providing a direct measure of the plant's potential to capture solar energy and contribute to ecosystem productivity | (Bhagsari and Brown, 1986) |
| Leaf mass per area | LMA | $g \cdot cm^{-2}$ | The ratio between leaf dry mass and leaf area | Higher LMA suggests thicker, denser leaves with potentially lower photosynthetic rates, and lower LMA indicates thinner, more photosynthetically active leaves | (Hassiotou et al., 2010) |
| Leaf dry matter content | LDMC | $mg \cdot g^{-1}$ | The ratio of leaf dry mass to fresh mass | LDMC evaluates the proportion of dry matter in the leaf, providing insights into leaf structure, resource-use strategies, and the plant's its ability to retain water | (Vendramini et al., 2002) |
| Trunk wood density | TWD | $g \cdot cm^{-3}$ | The dry weight per unit volume of wood, the amount of wood in a unit measured at trunks | TWD assesses the structural and mechanical properties of tree trunks, wood strength, resource allocation strategies, and potential resistance to mechanical stresses and environmental pressures | (Larjavaara and Muller-Landau, 2010) |
| Branch wood density | BWD | $g \cdot cm^{-3}$ | The dry weight per unit volume of wood, the amount of wood in a unit measured at branches | BWD evaluates the structural characteristics of tree branches, branch strength, mechanical stability, and resource allocation within the canopy, influencing overall tree architecture and ecological interactions | (Meinzer et al., 2008) |

**Table 18.** Description of all nutrient traits measured in Chile and the reasons why they were measured in this study.

| Trait | Abbreviation | Unit | Description | Trait selection justification | Reference |
|---|---|---|---|---|---|
| Leaf calcium content | Ca | % | Calcium content per unit dry leaf mass | Leaf nutrients comprehensively assess plant nutritional status, nutrient interactions, and potential impacts ecosystem dynamics | (Furey and Tilman, 2023) |
| Leaf potassium content | K | % | Potassium content per unit dry leaf mass | | |
| Leaf magnesium content | Mg | % | Magnesium content per unit dry leaf mass | | |
| Leaf nitrogen content | N | % | Nitrogen content per unit dry leaf mass | | |
| Leaf phosphorus content | P | % | Phosphorus content per unit dry leaf mass | | |
| Ratio of leaf nitrogen and phosphorus content | N/P | Unitless | Ratio of leaf nitrogen and phosphorus content per unit dry leaf mass | | |

**Table 19.** Description of all hydraulic traits measured in Chile and the reasons why they were measured in this study.

| Trait | Abbreviation | Unit | Description | Trait selection justification | Reference |
|---|---|---|---|---|---|
| P50 | P50 | MPa | Water potential at which 50% and 88% of hydraulic conductivity is lost | P50 and P88 evaluate the plant's tolerance to water stress and its ability to maintain hydraulic conductivity under drought conditions | (Brum *et al*., 2023) |
| P88 | P88 | MPa | | | |
| WPmd | WPmd | MPa | Minimum water potential (midday water potential at the driest month) | WPmd assesses the plant's drought tolerance and capacity to withstand water stress, providing critical information on its ability to maintain water balance during periods of limited water availability | (Markesteijn *et al*., 2010) |
| SM_P50 | SM50 | MPa | Safety Margin P50 and P88 | SM50 and SM88 serve as important indicators predicting the vulnerability of plants to drought-induced mortality | (Tavares *et al*., 2023) |
| SM_P88 | SM88 | MPa | | | |

**Table 20.** Description of all photosynthetic traits measured in Chile and the reasons why they were measured in this study.

| Trait | Abbreviation | Unit | Description | Trait selection justification | Reference |
|---|---|---|---|---|---|
| TmaxL | TmaxL | °C | Temperature at carbon compensation point | TmaxL assesses the minimum temperature required for a plant to achieve carbon balance, providing insights into a species' thermal tolerance, metabolic performance, and adaptation to specific environmental conditions | (Walker and Cousins, 2013) |
| Temperature of Optimum Photosynthesis | Topt | °C | Temperature of Optimum Photosynthesis | Topt evaluates the temperature range at which a plant achieves maximal photosynthetic efficiency, species' thermal adaptation, growth potential, and responsiveness to changing environmental conditions | (Sage and Kubien, 2007) |
| Photosynthesis rate at optimum temperature | Aopt | $\mu mol \cdot CO_2 \cdot m^{-2} \cdot s^{-1}$ | Photosynthesis rate at optimum temperature | Aopt assesses the species' potential for carbon assimilation and overall growth performance | (Sage and Kubien, 2007) |
| TspanL | TspanL | °C | Breadth of temperature optimum | TspanL characterised the range of temperatures over which a plant exhibits optimal photosynthetic rates and the species' ability to perform efficiently across varying environmental conditions | (Rohr *et al*., 2018) |
| T50 | T50 | °C | Temperature at which the maximum quantum yield of the photosystem II declines to 50% | T50 assesses the species' vulnerability to temperature-induced reductions in photosynthetic efficiency | (Perez and Feeley, 2020) |

**Table 21.** Description of multispectral images collected for each plot. Note: In the below table, B, G, R, RE, and NIR are abbreviations for the spectral bands blue, green, red, red edge, and near-infrared, respectively.

| Plot | Instrument type | Imaging date | Scenes tiles | Spectral bands (centre bandwidth (nm)) | Spatial resolution (m) |
|---|---|---|---|---|---|
| CAB1 | MicaSense Altum-PT | 10/01/2020 | 618 | B (475), G (560), R (668), RE (717), NIR (842) | $5.28\times10^{-2}$ |
| CAB2 | MicaSense Altum-PT | 10/01/2020 | 1,200 | B (475), G (560), R (668), RE (717), NIR (842) | $5.28\times10^{-2}$ |
| CAB3 | MicaSense Altum-PT | 10/01/2020 | 799 | B (475), G (560), R (668), RE (717), NIR (842) | $5.28\times10^{-2}$ |
| RAD3 | MicaSense Altum-PT | 11/01/2020 | 1,194 | B (475), G (560), R (668), RE (717), NIR (842) | $5.28\times10^{-2}$ |
| RAD2 | MicaSense Altum-PT | 11/01/2020 | 1,356 | B (475), G (560), R (668), RE (717), NIR (842) | $5.28\times10^{-2}$ |
| RAD1 | MicaSense Altum-PT | 11/01/2020 | 1,188 | B (475), G (560), R (668), RE (717), NIR (842) | $5.28\times10^{-2}$ |
| SPT1 | SuperDove | 09/11/2021 | 2 | Coastal blue (431-452), B (465-515), Green I (513-549), G (547-583), Yellow (600-620), R (650-680), RE (697-713), NIR (845-885) | 3 |
| SPT3 | MicaSense Altum-PT | 03/03/2020 | 390 | B (475), G (560), R (668), RE (717), NIR (842) | $5.28\times10^{-2}$ |
| ALE2 | SuperDove | 18/09/2020 | 2 | B (465-515), G (547-583), R (650-680), RE (697-713), NIR (845-885) | 3 |
| ALE3 | MicaSense Altum-PT | 04/03/2020 | 468 | B (475), G (560), R (668), RE (717), NIR (842) | $5.28\times10^{-2}$ |
| COR1 | SuperDove | 24/03/2021 | 1 | B (465-515), G (547-583), R (650-680), RE (697-713), NIR (845-885) | 3 |
| COR3 | SuperDove | 24/03/2021 | 1 | B (465-515), G (547-583), R (650-680), RE (697-713), NIR (845-885) | 3 |
| TRA1 | MicaSense Altum-PT | 20/01/2020 | 498 | B (475), G (560), R (668), RE (717), NIR (842) | $5.28\times10^{-2}$ |
| TRA2 | MicaSense Altum-PT | 20/01/2020 | 732 | B (475), G (560), R (668), RE (717), NIR (842) | $5.28\times10^{-2}$ |
| MAG1 | SuperDove | 17/01/2023 | 1 | Coastal blue (431-452), B (465-515), Green I (513-549), G (547-583), Yellow (600-620), R (650-680), RE (697-713), NIR (845-885) | 3 |
| MAG2 | MicaSense Altum-PT | 07/03/2020 | 780 | B (475), G (560), R (668), RE (717), NIR (842) | $5.28\times10^{-2}$ |

**Table 22.** Description of LiDAR data collected for each plot.

| Plot | Instrument type | Imaging date | Scanner points per second | Spatial resolution (m) |
|---|---|---|---|---|
| CAB1 | ZEB1 handheld 3D scanner | 06/01/2020 | 43,000 | 0.01-0.03 (environment dependant) |
| CAB2 | ZEB1 handheld 3D scanner | 06/01/2020 | 43,000 | 0.01-0.03 (environment dependant) |
| CAB3 | ZEB1 handheld 3D scanner | 06/01/2020 | 43,000 | 0.01-0.03 (environment dependant) |
| RAD3 | ZEB1 handheld 3D scanner | 17/01/2020 | 43,000 | 0.01-0.03 (environment dependant) |
| RAD2 | ZEB1 handheld 3D scanner | 17/01/2020 | 43,000 | 0.01-0.03 (environment dependant) |
| RAD1 | ZEB1 handheld 3D scanner | 17/01/2020 | 43,000 | 0.01-0.03 (environment dependant) |
| SPT1 | ZEB1 handheld 3D scanner | 28/02/2020 | 43,000 | 0.01-0.03 (environment dependant) |
| SPT3 | ZEB1 handheld 3D scanner | 28/02/2020 | 43,000 | 0.01-0.03 (environment dependant) |
| ALE2 | ZEB1 handheld 3D scanner | 29/02/2020 | 43,000 | 0.01-0.03 (environment dependant) |
| ALE3 | ZEB1 handheld 3D scanner | 29/02/2020 | 43,000 | 0.01-0.03 (environment dependant) |
| COR1 | Global Forest Canopy Height | / | / | 30 |
| COR3 | Global Forest Canopy Height | / | / | 30 |
| TRA1 | ZEB1 handheld 3D scanner | 15/01/2020 | 43,000 | 0.01-0.03 (environment dependant) |
| TRA2 | ZEB1 handheld 3D scanner | 15/01/2020 | 43,000 | 0.01-0.03 (environment dependant) |
| MAG1 | ZEB1 handheld 3D scanner | 02/03/2020 | 43,000 | 0.01-0.03 (environment dependant) |
| MAG2 | ZEB1 handheld 3D scanner | 03/03/2020 | 43,000 | 0.01-0.03 (environment dependant) |

**Table 23.** Vegetation indices generated from spectral bands.

| Vegetation indices | Abbreviation | Equation | Description | Reference |
|---|---|---|---|---|
| Normalised Difference Vegetation Index | NDVI | $\frac{\rho_{NIR} - \rho_{Red}}{\rho_{NIR} + \rho_{Red}}$ | Used to estimate the amount and health of vegetation | (Rouse *et al*., 1974) |
| Normalised Difference Red Edge Index | NDRE | $\frac{\rho_{NIR} - \rho_{Red\ Edge}}{\rho_{NIR} + \rho_{Red\ Edge}}$ | Particularly useful for assessing subtle changes in vegetation health and stress | (Fitzgerald *et al*., 2010) |
| Soil-Adjusted Vegetation Index | SAVI | $1.5\times\frac{\rho_{NIR} - \rho_{Red}}{\rho_{NIR} + \rho_{Red} + 0.5}$ | An improvement over NDVI, designed to minimise the influence of soil brightness | (Huete, 1988) |
| Modified Soil-Adjusted Vegetation Index | MSAVI | $\frac{2\times\rho_{NIR} + 1 - \sqrt{(2\times\rho_{NIR} + 1)^2 - 8\times(\rho_{NIR} - \rho_{Red})}}{2}$ | Aiming to provide a more accurate representation of vegetation cover | (Qi *et al*., 1994) |
